# Solid Phase Synthesis of Fluorosulfate Containing Macrocycles for Chemoproteomic Workflows

**DOI:** 10.1101/2023.02.17.529022

**Authors:** Franco F. Faucher, Daniel Abegg, Phillip Ipock, Alexander Adibekian, Scott Lovell, Matthew Bogyo

**Affiliations:** Department of Chemistry, Stanford University; Department of Chemistry, University of Illinois Chicago, Chicago, Illinois 60607, USA; Department of Life Sciences, University of Bath, Bath BA2 7AX, U.K.; Department of Pathology, Stanford University School of Medicine, Stanford, CA 94305; Department of Chemical and Systems Biology, Stanford University School of Medicine, Stanford, CA 94305; Department of Microbiology and Immunology, Stanford University School of Medicine, Stanford, CA 94305

**Keywords:** Fluorosulfate, macrocycles, chemoproteomics, SuFEx, electrophile

## Abstract

Macrocyclic peptides are attractive for chemoproteomic applications due to their modular synthesis and potential for high target selectivity. We describe a solid phase synthesis method for the efficient generation of libraries of small macrocycles that contain an electrophile and alkyne handle. The modular synthesis produces libraries that can be directly screened using simple SDS-PAGE readouts and then optimal lead molecules applied to proteomic analysis. We generated a library of 480 macrocyclic peptides containing the weakly reactive fluorosulfate (OSF) electrophile. Initial screening of a subset of the library containing each of the various diversity elements identified initial molecules of interest. The corresponding positional and confirmational isomers were then screened to select molecules that showed specific protein labeling patterns that were dependent on the probe structure. The most promising hits were applied to standard chemoproteomic workflows to identify protein targets. Our results demonstrate the feasibility of rapid, on-resin synthesis of diverse macrocyclic electrophiles to generate new classes of covalent ligands.

## Introduction

Macrocycles are an important class of drugs that have found success in the clinic due to their high specificity, affinity, and stability.^[1]^ Although there are many examples of macrocyclic drugs, most of these are derived from natural product scaffolds.^[1]^ However, recent advances in diversity-oriented synthesis have enabled the synthesis and screening of large libraries of macrocycles to discover new bioactive molecules.^[2]^ In particular, combinatorial libraries made from high-throughput step-wise syntheses allow the generation of large libraries that can be screened directly from crude mixtures.^[3–6]^ While many libraries include compounds with electrophilic functionalities that can covalently target reactive amino acids, macrocyclic libraries have yet to include such fragments as many electrophiles are unstable to solid phase peptide synthesis (SPPS) or synthetic conditions used to close the macrocycle.^[2,3]^ Thus, there remains a need to develop synthesis methods that allow rapid and efficient incorporation of reactive electrophiles into libraries of complex molecules such as macrocycles to enable the synthesis of libraries with the potential to yield new covalent drug leads.

Covalent drugs have a number of significant benefits over traditional reversible binding drugs. This includes decreased propensity of resistance, increased efficacy, and shortened dosing intervals as a result of their permanent target engagement.^[7]^ In addition to traditional inhibition assays, covalent drug discovery platforms often leverage chemoproteomics to characterize the selectivity of lead molecules and identify off targets.^[8,9]^ Both ligand-first (electrophile is attached to a known reversible binder) and electrophile-first (ligand is built around a covalent fragment as a starting point) approaches have proven successful in identifying inhibitors for disease relevant targets.^[8,10]^ The value of these approaches for drug development has recently been demonstrated with the FDA approval of adagrasib, which forms an irreversible covalent bond with the mutated cysteine of KRAS (G12-C) in metastatic non-small cell lung cancer.^[11]^ Electrophiles which target cysteine, such as acrylamides and halomethylketones, are commonly used in many fragment libraries. However, these reactive groups are also often used for cyclizing macrocycles due to their compatibility with free cysteine nucleophiles.^[2,12,13]^ While there are many synthetic methods for cyclization of compounds, the use of electrophile-mediated linkages with cysteines are usually the most robust.^[3,14]^ Thus, it is important to identify electrophiles that can be used in macrocycle scaffolds without interfering with cysteine mediated ring closing reactions. Recently, a number of electrophiles that target residues other than cysteine such as lysine, histidine, and tyrosine have been described and are ideal candidates for use in the construction of macrocycle libraries due to their overall low reactivity.^[15–18]^

Sulfonyl fluoride exchange (SuFEx) electrophiles are a biologically important class of fragments due to their context dependent reactivity with non-catalytic amino acid residues such as lysine, histidine, and tyrosine.^[17,19]^ This latent reactivity, which is dependent on reversible target binding, has been leveraged in “inverse drug discovery approaches”.^[20,21]^ In this process, low complexity ligands containing SuFEx electrophiles and an alkyne handle are screened in lysates to identify ligand-able proteins through a combination of SDS-PAGE and chemoproteomics. The current landscape of synthetically tractable SuFEx electrophiles covers a wide range of intrinsic reactivity and stability, thus enabling the selection of molecules compatible with chemistries such as those used for SPPS.^[17,22]^ In particular, the fluorosulfate (OSF) electrophile has recently been shown to be an effective covalent electrophile that can be used to selectively label protein targets *in vivo* due to its high aqueous stability, low off target reactivity, and synthetic accessibility.^[17,21,23]^ Therefore, the OSF electrophile is a prime candidate for incorporation into macrocycle libraries that are compatible with chemoproteomic workflows.

Here we present a fully solid-phase synthesis strategy for the rapid and efficient generation of alkyne labeled covalent OSF macrocycles compatible with proteomic workflows. Using two benchmark alkyne probes, we compared the intrinsic reactivity/selectivity of the OSF electrophile to the more commonly used sulfonyl fluoride (SF) to assess the overall reactivity of these two electrophiles in complex biological samples. We ultimately selected the OSF electrophile due to reduced general reactivity and developed a synthesis for a fluorenylmethyloxycarbonyl (Fmoc) protected amino acid containing the OSF electrophile that avoids the use of toxic sulfuryl fluoride gas.^[24]^ This modified amino acid enabled high throughput SPPS synthesis of OSF macrocycles containing alkyne handles based on previously described libraries from the Heinis group.^[3–6]^ Using this synthesis method, we were able to generate a library of 480 macrocycles using a series of 5 linkers, 24 amino acids, 2 cysteine isomers and alternate positioning of the OSF containing amino acids. Because all of the resulting molecules contained an alkyne tag suitable for use in click chemistry, we were able to rapidly screen the library in crude cellular lysates using simple SDS-PAGE readouts. This screening resulted in a small number of compounds that showed specific labeling of protein targets that could be identified by direct chemoproteomic analysis using the selected library hits. These results demonstrate the feasibility of generating libraries of electrophile containing macrocycles using solid phase synthesis methods and further confirm that the OSF electrophile has overall low reactivity in complex proteomes making it an ideal starting point for the design of next generation covalent drugs.

## Results and Discussion

In order to identify an electrophile compatible with both SPPS and cysteine-based cyclizations, we focused on SuFEx electrophiles as their latency provides greater chance for compatibility with other diverse chemistries used in peptide synthesis. We first assessed the reactivity of two commonly used SuFEx electrophiles in complex proteomic samples. To mimic the fragment that would be contained in the final macrocycles, we synthesized probes that contain the SF (**Figure 1A**) and OSF (**Figure 1B**) electrophiles attached to a phenyl ring with an alkyne handle for click chemistry. We then used each probe to label human embryonic kidney 293 (HEK 293) lysates at a high probe concentration (100 μM) for two hours at 37°C followed by copper-catalyzed azide–alkyne cycloaddition (CuAAC) click reaction with desthiobiotin. After affinity purification and trypsinization, we performed LC-MS/MS analysis to identify protein adducts. Overall, we identified 4,504 unique adducts for the SF probe and 181 unique adducts for OSF probe (**Tables S1 and S2**), confirming that the OSF electrophile is intrinsically much less reactive than the SF electrophile. Interestingly, we found that the two probes showed a different pattern of reactivity towards amino acids in proteins (**Figure 1C**). As previously reported, the SF electrophile modified both tyrosine and lysine residues with a slight preference for tyrosine (∼60%) while the OSF had minimal lysine reactivity and mostly targeted histidine and tyrosine (∼50% each).^[18]^ These findings were surprising given that OSF has been shown to react with lysine residues when targeted with a ligand directing group.^[21,25]^ This general lack of lysine modification by our simple OSF fragment probe may be due to the relative small size of the fragment and limited labeling time (2 hours).

**Figure 1.**
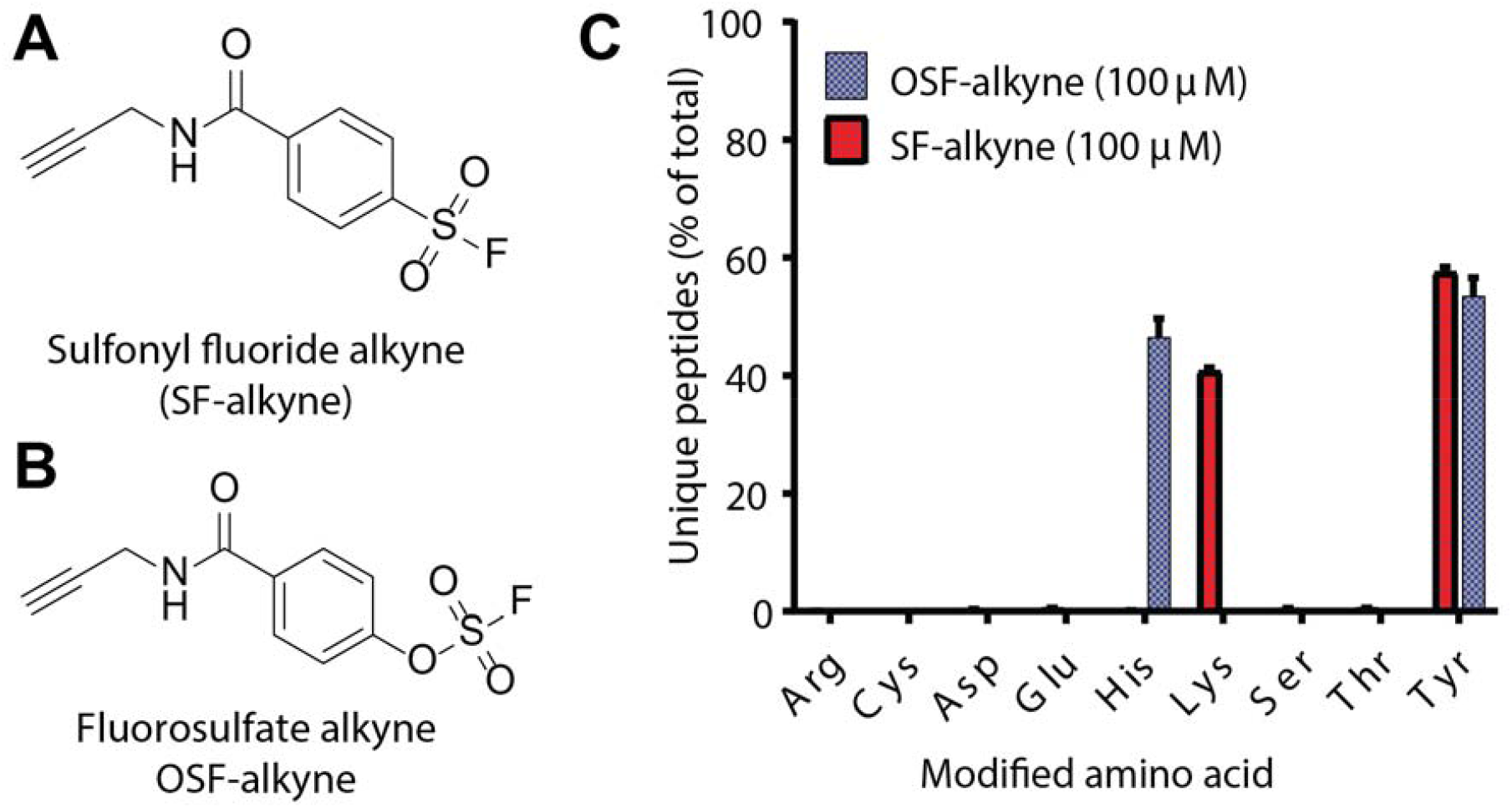
Chemoproteomics with two common SuFEx electrophiles. (**A**) Structure of sulfonyl fluoride alkyne probe (SF-alkyne). (**B**) Structure of fluorosulfate alkyne probe (OSF-alkyne). (**C**) Plot of percent of unique peptides versus amino acid modified for both SF-alkyne and OSF-alkyne. HEK lysate (5 mg/mL, 2.5 mL) was treated with 100 μM OSF or 100 μM SF for two hours at 37 °C followed by site specific chemoproteomic analysis.

Given the results from our chemoproteomics, we decided to use the OSF electrophile in our macrocycle library synthesis. We had to first synthesize the Fmoc-protected unnatural amino acid building block containing the OSF electrophile. We were able to synthesize Fmoc-fluorosulfated tyrosine (Fmoc-Tyrosine(OSF)-OH) using the shelf-stable, crystalline reagent [4-(acetylamino)phenyl]imidodisulfuryl difluoride (AISF) which can be purchased commercially or made in bulk using a one-step, chromatography-free protocol.^[24]^ We initially attempted direct reaction of AISF with the phenol of (tert-butoxycarbonyl)-L-tyrosine as Fmoc protected tyrosine could not be used due to the 1,8-Diazabicyclo[5.4.0]undec-7-ene (DBU) base needed for AISF reactions.^[24]^ However, the presence of the free carboxylic acid led to unwanted side products and low yield. We therefore used, tert-butyl (tert-butoxycarbonyl)-L-tyrosinate to produce the fluorosulfate Intermediate I (**Scheme 1**). To generate the final Fmoc-Tyrosine(OSF)-OH we removed protecting groups from the N and C terminus followed by Fmoc reprotection of the N terminus (**Scheme 1**).

When designing a macrocyclic library that uses cysteine mediated macrocyclization, the choice of cysteine protecting group can dramatically impact the purity of the library. Standard protecting groups, such as tert-butylthio (StBu), are commercially available and stable to SPPS conditions but are sluggish to remove with mild reducing agents.^[26]^ This can lead to incomplete deprotection while also leaving behind thiols or phosphines which may react with the electrophilic linker or covalent fragment. Thus, we chose to use the Trimethoxyphenylthio (S-Tmp) protecting group, which can be removed rapidly and efficiently using 0.1 M N-methylmorpholine (NMM) and dithiothreitol (DTT) (5%) in DMF (3X 5 minutes treatments).^[26]^ This protecting group is ideal as it can be removed on resin during SPPS and the deprotection mixture can be washed away. We synthesized both L and D versions of this amino acid in bulk using reported procedures.^[26]^

Taking inspiration from the macrocyclic libraries reported by the Heinis lab, we designed a synthetic method for the generation of macrocycles which contain a OSF and alkyne handle.^[3–6]^ We designed our libraries to generate diversity by including the Tyr-OSF amino acid and second variable amino acid placed between the cysteine and amino terminus used for cyclization. Further diversity could be created by switching the position of the Tyr-OSF relative to the variable amino acid, by using either the D or L isomer of cysteine and by using a range of different cyclization linkers. This resulted in an initial set of 5 sub-libraries (one for each linker) made up of 96 macrocycles each built from 24 variable amino acids, two cysteine residues (D and L isomers), and 2 positions of the variable amino acids relative to the OSF amino acid. To generate this library, we coupled the Fmoc-(D/L)Cys(S-TMP)-OH to rink amide resin followed by deprotection and coupling of Fmoc-Tyrosine(OSF)-OH and a variable amino acid using standard SPPS conditions (**Figure 2A**). The position of the Fmoc-Tyrosine(OSF)-OH and variable amino acid was also swapped (**Figure 2B**). We then coupled a bromoacetate to the peptide free amino terminus, which allows addition of primary amines to produce capped peptides with a secondary amine that could be used to cyclize the molecule (**Figure 2A**). It also allowed the incorporation of an additional diversity position on the main macrocyclic scaffold. For this library, we used propargyl alkyne which resulted in final macrocycles that contained a tag for labeling and chemoproteomic analysis. To generate untagged compounds for competition labeling studies we replaced the propargyl amine with propyl amine. As a final step, we removed the S-Tmp protecting group followed by multiple wash steps to remove excess NMM/DTT and finally cyclized the molecules using multiple different bis-electrophilic linkers (**Figure 2A**). We had to optimize the reactions conditions for each of the bis-electrophilic linkers (**Figure 2D)** as initial testing showed no single optimal condition (see Chemistry Methods in **Supporting Information** for details). Using this approach and the components outlined in **Figure 2C**, we synthesized 96 macrocycles for each of the 5 main linkers in **Figure 2D** using 24 overall variable amino acids which included 6 unnatural amino acids.

**Figure 2.**
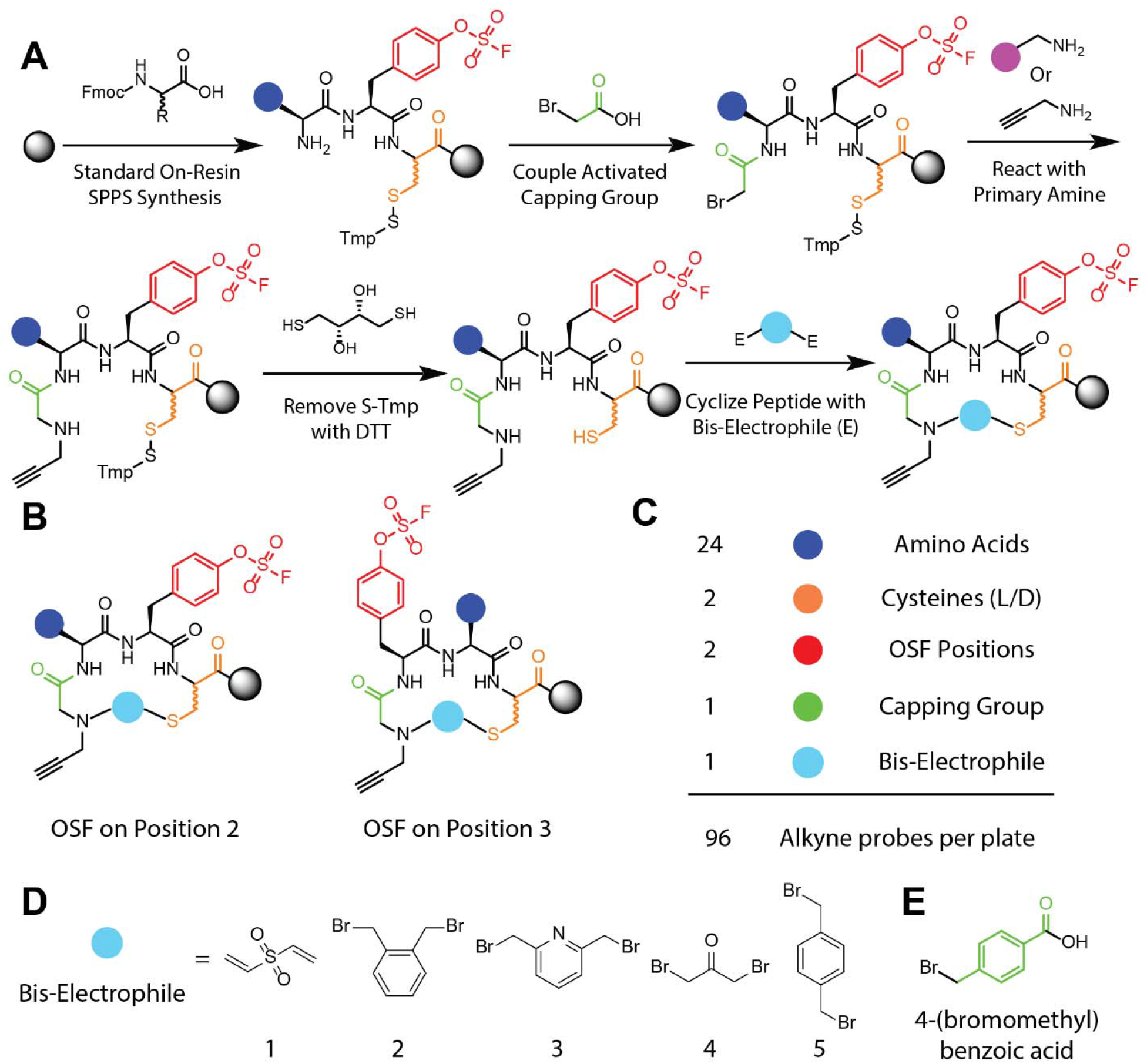
Synthetic method and components for generation of covalent macrocycles for chemoproteomic applications (**A**) Simplified scheme for the solid phase synthesis of macrocycles which contain a OSF and alkyne handle. (**B**) General structures for macrocycles in the library. The position of the electrophilic Tyrosine-(OSF) and variable amino acid was switched to generate additional diversity. (**C**) List of the diversity of each component used to generate the 96 unique macrocyclic probes per bis-electrophile cyclizing agent used. **(D**) Structures of the bis-electrophiles used to generate libraries. (**E**) Structure of additional capping group 4-(Bromomethyl)benzoic acid which formed significant side products during the ring closing reaction.

We used all five linkers with both a bromoacetate and 4-(Bromomethyl)benzoate capping group. However, preliminary tests indicated that the 4-(Bromomethyl)benzoate cap (**Figure 2E**) tended to form significant side products during the ring closing with all the linkers except linker 1 and 2. Therefore, we synthesized five macrocyclic libraries with a bromoacetate capping group, with an additional two libraries synthesized with the 4-(Bromomethyl)benzoate cap (**Figure 2E**) and linker 1-2. Overall, we synthesized the seven OSF macrocyclic sub-libraries using automated high-throughput SPPS on an automated synthesizer and cleaved these libraries using standard cleavage mixture of triisopropyl silane, water, and trifluoroacetic acid, followed by precipitation in ether directly in 96 well plates. We randomly selected 20 peptides from each sub-library and assayed purity by liquid chromatography mass spectrometry (LC-MS) analysis. The majority of samples produced signals corresponding to the predicted mass and the overall purity was assessed by LC/MS analysis (see **Supplementary Spectra)**.

After synthesis of the sub-libraries, we sought to rapidly screen compounds in crude cellular lysates to identify macrocycles which showed specific labeling of proteins. We screened the macrocycles in HEK 293 lysates using a final probe concentration of 10 μM for two hours at 37°C followed by CuAAC click reaction with tetramethylrhodamine azide for visualization. To avoid SDS-PAGE analysis of all 480 molecules, we selected and tested 24 macrocycles each with a unique amino acid on a specific scaffold to identify compounds that gave specific labeling patterns of proteins (**Figure 3A**). We choose to use the base scaffold (**Figure 3B**) of L-Cysteine and OSF on position 2 to identify amino acid specific labelling. If a specific amino acid produced a unique labelling profile, we performed a secondary screen (**Figure 3D**) in which we held that amino acid of interest constant and compared the impact of altering the electrophile position and cysteine stereochemistry (**Figure 3C**). In general, the macrocycles showed relatively faint labeling of a few proteins, consistent with the general low reactivity of the OSF electrophile (**Figure 3A**). Furthermore, there were very little differences in labeling patterns across the various amino acids with unique labeling patterns occurring rarely (**Figure 3A** column G, glycine). When we identified labeling patterns that appeared specific for a given macrocycle, we then followed up by assessing the labeling of the closest relatives in the sub-library in which the position of the electrophile and the stereochemistry of the cysteine were varied. This allowed us to focus on compounds that labeled proteins in a manner that was specific to the cyclic peptide scaffold (**Figure 3D**).

**Figure 3.**
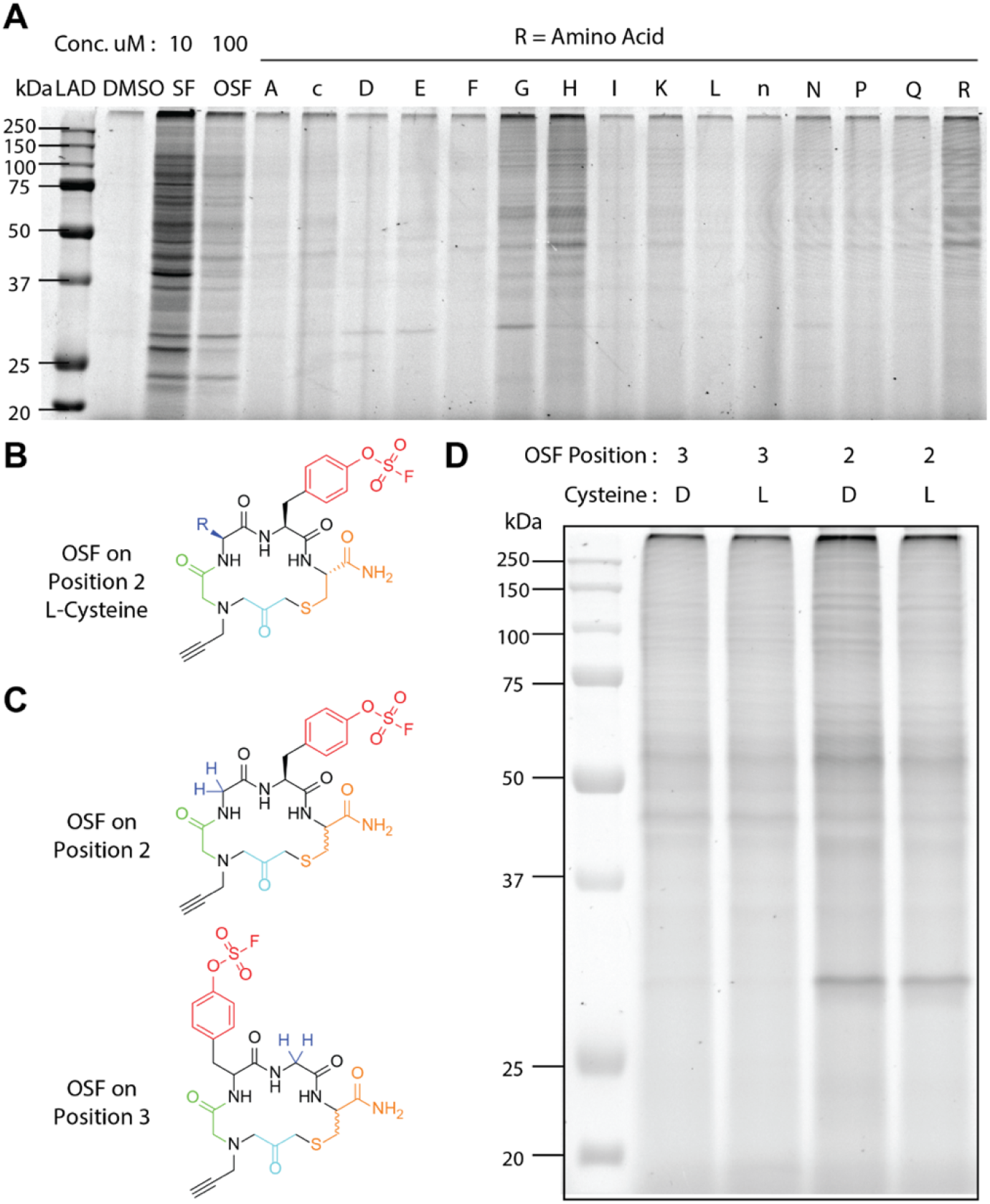
Rapid *in vitro* screening of covalent macrocycles in HEK 293 lysates by SDS-PAGE. Probes were incubated at 10 μM for two hours at 37°C in HEK 293 lysate then labeled by CLICK reaction with TMR-azide for one hour at r.t. and then 25 μg protein was analyzed by SDS-PAGE. Fluorescent labeling was visualized with an Azure Biosystems Sapphire Biomolecular Imager. (**A**) Representative gel showing screen of a set of macrocycles with the core structure shown in panel B containing a range of amino acids indicated by the one letter codes. Letters represent the one letter amino acid code. c is the unnatural amino acid O-benzyl-L-homoserine and n is norleucine (**B**) General structures for macrocycles used for labeling in panel A with the R group representing the variable amino acids. (**C**) Core structure for probes in panel D. The electrophile position and cysteine stereochemistry are varied. **(D**) Gel labeling image for probes from panel C showing the change in labeling upon change of the position of the electrophile.

After this initial screening, we identified six macrocycles of interest which produced distinct labeling patterns that were dependent on the position of the electrophile and the stereochemistry of the cysteine used for the cyclization. We resynthesized these hits and purified them via High Performance Liquid Chromatography (HPLC). Two probes (**Figure 4A**) showed intense labelling compared to the OSF and SF alkyne probes (**Figure 4C**). However, we determined that this labeling was specific to the vinyl sulfone linker used for cyclization and was due to incomplete cyclization of the compounds. This resulted in a linear compound containing a free vinyl sulfone that reacted with proteins independently of the OSF electrophile. This problem was missed in our quality control analysis of the libraries due to the fact that the single adduct of the linker was the exact same mass as the cyclized product. To confirm that the labeling was due to the free vinyl sulfone and not the OSF electrophile we synthesized several compounds with the vinyl sulfone linker but lacking the OSF electrophile. Those compounds also produced strong labeling patterns in lysates, confirming that the labeling was not the result of the OSF electrophile (data not shown). We therefore eliminated all compounds from these two sub-libraries from further analysis. We therefore ultimately selected the P4:30, P2:9, P2:12, and P2:15 probes (**Figure 4B**) as they also produced specific labeling patterns dependent on the electrophile position and cysteine stereochemistry. Using these purified probes, we repeated the labeling of HEK 293 lysates and found good correlation with the previous labelling patterns (**Supplementary Figures 1-3**). For all four probes, we also synthesized control macrocycles using propyl amine in place of the propargyl amine for competitive labelling experiments. We chose to further pursue probe P4:30 for chemoproteomics to identify potential targets because it showed the strongest competition with the no alkyne control macrocycles, (**Supplementary Figures 1-3**).

**Figure 4.**
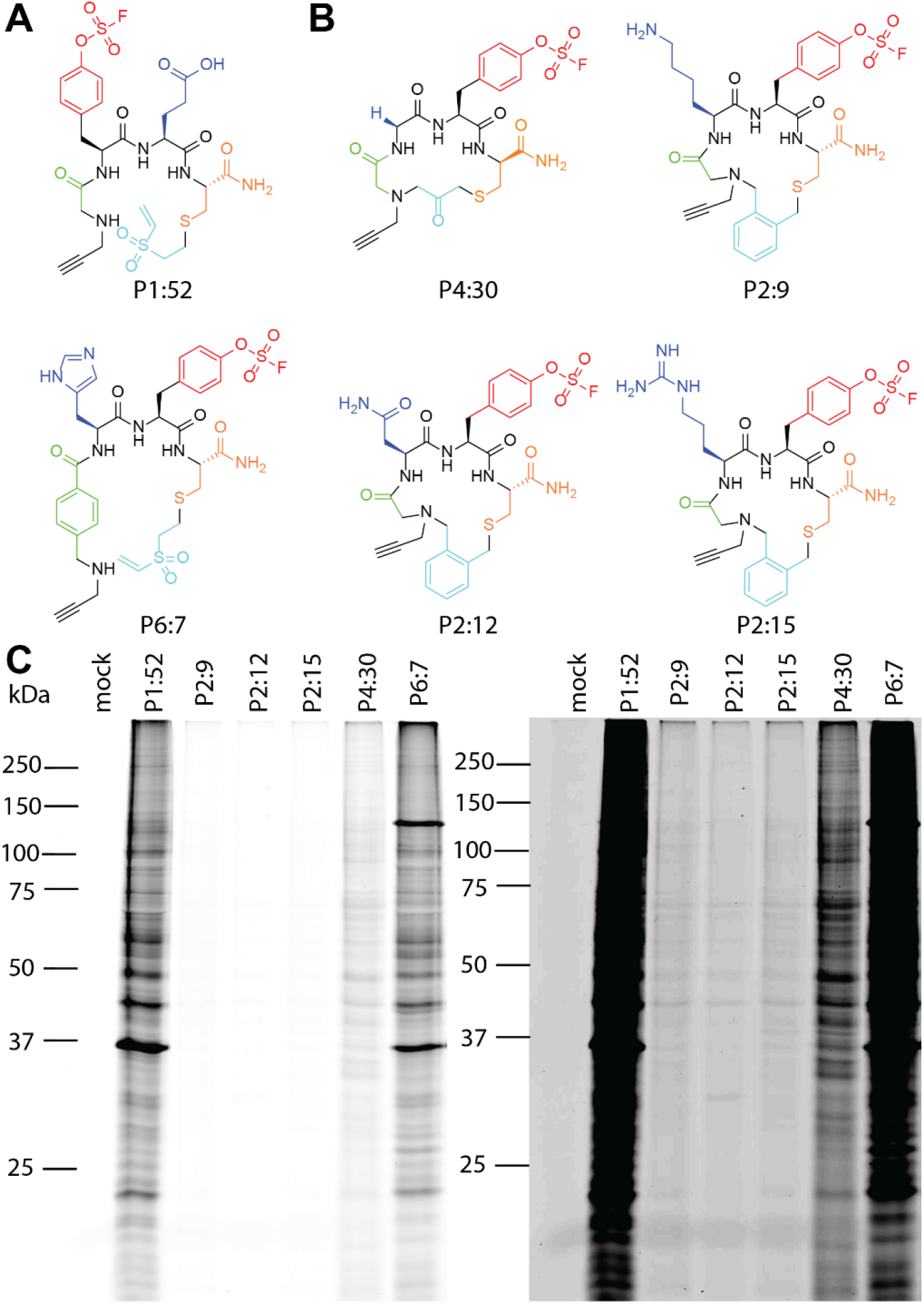
Protein labeling of selected hits from the library of OSF macrocycles. (**A**) Structures of the linear vinyl sulfone molecules which failed to cyclize and produced intense labeling profiles in the crude extracts. (**B**) Structures of macrocycles P4:30, P2:9, P2:12, and P2:15 used for labelling in panel D. **(C**) *In vitro* labelling of HEK 293 lysates with purified and concentration matched probes shown in A and B. Probes were incubated at 10 μM for two hours at 37°C in HEK 293 lysate then labeled by CLICK reaction with TMR-azide for one hour at r.t. and then 25 μg protein was analyzed by SDS-PAGE. Fluorescent labeling was visualized with an Azure Biosystems Sapphire Biomolecular Imager.

To identify targets of P4:30 we performed *in vitro* LC-MS/MS chemoproteomic analysis. We included samples in which we incubated HEK 293 lysates with the no alkyne control macrocycle at varying concentrations prior to labeling with P4:30 (**Supplementary Figure 3**). The top hits for probe P4:30 were translin (TSN) and the proteasome subunit alpha type-5 (PSMA5), with both being competed at 30 μM using a 4-fold enrichment cutoff. We identified an additional six proteins targets (AHSA1, PDCD6IP, ERP29, HYOU1, RAD23B, and NUDT21) that were competed at 100 μM of the control no-alkyne probe (**Figure 5A and Table S3**). Interestingly, we identified tyrosine adducts to AHSA1, PDCD6IP, ERP29, HYOU1, RAD23B, and NUDT21 using the general SF alkyne probe but not with the general OSF alkyne probe (**Tables S1-S3**). This data indicates that proteins that contain residues that can be targeted only with a highly reactive SuFEx electrophile can also be targeted with less reactive electrophiles if the ligand imparts enough binding energy for the SuFEx reaction to take place. To further validate the top hits (TSN and PSMA5) as targets of the P4:30 probe, we performed direct labeling experiments using purified commercial protein (**Figure 5B**). These results confirmed that P4:30 efficiently labelled translin with improved potency compared to OSF alkyne which was almost undetectably labeled even at 500 μM OSF alkyne. Interestingly, labelling of human 20S proteasomes with P4:30 in two different buffer systems failed to produce specific labeling of the PSMA5 subunit. This result may be due to the fact that the proteasome is highly abundant in cells and was non-specifically modified by the probe. It is also possible that the PSMA5 subunit is specifically labeled but only in a subset of proteasomes in the lysates that require capping groups or alternate modification which is not present in the purified 20S sample.

**Figure 5.**
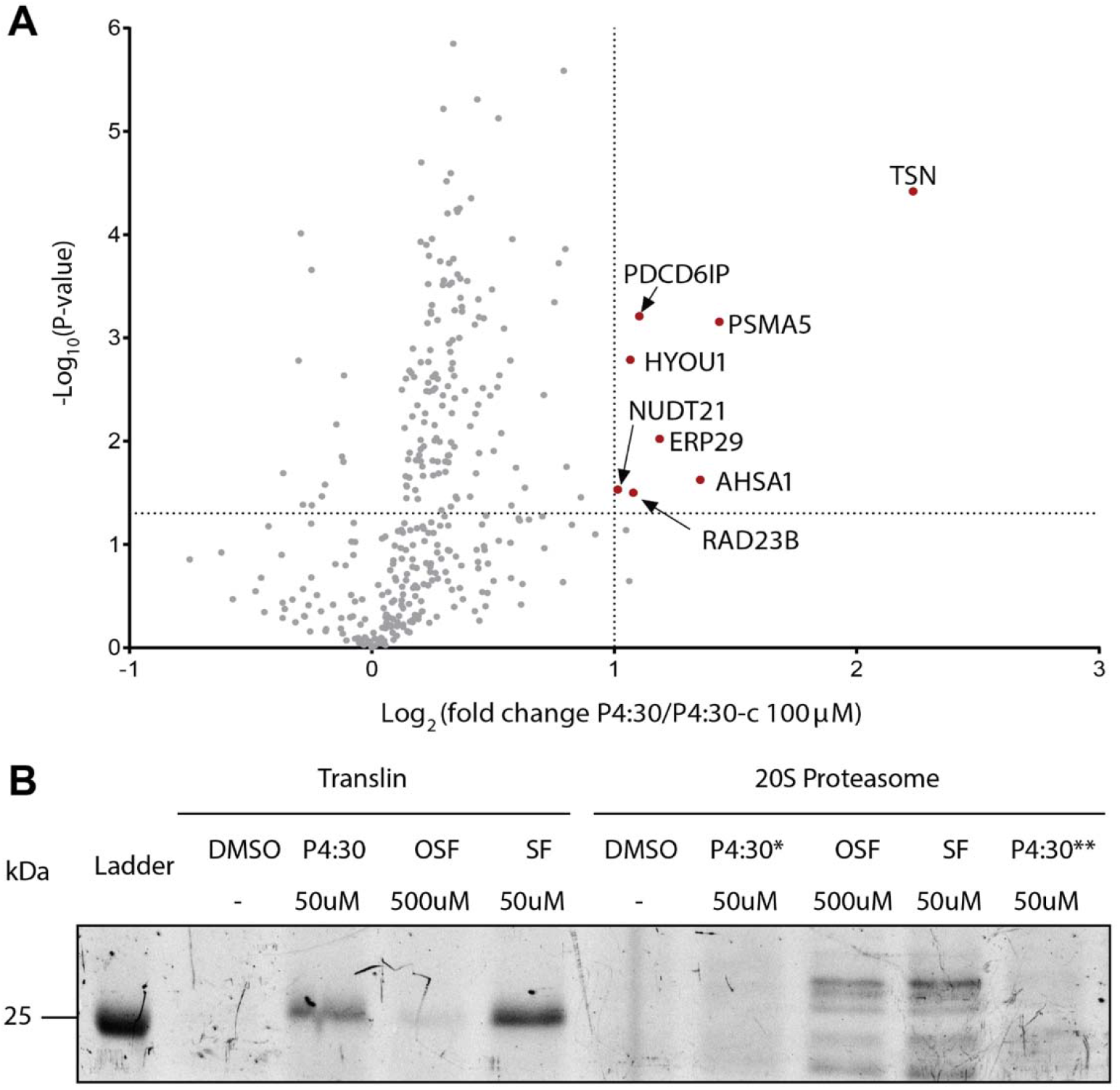
Chemoproteomic identification of targets of the macrocycle P4:30 (**A**) Volcano plot generated from *in vitro* LC-MS/MS chemoproteomic analysis of probe P4:30 (10 μM) and no alkyne control P4:30c (100 μM) in HEK 293 lysate (n = 4). X-axis represents the ratio of labeling with and without pretreatment with the no-alkyne control. Y-axis indicates P value. Dashed lines represent 50% competition and 5% P-value (**B**) Direct labeling experiments using purified commercial translin and 20S proteasome with probes P4:30, SF alkyne, and OSF alkyne in HEK 293 lysate. Probes were incubated for 2 hours at 37°C at 50 μM or 500 μM with 50 ng translin or 1500 ng of 20S Proteasome then labeled by CLICK reaction with TMR-azide for one hour at r.t. and analyzed by SDS-PAGE. Fluorescent labeling was visualized with an Azure Biosystems Sapphire Biomolecular Imager. Both a Tris Base (*) and HEPES (**) buffer were used for 20S proteasome labelling with probe P4:30.

Overall, this work provides evidence that the OSF electrophile can be coupled to macrocyclic scaffolds to generate new covalent binding molecules with defined target selectivity. Our optimized synthetic strategy enables fully on-resin synthesis to generate macrocycles that are suitable for rapid screening in lysates using SDS-PAGE methods and for target identification using standard chemoproteomic workflows. We identified challenges in the synthesis design and implementation that will inform future scaffold designs. Specifically, we identified conditions and multiple building blocks that were compatible with the SPPS approach, as well as one linker which requires further optimization. Importantly, this dataset demonstrates that linker diversity and structural permutations could be modulated to change the target reactivity of the OSF electrophile. Furthermore, while including an alkyne in the N-terminal position of all macrocycles enabled an efficient drug discovery workflow, it also limited chemical diversity.^[3–6]^ Future libraries could overcome this limitation by use of a diverse array of amines as the N-terminal capping group and inclusion of the alkyne group at the C-terminal end of the peptide. Overall, the methods developed here can be translated to other electrophiles compatible with SPPS workflows.

## Conclusion

In this report, we developed a solid phase strategy to generate libraires of macrocycles containing the fluorosulfonate (OSF) electrophile that are compatible with drug discovery workflows. We explored the reactivity landscape of the OSF electrophile compared to the more commonly used sulfonyl fluoride (SF) electrophile. Furthermore, we tested a variety of chemical components for compatibility with this library synthesis and identified areas of improvement for library design. We also developed a screening strategy in which we analyzed a sub-set of macrocycles for their labeling of protein targets in a complex proteome using simple SDS-PAGE analysis and then identified specific scaffolds of interest for chemoproteomic analysis. We found that the macrocyclic scaffolds directed target selectivity and potency of the OSF electrophile suggesting that these scaffolds should be further investigated for use with latent electrophiles. Ultimately, we confirmed that the overall low reactivity of the OSF electrophile results in a very narrow set of protein targets but this target set can be modulated through the use of structurally distinct binding scaffolds such as the macrocycles described here. We believe that the OSF electrophile is an optimal choice for drug development applications as it can be directed to react with specific protein targets while its overall low reactivity prevents unwanted modification of off targets.

## Supporting information

Supplemental Spectra

## Acknowledgments

This work was supported by NIH grants (R01 EB026285, to M.B.), the National Institutes of Health (R01 GM145886, to A.A.), the National Science Foundation Graduate Research Fellowship under grant no. DGE-1656518 (F.F.), Stanford ChEM-H O’Leary-Thiry Graduate Fellowship (F.F.), NIH Stanford Graduate Training Program in Biotechnology T32GM141819 (F.F.), International Alliance for Cancer Early Detection (ACED) Graduate Fellowship (F.F.).

## Author contributions

S.L., A.A., M.B. designed the research project. F.F., and D.A. carried out all the experiments. P.I. aided with synthesis and labelling. F.F., and M.B. wrote and edited the manuscript with input from all authors.

## Data Availability Statement

The data that support the findings of this study are available in the supplementary material of this article.

---

**Scheme 1.**
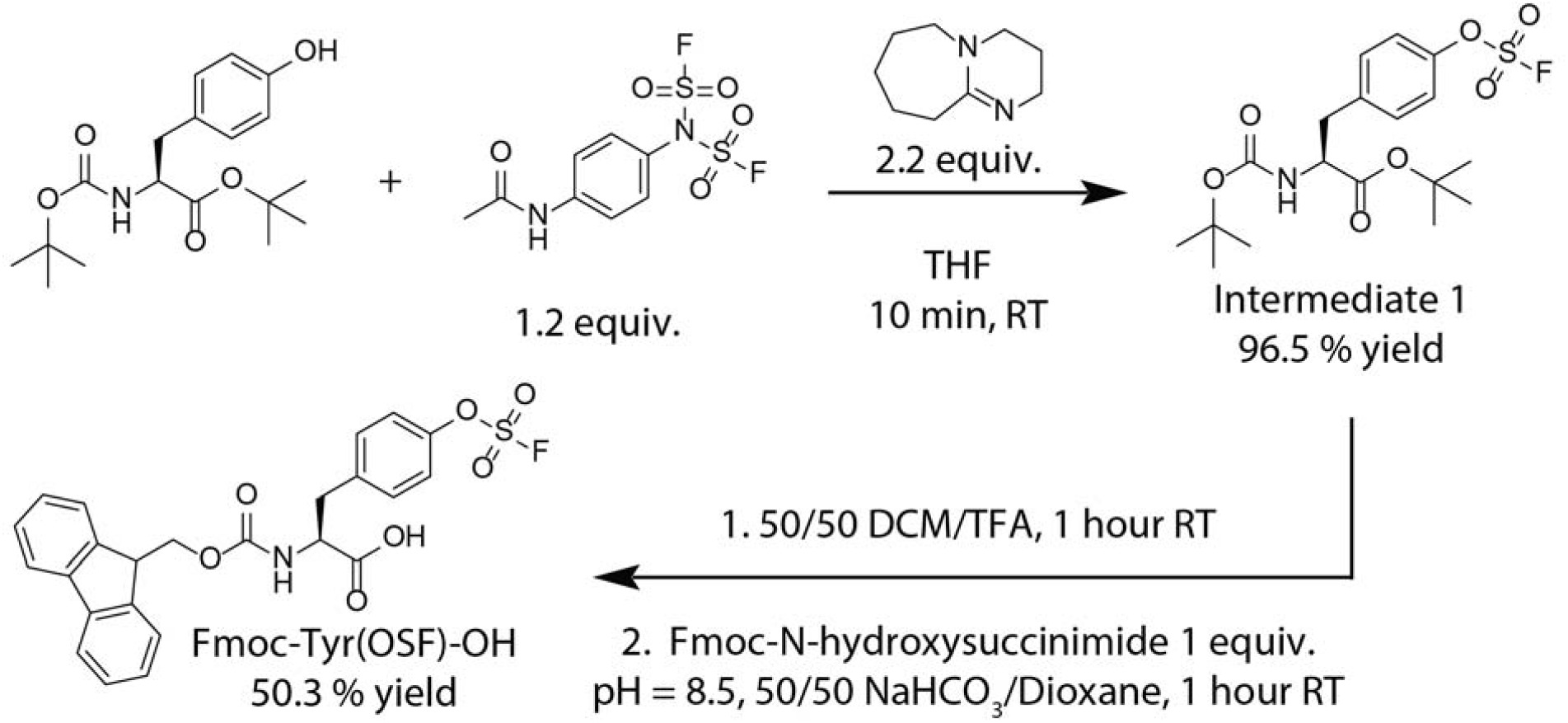
Synthesis of Fmoc-fluorosulfated tyrosine (Fmoc-Tyrosine(OSF)-OH) using the shelf-stable, crystalline reagent [4-(acetylamino)phenyl]imidodisulfuryl difluoride (AISF).

