## Supplemental figures and methods for "Solid Phase Synthesis of Fluorosulfate Containing Macrocycles for Chemoproteomic Workflows"

### Supplementary Figures

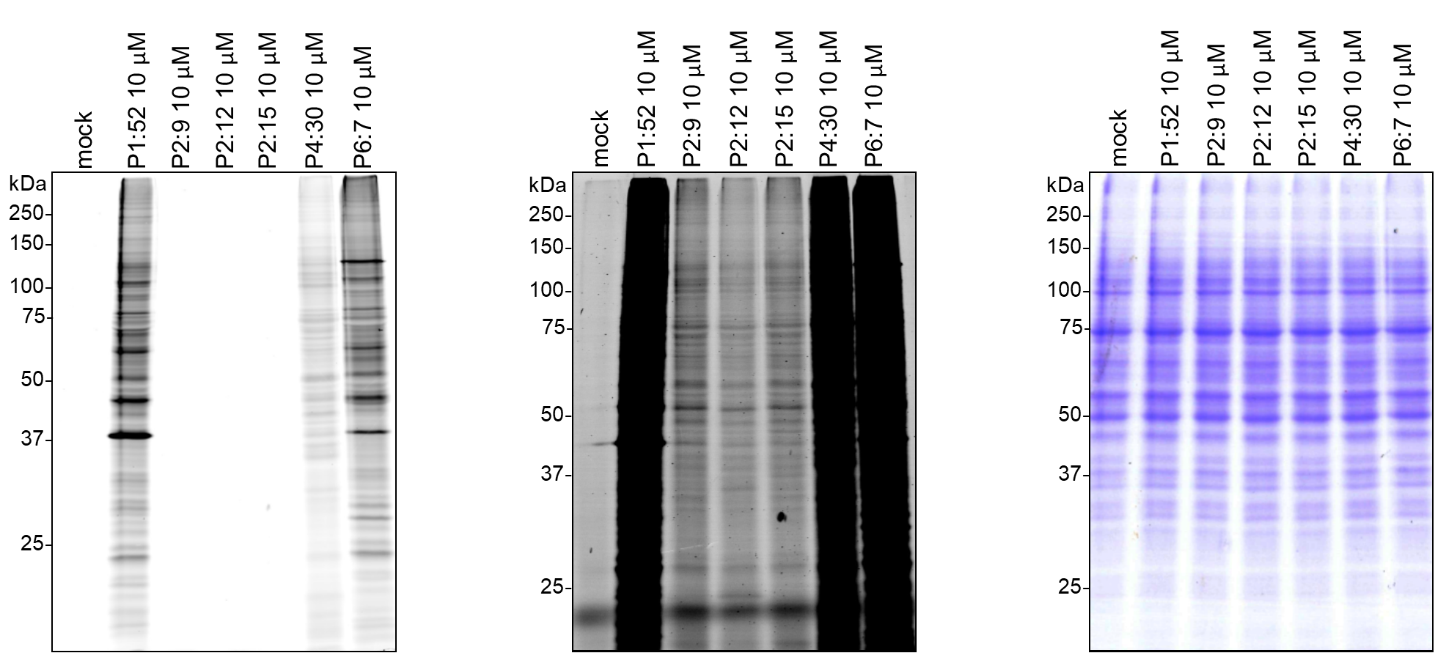

**Supplementary Figure 1** Protein labeling patterns of selected hits from the library of OSF macrocycles. HEK 293 lysate (50 µg in 25 µL) were treated with DMSO for two hours at 37°C, then with indicated probes for two hours at 37°C, followed by click chemistry with 50 µM TMR-azide for one hour at r.t. Reducing loading buffer was added, proteins were separated by SDS-PAGE and visualized with an Azure Biosystems Sapphire Biomolecular Imager, then stained with Coomassie.

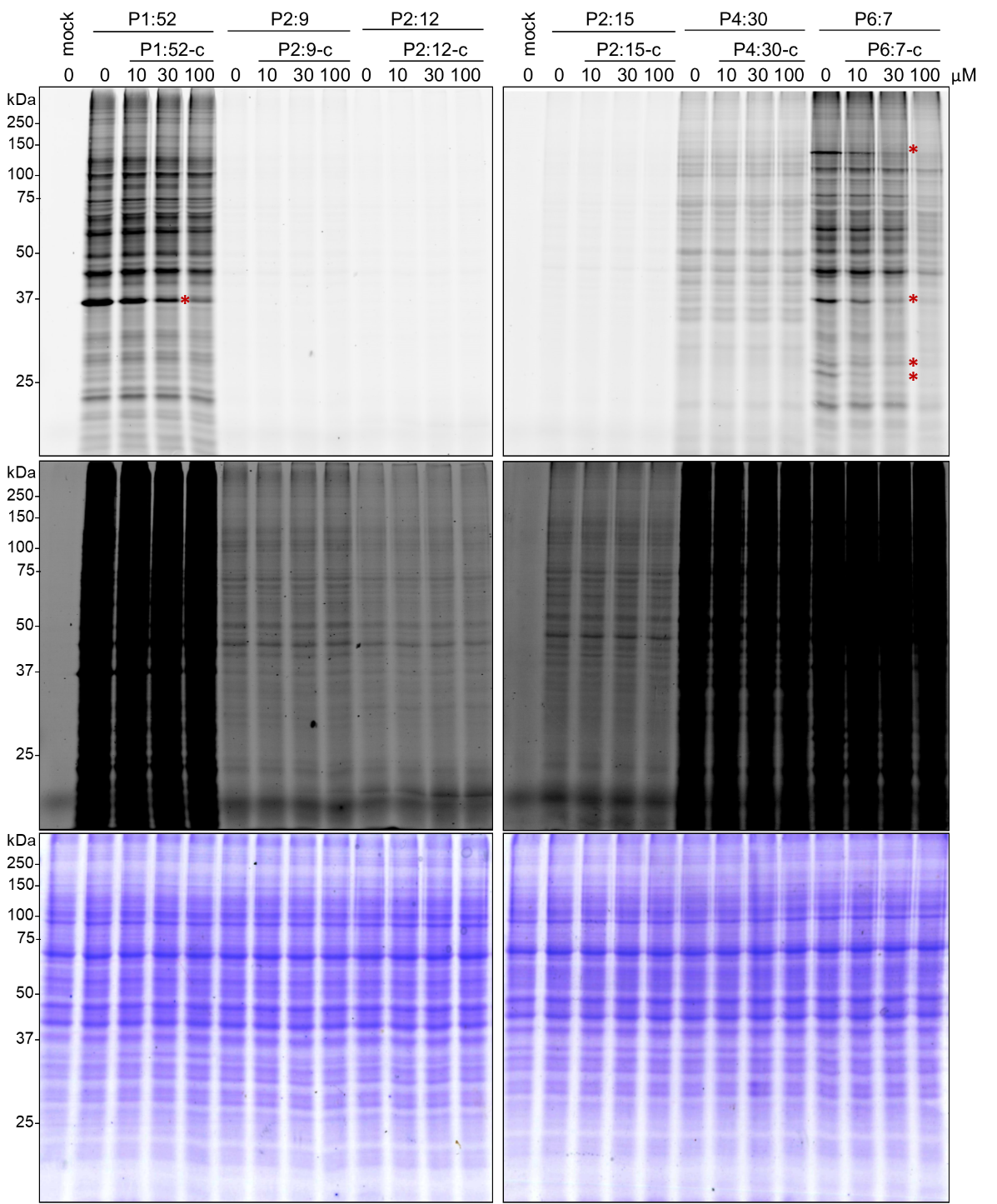

**Supplementary Figure 2** Competitive protein labeling patterns of selected hits from the library of OSF macrocycles. c indicates control compound. HEK 293 lysate (50 µg in 25 µL) were treated with indicated compounds for two hours at 37°C, then with indicated probes for two hours at 37°C, followed by click chemistry with 50 µM TMR-azide for one hour at r.t. Reducing loading buffer was added, proteins were separated by SDS-PAGE and visualized with an Azure Biosystems Sapphire Biomolecular Imager, then stained with Coomassie. * Indicates competition with probes that failed to cyclize.

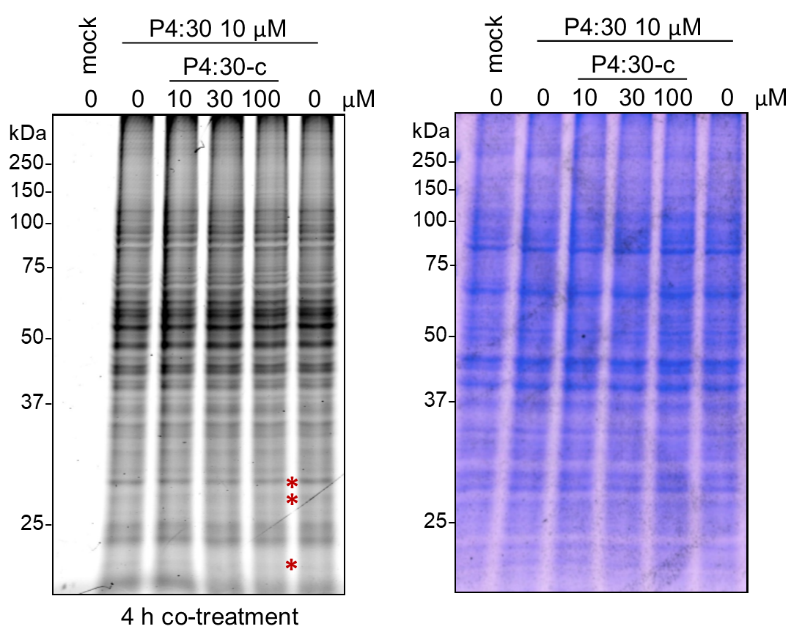

**Supplementary Figure 3** P4:30 *in-vitro* competition optimized. HEK 293 lysate (50 µg in 25 µL) were co-treated with indicated concentration of P4:30-c and 10 µM P4:30 for four hours at 37°C, followed by click chemistry with 50 µM TMR-azide for one hour at r.t. Reducing loading buffer was added, proteins were separated by SDS-PAGE and visualized with an Azure Biosystems Sapphire Biomolecular Imager, then stained with Coomassie. * Indicates competition.

### Rapid Screen SDS-PAGE Gels

HEK 293 lysate were treated with DMSO for 2h at 37°C, then with the indicated probes for two hours at 37°C, followed by click chemistry with TMR-azide for 30 min at r.t. Reducing loading buffer was added, proteins were separated using a 12% SDS-PAGE gel. Gels were visualized using a GE Typhoon FLA 9000, then stained using Coomassie.

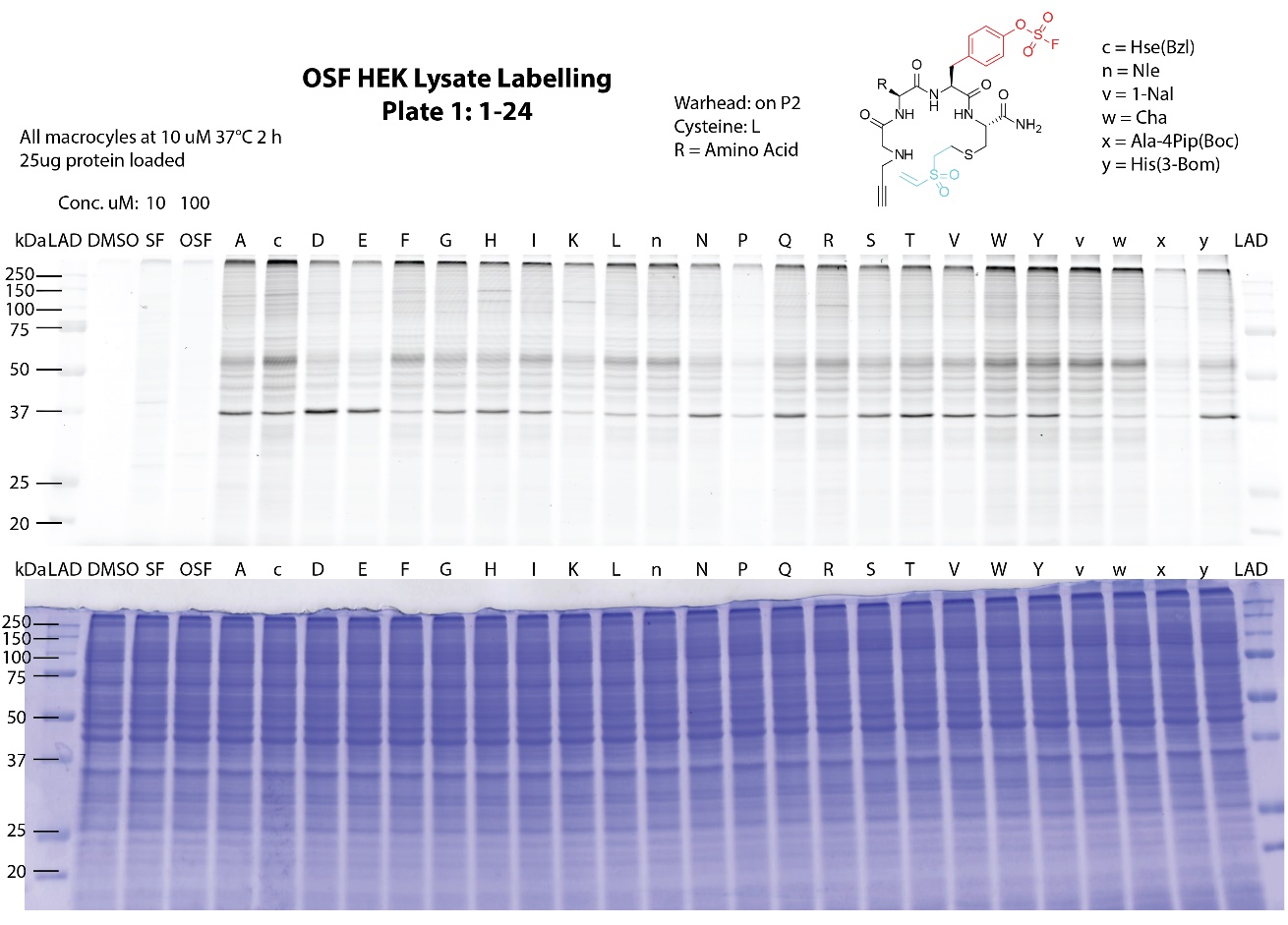

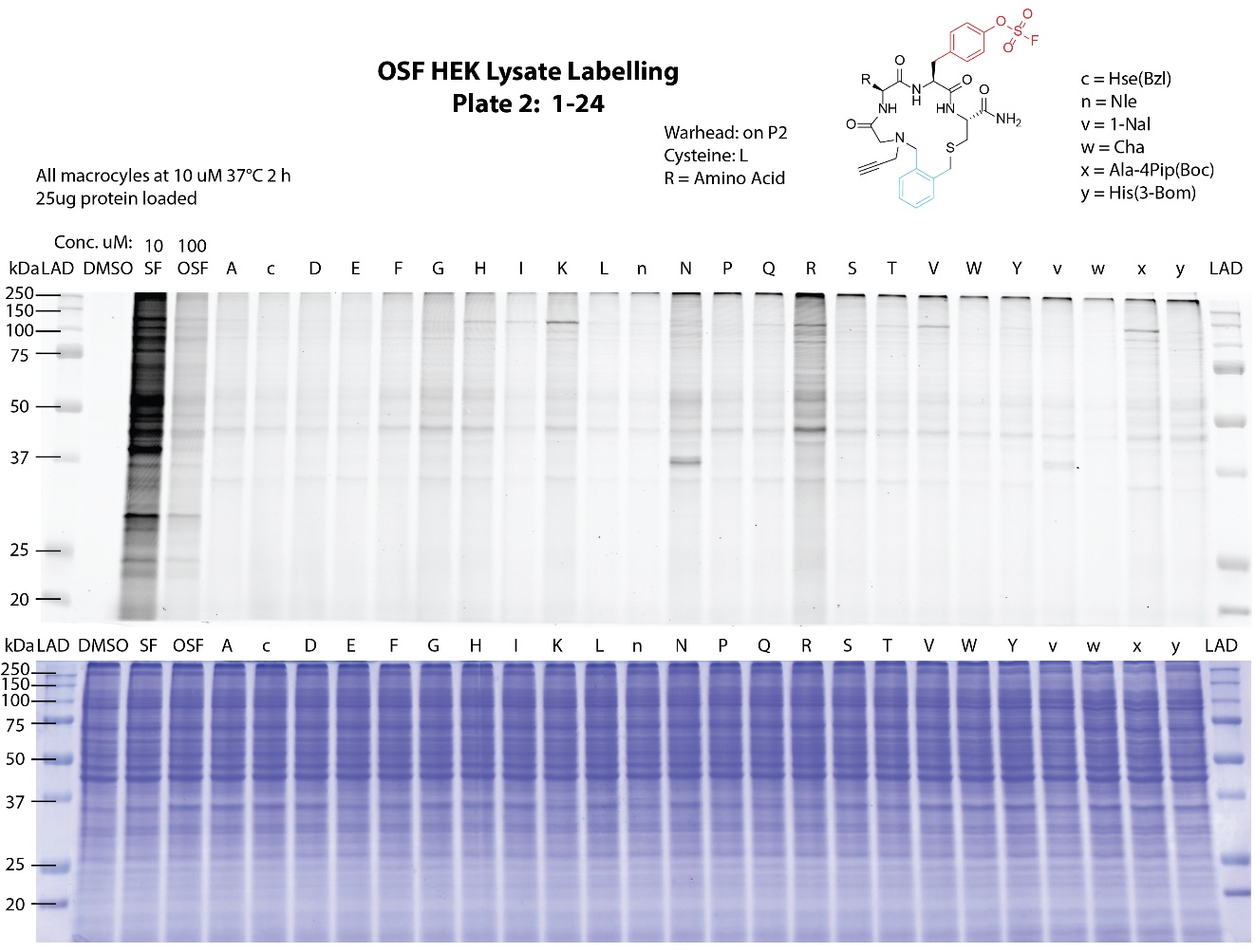

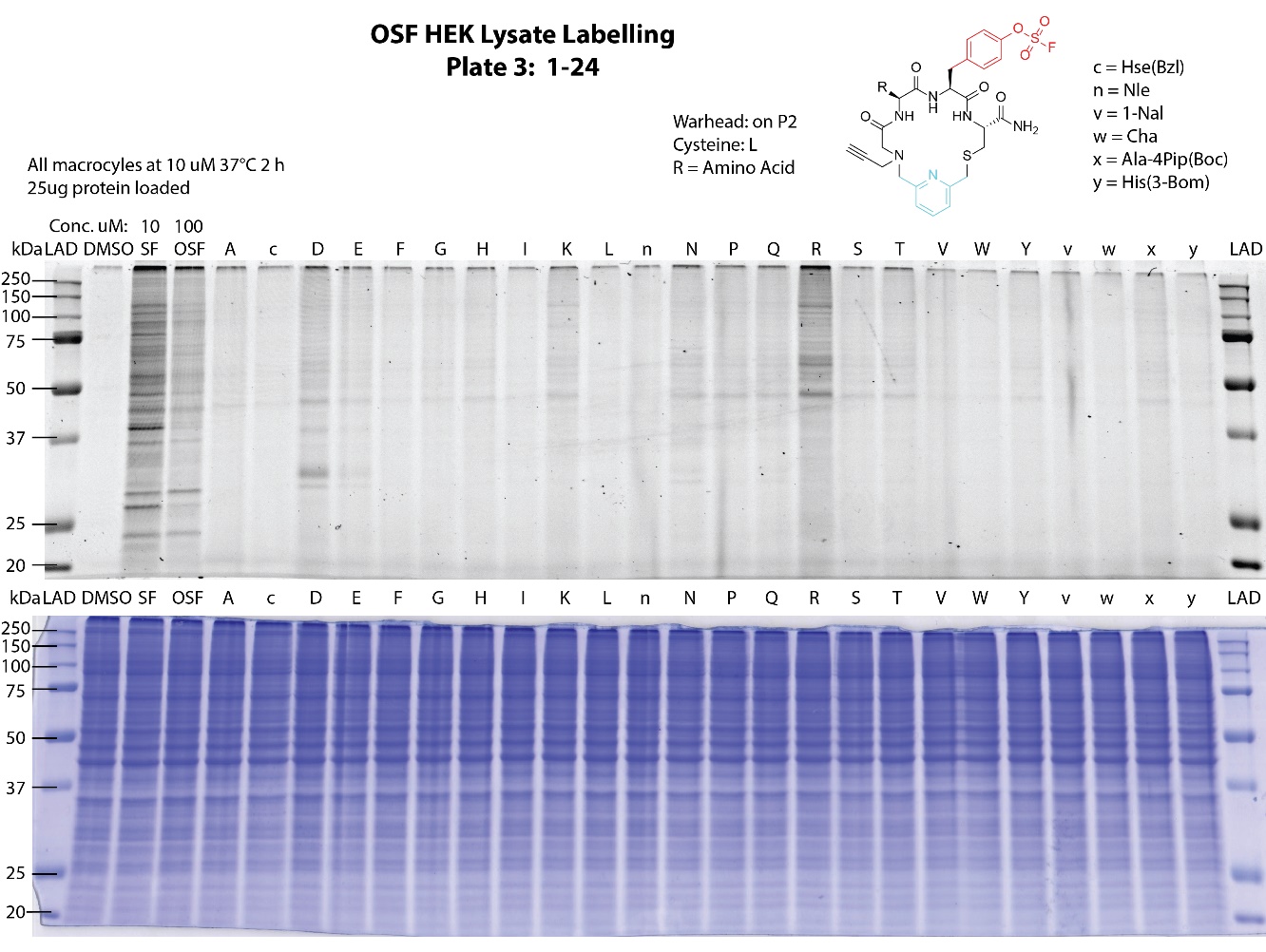

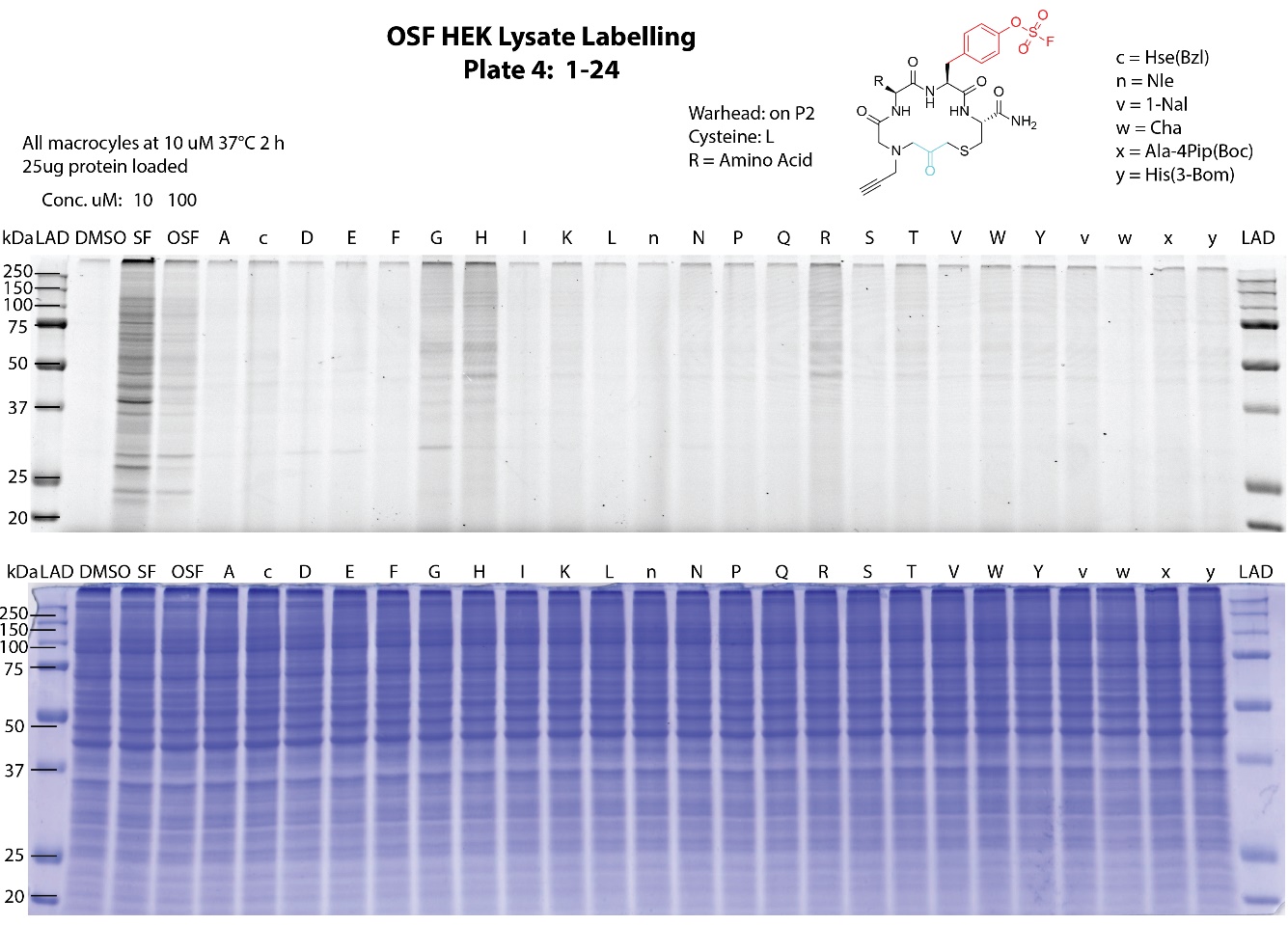

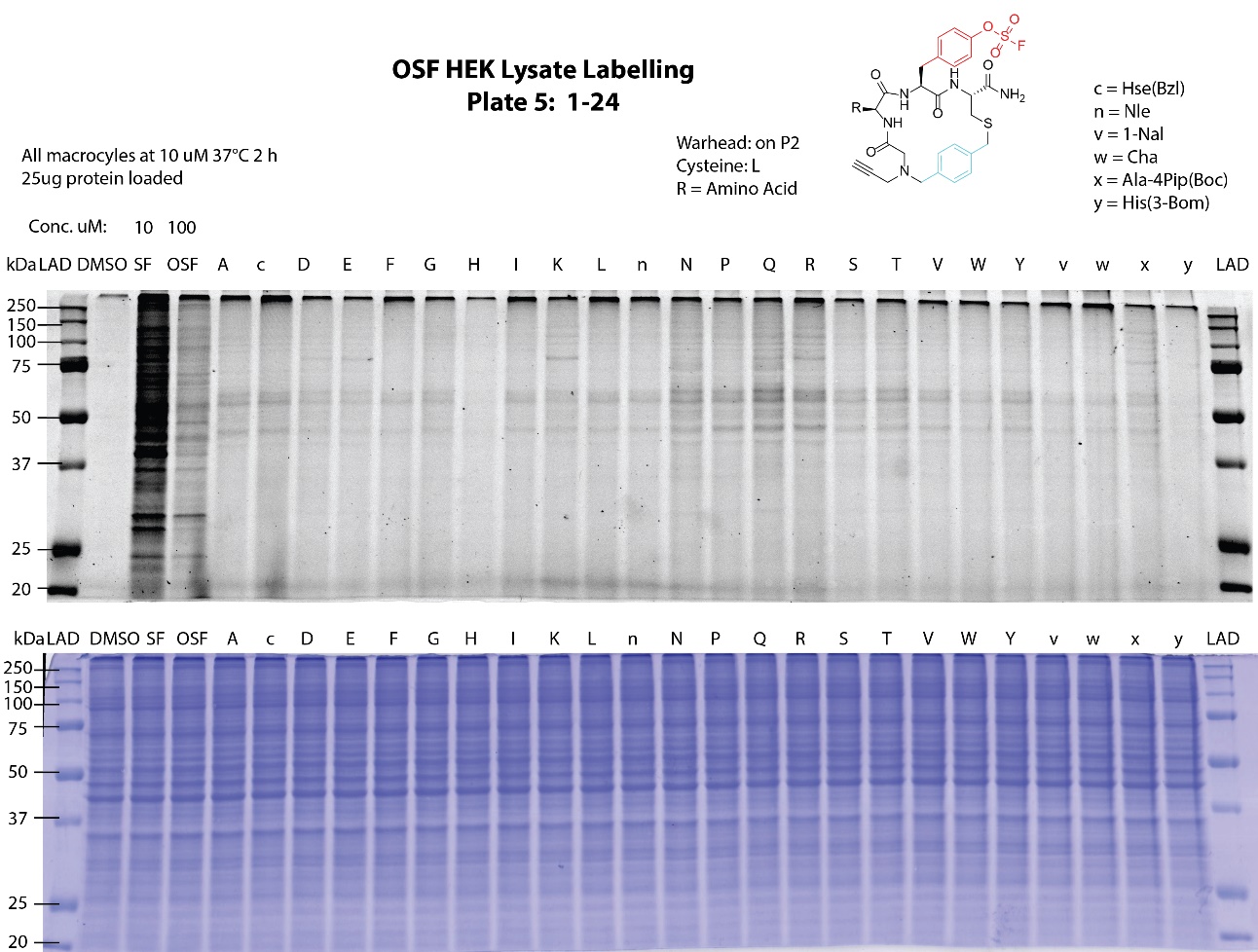

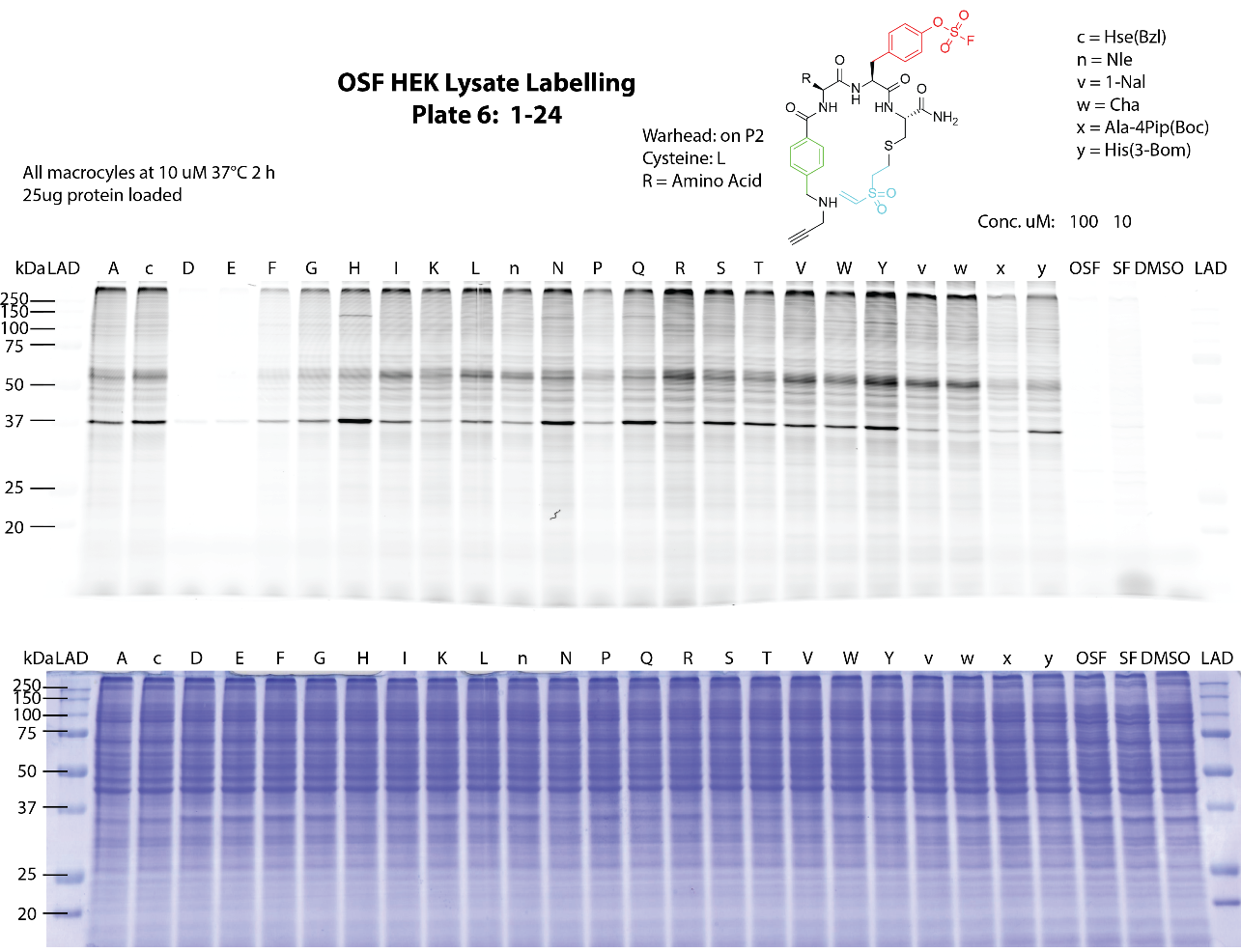

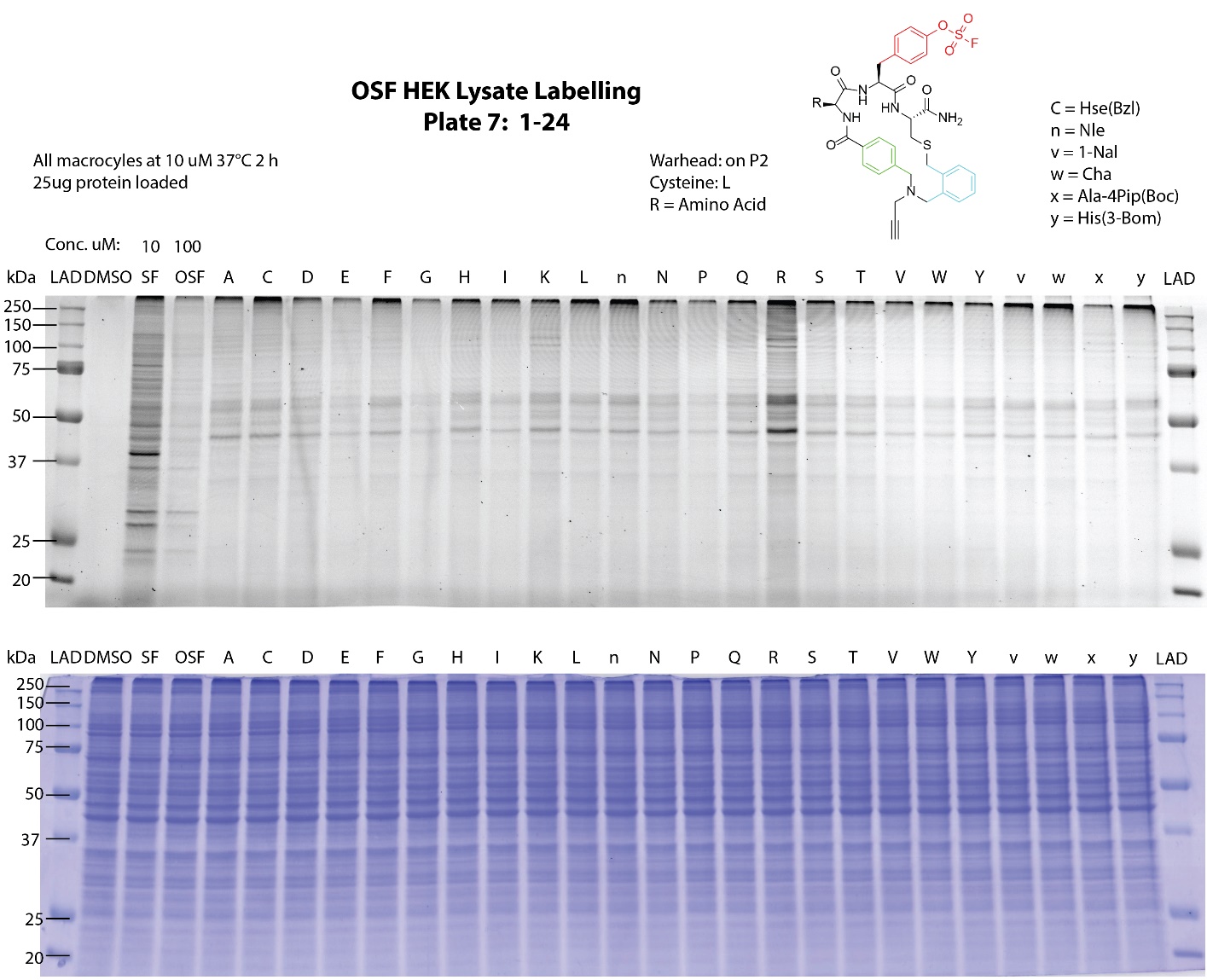

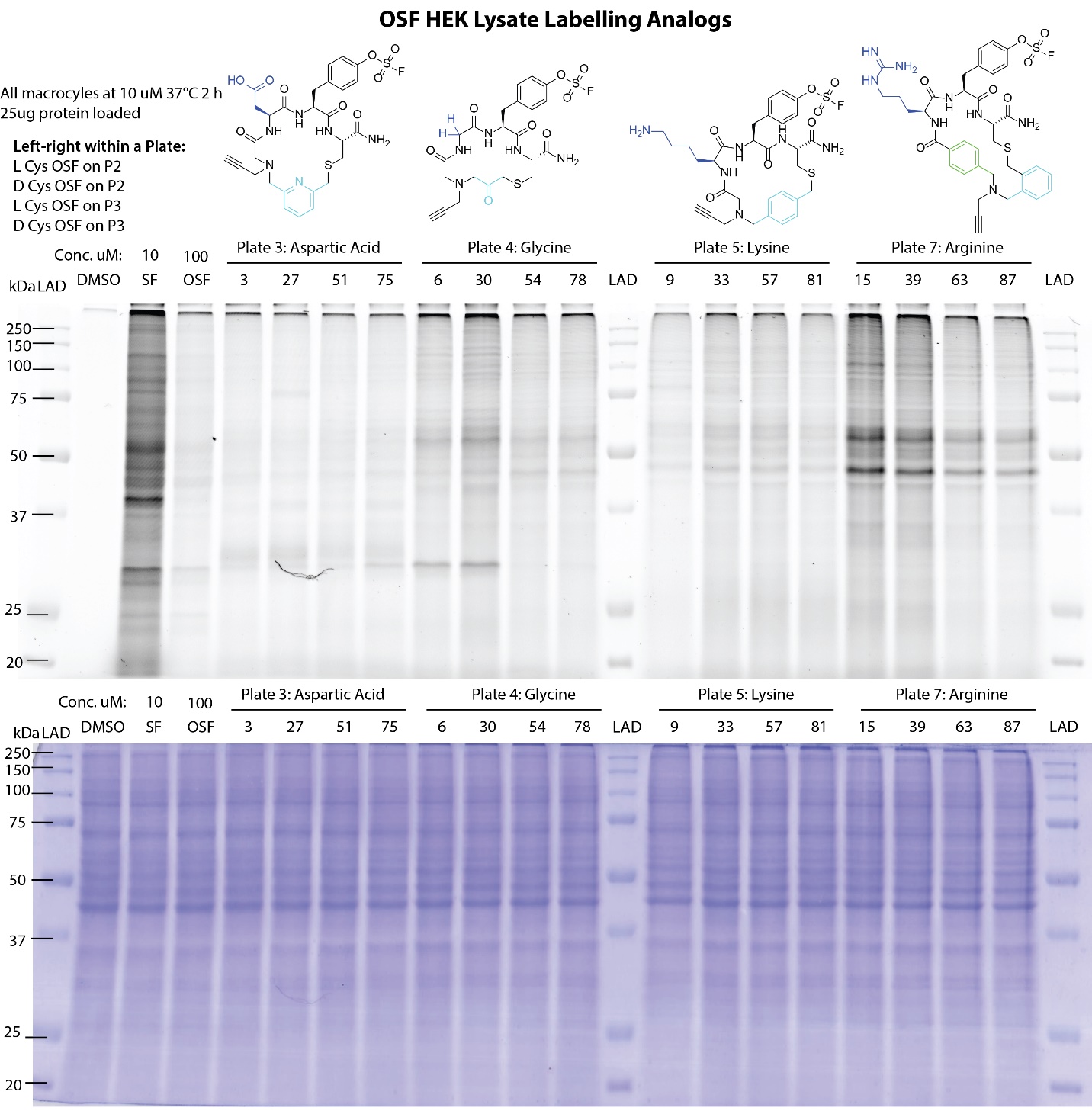

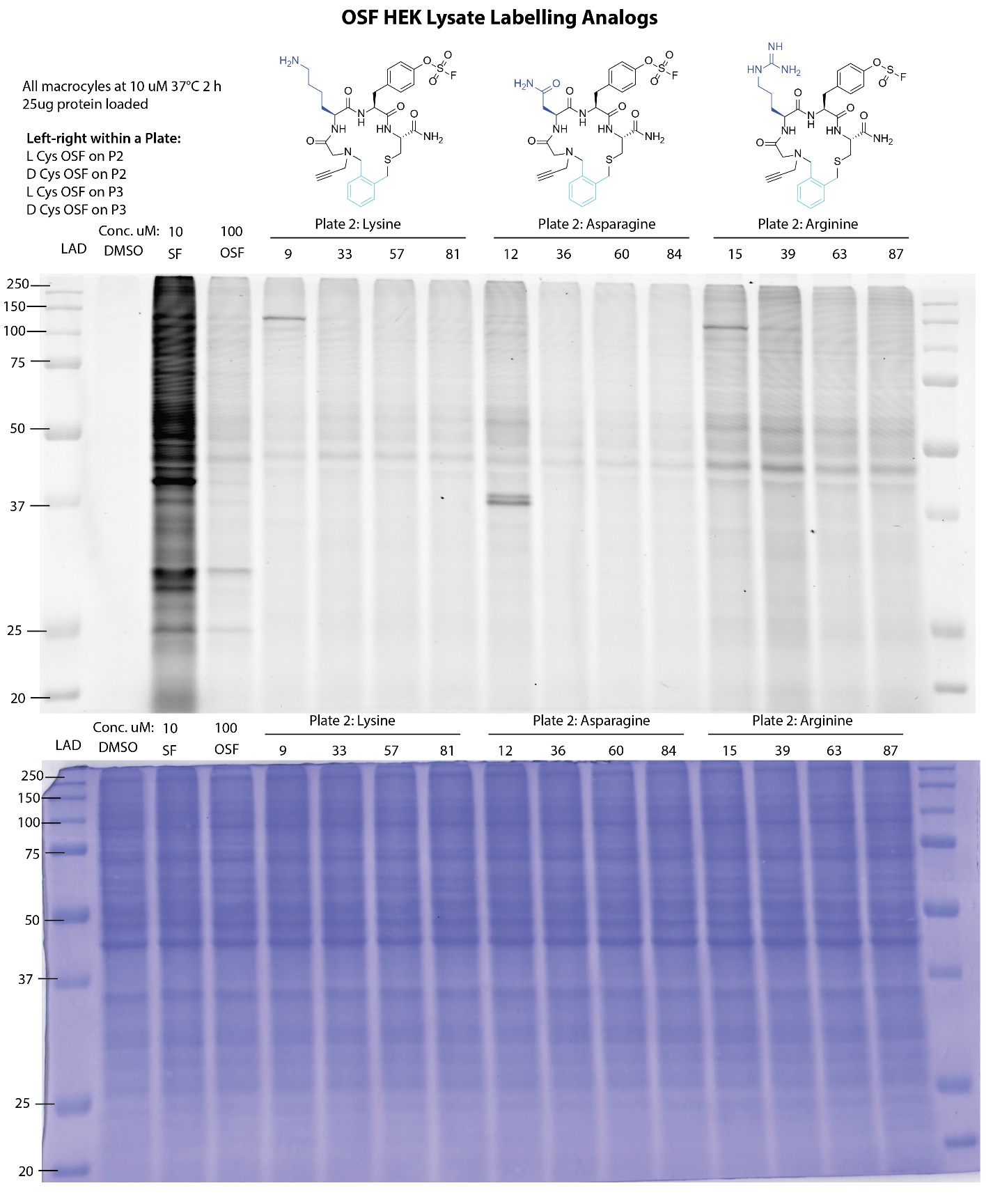

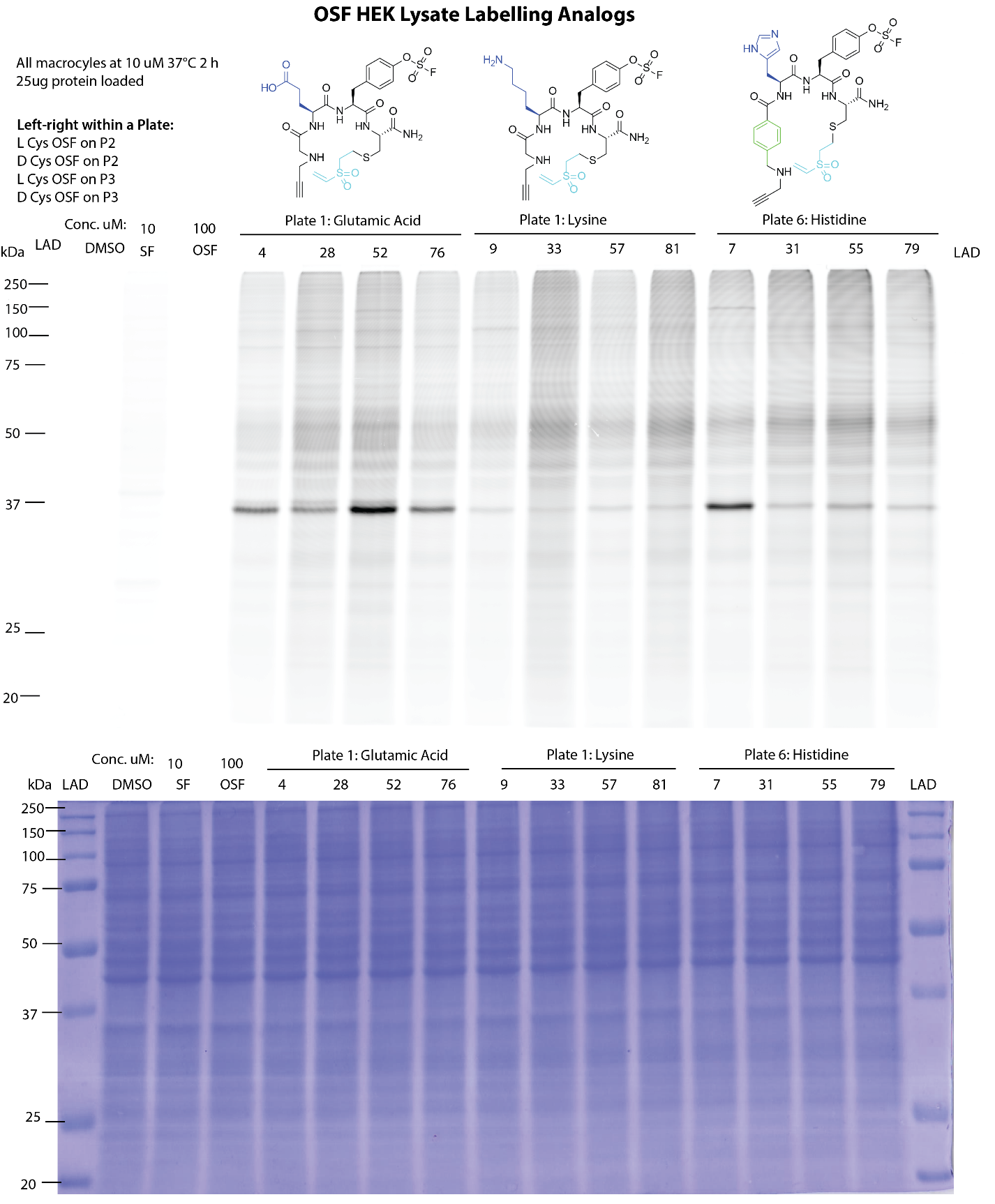

### Methods

#### **Chemistry Methods**

**Materials and Synthetic Methods**. All reactions were performed exposed to atmospheric air and with solvents not previously dried over molecular sieves or other drying agents, unless specified. The ACS reagent grade N,N’- dimethylformamide (DMF), tetrahydrofuran (THF) containing 250 ppm of butylated hydroxy toluene (BHT), molecular biology grade dimethyl sulfoxide (DMSO), and all other commercially available chemicals were used without further purification. All reactions were performed at room temperature unless specified. Reaction progress and purity analysis was monitored using an analytical LC-MS. The LC-MS systems used was either an Agilent 1200 HPLC equipped with an Agilent Zorbax SB-C18 column (1.8 μm, 2.1 x 50 mm) coupled to an Agilent 6125B Single Quad Mass Spectrometer or an Agilent 1100 Series HPLC equipped with a Luna 4251-E0 C_18_ column (3 μm, 4.6 x 150 mm) coupled to a PE SCIEX API 3000 mass spectrometer (wavelengths monitored = 215, 254, 600 nm). Purification of intermediates and final compounds was carried out using a CombiFlash Rf + packed with silica gel (wavelengths monitored = 215 & 254 nm). Information regarding gradient programs for purifications can be found in the Chemistry Protocols section below. Intermediates were identified by their expected m/z using LC-MS.

**Chemistry Protocols.**

*The following protocols were optimized for use with the resins or reaction vessels stated.*

**Solid Phase Peptide Synthesis**. Macrocycles were synthesized on rink amide resin (loading 100-200 mesh, 0.68 meq/g, 1% DVB) (Chem-Impex Int’L INC, Cat. # 02900) using standard Fmoc chemistry. A Syro II (Biotage) fully automated parallel peptide synthesizer with 96 tips synthesis module (PP-Reactor Tip, 0.4 mL with PE Frit, Cat. # V004PE050) and standard reactor block with 2 mL reaction vessel (PP-Reactor, 2 mL, with PE Frit, Cat. # V020PE051) (2mL plunger, Cat. # V020ST020) were used for library synthesis in addition to manual synthesis for larger purified peptide stocks. Average resin in tips per 96 was calculated to be approximately 6 mg and reagents were adjusted accordingly. Peptides were cleaved from resin using a mixture of 95% trifluoroacetic acid (Chem-Impex Int’L INC, Cat. # 00289), 2.5 % triisopropylsilane (Sigma Aldrich, Cat # 233781), 2.5% MilliQ water for 2 hours at RT. The cleavage mixture was drained and collected. The resin was then washed with additional cleavage mixture, drained, and collected. TFA was concentrated through evaporation with air stream in a ventilated hood. The residual cleavage mixture was precipitated in diethyl ether and allowed to cool at -20°C for 2 hours. The ether was then removed, and this process was completed three times. After the third ether wash, the residual ether was allowed to evaporate, and the compounds were dissolved in DMSO into 10mM stocks.

**General Procedure A**: Fmoc Deprotection. Peptides were deprotected using a 20% 2-methylpiperidine (TCI, Cat # 203-642-1) in DMF solution. Peptides were deprotected for 4 min at RT three times.

**General Procedure B**: Amide bond coupling for amino acids. The coupling reagent (2-(1H-benzotriazol-1-yl)-1,1,3,3-tetramethyluronium hexafluorophosphate (HBTU) (Peptides International, Cat # KHB-1065-PI) was pre-dissolved in DMF. The coupling reagent 2,4,6-collidine (Alfa Aesar, Cat # A11058) was pre-dissolved in DMF. All coupling reagent amino acids were pre-dissolved in DMF. 2.5 equivalents of amino acid and 2.5 equivalents HBTU, and 4.0 equivalents of 2,4,6-collidine were preactivated and added to the reaction vessel using minimal DMF. The coupling reaction was allowed to react for 1 hour at RT.

**General Procedure C**: Washing Step. The reaction vessel was drained followed by addition of DMF and allowed to sit at RT for 1 min, this process was repeated two more times. The reaction vessel was drained followed by addition of DCM and allowed to sit at RT for 1 min, this process was repeated two more times. The reaction vessel was drained followed by addition of DMF and allowed to sit at RT for 1 min, this process was repeated two more times. Finally, the reaction vessel was drained.

**General Procedure for Macrocycle Synthesis**

Step 1: The resin was allowed to swell for 10 min in DCM and drained. Using general procedure A, the N-terminal Fmoc of the rink amide resin was removed. Then general procedure C was used to wash the resin.

Step 2: Using general procedure B, Fmoc-(L/D)Cys(S-Tmp)-OH was coupled to the resin. Then general procedure C was used to wash the resin. Using general procedure A, the N-terminal Fmoc of the amino acid was removed. Then general procedure C was used to wash the resin.

Step 3: Using general procedure B, either a variable amino acid or Fmoc-Tyr(OSF)OH was coupled to the resin. Then general procedure C was used to wash the resin. Using general procedure A, the N-terminal Fmoc of the amino acid was removed. Then general procedure C was used to wash the resin.

Step 4: Using general procedure B, either a variable amino acid or Fmoc-Tyr(OSF)OH was coupled to the resin. Then general procedure C was used to wash the resin. Using general procedure A, the N-terminal Fmoc of the amino acid was removed. Then general procedure C was used to wash the resin.

Step 5: Amide bond coupling for capping group. The coupling reagent N,N′-Diisopropylcarbodiimide (DIC) (Chem-Impex Int’L INC, Cat. # 00110) was pre-dissolved as a 0.5 M solution in DMF. The capping group bromoacetic acid (Sigma-Aldrich, Cat # B56307) was pre-dissolved as a 0.5 M solution in DMF. The capping group 4-Bromophenylacetic acid (Sigma-Aldrich, Cat # 138673) was pre-dissolved as a 0.5 M solution in DMF. Equal volumes of capping group and DIC were added to the reaction vessel and added to the reaction vessel (0.25 M reaction) for 30 min at RT. The reaction vessel was drained, then the coupling reaction was repeated once more, and finally the reaction vessel drained. Then general procedure C was used to wash the resin.

Step 6: Reaction with primary amine. The primary amine, propargylamine (Chem-Impex Int’L INC, Cat. # 39688) or propylamine (Sigma-Aldrich, Cat # 109819) was pre-dissolved in DMF. 15 equivalents of primary amine were allowed to react with the resin for two hours at RT. Then general procedure C was used to wash the resin.

Step 7: Cysteine deprotection, as described by Postma et. al. N-methylmorpholine (NMM) (Sigma-Aldrich, Cat # M56557) (0.1 M) and DL-dithiothreitol (DTT) (AK Scientific, Cat # J55598) (5%) in DMF were made fresh prior to usage.^[1]^ The deprotection solution was added to the reaction vessel and allowed to react for 5 min at RT and then drained. This was repeated two more times; 3 total. Then general procedure C was used to wash the resin.

Step 8: Bis-electrophile linker cyclization. X equivalents of linker was pre-dissolved in DMF or 40/60 DCM/DMF mixture. Linker and 5 equivalents of 2,4,6-collidine was added to the reaction vessel and allowed to react for two hours with or without shaking. The reaction vessel was then drained. Then 5 equivalents of 2,4,6-collidine was added and allowed to react for six hours with or without shaking. Then resin was then drained. Then general procedure C was used to wash the resin. DCM was added to the reaction vessel and allowed to sit at RT for 1 min and then drained.

Equivalents of linker, solvent mixture, and mixing method used:

Linker 1: (vinylsulfonyl)ethene (Oakwood Chemical, Cat # 008036), 5 equivalents in DMF mixture with/without shaking. Note these failed to cyclize.

Linker 2: 1,2-bis(bromomethyl)benzene (Sigma-Aldrich, Cat # D44405), 6 equivalents in DCM/DMF mixture with shaking.

Linker 3: 2,6-bis(bromomethyl)pyridine (Sigma-Aldrich, Cat # 405426), 2.5 equivalents in DCM/DMF mixture with shaking.

Linker 4: 1,3-dibromopropan-2-one (AK Scientific, Cat # J97961), 2.5 equivalents in DMF mixture with/without shaking.

Linker 5: 1,4-bis(bromomethyl)benzene (Sigma-Aldrich, Cat # D44804), 2.5 equivalents in DCM/DMF mixture with shaking.

Step 9: Peptides were then cleaved, precipitated, and dissolved in DMSO according to the methods outlined in solid phase peptide synthesis.

**Library Composition**

*96 well plates (P) were divided into four parts as shown below.*

PX:1-24 - L cysteine, OSF on Position 2, Amino Acids 1-24

PX: 25-48 - D cysteine, OSF on Position 2, Amino Acids 1-24

PX: 49-72 - L cysteine, OSF on Position 3, Amino Acids 1-24

PX:73-96 - D cysteine, OSF on Position 3, Amino Acids 1-24

*7 libraries were synthesized in 96 well-plates with 5 forming macrocycles.*

P1: Bis-electrophile linker: 1, capping group: bromoacetic acid (note: failed to cyclize)

P2: Bis-electrophile linker: 2, capping group: bromoacetic acid

P3: Bis-electrophile linker: 3, capping group: bromoacetic acid

P4: Bis-electrophile linker: 4, capping group: bromoacetic acid

P5: Bis-electrophile linker: 5, capping group: bromoacetic acid

P6: Bis-electrophile linker: 1, capping group: 4-Bromophenylacetic acid (note: failed to cyclize)

P7: Bis-electrophile linker: 2, capping group: 4-Bromophenylacetic acid

*24 amino acids were used (18 natural, 6 unnatural*), purchased from CreoSalus.*

1 Fmoc-Ala-OH

2 *Fmoc-Hse(Bzl)-OH (N-alpha-(9-Fluorenylmethyloxycarbonyl)-O-benzyl-L-homoserine)

3 Fmoc-Asp(OtBu)-OH

4 Fmoc-Glu(OtBu)-OH

5 Fmoc-Phe-OH

6 Fmoc-Gly-OH

7 Fmoc-His(Trt)-OH

8 Fmoc-Ise-OH

9 Fmoc-Lys(Boc)-OH

10 Fmoc-Leu-OH

11 *Fmoc-Nle-OH (N-α-Fmoc-L-norleucine)

12 Fmoc-Asn(Trt)-OH

13 Fmoc-Pro-OH

14 Fmoc-Gln(Trt)-OH

15 Fmoc-Arg(Pbf)-OH

16 Fmoc-Ser(Tbu)-OH

17 Fmoc-Thr(Tbu)-OH

18 Fmoc-Val-OH

19 Fmoc-Trp(Boc)-OH

20 Fmoc-Tyr(tBu)-OH

21 *Fmoc-1-Nal-OH (Fmoc-3-(1-naphthyl)-L-alanine)

22 *Fmoc-Cha-OH (N-α-Fmoc-β-cyclohexyl-L-alanine)

23 *Fmoc-Ala-4Pip(Boc)-OH (2-N-Fmoc-amino-3-(4-N-boc-piperidinyl)propionic acid)

24 *Fmoc-His(3-Bom)-OH (N-(fluorenylmethoxycarbonyl)-3-benzoxymethyl-L-histidine)

**Small Molecule Synthesis**

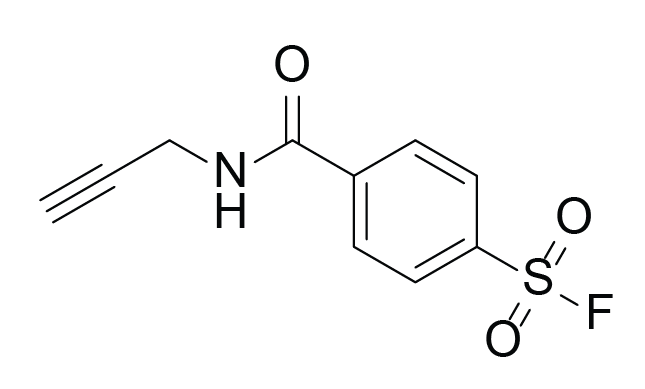

**Sulfonylfluoride Alkyne (SF-alkyne),** 4-(prop-2-yn-1-ylcarbamoyl)benzenesulfonyl fluoride: Was synthesized as previously reported in Hahm et. al.^[2]^

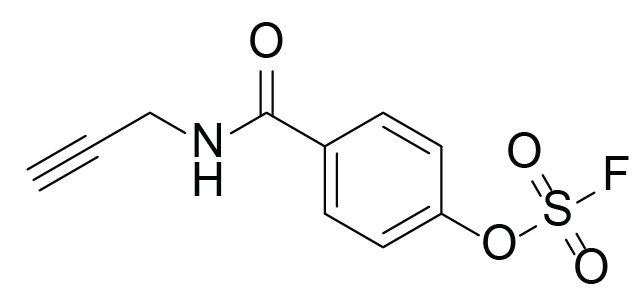

**Fluorosulfate Alkyne (OSF-alkyne),** 4-hydroxy-N-(prop-2-yn-1-yl)benzamide was synthesized as previously described in Gopin et. al.^[3]^ To a vial containing the 4-hydroxy-N-(prop-2-yn-1-yl)benzamide (55 mg, 314umol, 1 equiv.), AISF (118 mg, 377umol,1.2 equiv.) and stir bar was added tetrahydrofuran (1.5 mL) followed by 1,8-diazabicyclo[5.4.0]undec-7-ene (103uL, 691umol, 2.2 equiv.) over a period of 30 seconds. The reaction mixture was stirred at room temperature for 10 minutes and then diluted with ethyl acetate and washed with either 1 N HCl (2x) and brine (1x). The combined organic fraction was dried with anhydrous sodium sulfate and concentrated under reduced pressure. The crude residue was purified by silica gel flash chromatography 0-100% hexanes/ethyl acetate gradient. The purified fractions were collected and concentrated to produce a white powder (61.4 mg, 76% yield).

^1^H NMR (400 MHz, Chloroform-*d*) δ 7.96 – 7.85 (m, 2H), 7.48 – 7.39 (m, 2H), 6.31 (s, 1H), 4.26 (d, *J* = 2.6 Hz, 2H), 2.31 (t, *J* = 2.5 Hz, 1H).

^19^F NMR (376 MHz, Chloroform-*d*) δ 38.55.

MS(ESI)^+^ calculated for C_11_H_8_FNO_4_S^1+^: 258.0192, found 258.3

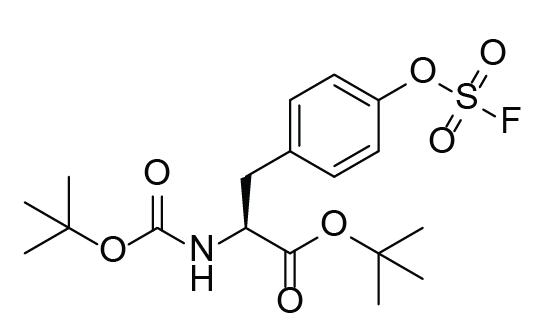

**Intermediate 1,** tert-butyl(S)-2-((tert-butoxycarbonyl)amino)-3-(4-((fluorosulfonyl)oxy)phenyl)propanoate: To a vial of tert-butyl (tert-butoxycarbonyl)-L-tyrosinate (675 mg, 2 mmol, 1 equiv.) and AISF (754 mg, 2.4 mmol,1.2 equiv.), was added tetrahydrofuran (9 mL) followed by 1,8-diazabicyclo[5.4.0]undec-7-ene (657uL, 4.4 mmol, 2.2 equiv.) over a period of 30 seconds. The reaction mixture was stirred at room temperature for 10 minutes and then diluted with ethyl acetate and washed with either 1 N HCl (2x) and brine (1x). The combined organic fraction was dried with anhydrous sodium sulfate and concentrated under reduced pressure. The crude residue was purified by silica gel flash chromatography 0-100% hexanes/ethyl acetate gradient. The purified fractions were collected and concentrated to produce a white powder (789 mg, 94% yield).

^1^H NMR (400 MHz, Chloroform-*d*) δ 7.96 – 7.85 (m, 2H), 7.48 – 7.39 (m, 2H), 6.31 (s, 1H), 4.26 (d, *J* = 2.6 Hz, 2H), 2.31 (t, *J* = 2.5 Hz, 1H).

^19^F NMR (376 MHz, Chloroform-*d*) δ 38.55.

MS(ESI)^+^ calculated for C_18_H_26_FNO_7_SNa^1+^: 442.1307, found 442.3

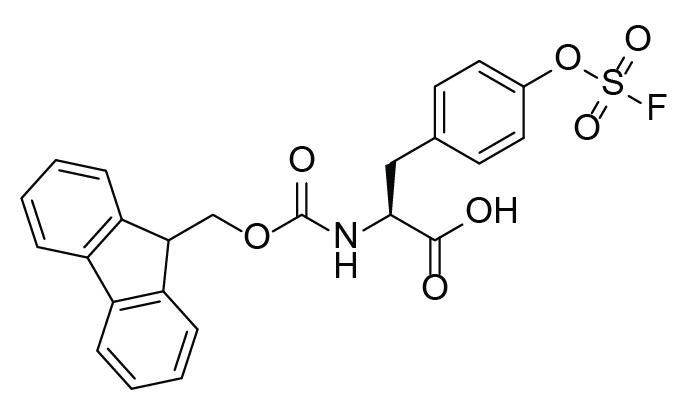

**Fmoc-Tyr(OSF)-OH,** (S)-2-((((9H-fluoren-9-yl)methoxy)carbonyl)amino)-3-(4-((fluorosulfonyl)oxy)phenyl)propanoic acid: To a flask with stir bar, Intermediate 1 (4.8 g, 11.4 mmol, 1 equiv.) was dissolved in 50/50 DCM/ trifluoroacetic acid and stirred for one hour at RT. The mixture was then concentrated under reduced pressure. The resulting residue was dissolved in 50/50 dioxane/ water. Sodium bicarbonate (approximately 1.9 g, 22.4 mmol, 2 equiv.) was added until the mixture reached pH 8.5. Then Fmoc-Succinimide (3.86 g, 11.4 mmol, 1 equiv.) was added and allowed to stir at RT for one hour. The reaction mixture was then diluted with ethyl acetate, acidified, and washed with 1 N HCl (2x) and brine (1x). The combined organic fraction was dried with anhydrous sodium sulfate and concentrated under reduced pressure. The crude residue was purified by silica gel flash chromatography 0-5% DCM/MeOH. The purified fractions were collected and concentrated to produce a white powder (2.794 g, 50.3% yield).

^1^H NMR (500 MHz, Methanol-*d*_4_) δ 7.79 (d, *J* = 7.6 Hz, 2H), 7.61 (t, *J* = 6.7 Hz, 2H), 7.44 – 7.25 (m, 8H), 4.44 (dt, *J* = 10.0, 5.7 Hz, 1H), 4.29 (qd, *J* = 10.6, 7.0 Hz, 2H), 4.16 (t, *J* = 7.0 Hz, 1H), 3.27 (d, *J* = 4.7 Hz, 1H), 3.00 (dd, *J* = 14.0, 9.8 Hz, 1H).

^19^F NMR (376 MHz, Methanol-*d*_4_) δ 35.19. C24H20FNO7S

MS(ESI)^+^ calculated for C_24_H_21_FNO_7_S: 486.0978, found 486.2

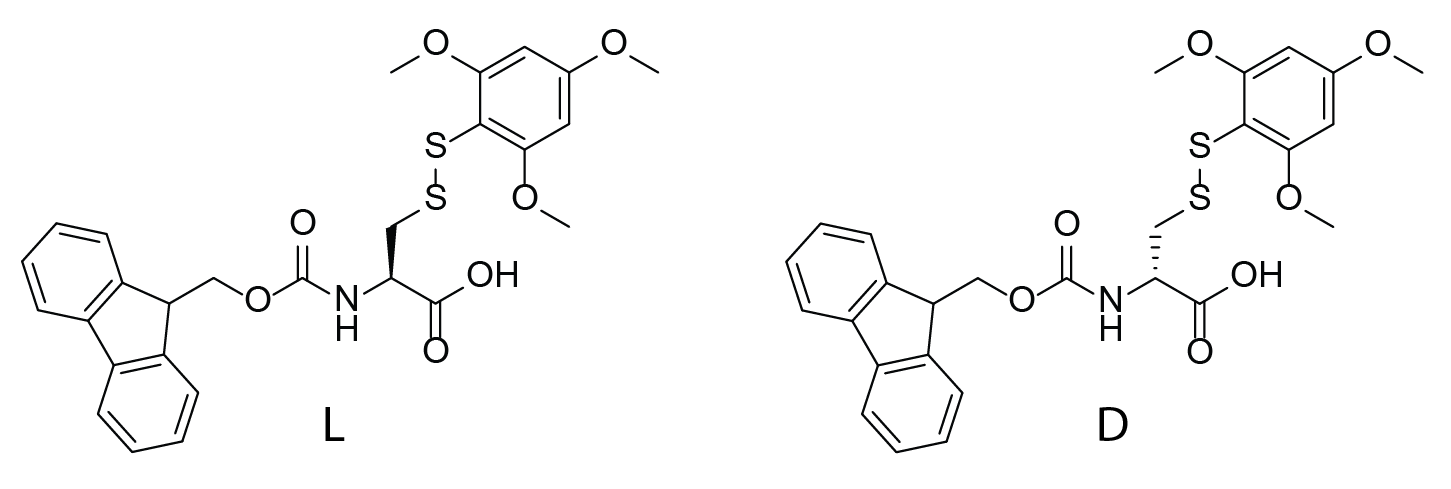

**Fmoc-(L/D)Cys(S-Tmp)-OH,** Fmoc-Cys(STmp)-OH, N-α-Fmoc-S-2,4,6-trimethoxyphenylthio-(L/D)-cysteine: Was synthesized as previously reported in Postma et. al.^[1]^

##### **HPLC Purity Analysis for Library Synthesis**

*Please see the Supplementary Spectra for TIC and UV data.*

*Purified Probes*

| **Probe Name** | **Expected Mass [M+1]^+1^** | **Found Mass** |
| --- | --- | --- |
| P1-52 | 708.1435 | 708.3 |
| P1-52-c | 712.1748 | 712.3 |
| P4-30 | 572.1240 | 572.4 |
| P4-30-c | 576.1553 | 576.2 |
| P6-7 | 792.1911 | 792.2 |
| P6-7-c | 796.2224 | 796.3 |
| P2-9 | 691.2339 | 691.5 |
| P2-9-c | 695.2652 | 695.4 |
| P2-12 | 677.1819 | 676.8 |
| P2-12-c | 681.2132 | 681.2 |
| P2-15 | 719.2401 | 719.5 |
| P2-15-c | 723.2714 | 723.5 |

#### **Biology methods**

**Cell culture and preparation of lysates**

HEK 293 cells were maintained in DMEM media supplemented with 10% fetal bovine serum (FBS), non-essential amino acids and penicillin streptomycin at 37 °C under 5% CO_2_ atmosphere. Cells were collected by trypsinization or scraping, washed with PBS, centrifuged at 1,500 x g for five minutes at 4 °C and resuspended in PBS (PBS with 0.5% triton for rapid screen). Cell lysates were obtained by sonication and protein concentration was determined.

**Gel-based *in-vitro* rapid screening**

HEK293 lysate (2 mg/mL, 25 µL) was treated with indicated compound at indicated concentration or DMSO for two hours at 37 °C. Click chemistry was initiated by the addition of TMR azide (Sigma-Aldrich, 10mM stock in DMSO, 20 μM final), copper(II) sulfate (50 mM fresh stock, 1mM final), 2-(4-((Bis((1-(tert-butyl)-1H-1,2,3-triazol-4-yl)methyl)amino)methyl)-1H-1,2,3-triazol-1-yl)acetic acid (BTTAA, Click Chemistry Tools, 100 mM stock in DMSO, 5mM final), and Sodium ascorbate (Sigma-Aldrich, 300 mM stock, 15mM final) to the lysate and incubated in the dark for 30 min at 37 °C SDS-PAGE reducing loading buffer (4x) was added and proteins were separated using an 12% SDS-PAGE gel. Gels were visualized using a GE Typhoon FLA 9000, then stained using Coomassie.

**Gel-based *in-vitro* labeling with purified probes**

HEK 293 lysate (2 mg/mL, 25 µL) was treated with indicated compound at indicated concentration or DMSO for two hours at 37 °C followed by indicated probe at 10 µM or DMSO for two hours at 37 °C. Click chemistry was initiated by the addition of TMR azide (Sigma-Aldrich, 50 µM, 25x stock in DMSO), TCEP (1 mM, fresh 50x stock in water), tris[(1-benzyl-1*H*-1,2,3-triazol-4-yl)methyl]amine (TBTA, Sigma-Aldrich, 100 µM, 16x stock in DMSO:*t*BuOH 1:4), and copper(II) sulfate (1 mM, 50x stock in water) to the lysate and incubated in the dark for one hour at r.t. SDS-PAGE reducing loading buffer (4x) was added and proteins were separated using an 11% SDS-PAGE gel. Gels were visualized using a Sapphire Biomolecular Imager (Azure Biosystems), then stained using Coomassie.

**OSF-alkyne and SF-alkyne capture-and-release experiment**

HEK 293 lysate (5 mg/mL, 2.5 mL) was treated with 100 µM OSF-alkyne or 100 µM SF-alkyne for two hours at 37 °C. Desthiobiotin azide (150 µM, 50x stock in DMSO), TCEP (1 mM, 50x fresh stock in water), TBTA (100 µM, 16x stock in DMSO:*t*Butanol 1:4), and copper(II) sulfate (1 mM, 50x stock in water) were added to the proteome and left to react for one hour at r.t.^[4]^ Protein was precipitated by adding MeOH (4 vol.), CHCl_3_ (1 vol.) and water (3 vol.) to the reaction mixture and the turbid mixture was centrifuged for 10 min at 20,000 x g at 4 °C yielding a protein layer between the aqueous and organic layers. The protein layer was isolated, washed with 3 vol. of MeOH, dried, solubilized in 6 M urea in 50 mM NH_4_HCO_3_ via sonication, reduced with 10 mM neutralized TCEP (20x fresh stock in water) for 30 min. at r.t., and alkylated with 25 mM iodoacetamide (400 mM fresh stock in water) for 30 min. at r.t. in the dark. Sample was diluted to 2 M urea with 50 mM NH_4_HCO_3_ and digested with trypsin (Thermo Scientific, 100 μL of 0.5 μg/μL) overnight in the presence of 1 mM CaCl_2_. Sample was diluted to 1 M urea with 50 mM NH_4_HCO_3_ and 500 µL of streptavidin agarose beads were added and the mixture was rotated for two hours at r.t. Beads were washed with PBS (3x 10 mL) and water (3x 10 mL). Beads-bound peptides were released with 50% acetonitrile 0.1% FA for 3 min (three times). Eluate was lyophilized, desalted over a self-packed C18 spin column and dried. Samples were analyzed by LC-MS/MS (see below). The MS data was processed with Proteome Discoverer (see below).

**Gel-based *in-vitro* competitive labeling with purified probes**

HEK 293 lysate (2 mg/mL, 25 µL) was treated with indicated compound at indicated concentration or DMSO for two hours at 37 °C followed by indicated probe at 10 µM or DMSO for two hours at 37 °C. Alternatively, compound and probe were co-treated for four hours at 37 °C. Click chemistry was initiated by the addition of TMR azide (Sigma-Aldrich, 50 µM, 25x stock in DMSO), TCEP (1 mM, fresh 50x stock in water), tris[(1-benzyl-1H-1,2,3-triazol-4-yl)methyl]amine (TBTA, Sigma-Aldrich, 100 µM, 16x stock in DMSO:tBuOH 1:4), and copper(II) sulfate (1 mM, 50x stock in water) to the lysate and incubated in the dark for one hour at r.t. SDS-PAGE reducing loading buffer (4x) was added and proteins were separated using an 11% SDS-PAGE gel. Gels were visualized using a Sapphire Biomolecular Imager (Azure Biosystems), then stained using Coomassie.

**P4:30 proteomics**

HEK 293 lysate (2 mg/mL, 1 mL) was co-treated with DMSO, 10 µM, 30 µM, or 100 µM of P4:30-c and 10 µM P4:30 for four hours at 37 °C or with DMSO and 10 µM P2:9, for four hours at 37 °C. Biotin azide (Sigma-Aldrich, 20 µM, 50x stock in DMSO), tris(2-carboxyethyl)phosphine hydrochloride (TCEP) (1 mM, 50x fresh stock in water), tris[(1-benzyl-1H-1,2,3-triazol-4-yl)methyl]amine (TBTA) (100 µM, 16x stock in DMSO:tButanol 1:4), and copper(II) sulfate (1 mM, 50x stock in water) were added and left to incubate for one hour at r.t. Protein was precipitated by adding 4 vol. of MeOH, 1 vol. of CHCl_3_ and 3 vol. of water to the reaction mixture and the turbid mixture was centrifuged for five min. at 14’000 x g at 4 °C yielding a protein layer between the aqueous and organic layers. The protein layer was isolated, dried and solubilized in 2% SDS in PBS via sonication. Tube was centrifuged at 4’700 x g for five min. and soluble fraction was transferred to a new tube. PBS was added to give a final SDS concentration of 0.2%. 120 µL of streptavidin agarose beads (ProteoChem) were added and the mixture was rotated for overnight at r.t. Beads were washed with 1% SDS in PBS (1x 10 mL), PBS (3x 10 mL), and water (3x 10 mL). Beads were resuspended in 6 M urea in PBS (500 μL), reduced with 10 mM neutralized TCEP (20x fresh stock in water) for 30 min. at r.t., and alkylated with 25 mM iodoacetamide (400 mM fresh stock in water) for 30 min. at r.t. in the dark. Beads were pelleted by centrifugation (1’400 x g, two min.) and resuspended in 150 μL of 2 M urea, 1 mM CaCl_2_ (100x stock in water) and trypsin (Thermo Scientific, 1 μL of 0.5 μg/μL) in 50 mM NH_4_HCO_3_. The digestion was performed for 6 hours at 37 °C. Samples were acidified to a final concentration of 5% acetic acid, desalted over a self-packed C18 spin column and dried. Samples were analyzed by LC-MS/MS (see below) and the MS data was processed with MaxQuant (see below).

**Purified protein labeling with P4:30**

Purified human Translin (50 ng) or 20S Proteasome (1500 ng) was treated with P4:30, OSF-alkyne, SF-alkyne, and at indicated concentration or DMSO for two hours at 37 °C. Click chemistry was initiated by the addition of TMR azide (Sigma-Aldrich, 10mM stock in DMSO, 20μM final), copper(II) sulfate (50 mM fresh stock, 1mM final), 2-(4-((Bis((1-(tert-butyl)-1H-1,2,3-triazol-4-yl)methyl)amino)methyl)-1H-1,2,3-triazol-1-yl)acetic acid (BTTAA, Click Chemistry Tools, 100 mM stock in DMSO, 5mM final), and Sodium ascorbate (Sigma-Aldrich, 300 mM stock, 15mM final) to the lysate and incubated in the dark for 30 min at 37 °C. SDS-PAGE reducing loading buffer (4x) was added and proteins were separated using an 12% SDS-PAGE gel. Gels were visualized using a GE Typhoon FLA 9000, then stained using Coomassie.

**LC-MS/MS analysis**

Peptides were resuspended in water with 0.1% formic acid (FA) and analyzed using Proxeon EASY-nLC 1200 nano-UHPLC coupled to QExactive HF-X Quadrupole-Orbitrap mass spectrometer (Thermo Scientific). The chromatography column consisted of a 50 cm long, 75 μm i.d. microcapillary capped by a 5 μm tip and packed with ReproSil-Pur 120 C18-AQ 2.4 μm beads (Dr. Maisch GmbH). LC solvents were 0.1% FA in H_2_O (Buffer A) and 0.1% FA in MeCN (Buffer B). Peptides were eluted into the mass spectrometer at a flow rate of 300 nL/min. over a 90 minutes linear gradient (5-35% Buffer B) at 65 °C. Data was acquired in data-dependent mode (top-20, NCE 28, R = 7,500) after full MS scan (R = 60,000, m/z 400-1,300). Dynamic exclusion was set to 10 s, peptide match to prefer and isotope exclusion was enabled.

**Proteome Discoverer analysis**

The MS data was analyzed with Proteome Discoverer using the Sequest HT algorithm and searched against the human proteome (Uniprot).^[5]^ For amino acid selectivity, OSF-desthiobiotin and SF-desthiobiotin were searched as dynamic modification on either Arg, Cys, Asp, Glu, His, Lys, Ser, Thr, Tyr together with methionines oxidation and carbamidomethylation of Cys. The minimum peptide length was set to six, maximum precursor mass to 5,000 Da, precursor mass tolerance to 10 ppm and fragment mass tolerance to 0.02 Da. Only peptide with PEP value ≤1% were considered.

**MaxQuant analysis**

The MS data was analyzed with MaxQuant and searched against the human proteome (Uniprot) and a common list of contaminants (included in MaxQuant).^[6]^ The first peptide search tolerance was set at 20 ppm, 10 ppm was used for the main peptide search and fragment mass tolerance was set to 0.02 Da. The false discovery rate for peptides, proteins and sites identification was set to 1%. The minimum peptide length was set to six amino acids and peptide re-quantification was enabled. The minimal number of peptides per protein was set to two. Methionine oxidation was searched as a variable modification and carbamidomethylation of cysteines was searched as a fixed modification.
