## Supplemental Spectra for "Solid Phase Synthesis of Fluorosulfate Containing Macrocycles for Chemoproteomic Workflows"

Printing Time: 2:53:06 PM
Printing Date: Tuesday, January 31, 2023
Service Engineer Chris Maier

Post PM
*Stanford U

Polarity/Scan Type: Positive Q1 MS
Sample Name: F7-49-pep1-fr35
*API 3000 SN D7640012

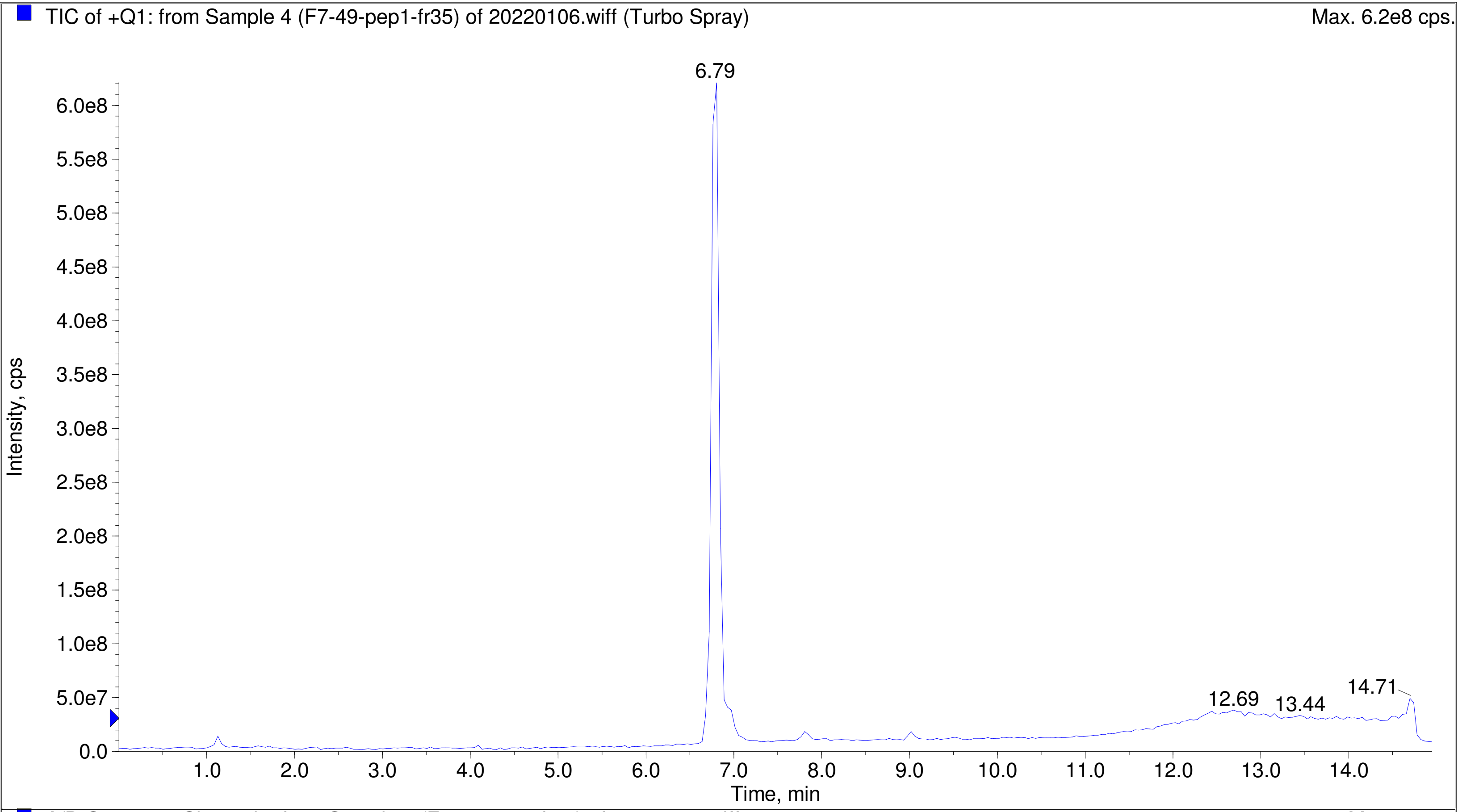


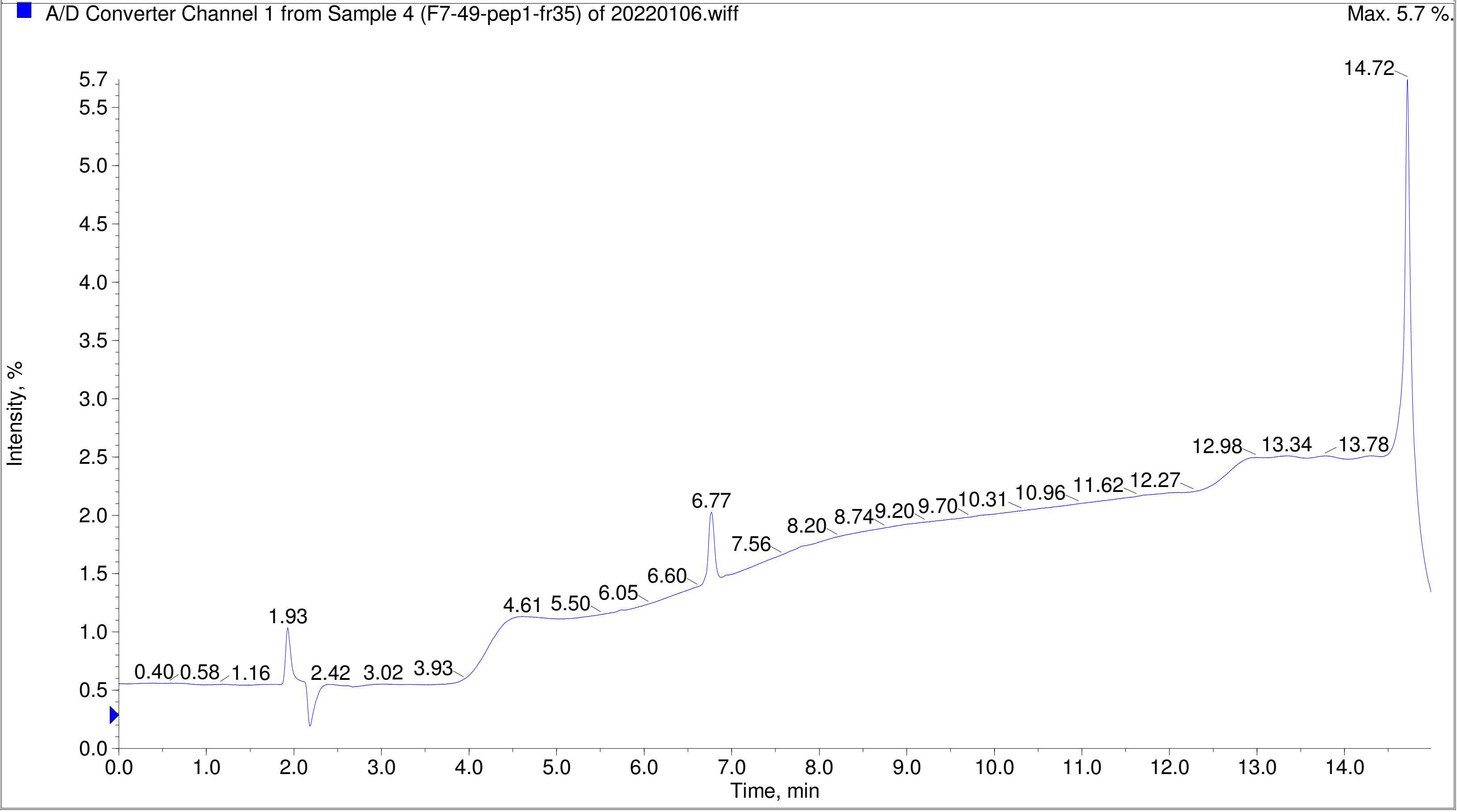

Printing Time: 2:55:13 PM Post PM Polarity/Scan Type: Positive Q1 MS
Printing Date: Tuesday, January 31, 2023 Sample Name: F7-49-pep7-fr33-36
Service Engineer Chris Maier *Stanford U *API 3000 SN D7640012

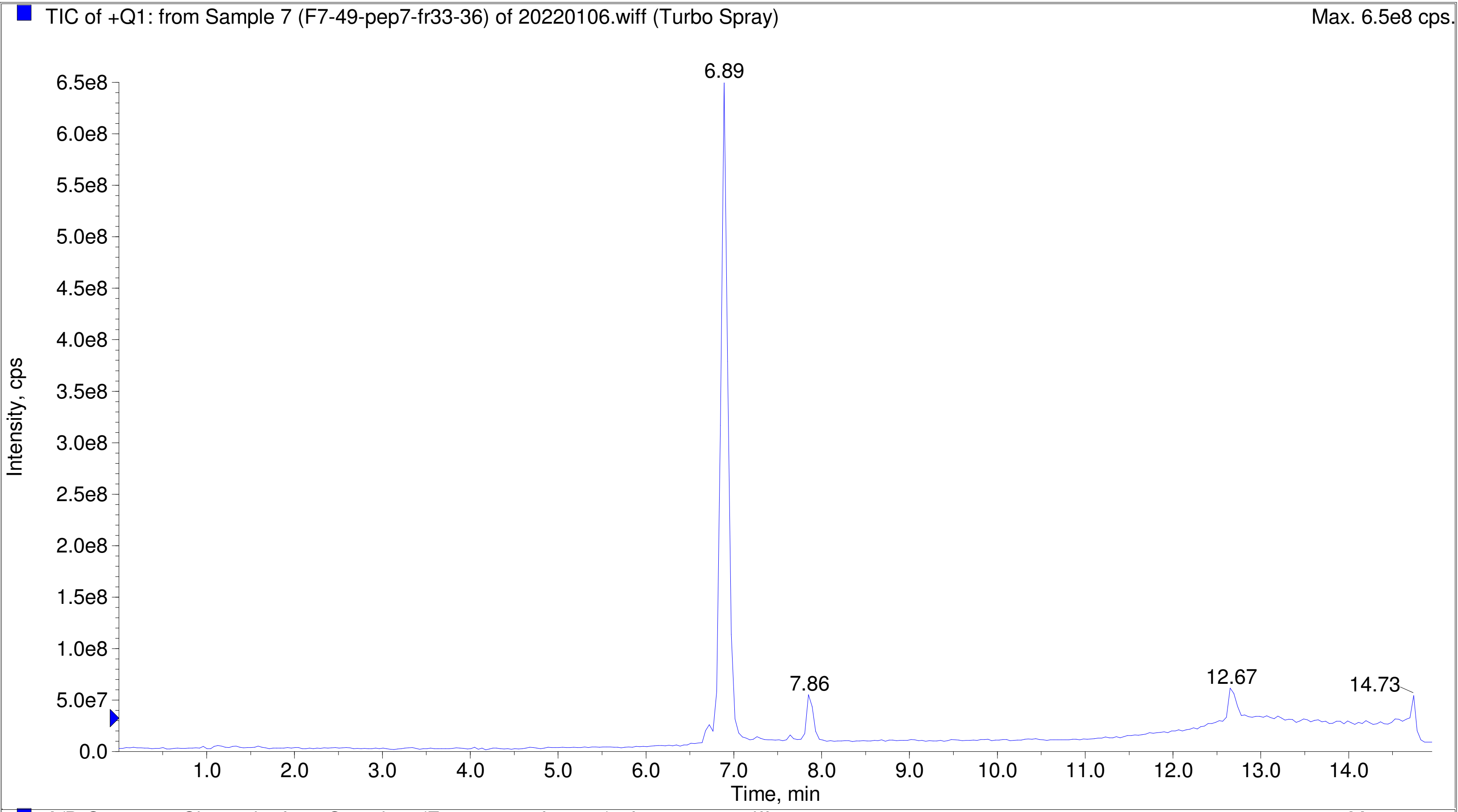


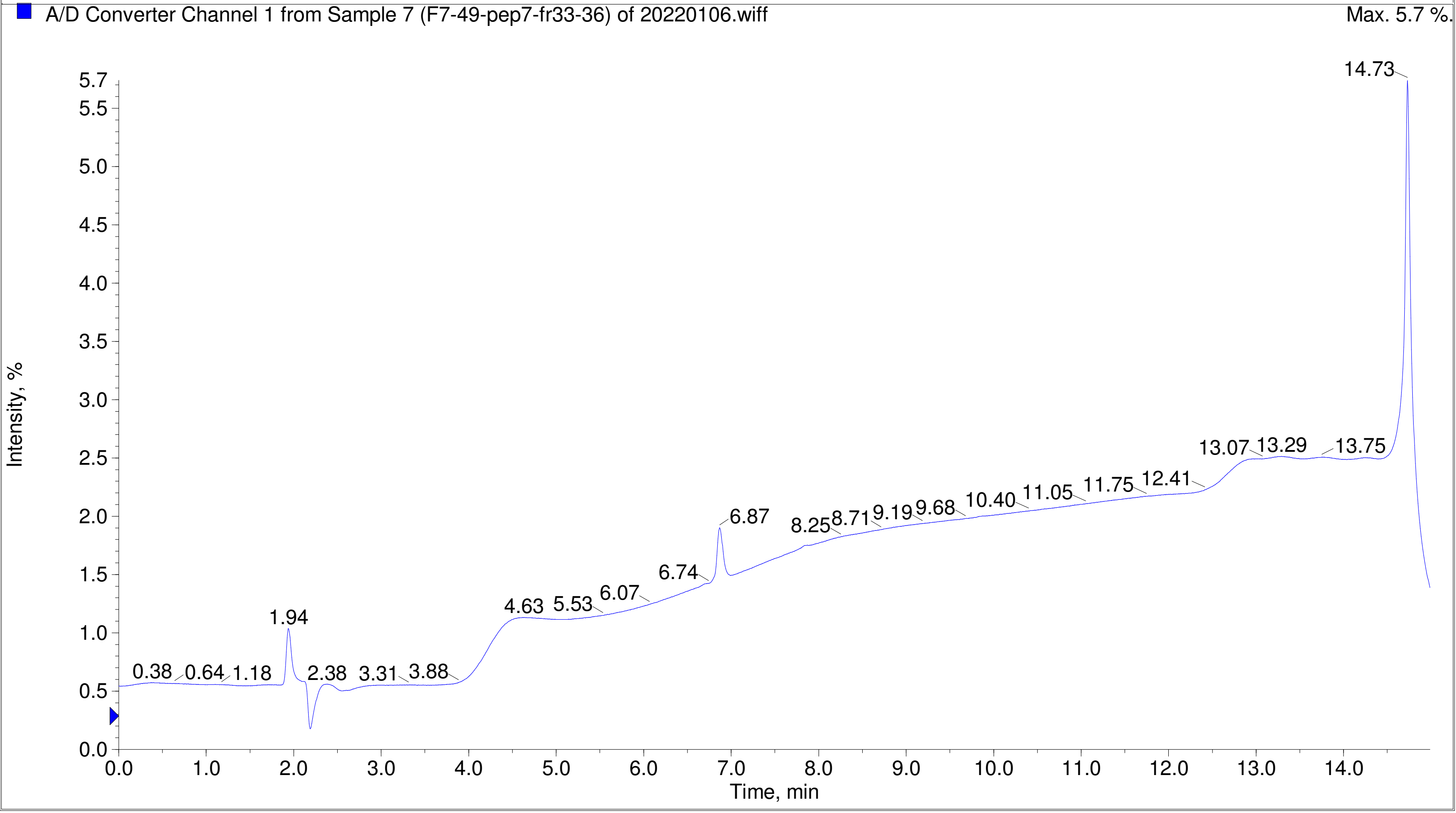

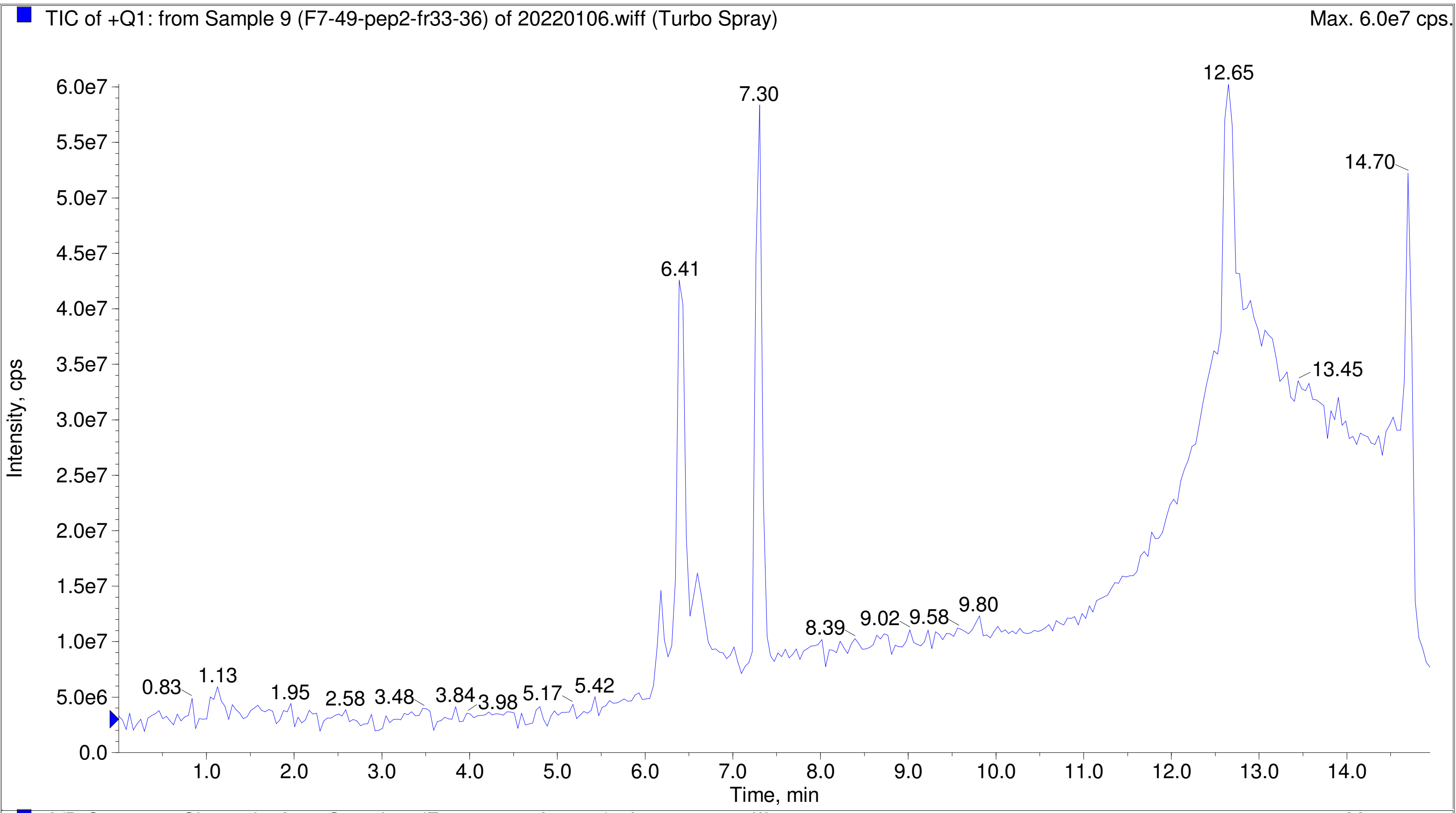
TIC of +Q1: from Sample 9 (F7-49-pep2-fr33-36) of 20220106.wiff (Turbo Spray)
Max. 6.0e7 cps.
Intensity, cps
Time, min
0.83
1.13
1.95
2.58
3.48
3.84
3.98
5.17
5.42
6.41
7.30
8.39
9.02
9.58
9.80
12.65
13.45
14.70

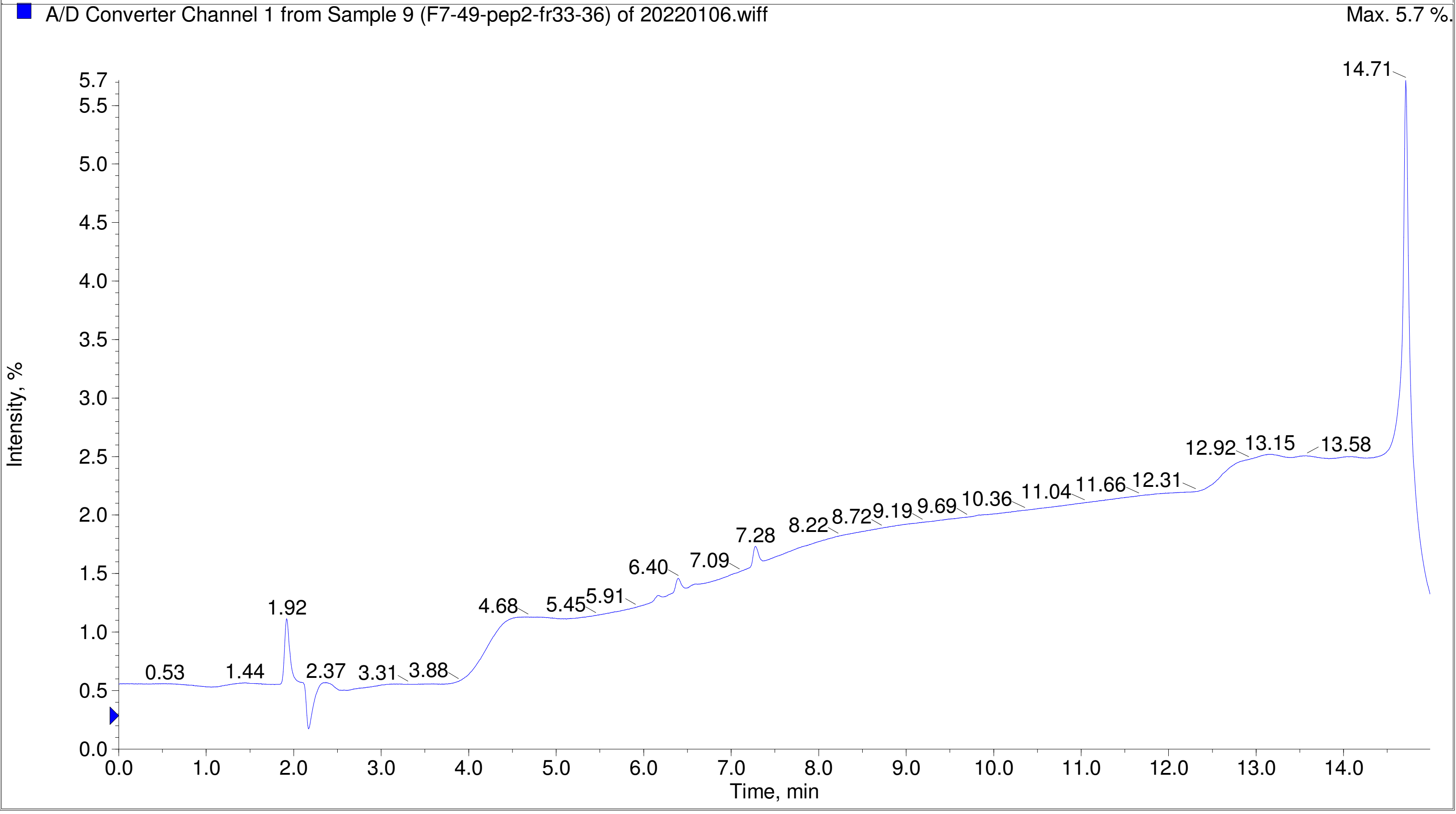
A/D Converter Channel 1 from Sample 9 (F7-49-pep2-fr33-36) of 20220106.wiff
Max. 5.7 %.
Intensity, %
Time, min
0.53
1.44
1.92
2.37
3.31
3.88
4.68
5.45
5.91
6.40
7.09
7.28
8.22
8.72
9.19
9.69
10.36
11.04
11.66
12.31
12.92
13.15
13.58
14.71

Printing Time: 2:58:47 PM | Post PM | Polarity/Scan Type: Positive Q1 MS
Printing Date: Tuesday, January 31, 2023 | | Sample Name: F7-49-pep8-fr33-36
Service Engineer Chris Maier | *Stanford U | *API 3000 SN D7640012

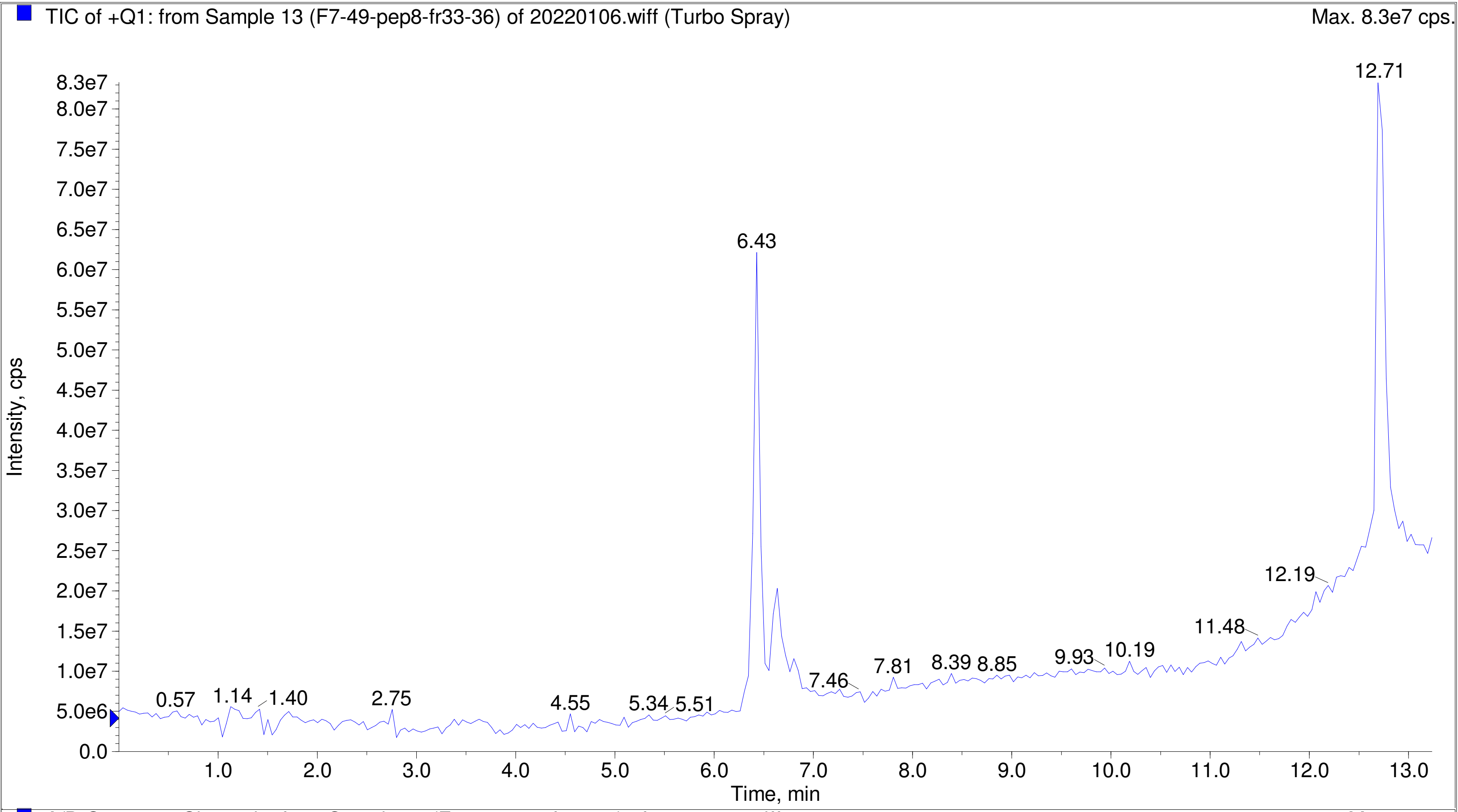


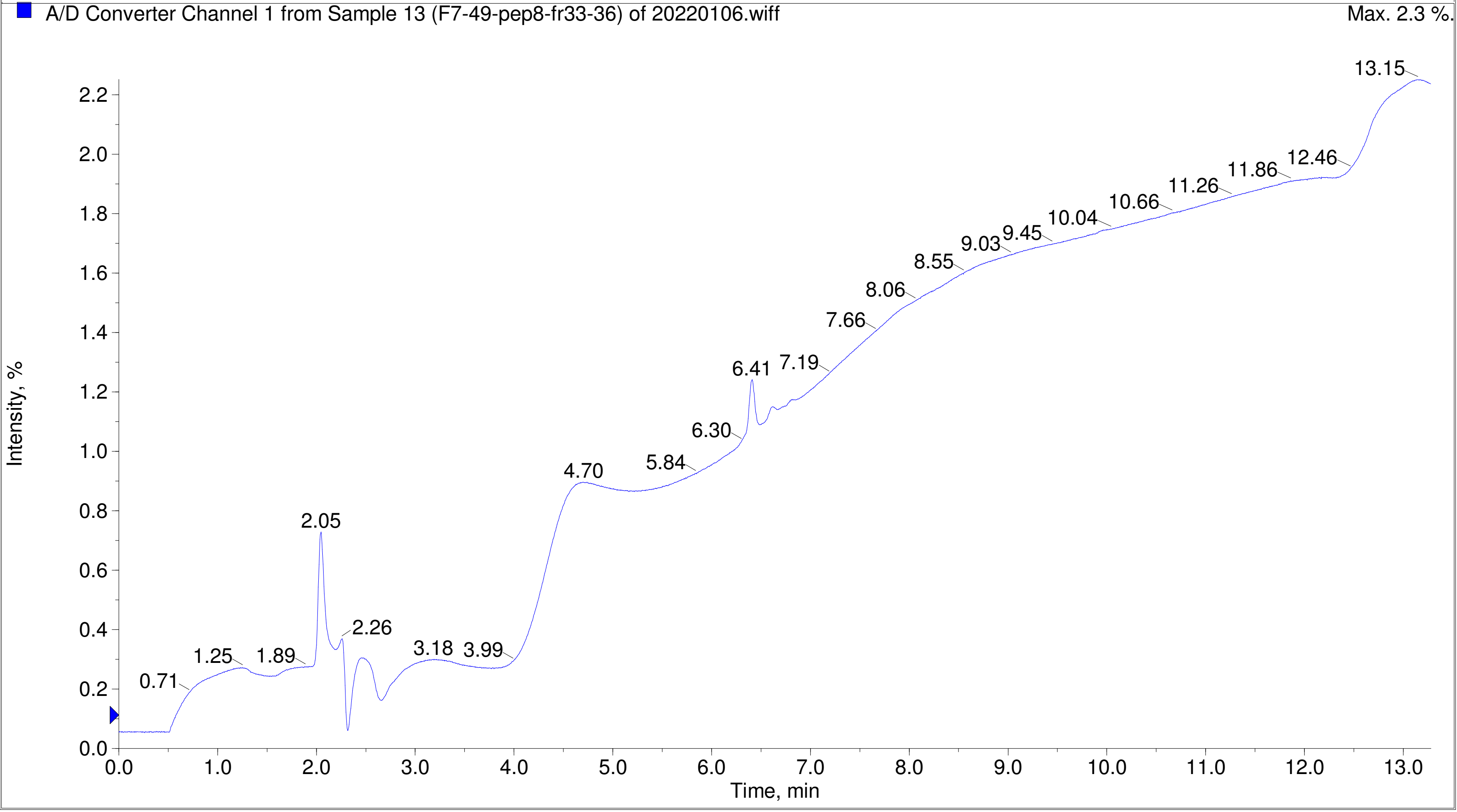

Printing Time: 3:12:47 PM
Printing Date: Tuesday, January 31, 2023
Service Engineer Chris Maier

Post PM
*Stanford U

Polarity/Scan Type: Positive Q1 MS
Sample Name: F7-49-ag-pep3-fr37
*API 3000 SN D7640012

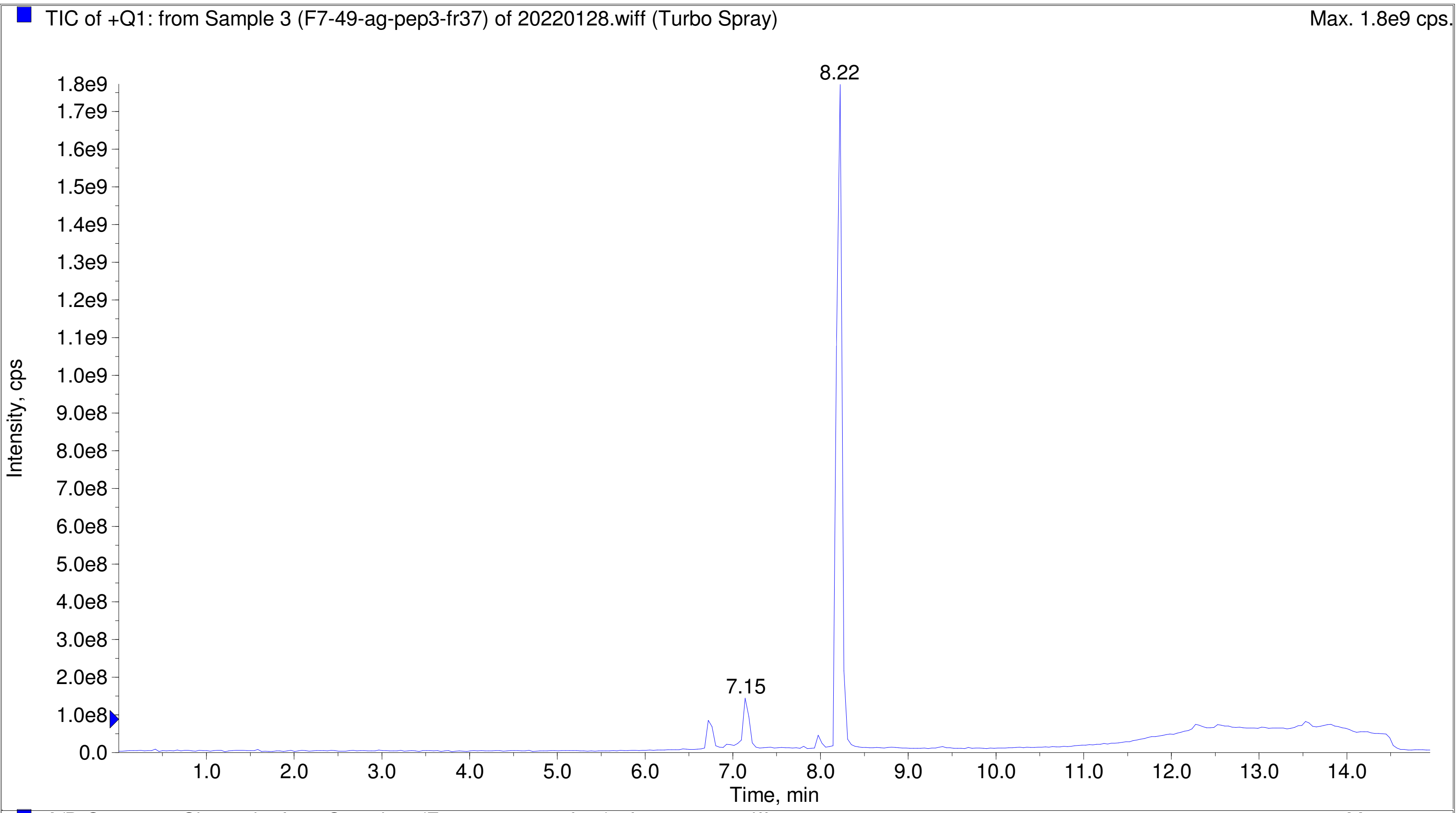


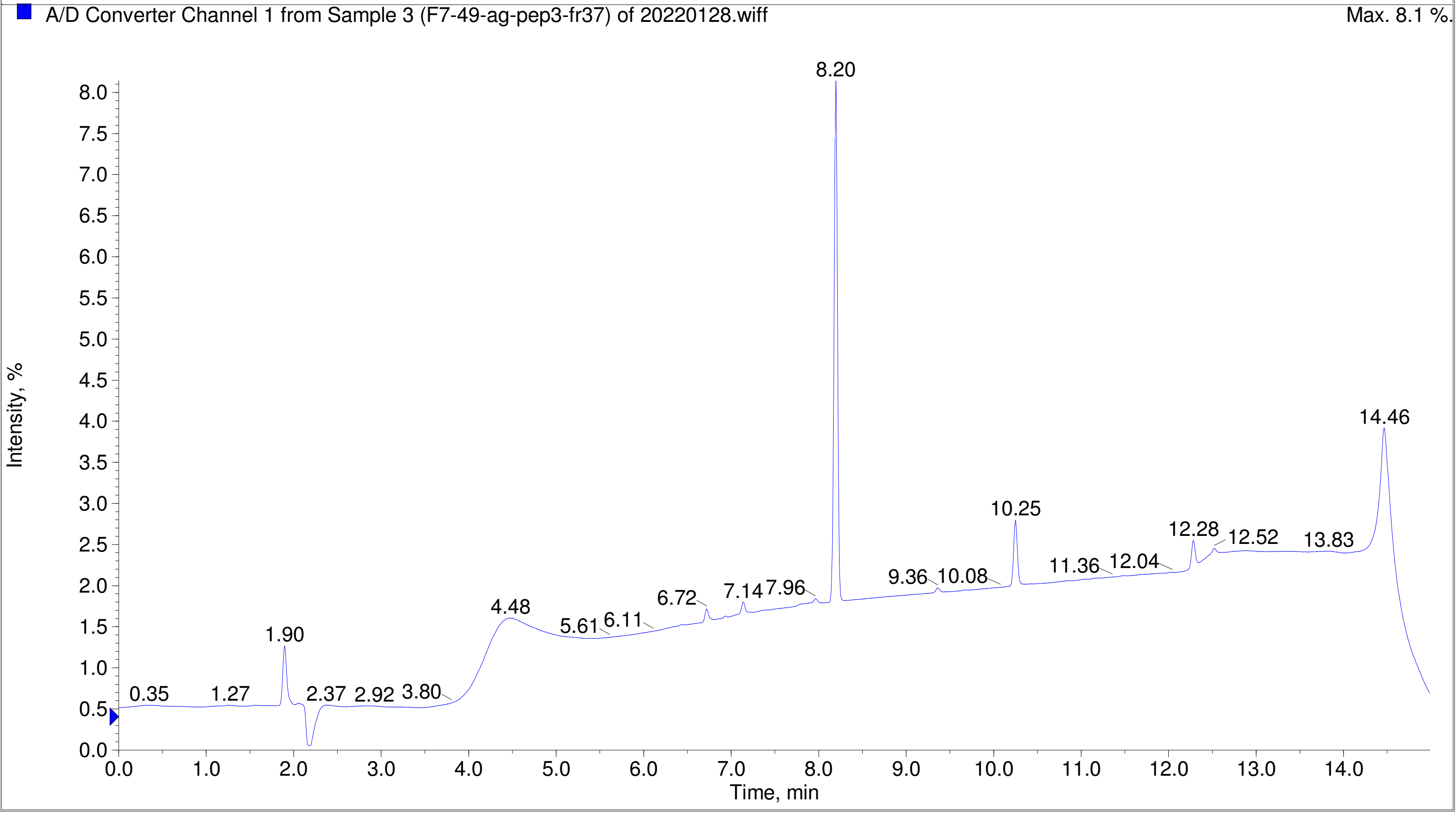

Printing Time: 3:13:57 PM
Printing Date: Tuesday, January 31, 2023
Service Engineer Chris Maier

Post PM
*Stanford U

Polarity/Scan Type: Positive Q1 MS
Sample Name: F7-49-ag-pep9-fr37
*API 3000 SN D7640012

TIC of +Q1: from Sample 4 (F7-49-ag-pep9-fr37) of 20220128.wiff (Turbo Spray) Max. 1.1e9 cps.

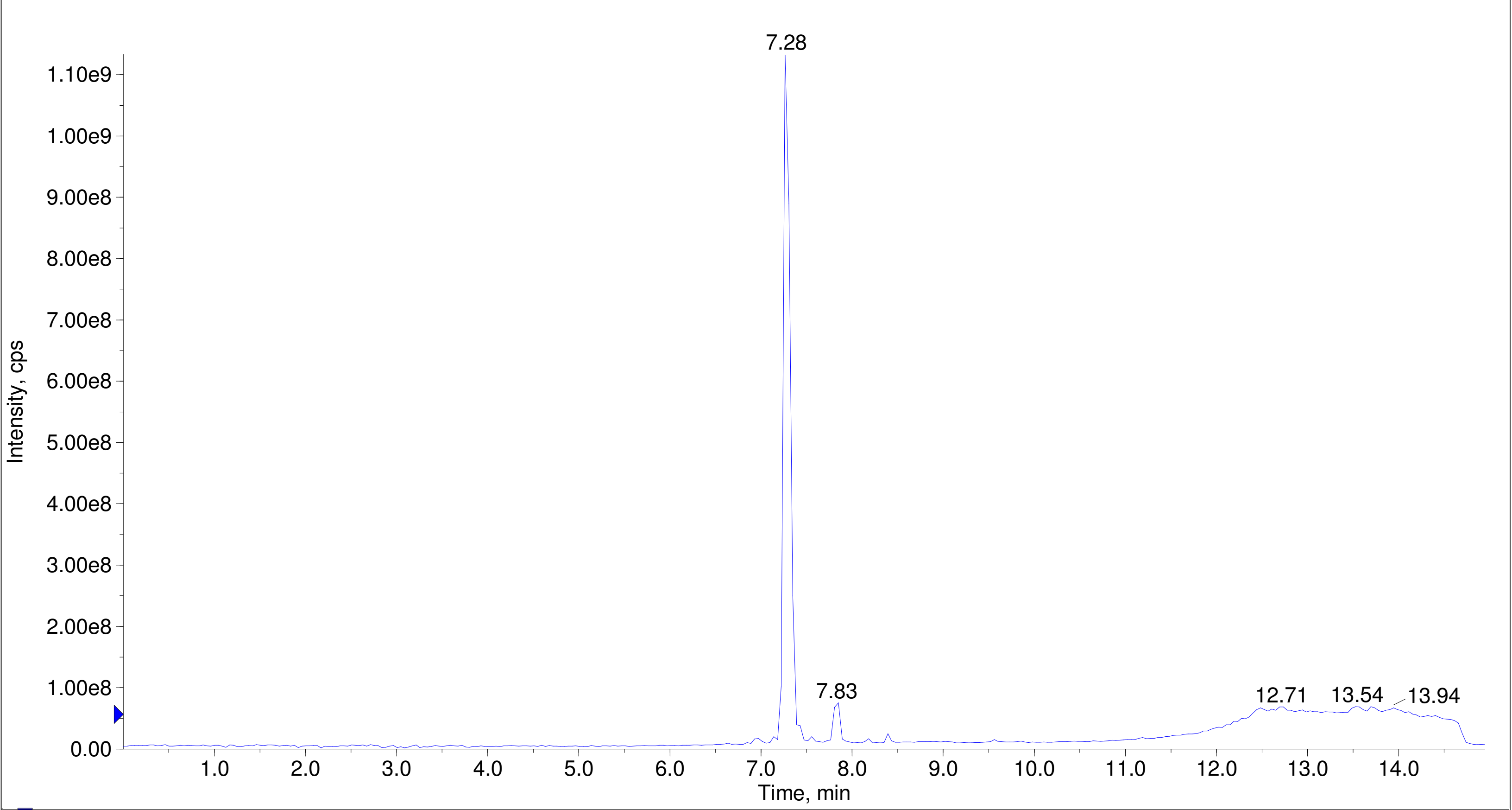


A/D Converter Channel 1 from Sample 4 (F7-49-ag-pep9-fr37) of 20220128.wiff Max. 6.0 %.

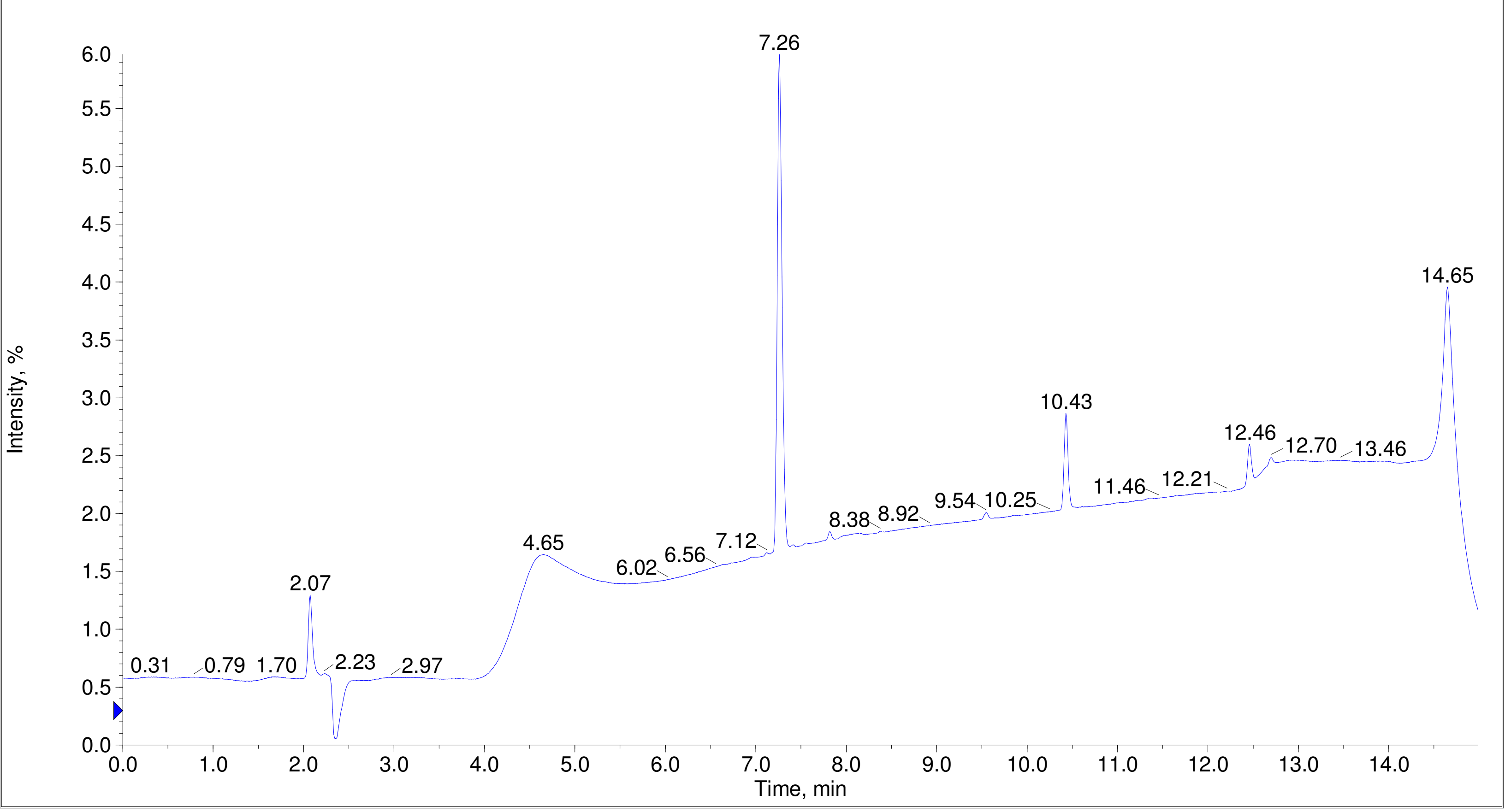

Printing Time: 3:15:02 PM Post PM Polarity/Scan Type: Positive Q1 MS
Printing Date: Tuesday, January 31, 2023 Sample Name: F7-49-ag-pep4-fr36
Service Engineer Chris Maier *Stanford U *API 3000 SN D7640012

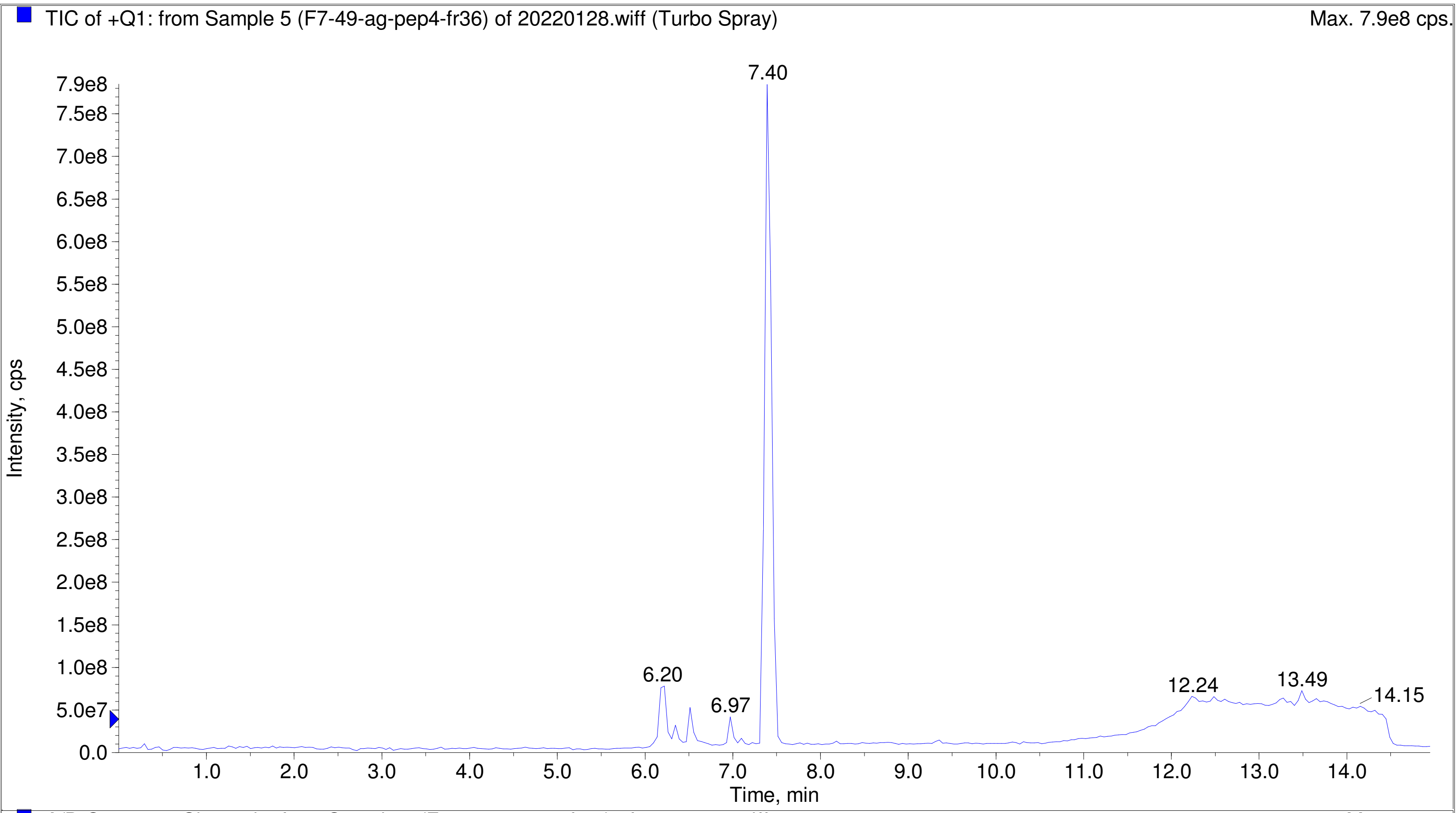


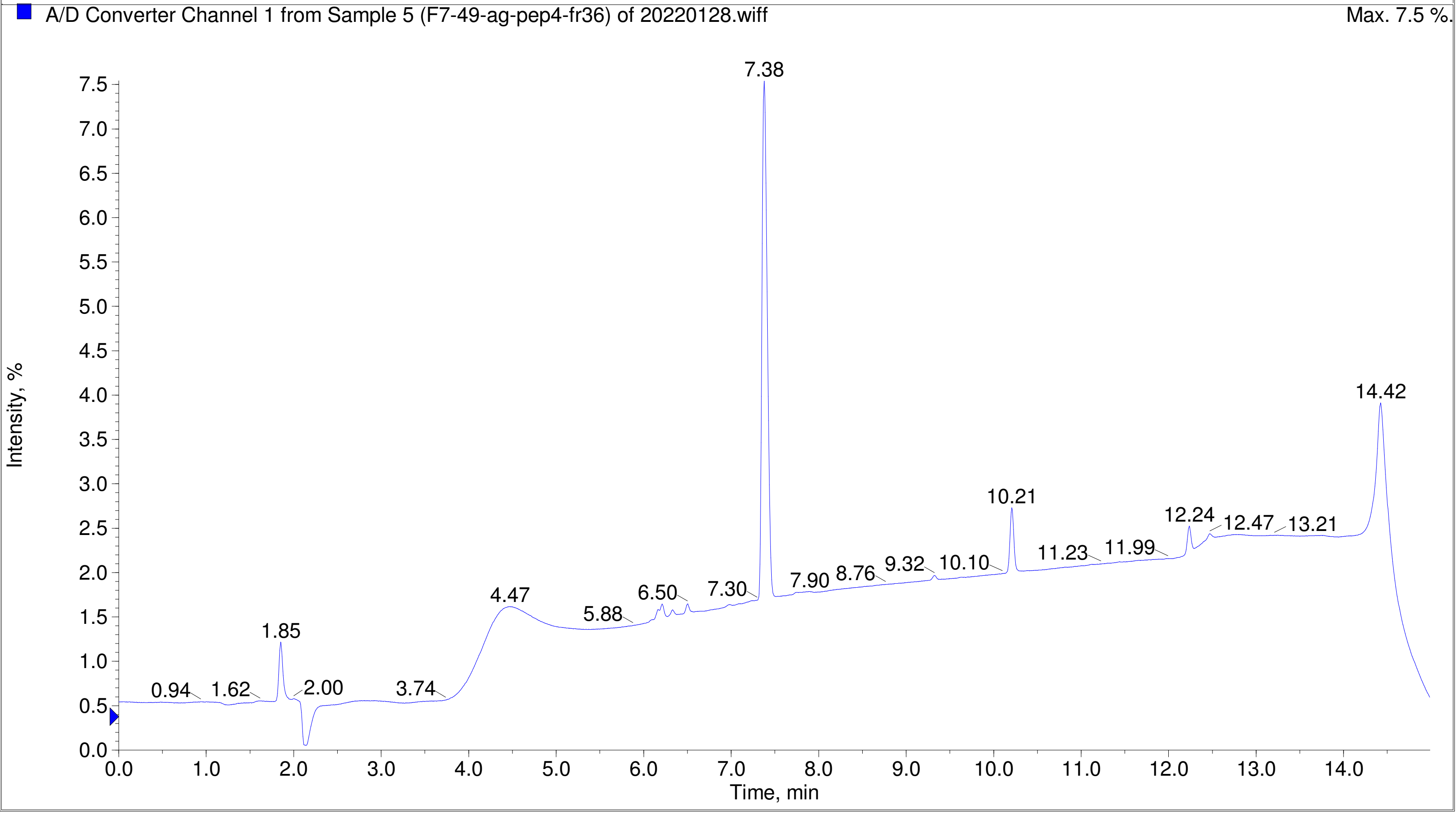

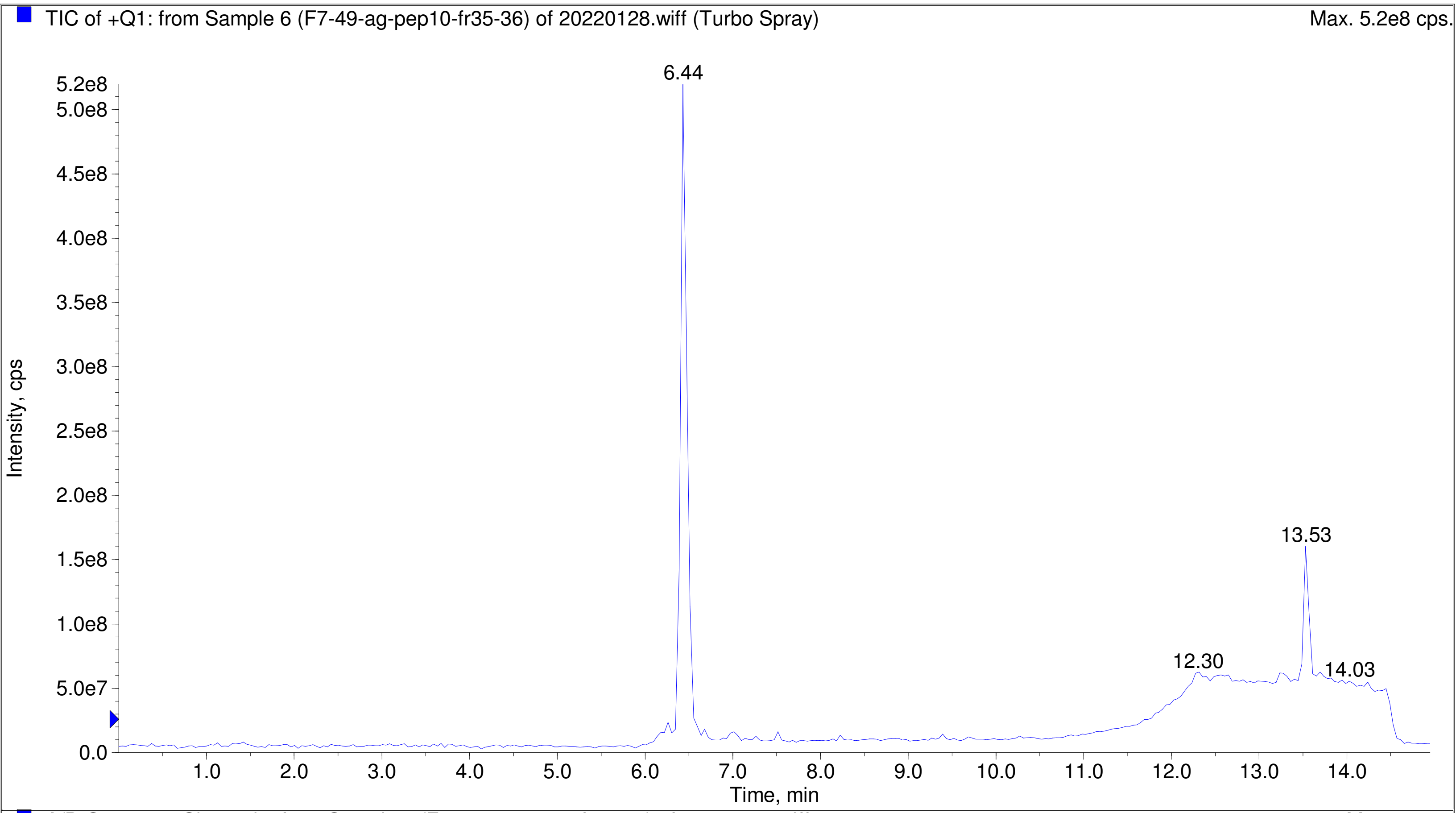
TIC of +Q1: from Sample 6 (F7-49-ag-pep10-fr35-36) of 20220128.wiff (Turbo Spray)
Max. 5.2e8 cps.
6.44
13.53
12.30
14.03
Intensity, cps
Time, min

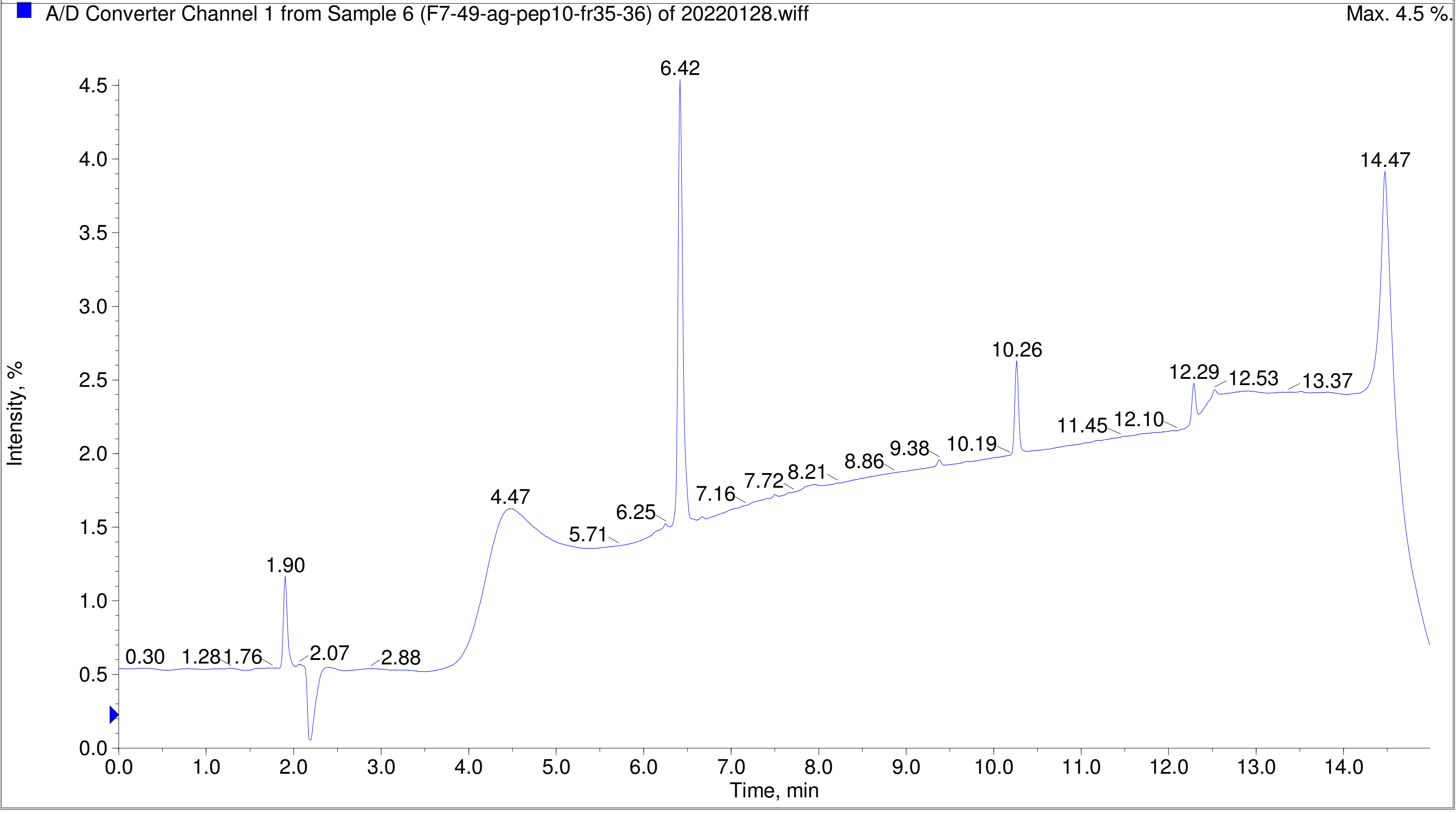
A/D Converter Channel 1 from Sample 6 (F7-49-ag-pep10-fr35-36) of 20220128.wiff
Max. 4.5 %.
6.42
14.47
10.26
12.29
12.53
13.37
11.45
12.10
10.19
9.38
8.86
8.21
7.72
7.16
6.25
5.71
4.47
1.90
0.30
1.28
1.76
2.07
2.88
Intensity, %
Time, min

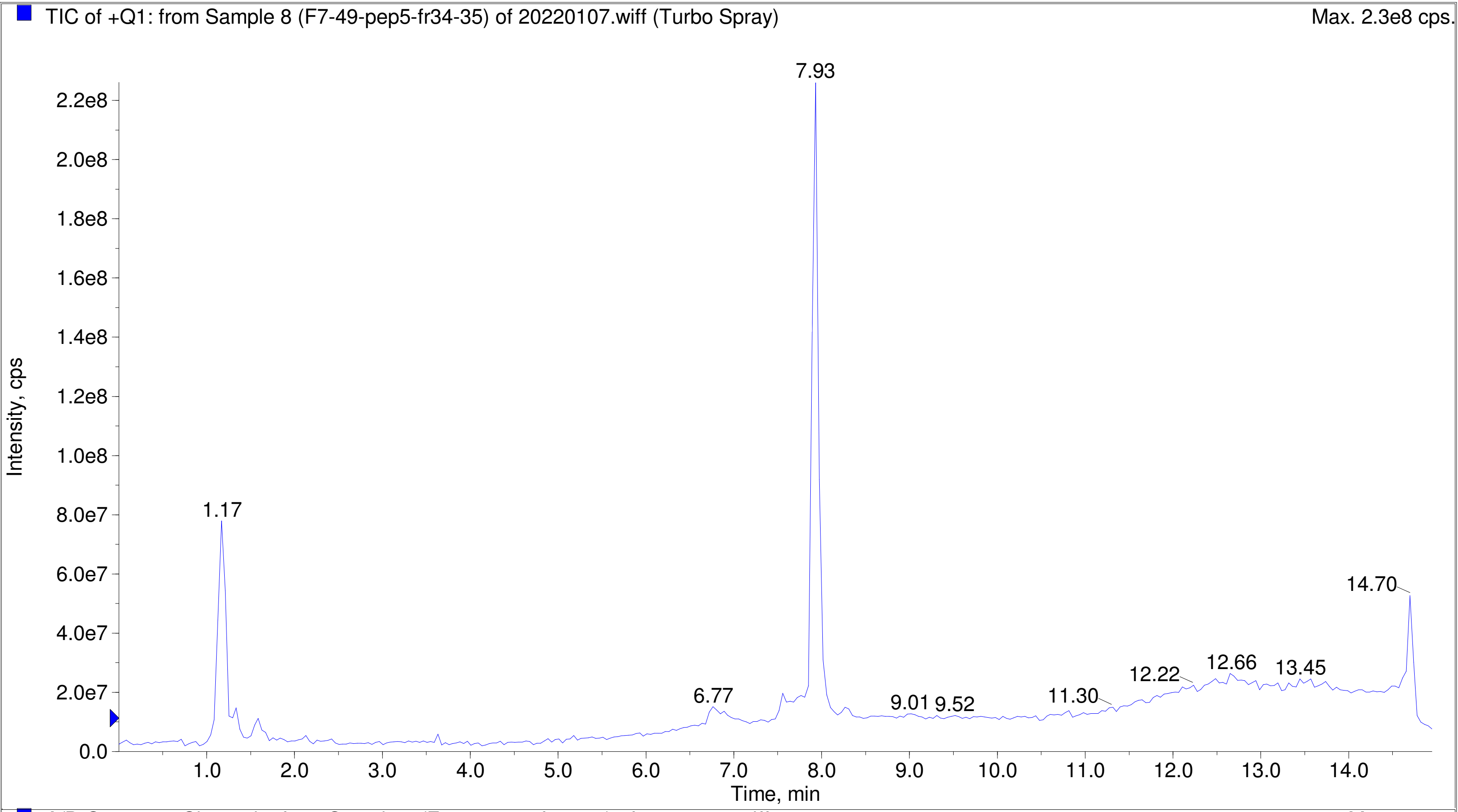
TIC of +Q1: from Sample 8 (F7-49-pep5-fr34-35) of 20220107.wiff (Turbo Spray)
Max. 2.3e8 cps.
Intensity, cps
Time, min
7.93
1.17
14.70
6.77
9.01
9.52
11.30
12.22
12.66
13.45
2.2e8
2.0e8
1.8e8
1.6e8
1.4e8
1.2e8
1.0e8
8.0e7
6.0e7
4.0e7
2.0e7
0.0
1.0
2.0
3.0
4.0
5.0
6.0
7.0
8.0
9.0
10.0
11.0
12.0
13.0
14.0

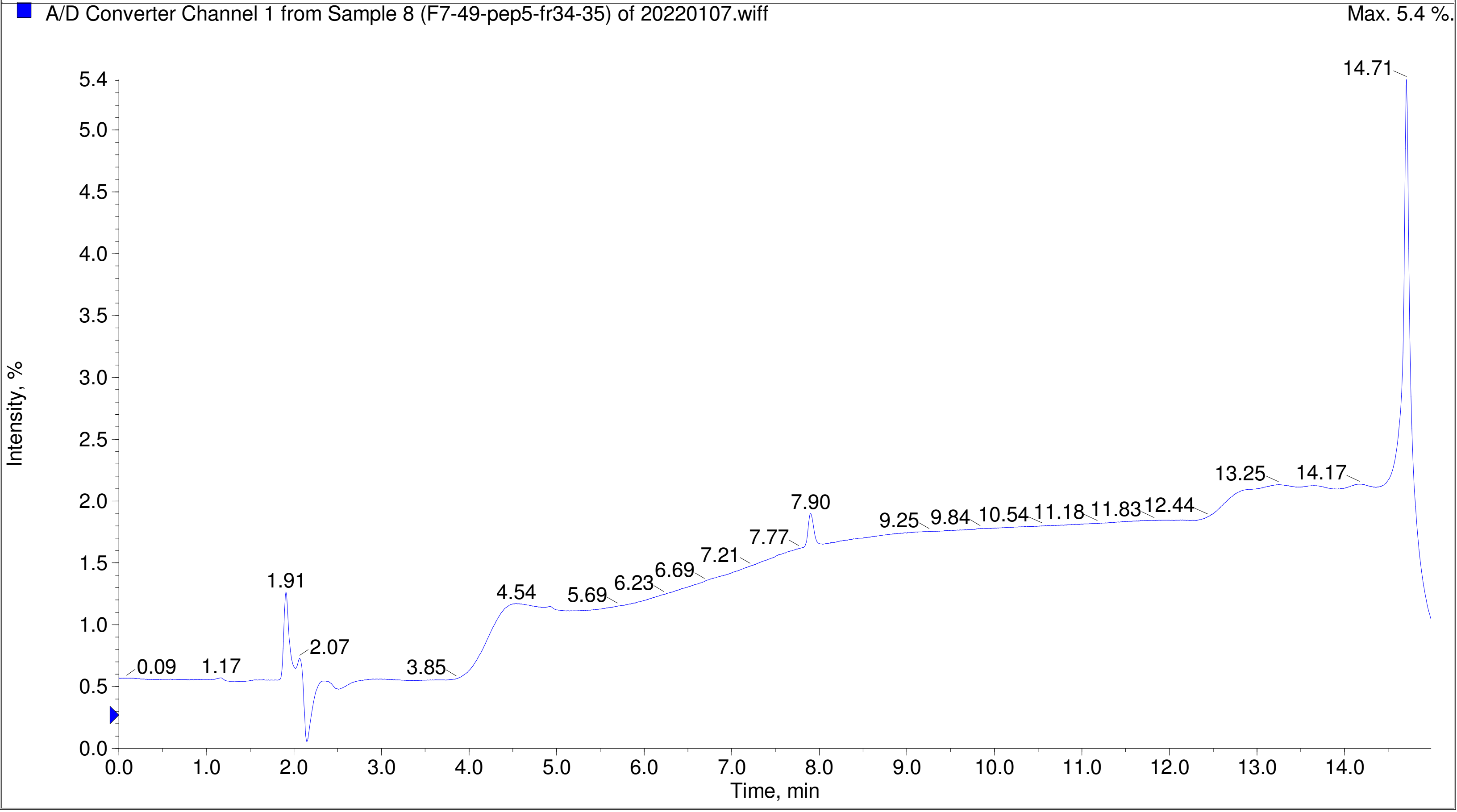
A/D Converter Channel 1 from Sample 8 (F7-49-pep5-fr34-35) of 20220107.wiff
Max. 5.4 %.
Intensity, %
Time, min
14.71
13.25
14.17
7.90
9.25
9.84
10.54
11.18
11.83
12.44
7.77
7.21
6.69
6.23
5.69
4.54
1.91
2.07
0.09
1.17
3.85
5.4
5.0
4.5
4.0
3.5
3.0
2.5
2.0
1.5
1.0
0.5
0.0
0.0
1.0
2.0
3.0
4.0
5.0
6.0
7.0
8.0
9.0
10.0
11.0
12.0
13.0
14.0

TIC of +Q1: from Sample 6 (F7-49-ag-pep11-fr35-36) of 20220122.wiff (Turbo Spray) Max. 1.0e8 cps.

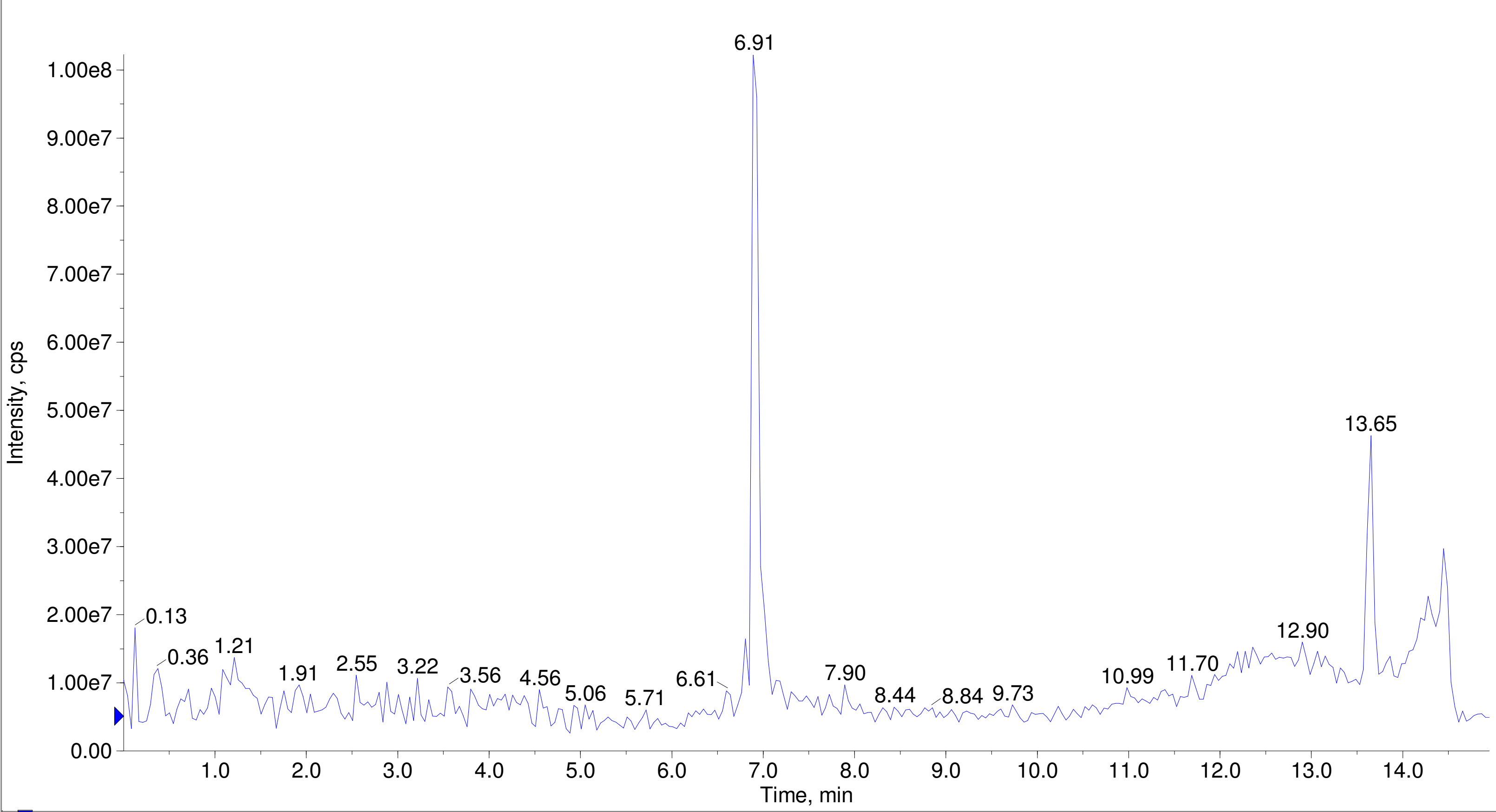


A/D Converter Channel 1 from Sample 6 (F7-49-ag-pep11-fr35-36) of 20220122.wiff Max. 3.8 %.

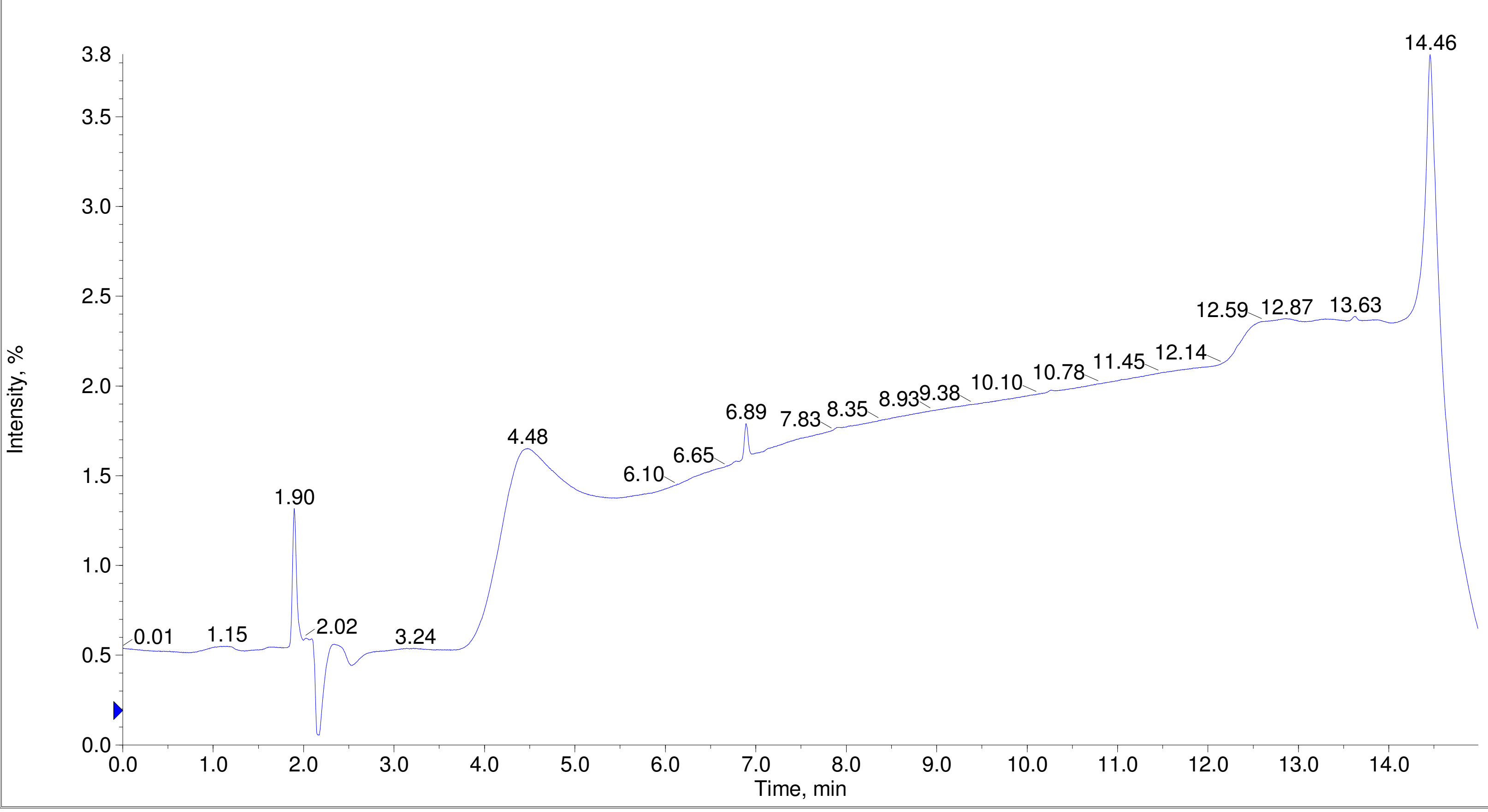

Printing Time: 3:01:15 PM    Post PM    Polarity/Scan Type: Positive Q1 MS
Printing Date: Tuesday, January 31, 2023    Sample Name: F7-49-pep6-fr13-14
Service Engineer Chris Maier    *Stanford U    *API 3000 SN D7640012

TIC of +Q1: from Sample 1 (F7-49-pep6-fr13-14) of 20220107.wiff (Turbo Spray)    Max. 1.7e8 cps.

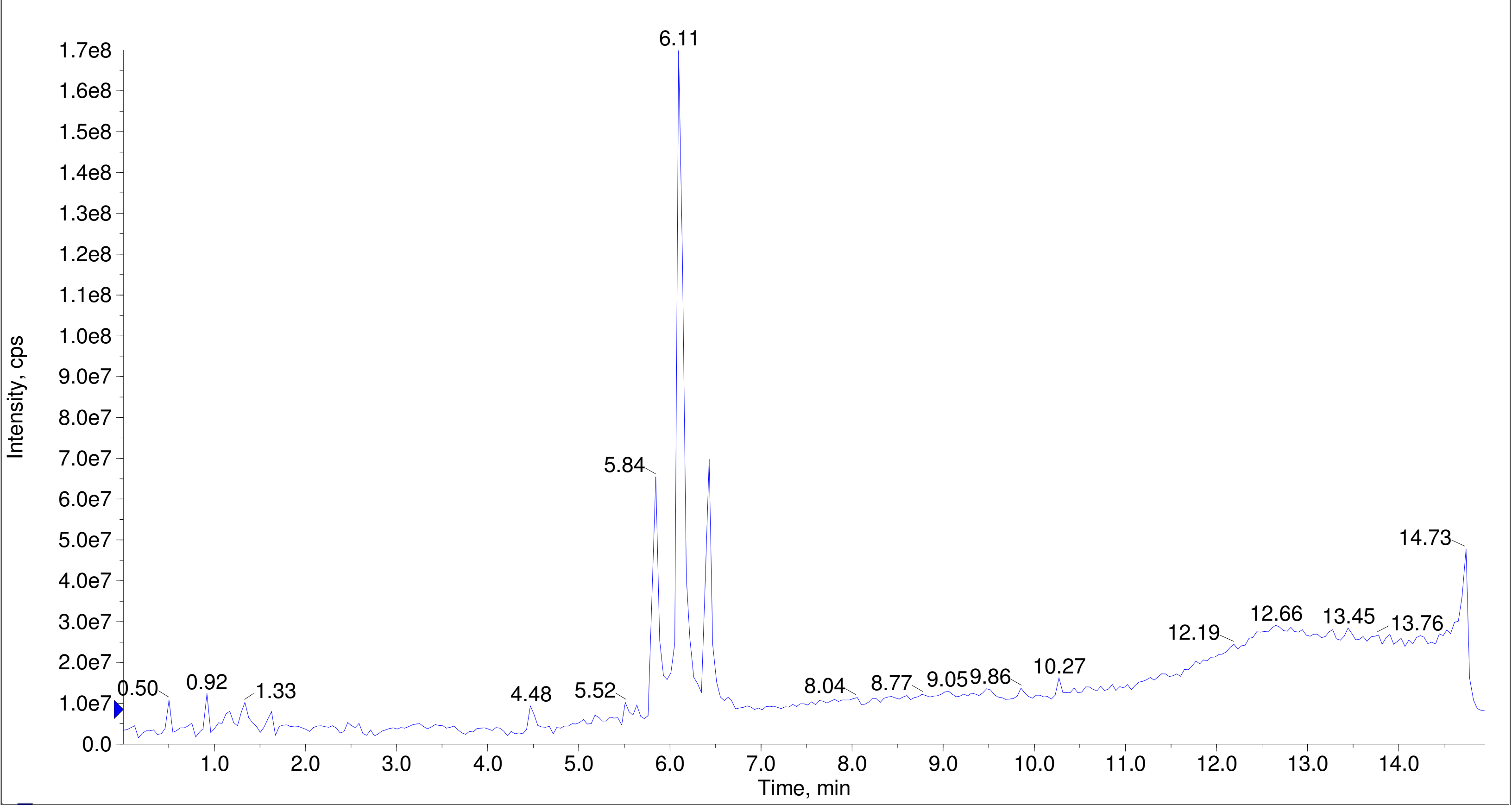


A/D Converter Channel 1 from Sample 1 (F7-49-pep6-fr13-14) of 20220107.wiff    Max. 5.4 %.

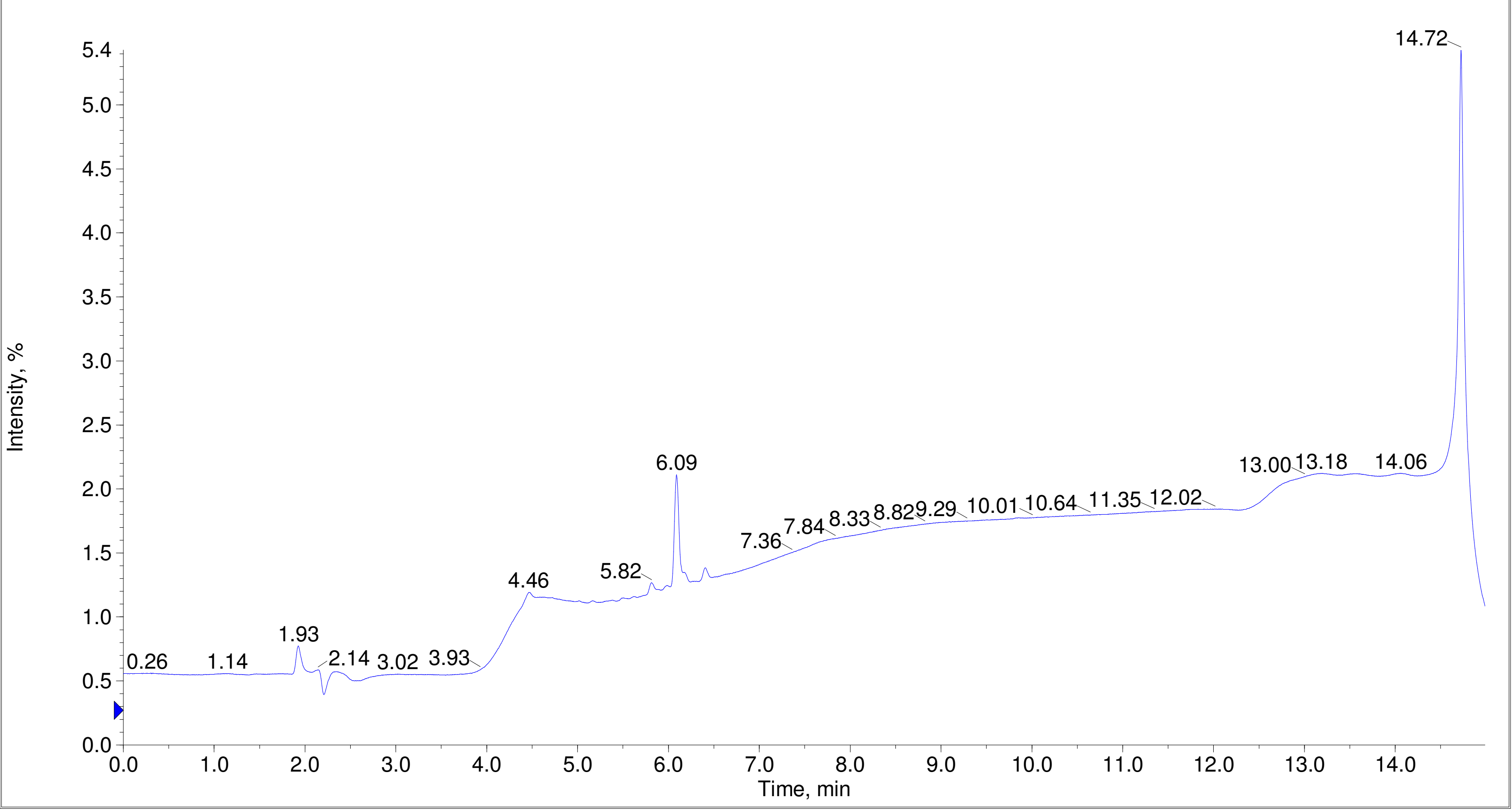

Printing Time: 3:02:38 PM
Printing Date: Tuesday, January 31, 2023
Service Engineer Chris Maier

Post PM
*Stanford U

Polarity/Scan Type: Positive Q1 MS
Sample Name: F7-49-pep12-fr13-14
*API 3000 SN D7640012

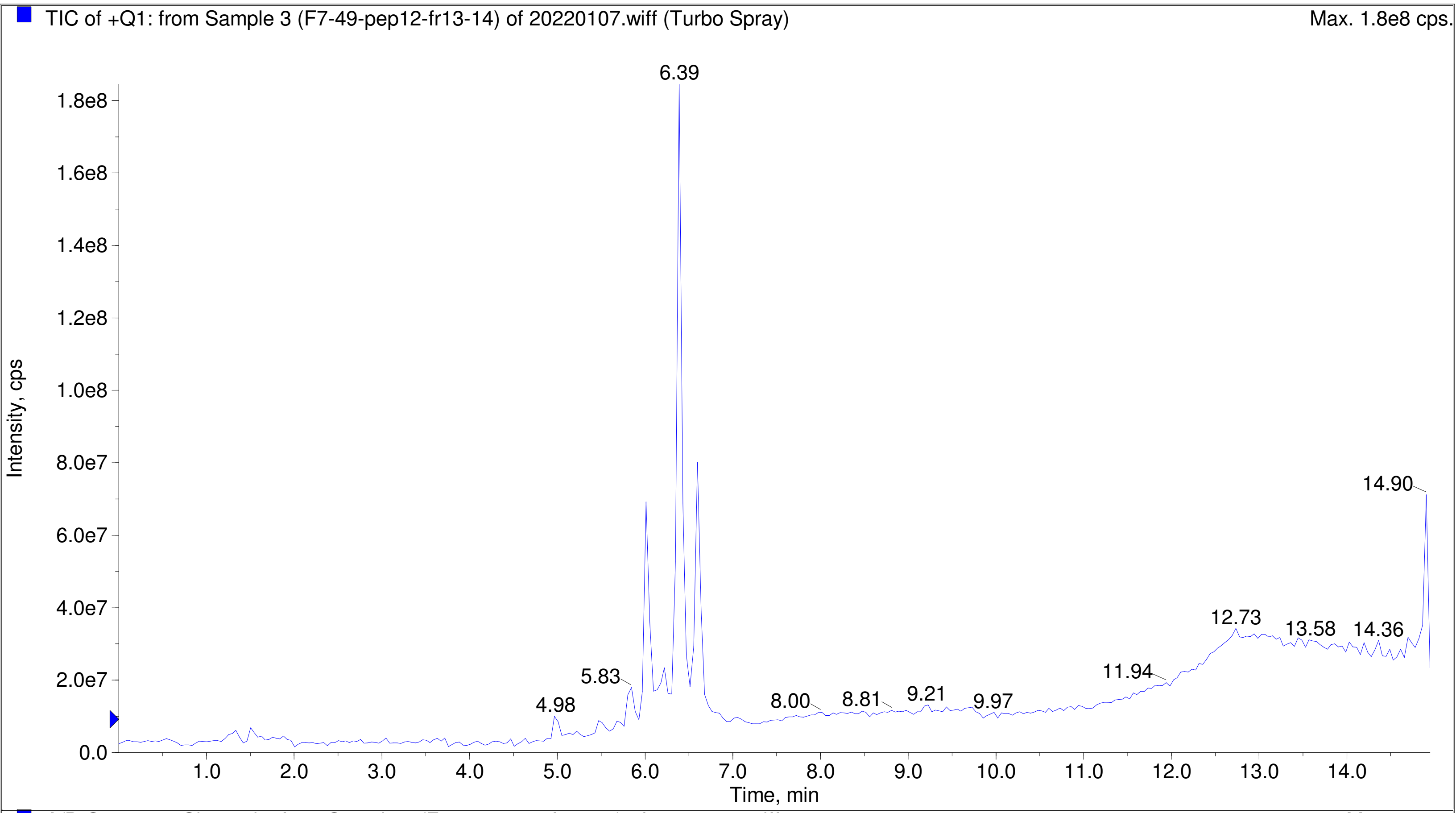


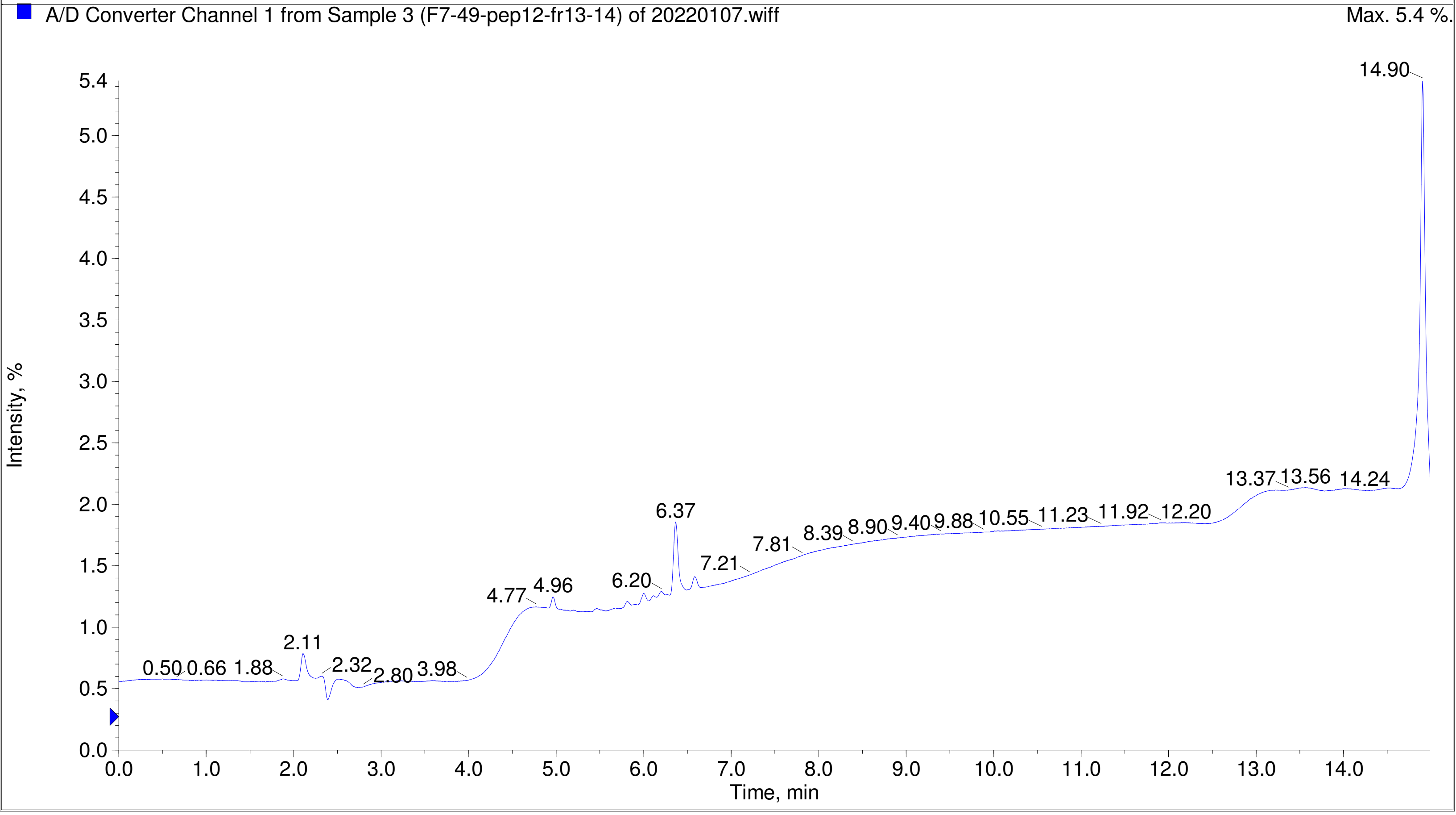

Printing Time: 3:33:30 PM    Post PM    Polarity/Scan Type: Positive Q1 MS
Printing Date: Monday, January 30, 2023    Sample Name:
Service Engineer Chris Maier    *Stanford U    *API 3000 SN D7640012

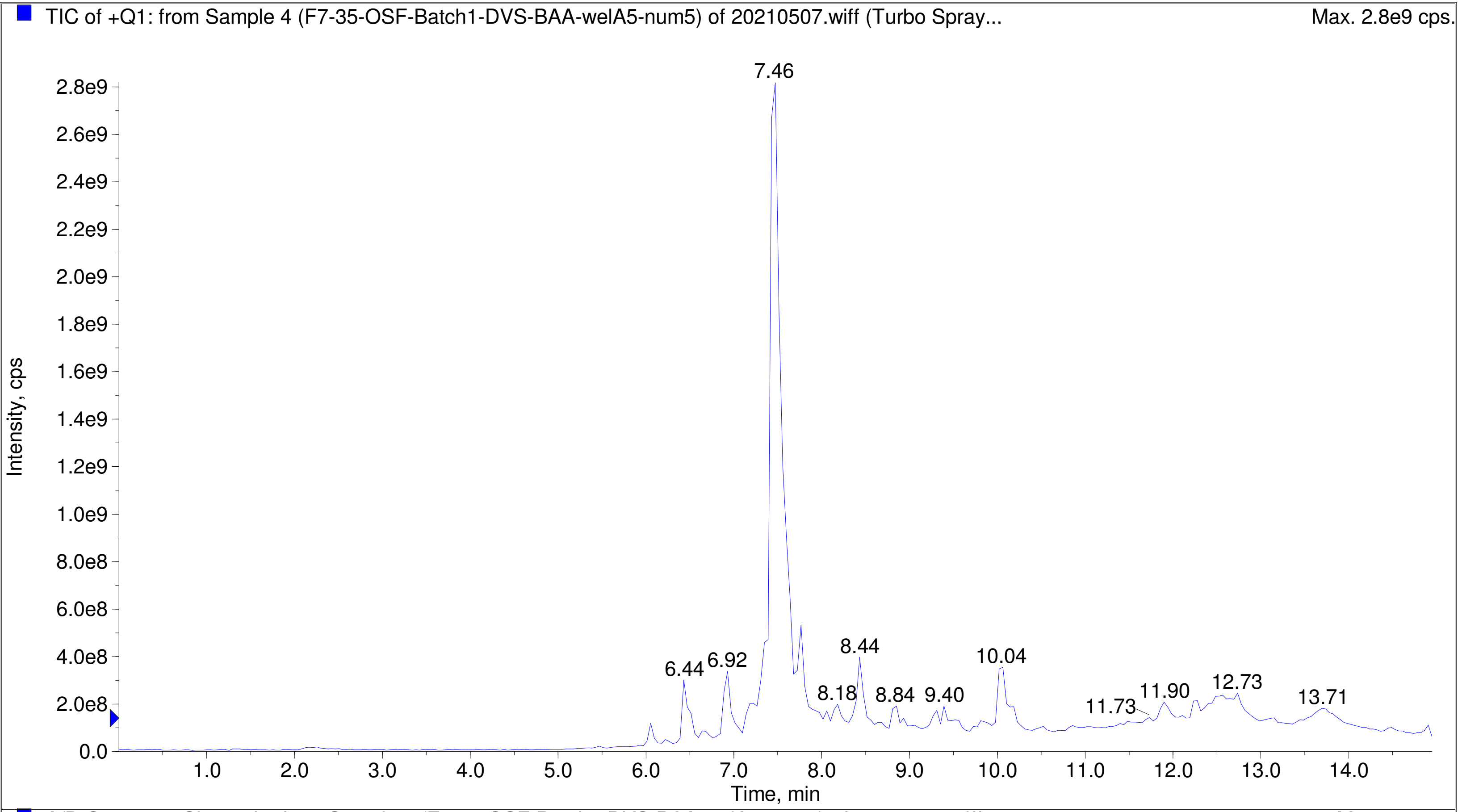


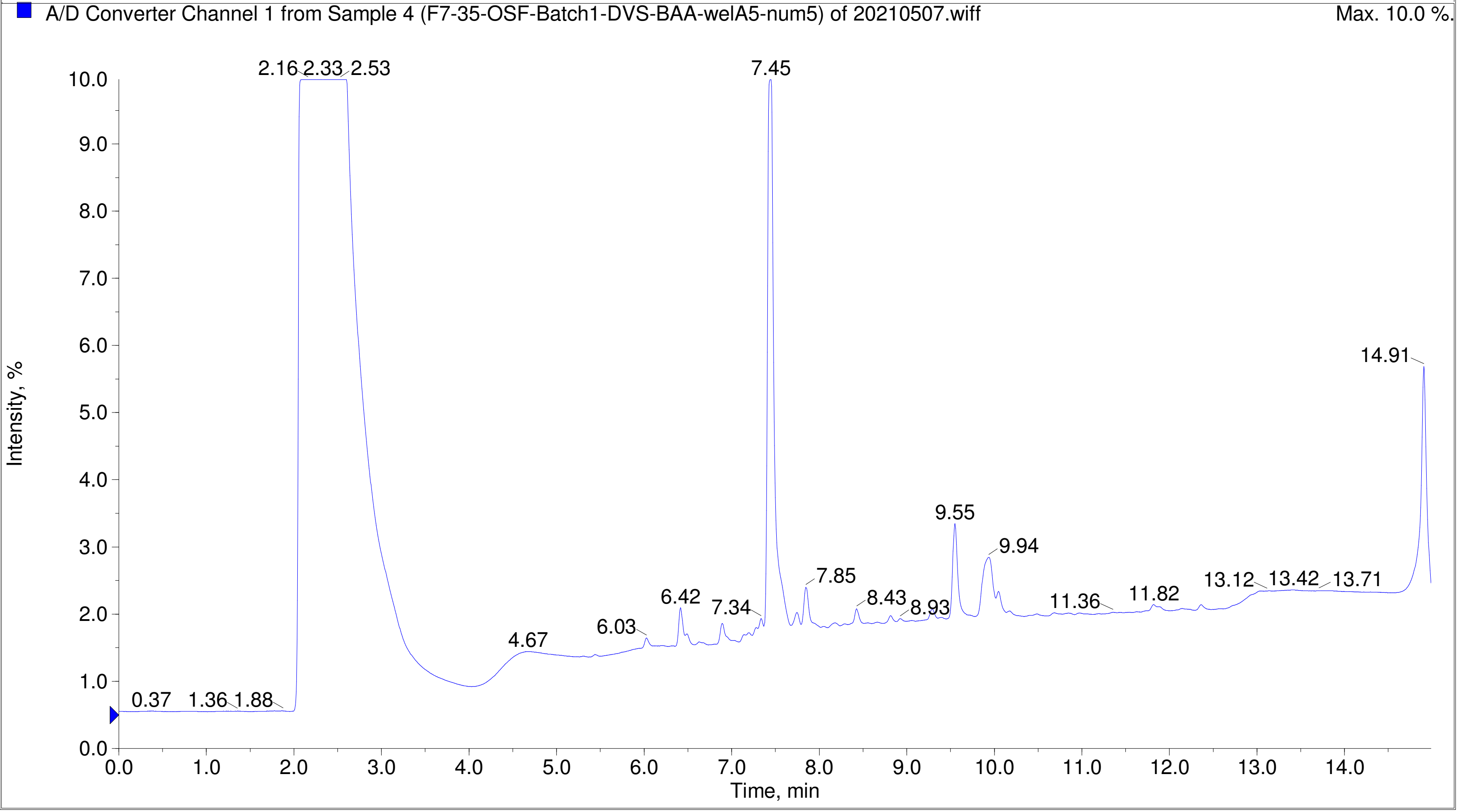

Printing Time: 3:35:23 PM    Post PM    Polarity/Scan Type: Positive Q1 MS
Printing Date: Monday, January 30, 2023    Sample Name:
Service Engineer Chris Maier    *Stanford U    *API 3000 SN D7640012

TIC of +Q1: from Sample 3 (F7-35-OSF-Batch1-DVS-BAA-blank-concag) of 20210507.wiff (Turbo Spray...    Max. 1.1e9 cps.

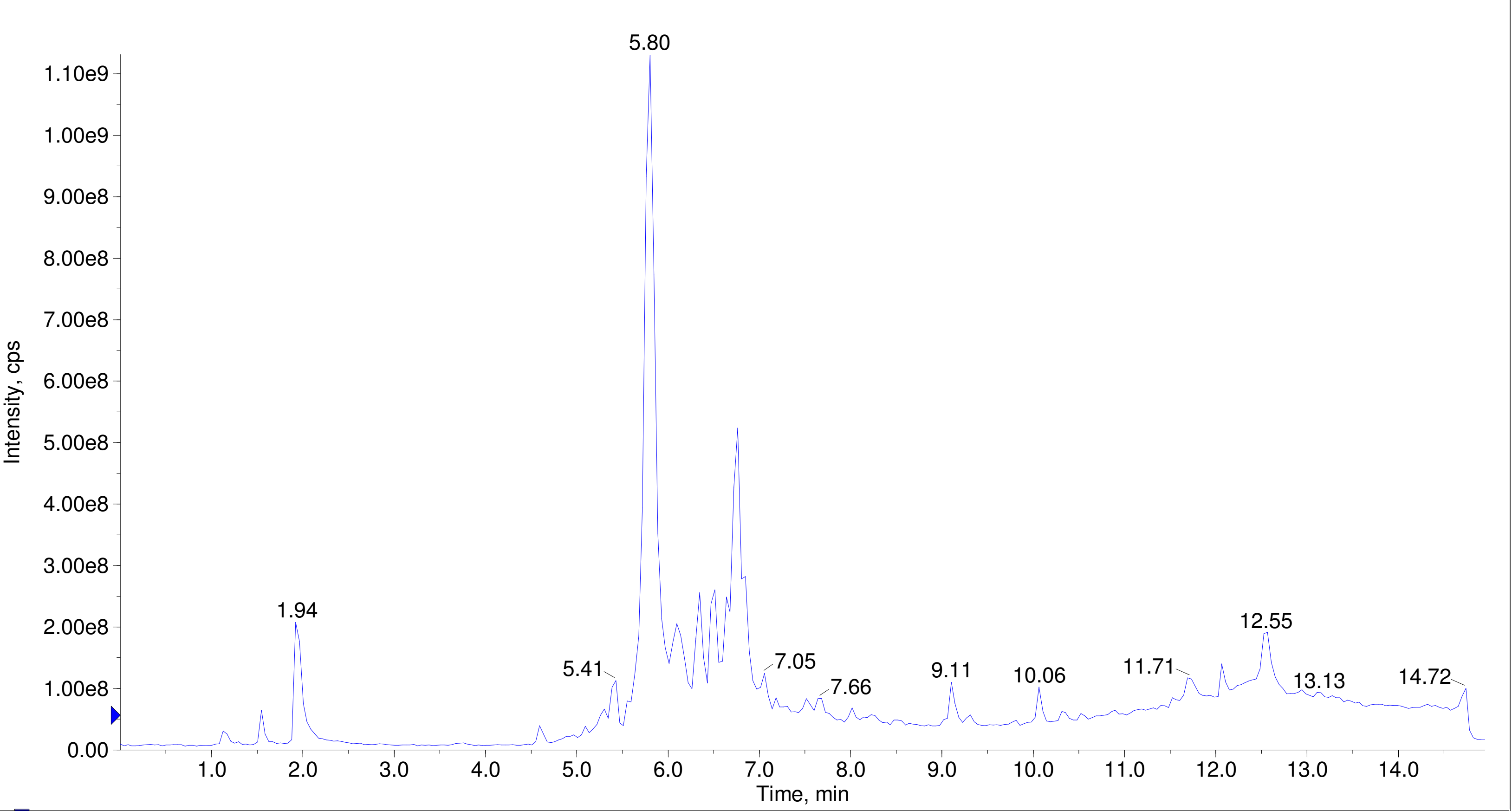


A/D Converter Channel 1 from Sample 3 (F7-35-OSF-Batch1-DVS-BAA-blank-concag) of 20210507.wiff    Max. 10.0 %.

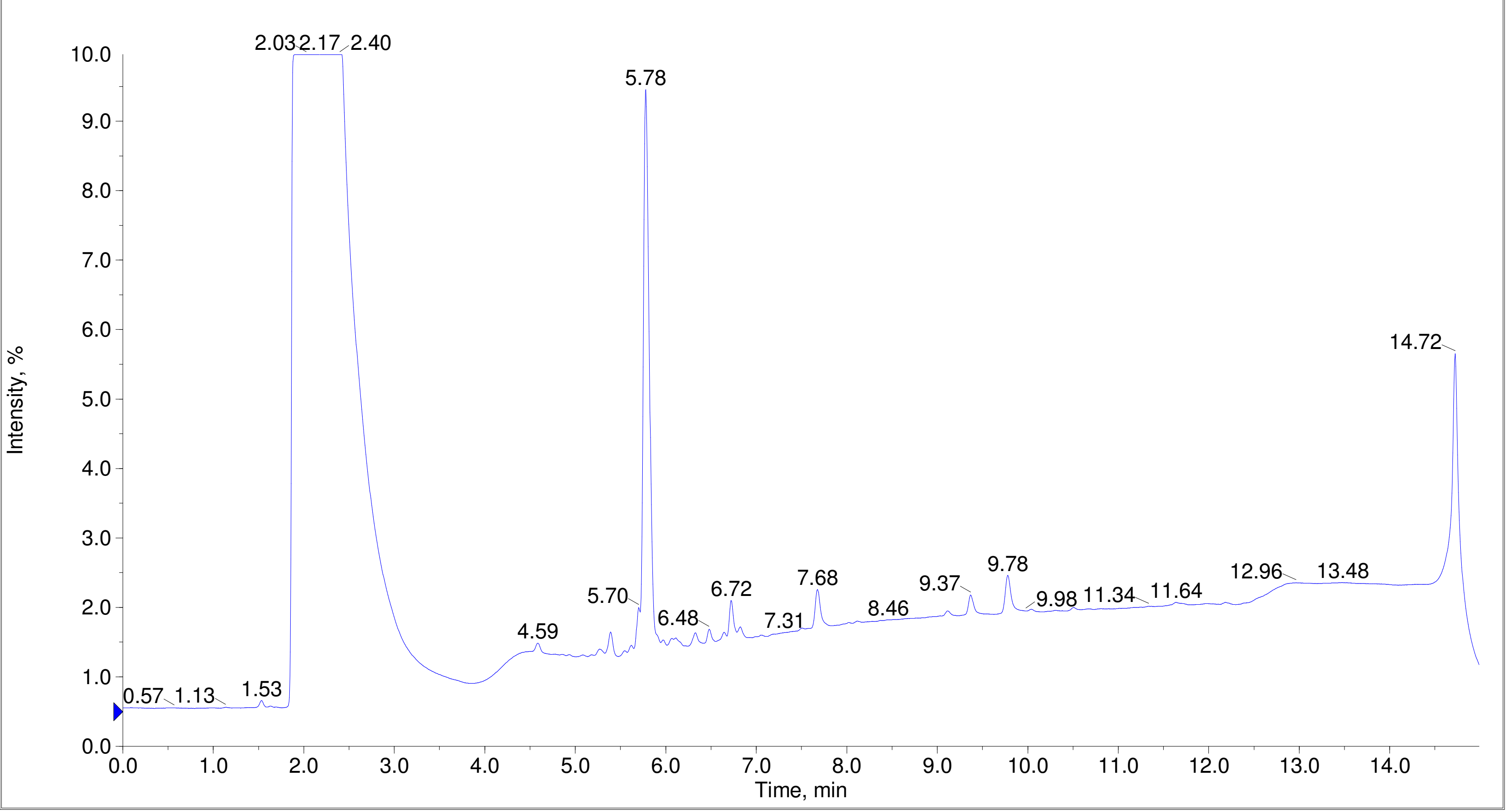

Printing Time: 3:32:26 PM Post PM Polarity/Scan Type: Positive Q1 MS
Printing Date: Monday, January 30, 2023 Sample Name:
Service Engineer Chris Maier *Stanford U *API 3000 SN D7640012

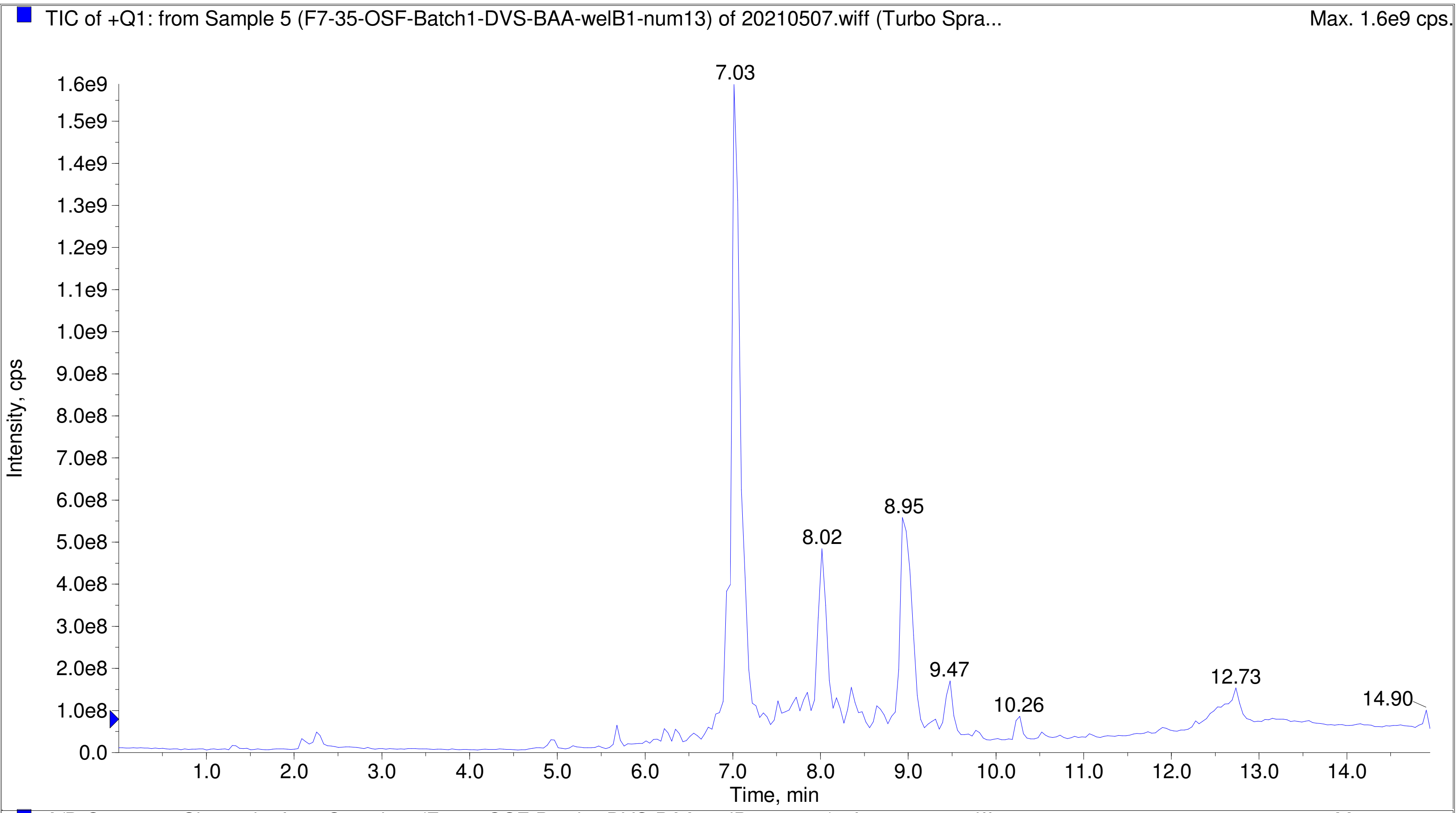


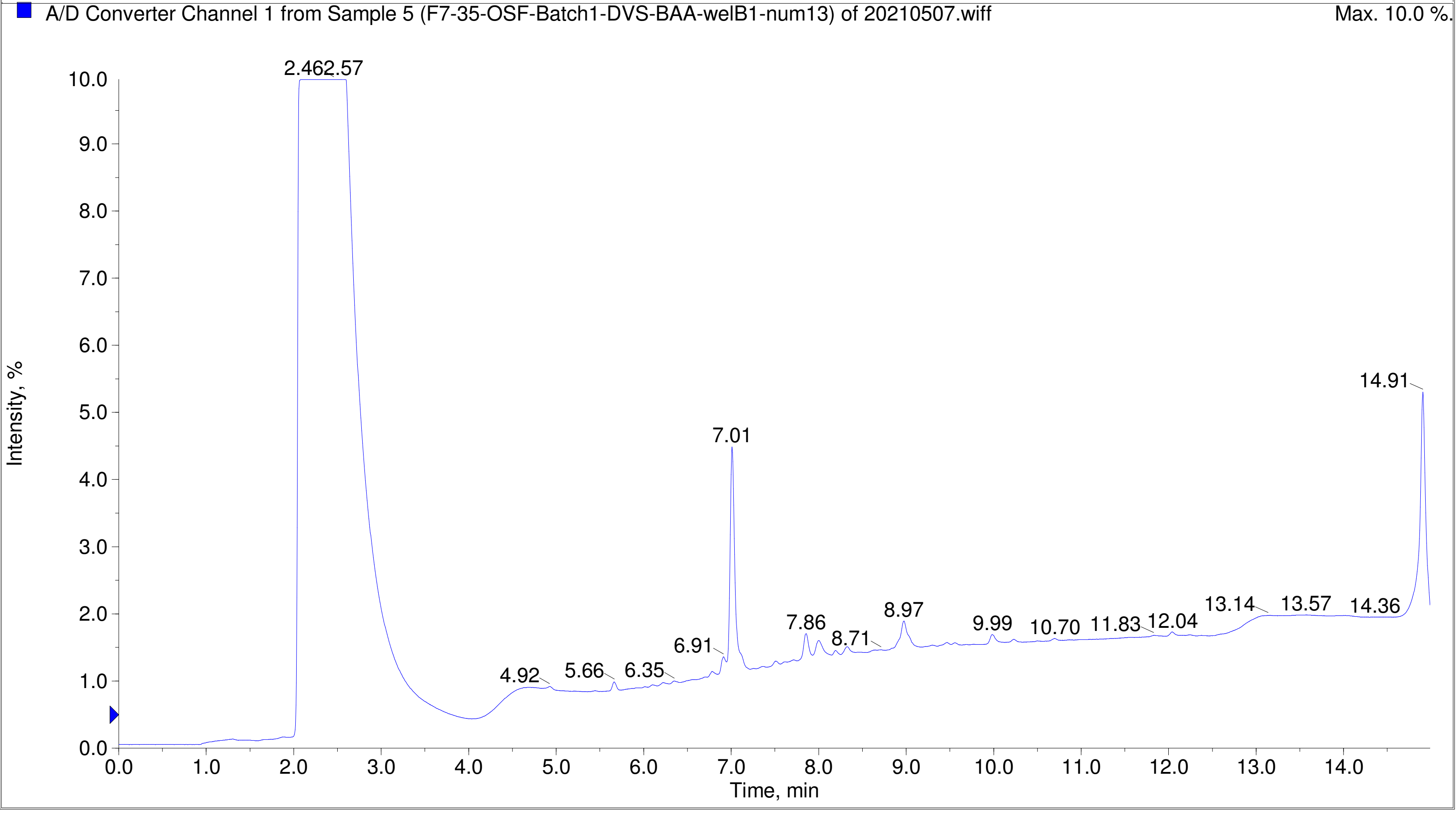

Printing Time: 3:31:12 PM  Post PM  Polarity/Scan Type: Positive Q1 MS
Printing Date: Monday, January 30, 2023  Sample Name:
Service Engineer Chris Maier  *Stanford U  *API 3000 SN D7640012

TIC of +Q1: from Sample 6 (F7-35-OSF-Batch1-DVS-BAA-welB7-num19) of 20210507.wiff (Turbo Spra...  Max. 2.5e9 cps.

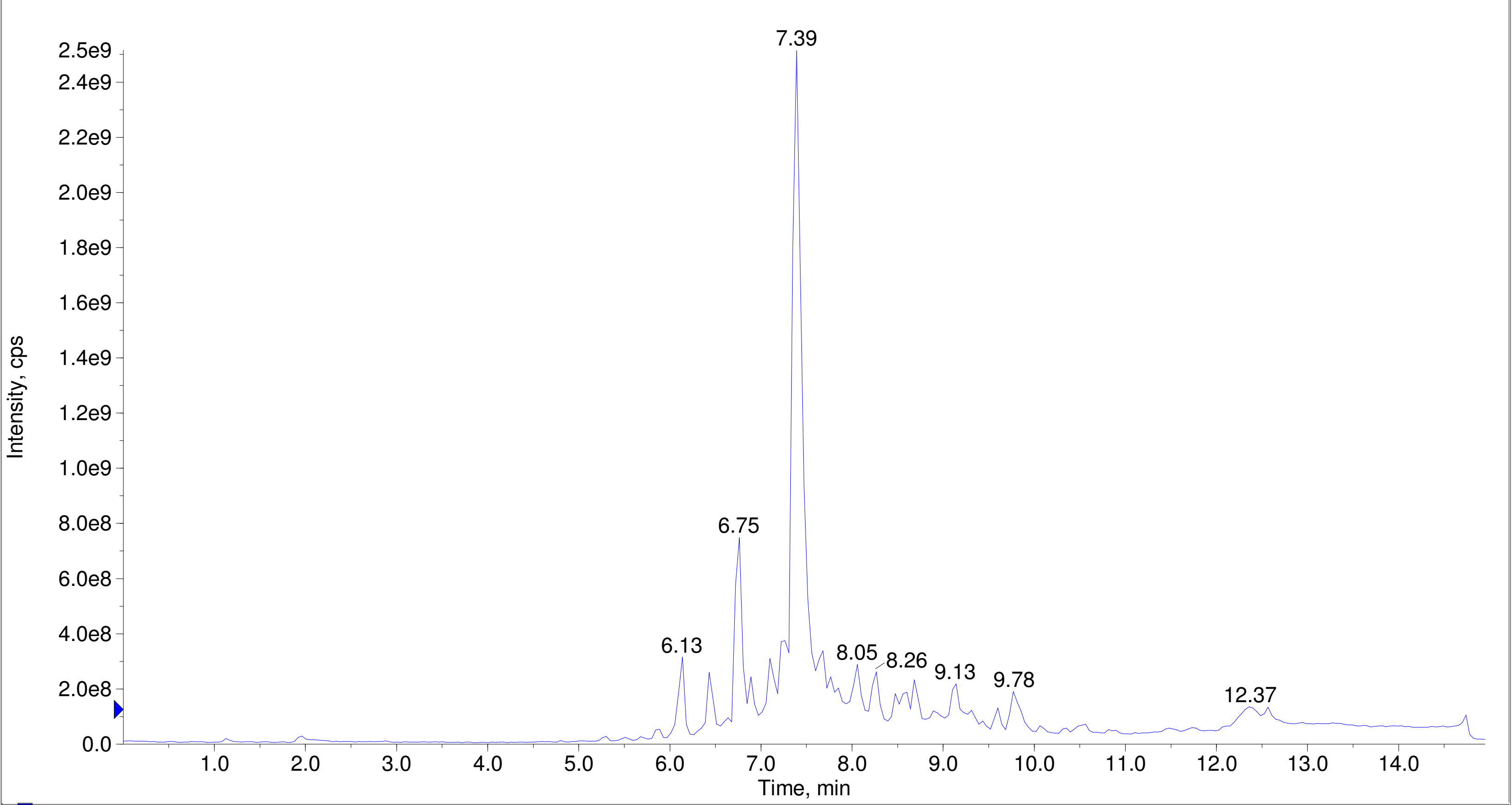


A/D Converter Channel 1 from Sample 6 (F7-35-OSF-Batch1-DVS-BAA-welB7-num19) of 20210507.wiff  Max. 10.0 %.

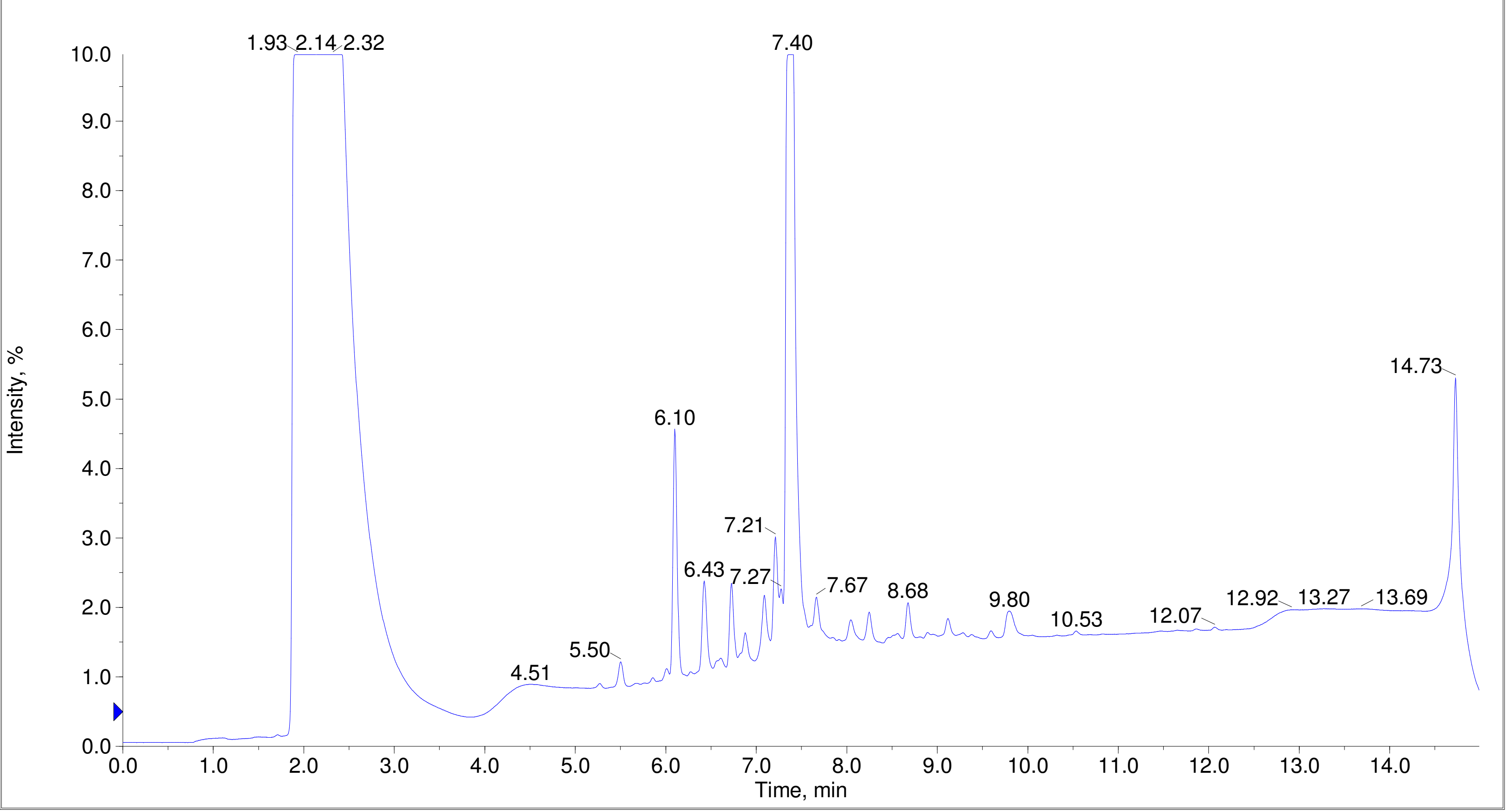

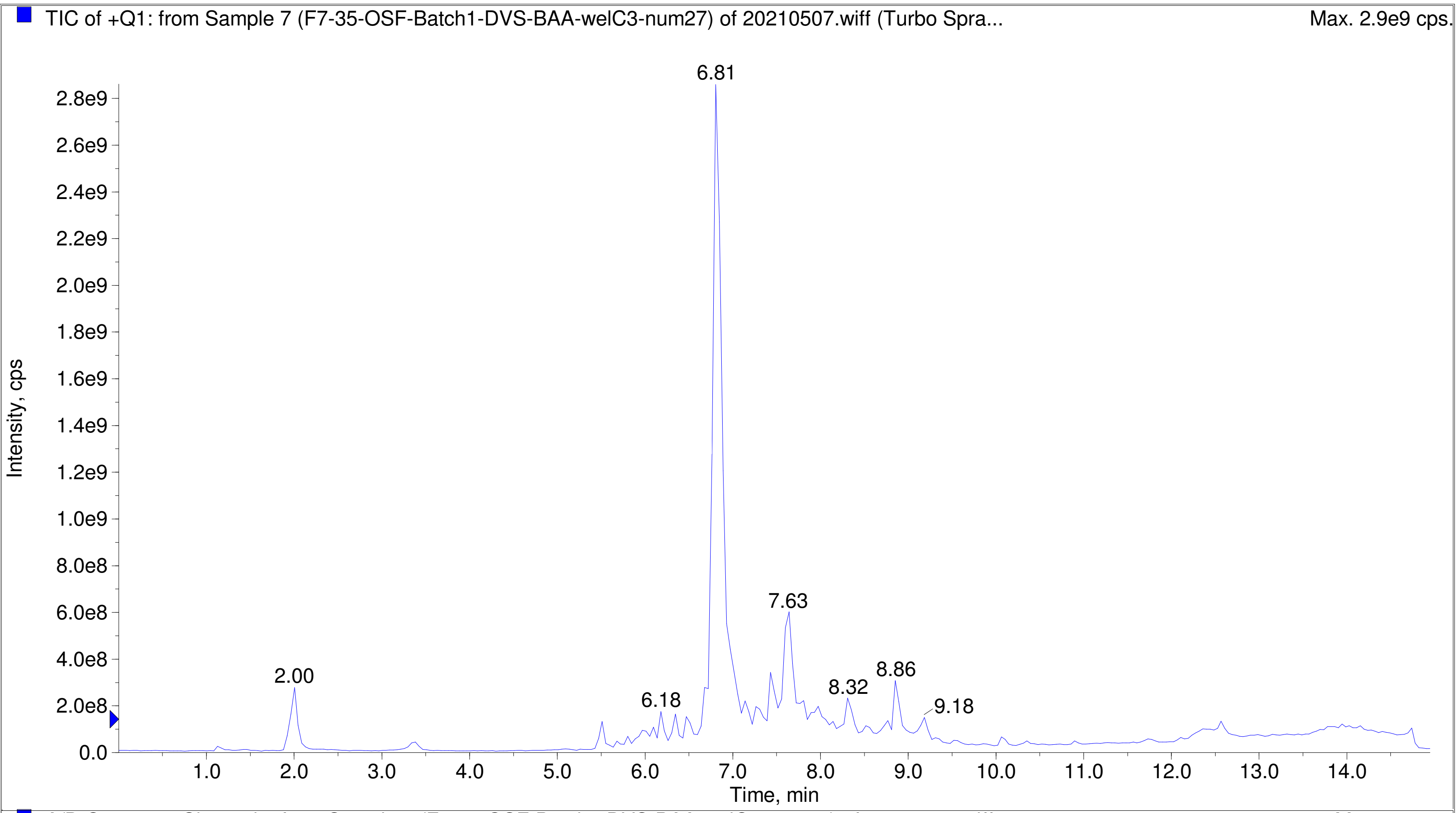
TIC of +Q1: from Sample 7 (F7-35-OSF-Batch1-DVS-BAA-welC3-num27) of 20210507.wiff (Turbo Spra...
Max. 2.9e9 cps.
Intensity, cps
Time, min
6.81
7.63
2.00
6.18
8.32
8.86
9.18

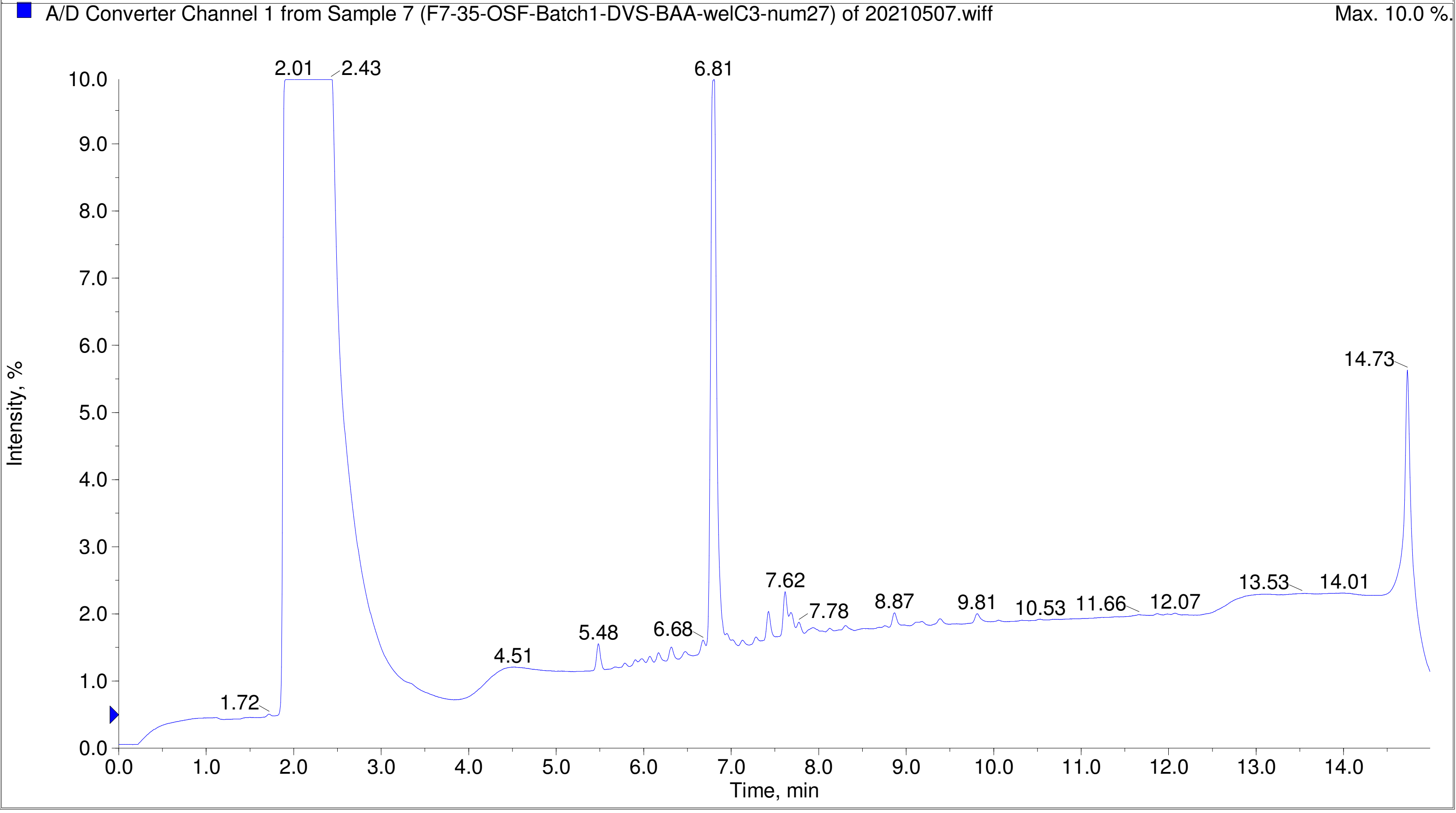
A/D Converter Channel 1 from Sample 7 (F7-35-OSF-Batch1-DVS-BAA-welC3-num27) of 20210507.wiff
Max. 10.0 %.
Intensity, %
Time, min
2.01
2.43
6.81
14.73
1.72
4.51
5.48
6.68
7.62
7.78
8.87
9.81
10.53
11.66
12.07
13.53
14.01

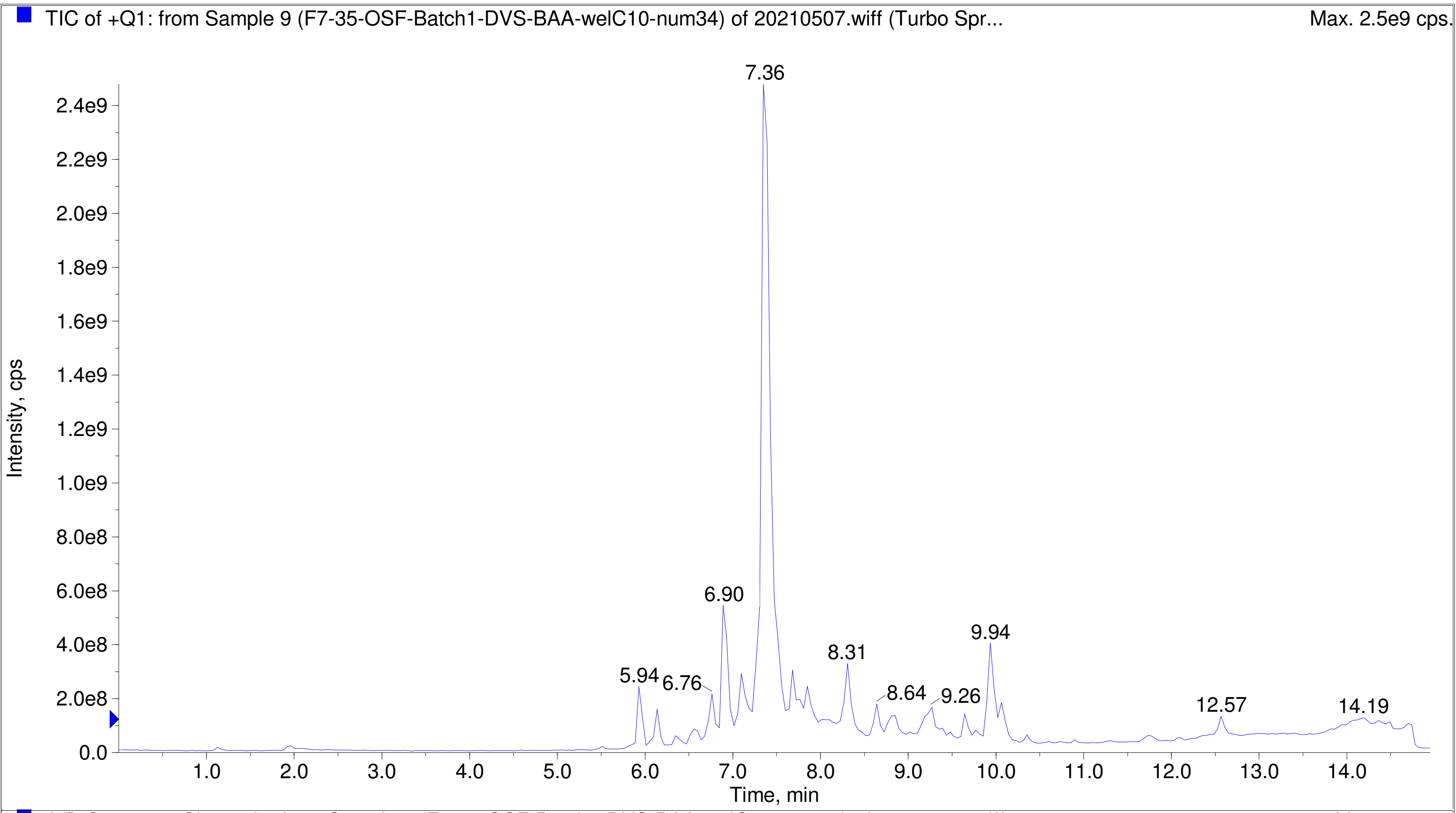
TIC of +Q1: from Sample 9 (F7-35-OSF-Batch1-DVS-BAA-welC10-num34) of 20210507.wiff (Turbo Spr...
Max. 2.5e9 cps.
Intensity, cps
Time, min
7.36
6.90
5.94
6.76
8.31
8.64
9.26
9.94
12.57
14.19

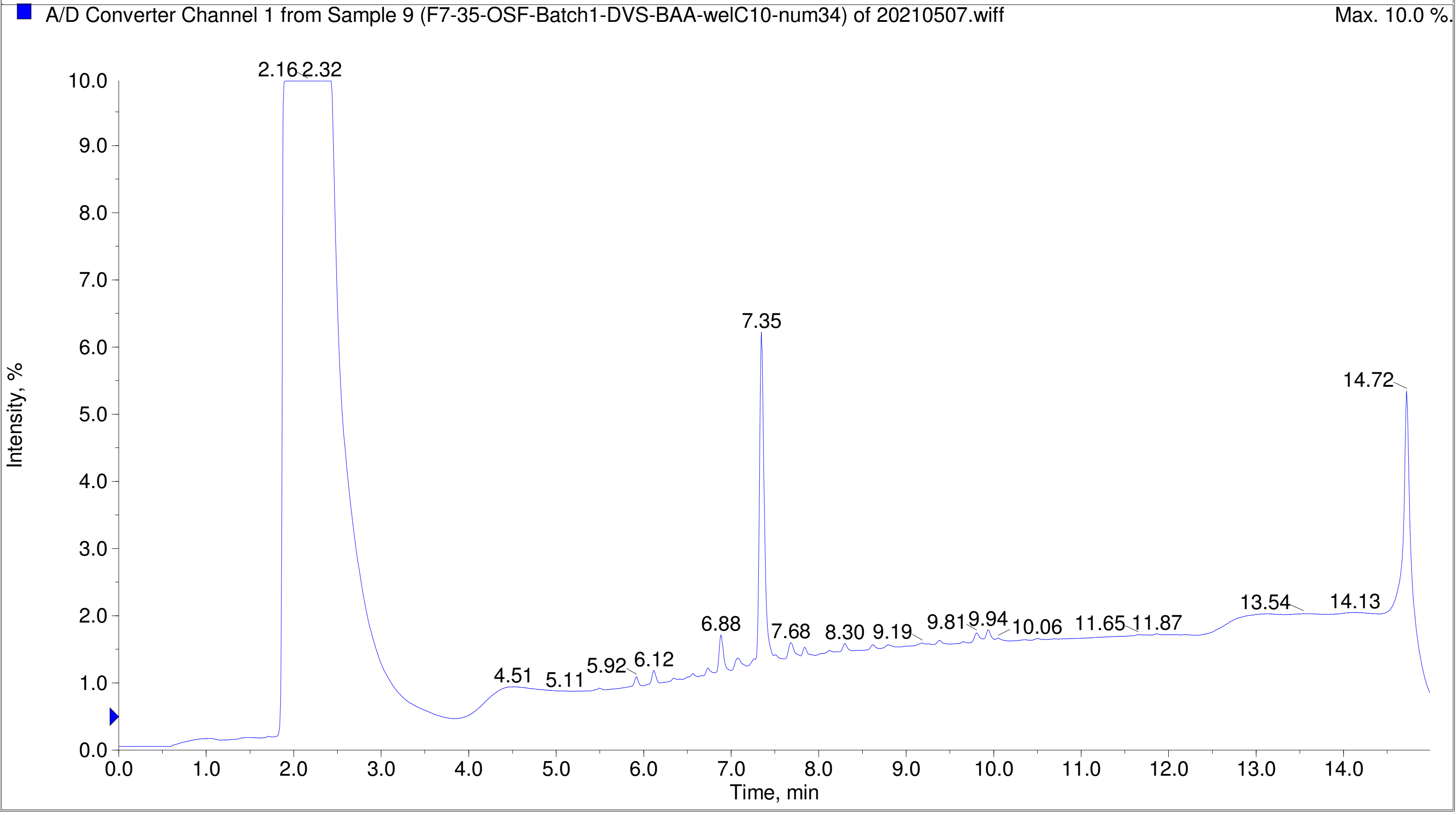
A/D Converter Channel 1 from Sample 9 (F7-35-OSF-Batch1-DVS-BAA-welC10-num34) of 20210507.wiff
Max. 10.0 %.
Intensity, %
Time, min
2.16
2.32
7.35
14.72
4.51
5.11
5.92
6.12
6.88
7.68
8.30
9.19
9.81
9.94
10.06
11.65
11.87
13.54
14.13

Printing Time: 7:33:17 PM
Printing Date: Monday, January 30, 2023
Service Engineer Chris Maier

Post PM

*Stanford U

Polarity/Scan Type: Positive Q1 MS
Sample Name:
*API 3000 SN D7640012

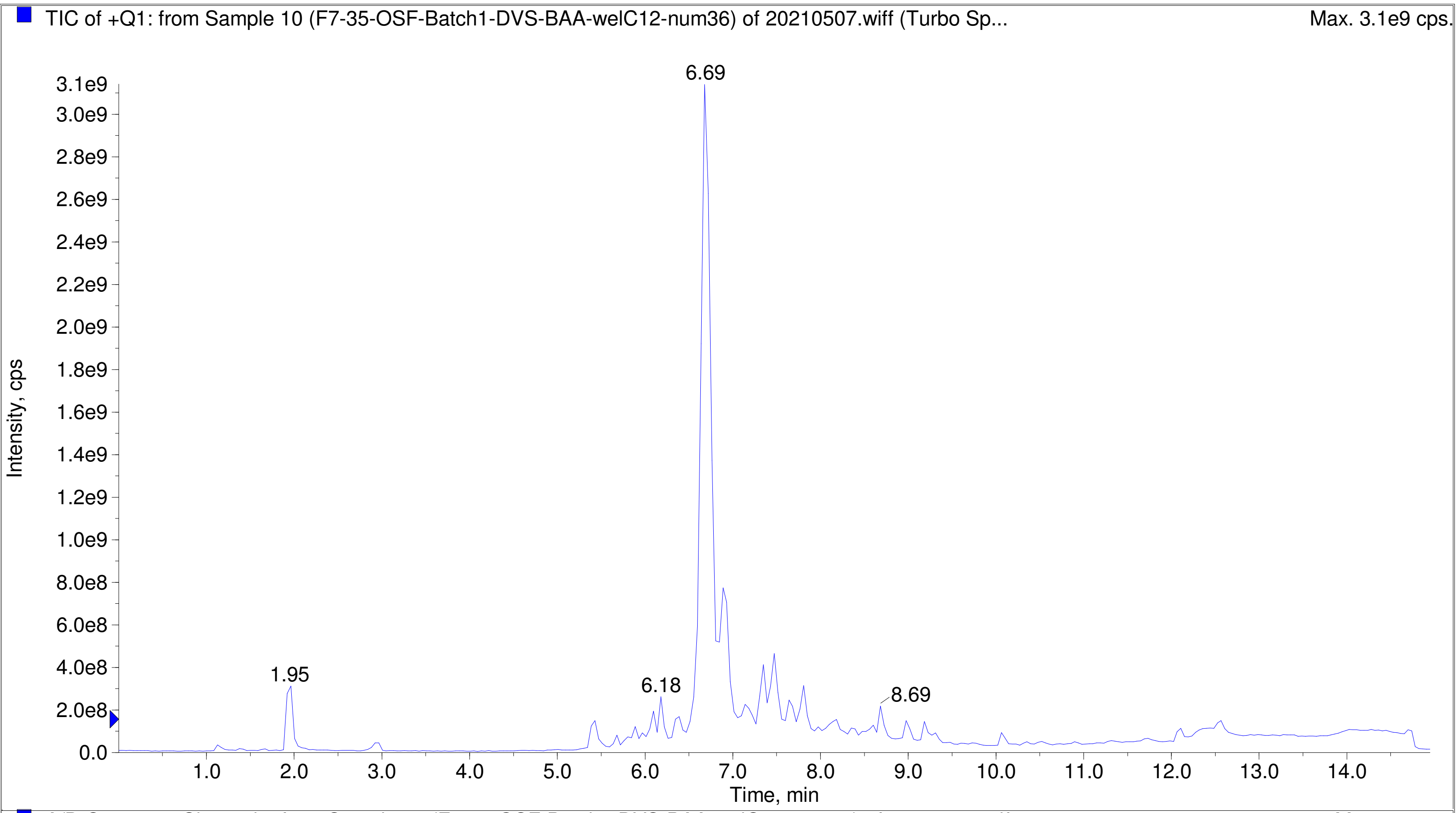


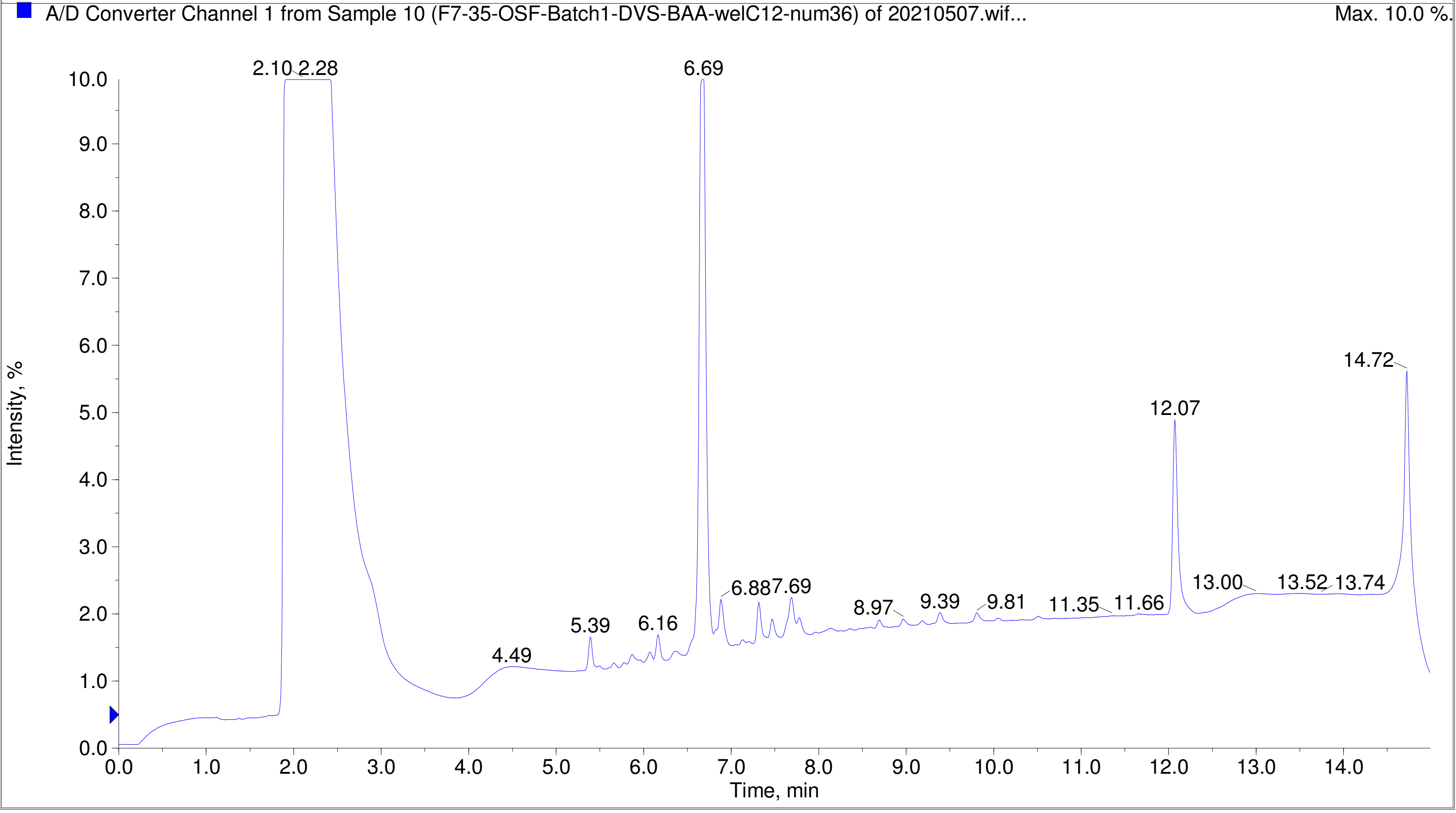

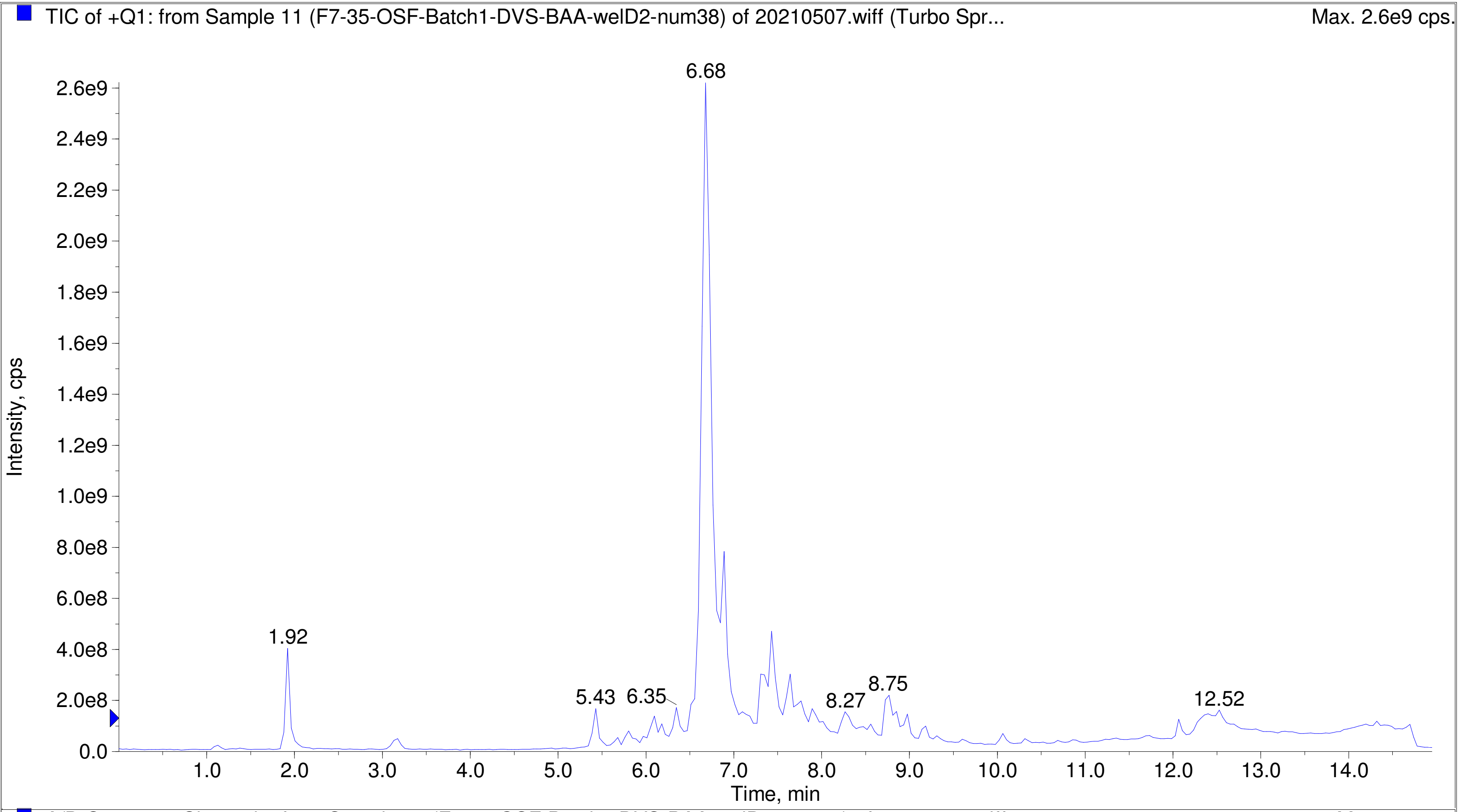
TIC of +Q1: from Sample 11 (F7-35-OSF-Batch1-DVS-BAA-welD2-num38) of 20210507.wiff (Turbo Spr...
Max. 2.6e9 cps.
Intensity, cps
Time, min
6.68
1.92
5.43
6.35
8.27
8.75
12.52
0.0
2.0e8
4.0e8
6.0e8
8.0e8
1.0e9
1.2e9
1.4e9
1.6e9
1.8e9
2.0e9
2.2e9
2.4e9
2.6e9
1.0
2.0
3.0
4.0
5.0
6.0
7.0
8.0
9.0
10.0
11.0
12.0
13.0
14.0

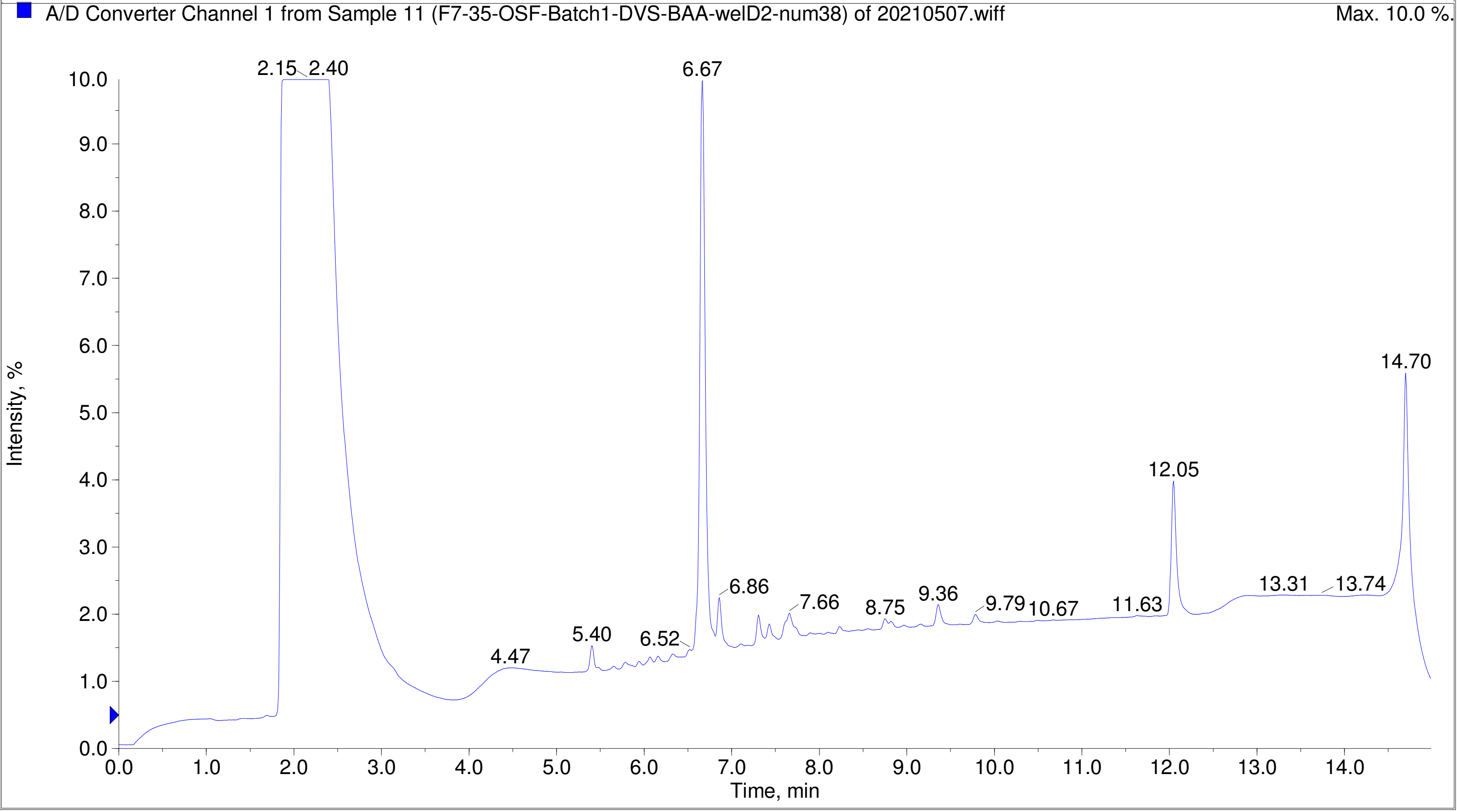
A/D Converter Channel 1 from Sample 11 (F7-35-OSF-Batch1-DVS-BAA-welD2-num38) of 20210507.wiff
Max. 10.0 %.
Intensity, %
Time, min
2.15
2.40
6.67
14.70
12.05
6.86
7.66
8.75
9.36
9.79
10.67
11.63
13.31
13.74
4.47
5.40
6.52
0.0
1.0
2.0
3.0
4.0
5.0
6.0
7.0
8.0
9.0
10.0
0.0
1.0
2.0
3.0
4.0
5.0
6.0
7.0
8.0
9.0
10.0
11.0
12.0
13.0
14.0

TIC of +Q1: from Sample 12 (F7-35-OSF-Batch1-DVS-BAA-welD6-num42) of 20210507.wiff (Turbo Spr... Max. 2.7e9 cps.

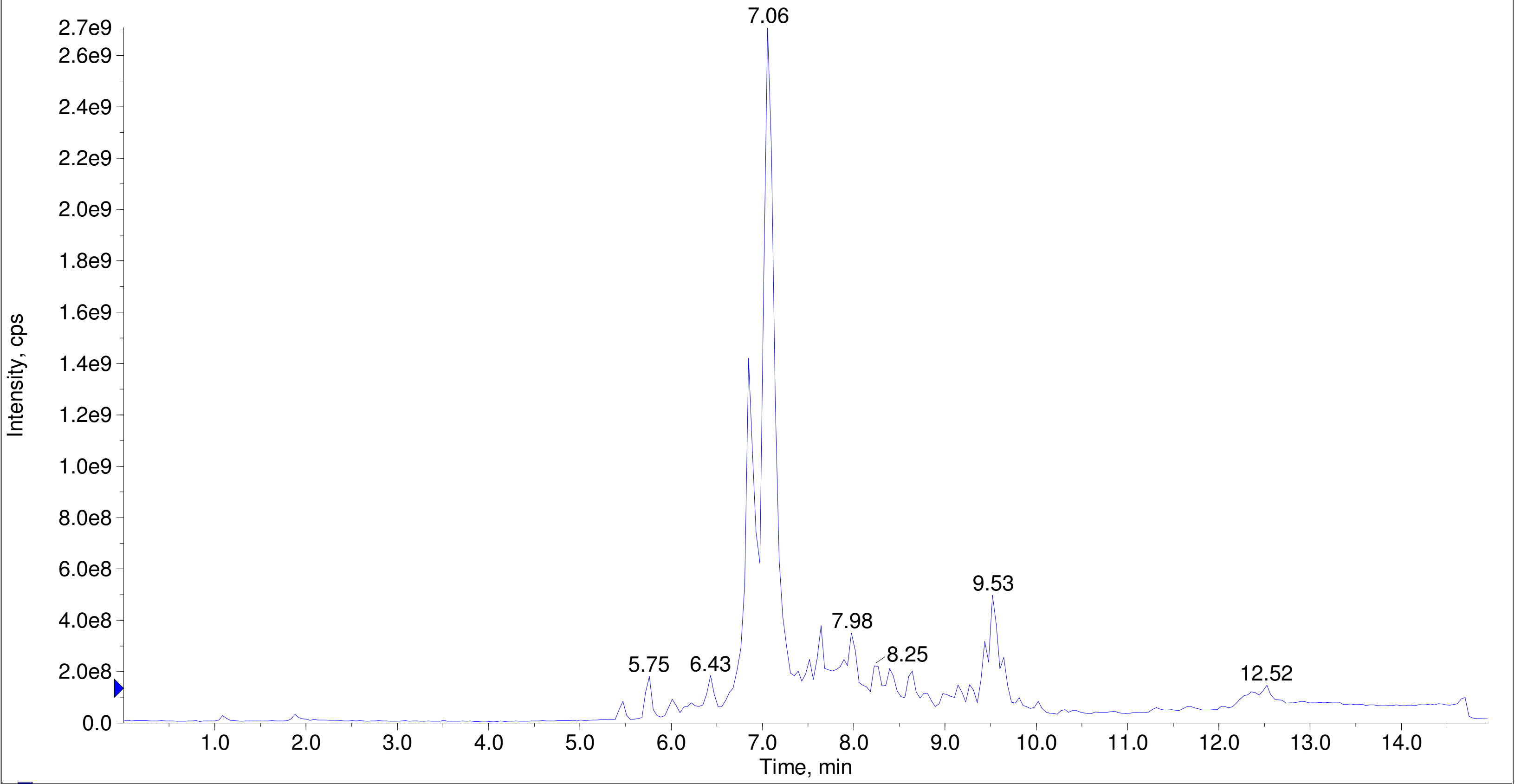


A/D Converter Channel 1 from Sample 12 (F7-35-OSF-Batch1-DVS-BAA-welD6-num42) of 20210507.wiff Max. 10.0 %.

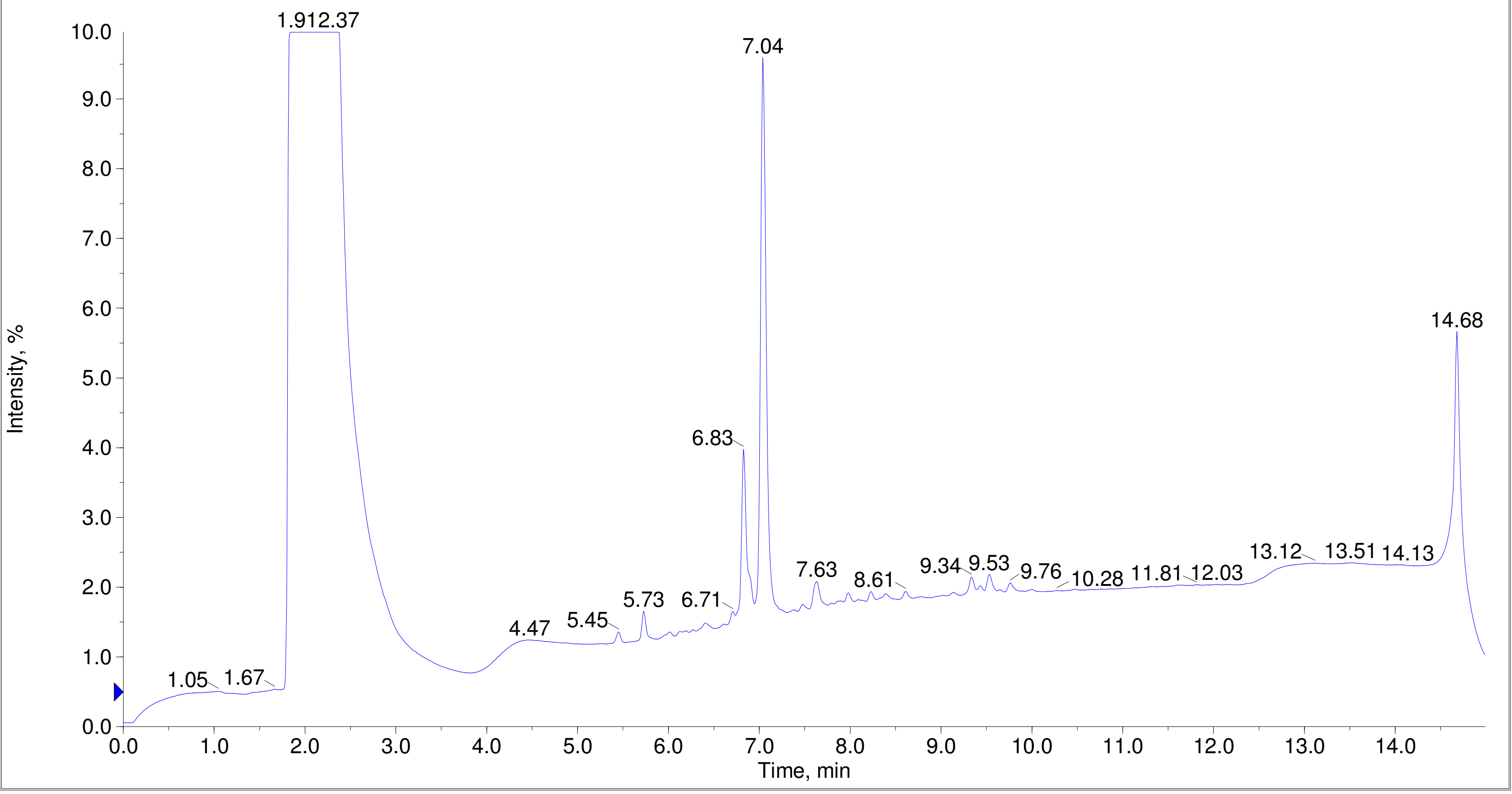

Printing Time: 7:37:50 PM  Post PM  Polarity/Scan Type: Positive Q1 MS
Printing Date: Monday, January 30, 2023  Sample Name:
Service Engineer Chris Maier  *Stanford U  *API 3000 SN D7640012

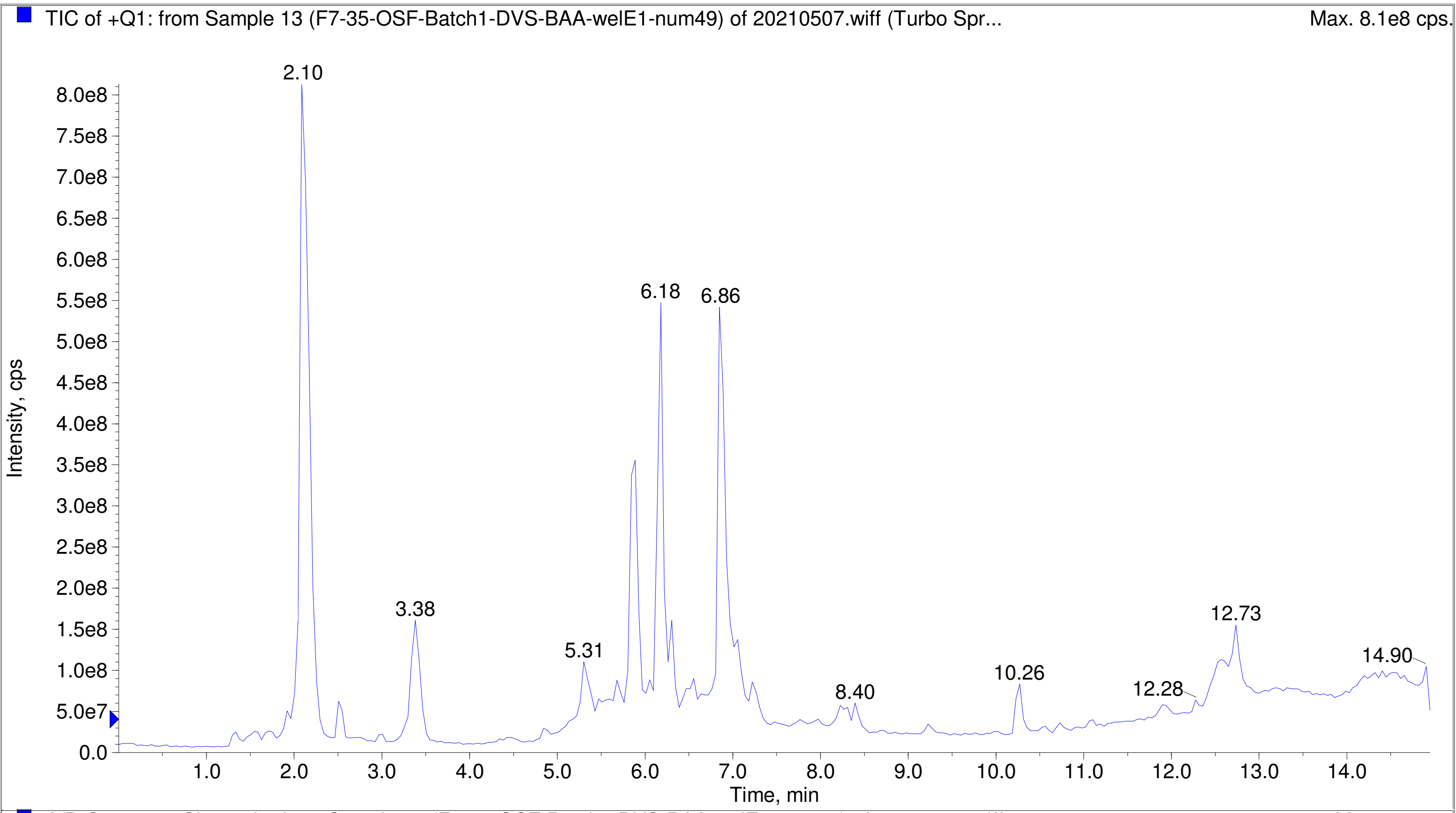


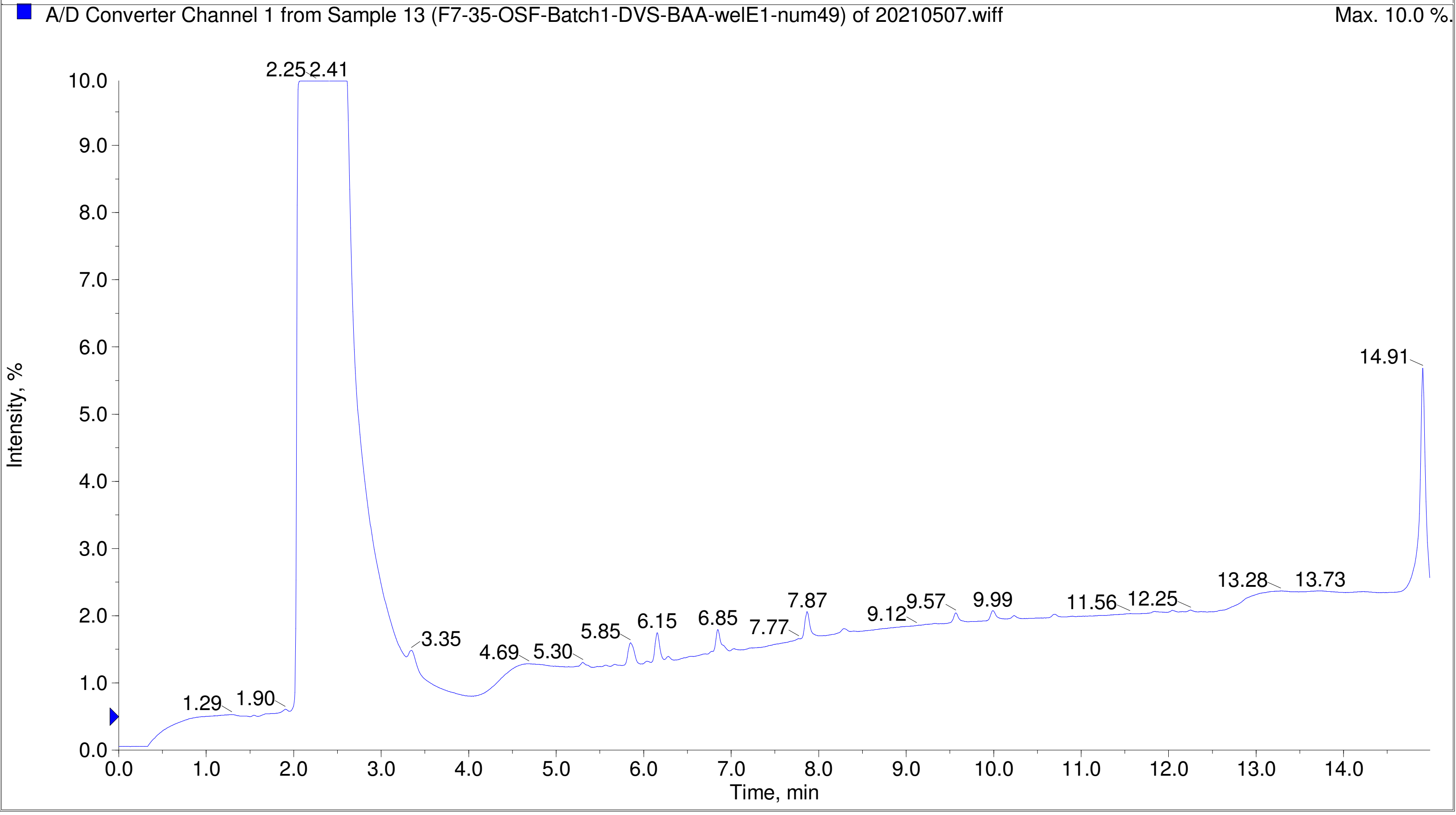

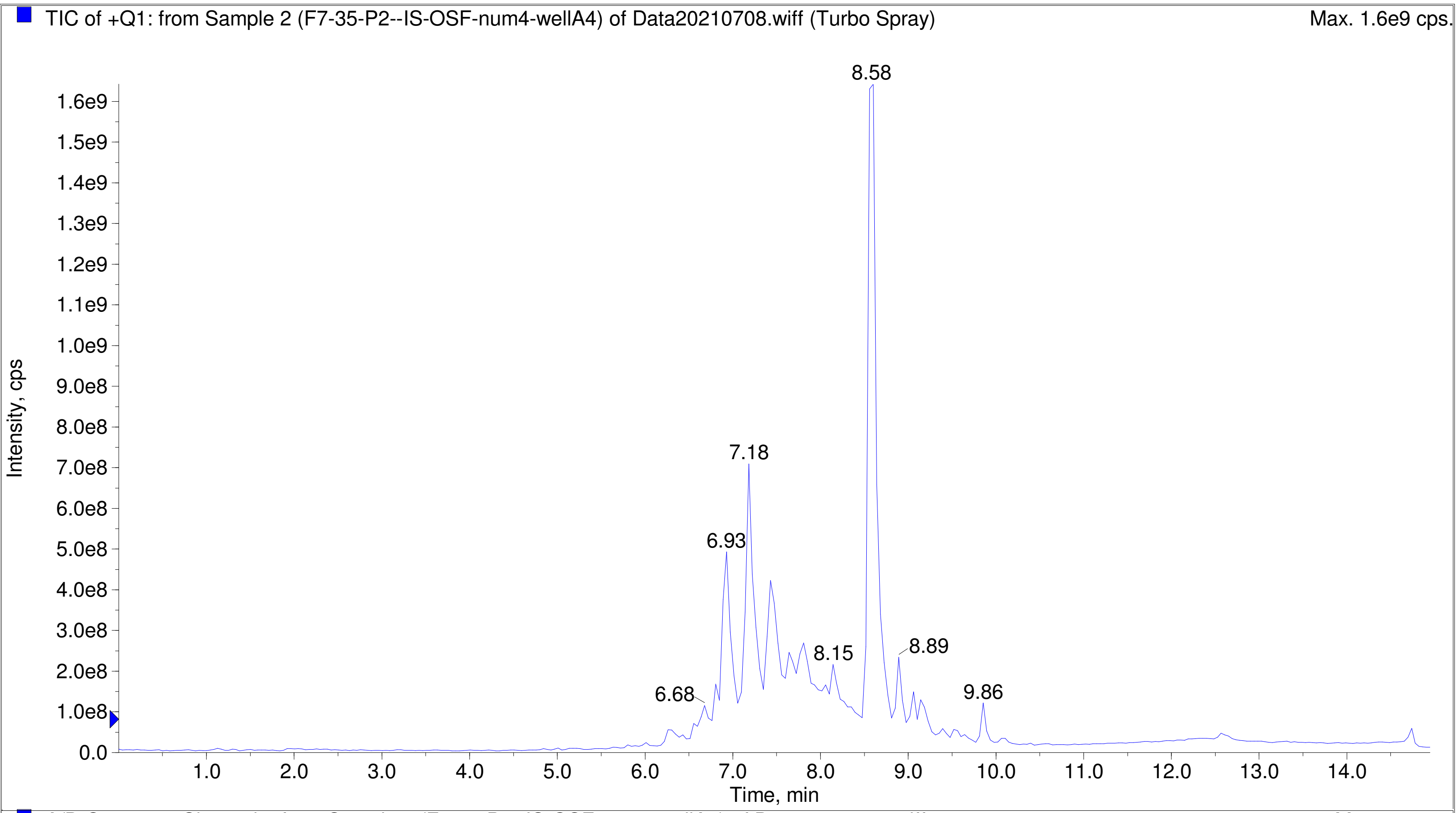
TIC of +Q1: from Sample 2 (F7-35-P2--IS-OSF-num4-wellA4) of Data20210708.wiff (Turbo Spray)
Max. 1.6e9 cps.
Intensity, cps
Time, min
8.58
7.18
6.93
8.15
8.89
6.68
9.86

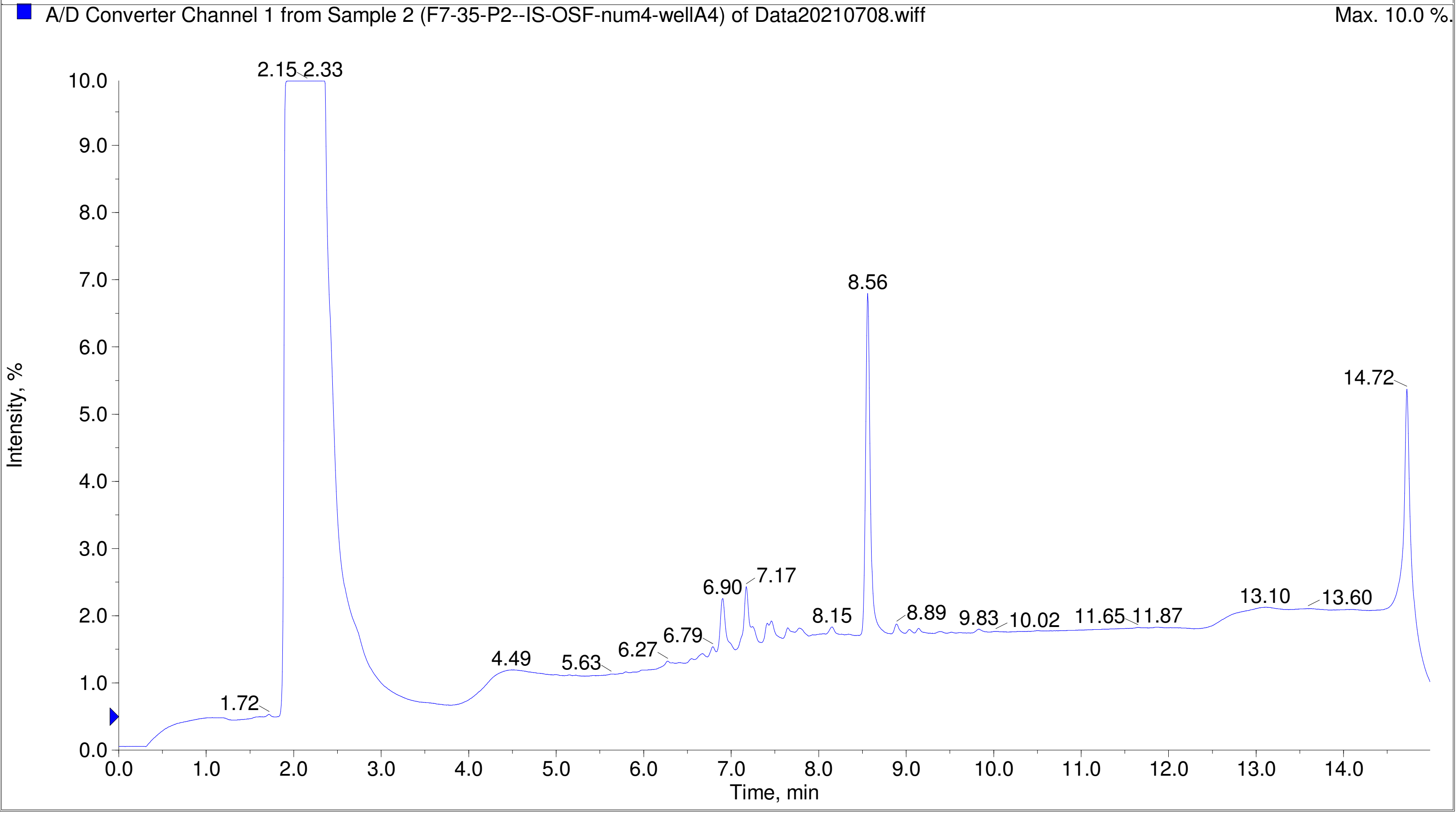
A/D Converter Channel 1 from Sample 2 (F7-35-P2--IS-OSF-num4-wellA4) of Data20210708.wiff
Max. 10.0 %.
Intensity, %
Time, min
2.15
2.33
8.56
14.72
6.90
7.17
8.15
8.89
9.83
10.02
11.65
11.87
13.10
13.60
6.79
6.27
4.49
5.63
1.72

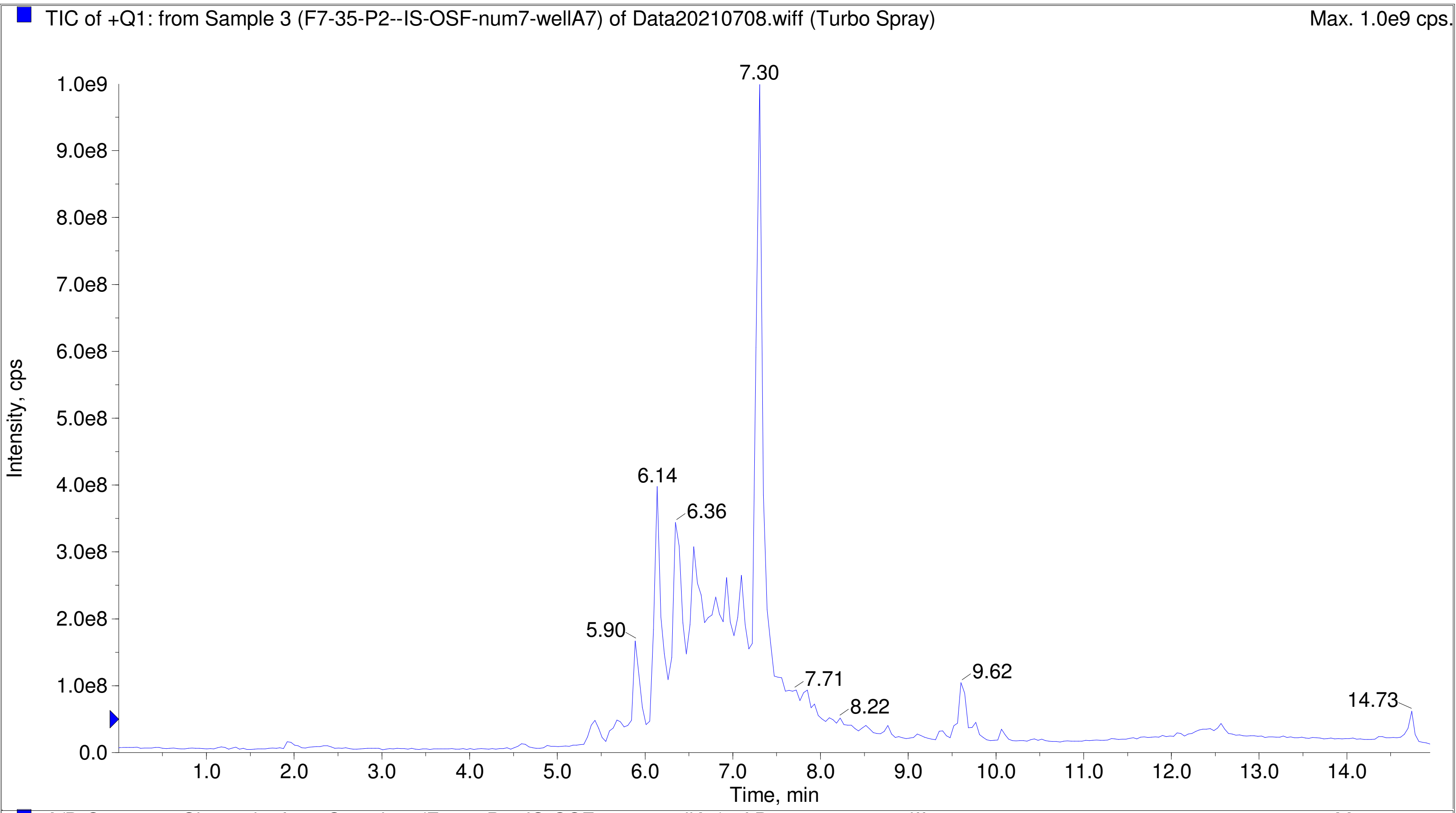
TIC of +Q1: from Sample 3 (F7-35-P2--IS-OSF-num7-wellA7) of Data20210708.wiff (Turbo Spray)
Max. 1.0e9 cps.
Intensity, cps
Time, min
7.30
6.14
6.36
5.90
7.71
8.22
9.62
14.73

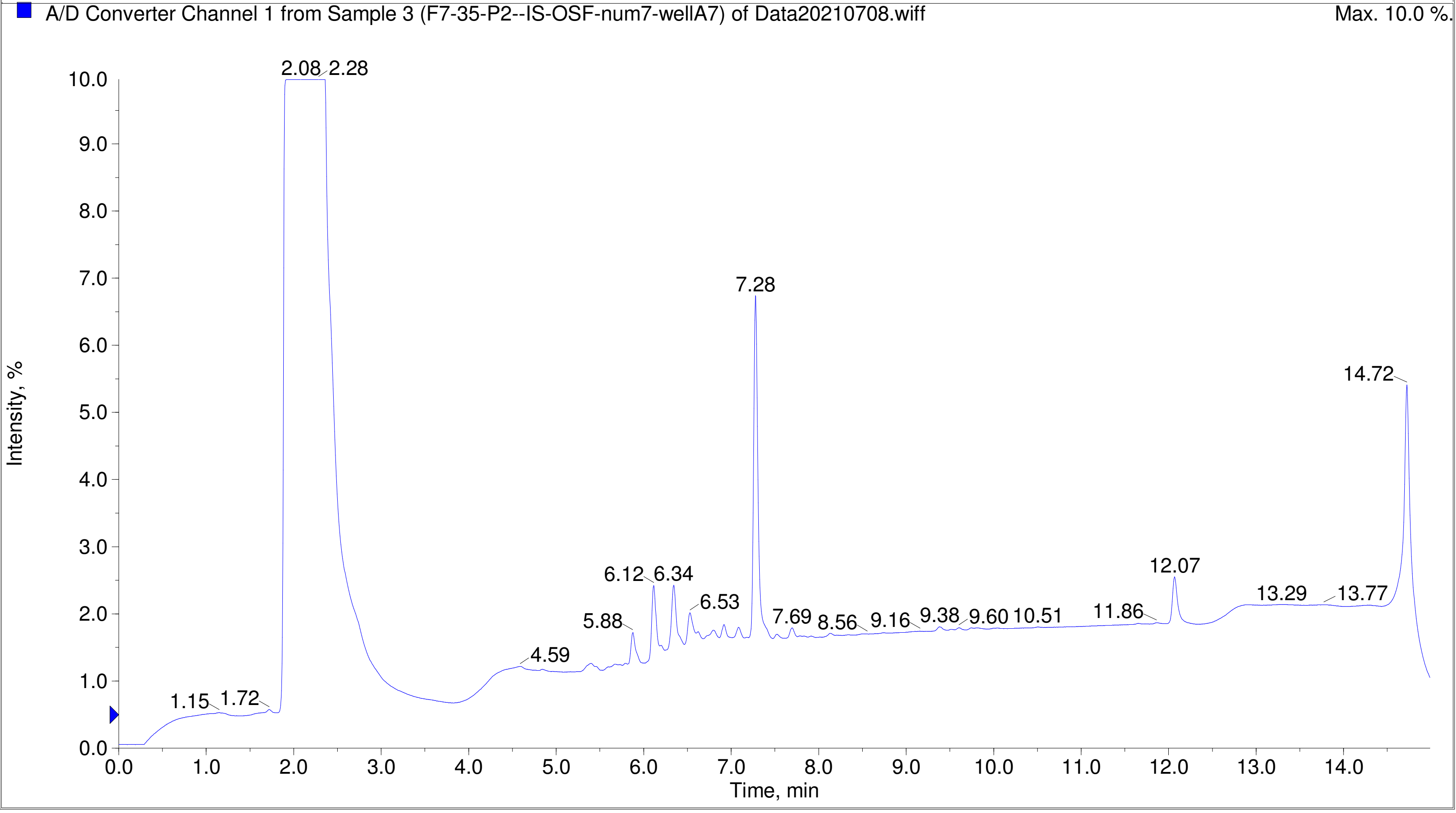
A/D Converter Channel 1 from Sample 3 (F7-35-P2--IS-OSF-num7-wellA7) of Data20210708.wiff
Max. 10.0 %.
Intensity, %
Time, min
2.08
2.28
7.28
14.72
6.12
6.34
6.53
5.88
4.59
1.15
1.72
7.69
8.56
9.16
9.38
9.60
10.51
11.86
12.07
13.29
13.77

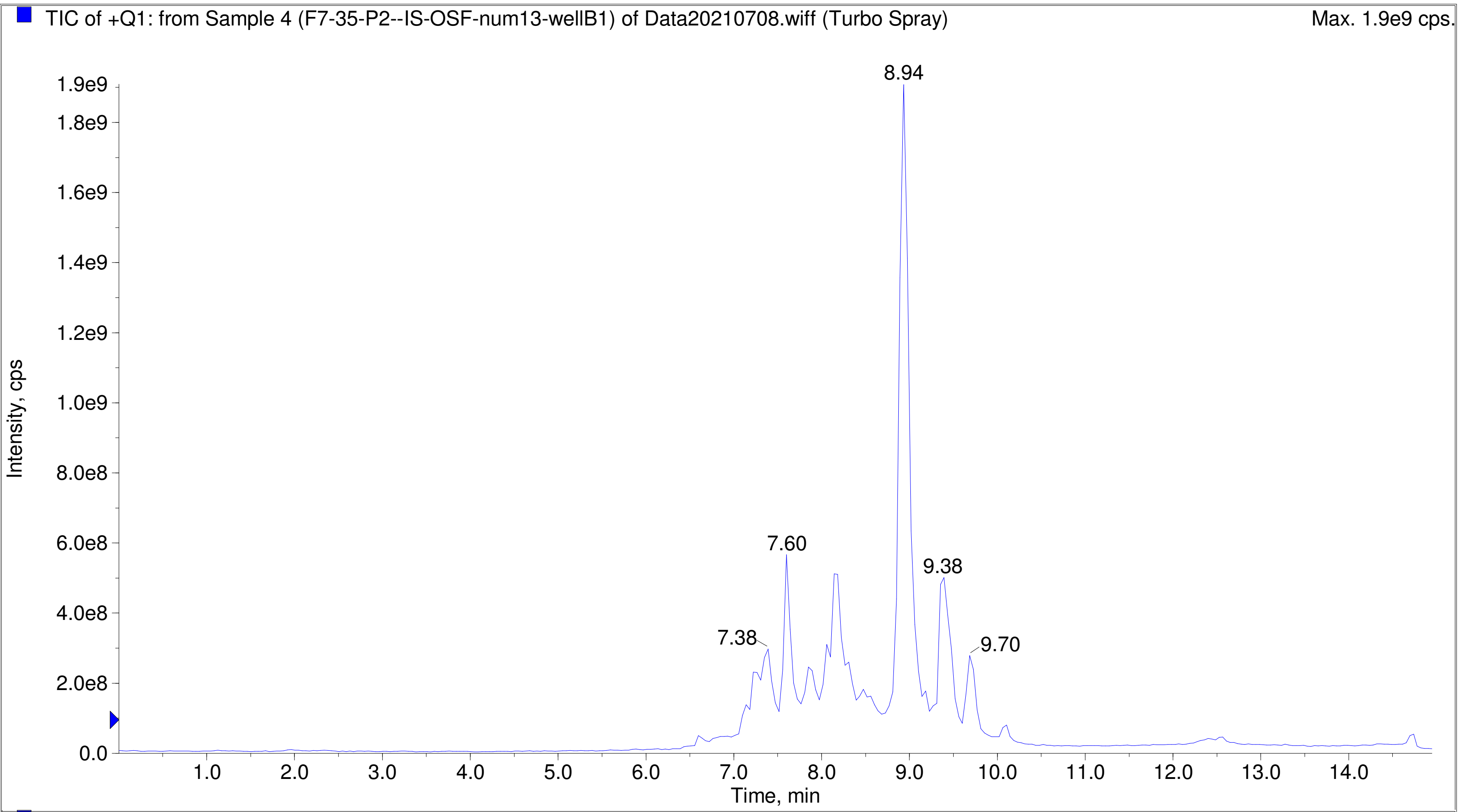
TIC of +Q1: from Sample 4 (F7-35-P2--IS-OSF-num13-wellB1) of Data20210708.wiff (Turbo Spray)
Max. 1.9e9 cps.
Intensity, cps
Time, min
8.94
7.60
9.38
7.38
9.70

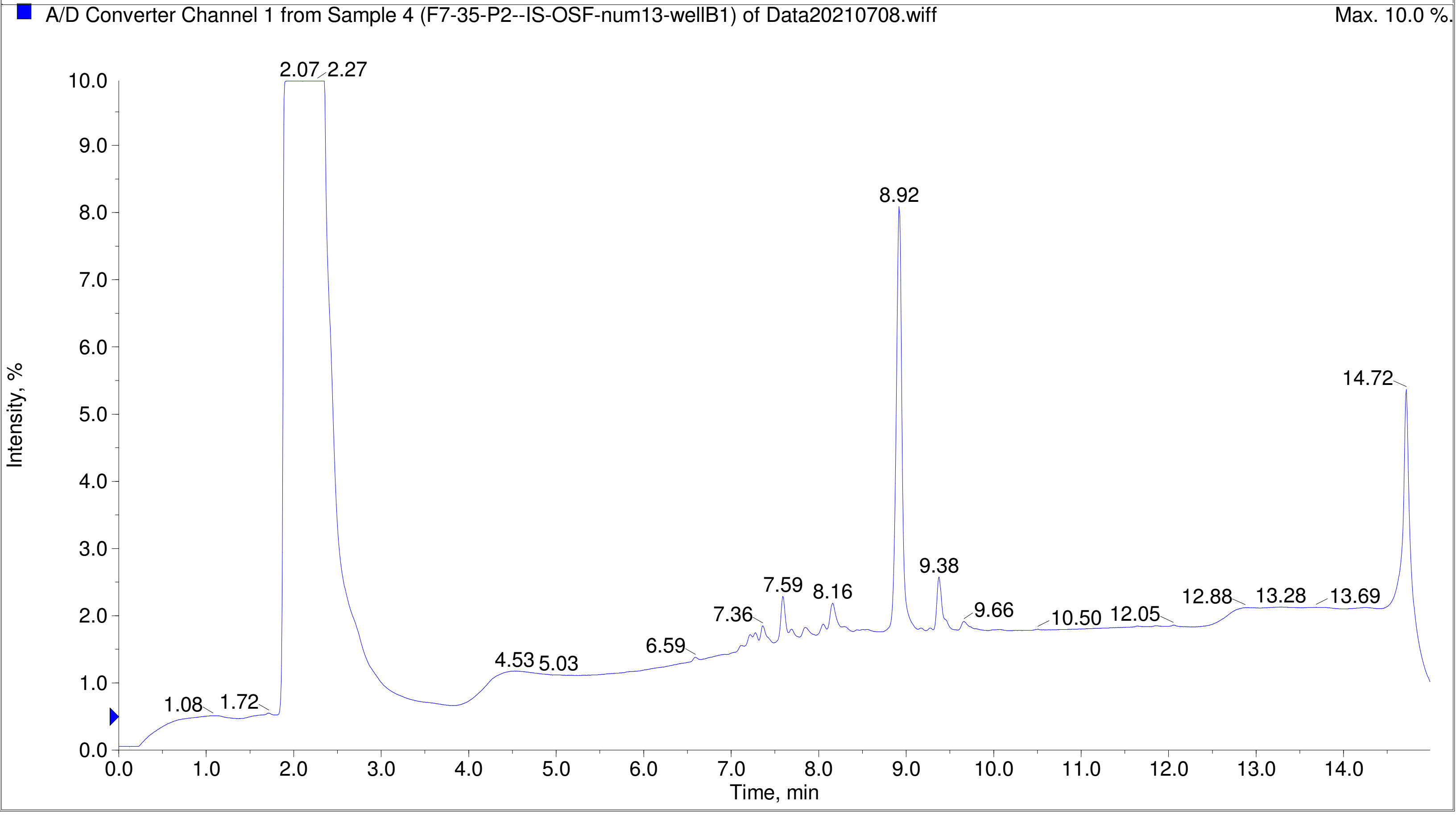
A/D Converter Channel 1 from Sample 4 (F7-35-P2--IS-OSF-num13-wellB1) of Data20210708.wiff
Max. 10.0 %.
Intensity, %
Time, min
2.07
2.27
8.92
14.72
9.38
7.59
8.16
7.36
9.66
10.50
12.05
12.88
13.28
13.69
6.59
4.53
5.03
1.08
1.72

Printing Time: 2:21:55 PM | Post PM | Polarity/Scan Type: Positive Q1 MS
Printing Date: Monday, January 30, 2023 | | Sample Name: F7-35-P2--IS-OSF-num15-wellB3
Service Engineer Chris Maier | *Stanford U | *API 3000 SN D7640012

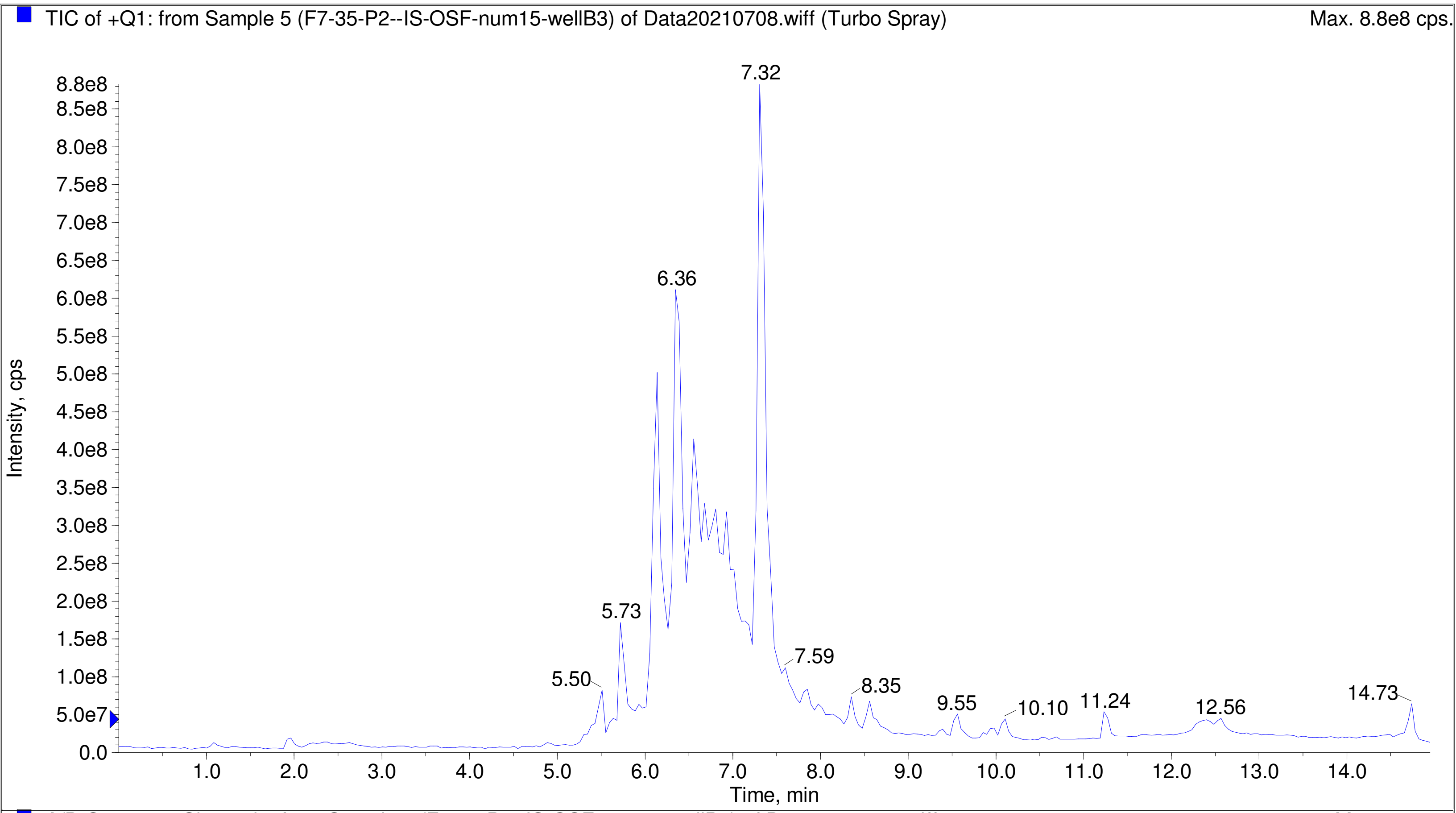


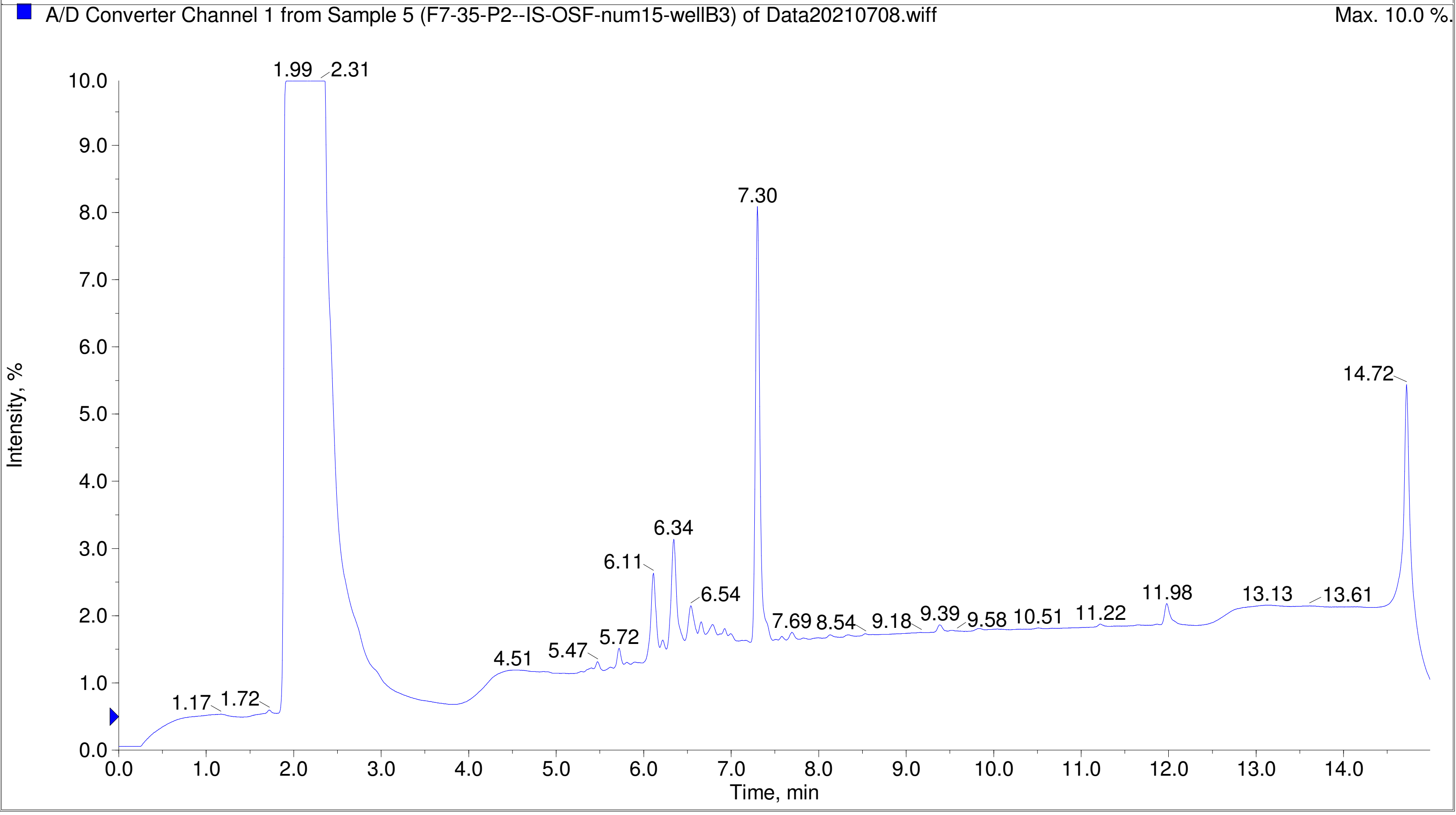

Printing Time: 2:22:44 PM    Post PM    Polarity/Scan Type: Positive Q1 MS
Printing Date: Monday, January 30, 2023    Sample Name: F7-35-P2--IS-OSF-num30-wellC6
Service Engineer Chris Maier    *Stanford U    *API 3000 SN D7640012

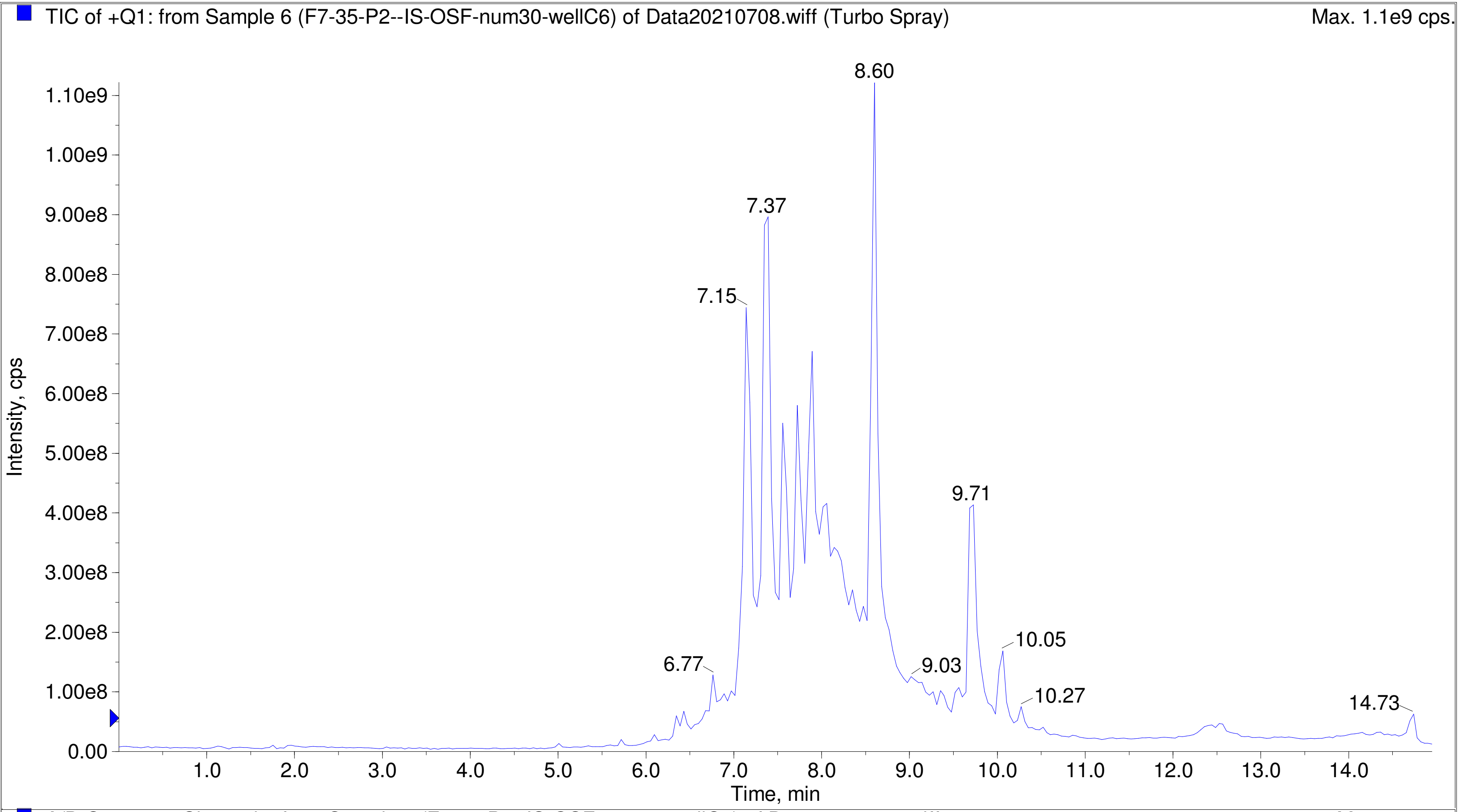


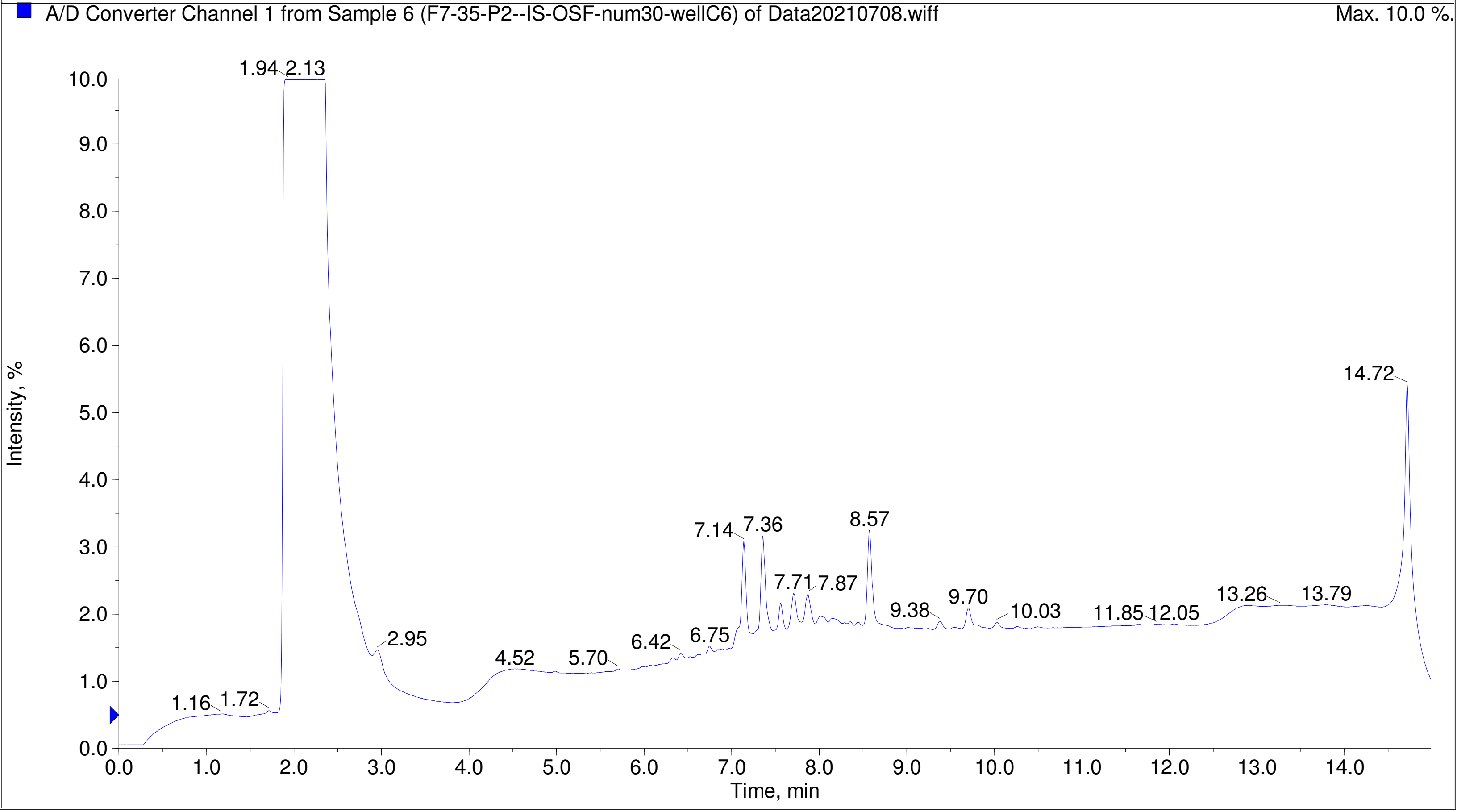

TIC of +Q1: from Sample 8 (F7-35-P2--IS-OSF-num32-wellC8) of Data20210708.wiff (Turbo Spray) Max. 1.0e9 cps.

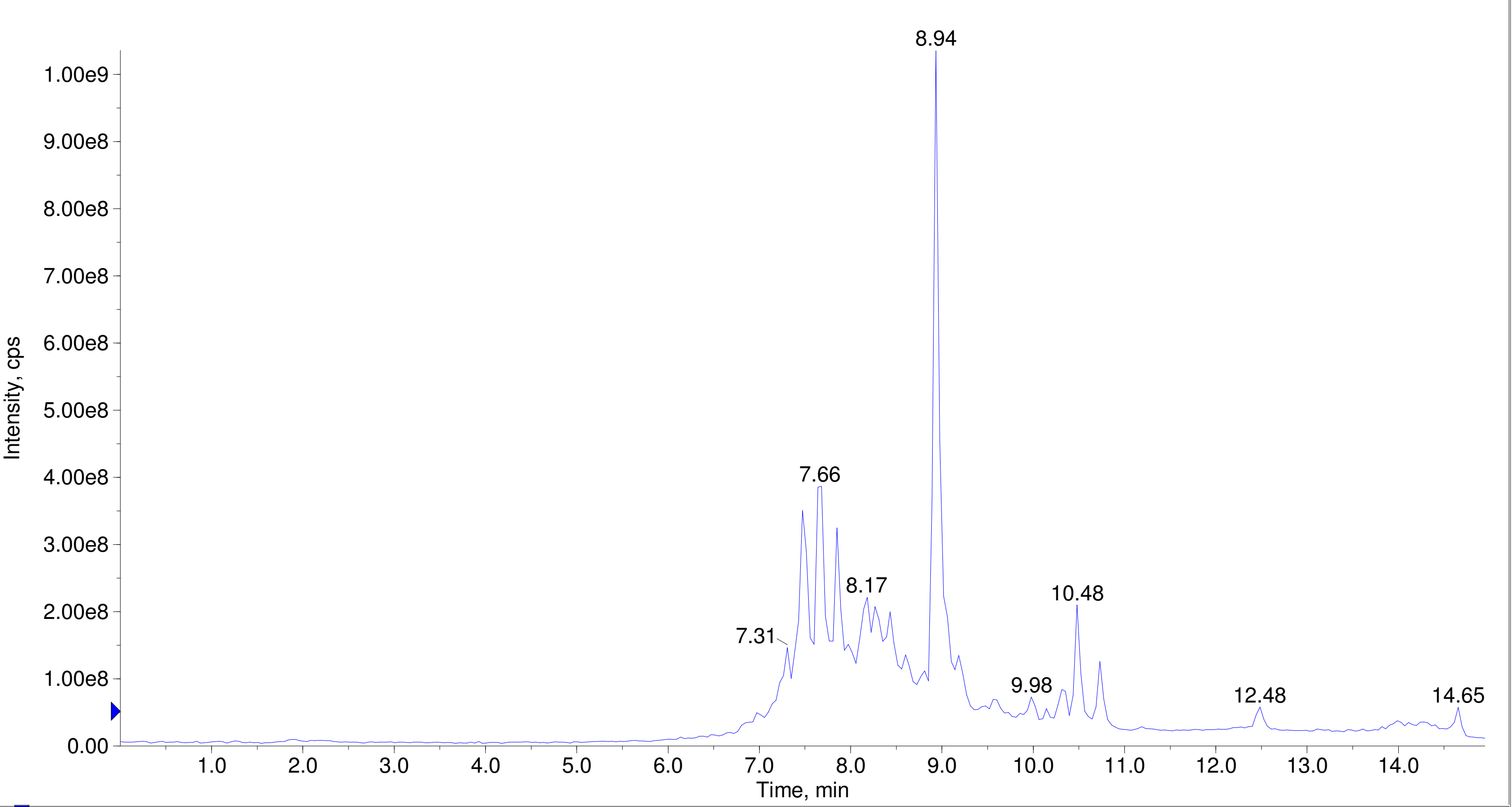


A/D Converter Channel 1 from Sample 8 (F7-35-P2--IS-OSF-num32-wellC8) of Data20210708.wiff Max. 10.0 %.

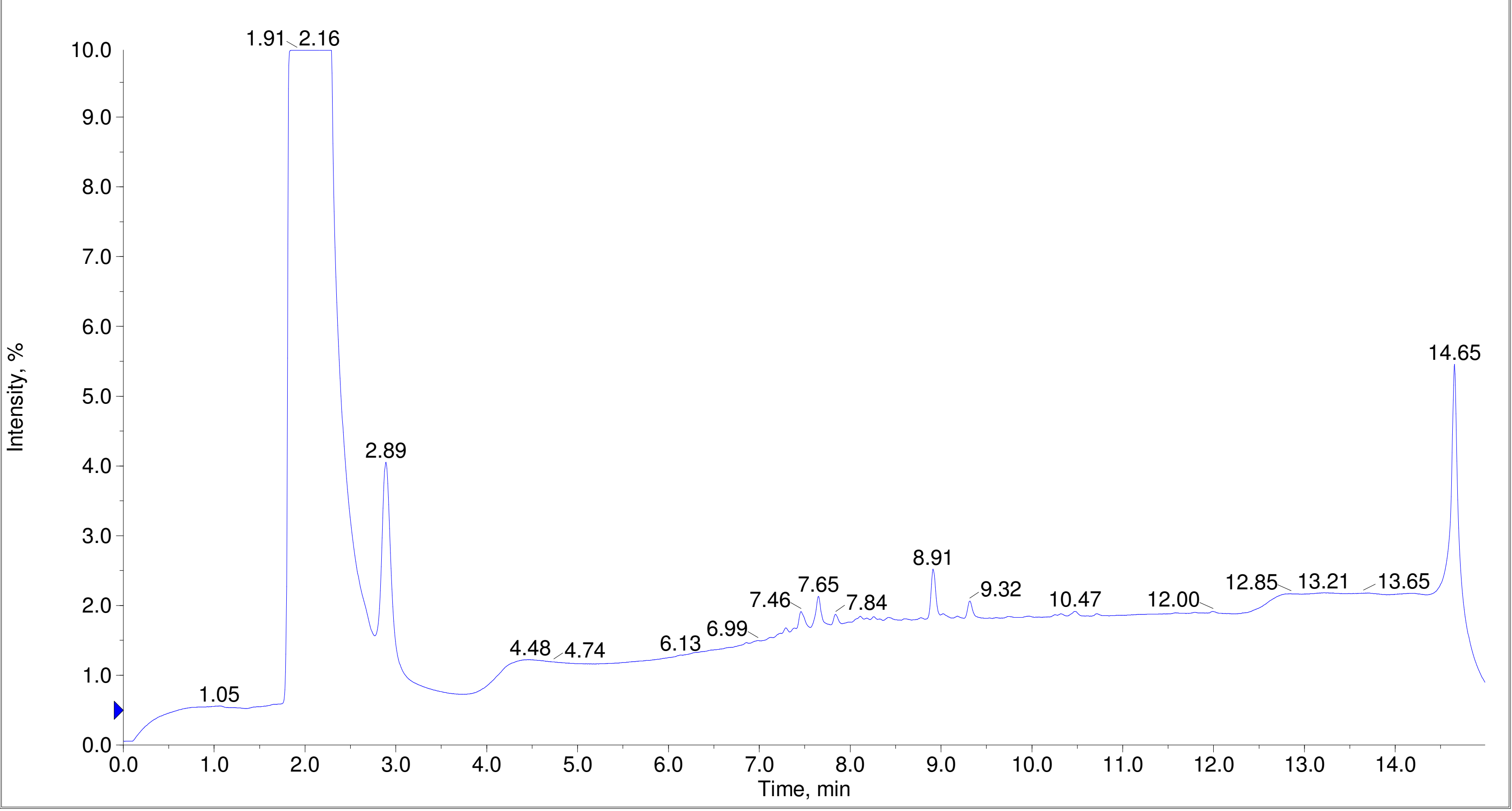

Printing Time: 3:44:57 PM
Printing Date: Tuesday, January 31, 2023
Service Engineer Chris Maier

Post PM

*Stanford U

Polarity/Scan Type: Positive Q1 MS
Sample Name: F7-35-P2--IS-OSF-num34-wellC10
*API 3000 SN D7640012

TIC of +Q1: from Sample 9 (F7-35-P2--IS-OSF-num34-wellC10) of Data20210708.wiff (Turbo Spray)    Max. 7.2e8 cps.

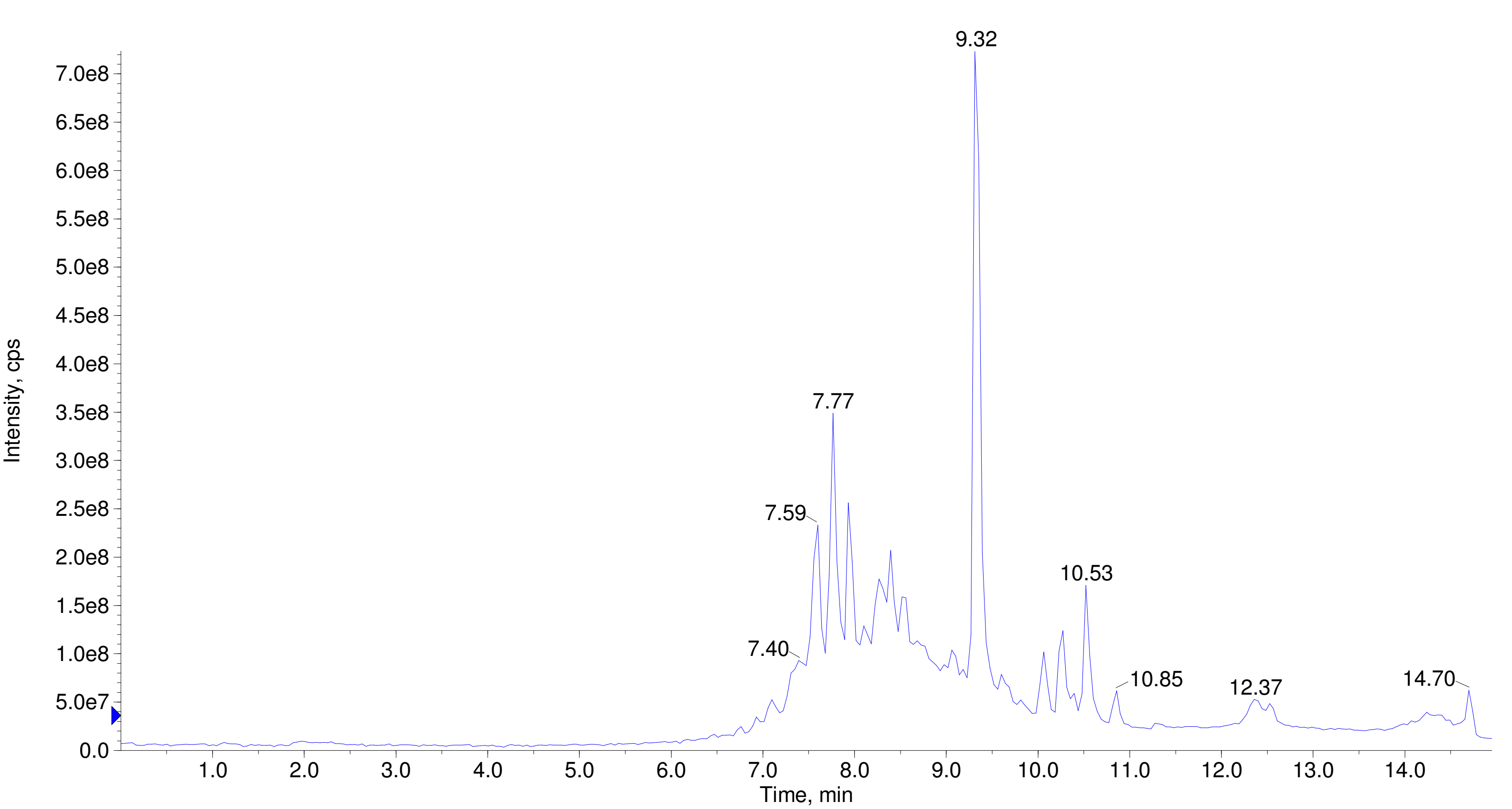


A/D Converter Channel 1 from Sample 9 (F7-35-P2--IS-OSF-num34-wellC10) of Data20210708.wiff    Max. 10.0 %.

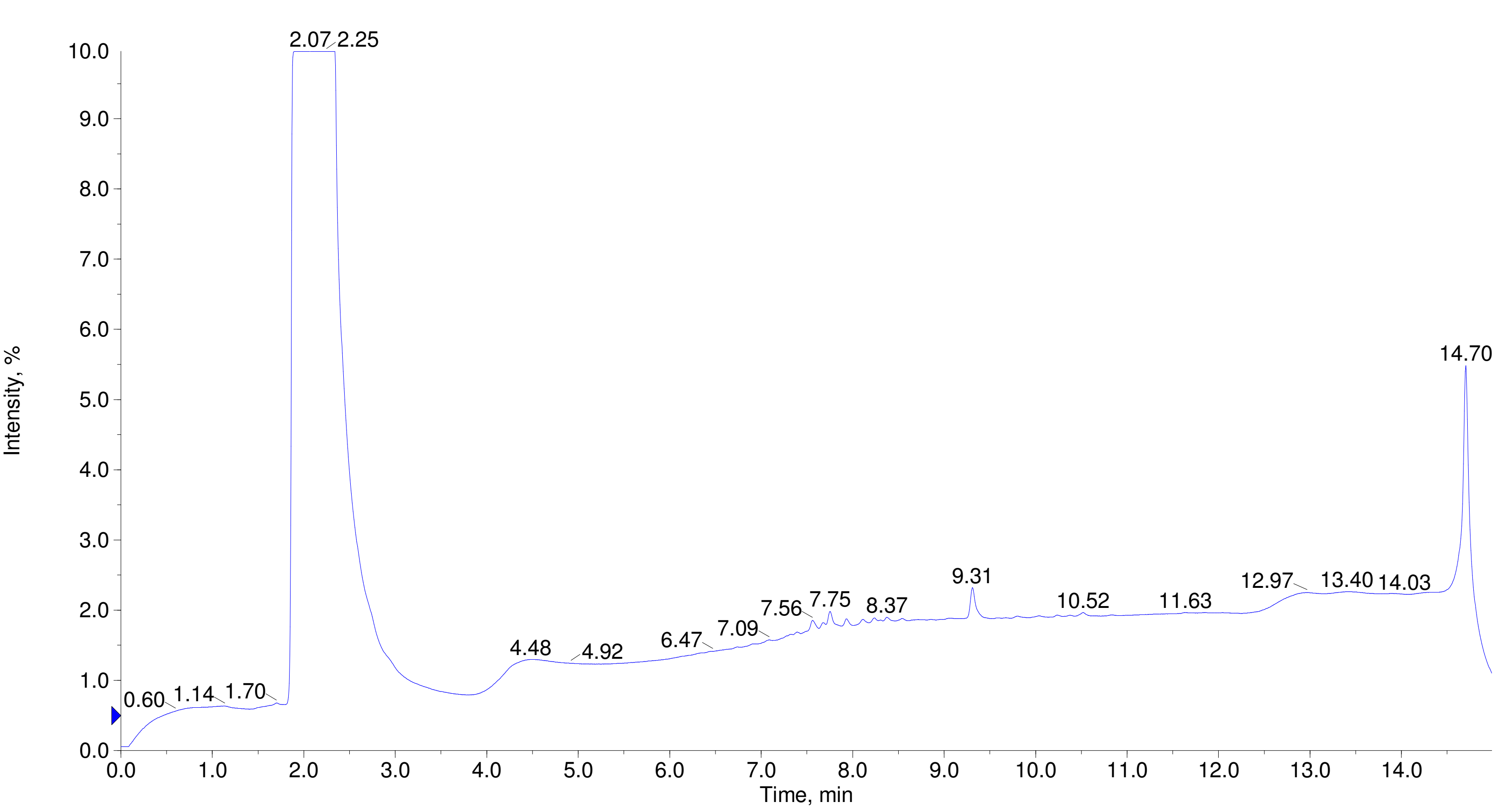

Printing Time: 3:46:21 PM
Printing Date: Tuesday, January 31, 2023
Service Engineer Chris Maier

Post PM
*Stanford U

Polarity/Scan Type: Positive Q1 MS
Sample Name: F7-35-P2--IS-OSF-num39-wellD3
*API 3000 SN D7640012

TIC of +Q1: from Sample 10 (F7-35-P2--IS-OSF-num39-wellD3) of Data20210708.wiff (Turbo Spray) Max. 7.7e8 cps.

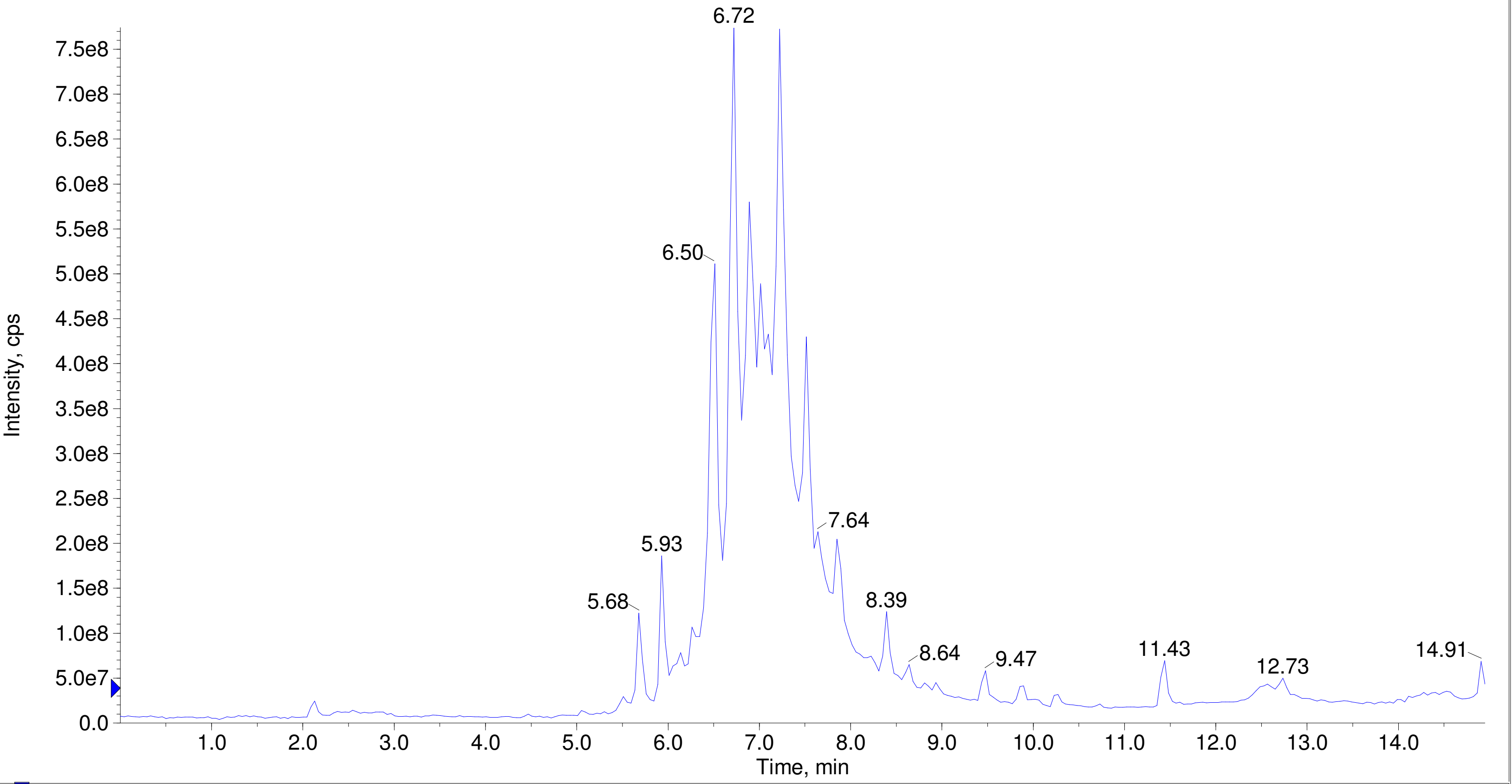


A/D Converter Channel 1 from Sample 10 (F7-35-P2--IS-OSF-num39-wellD3) of Data20210708.wiff Max. 10.0 %.

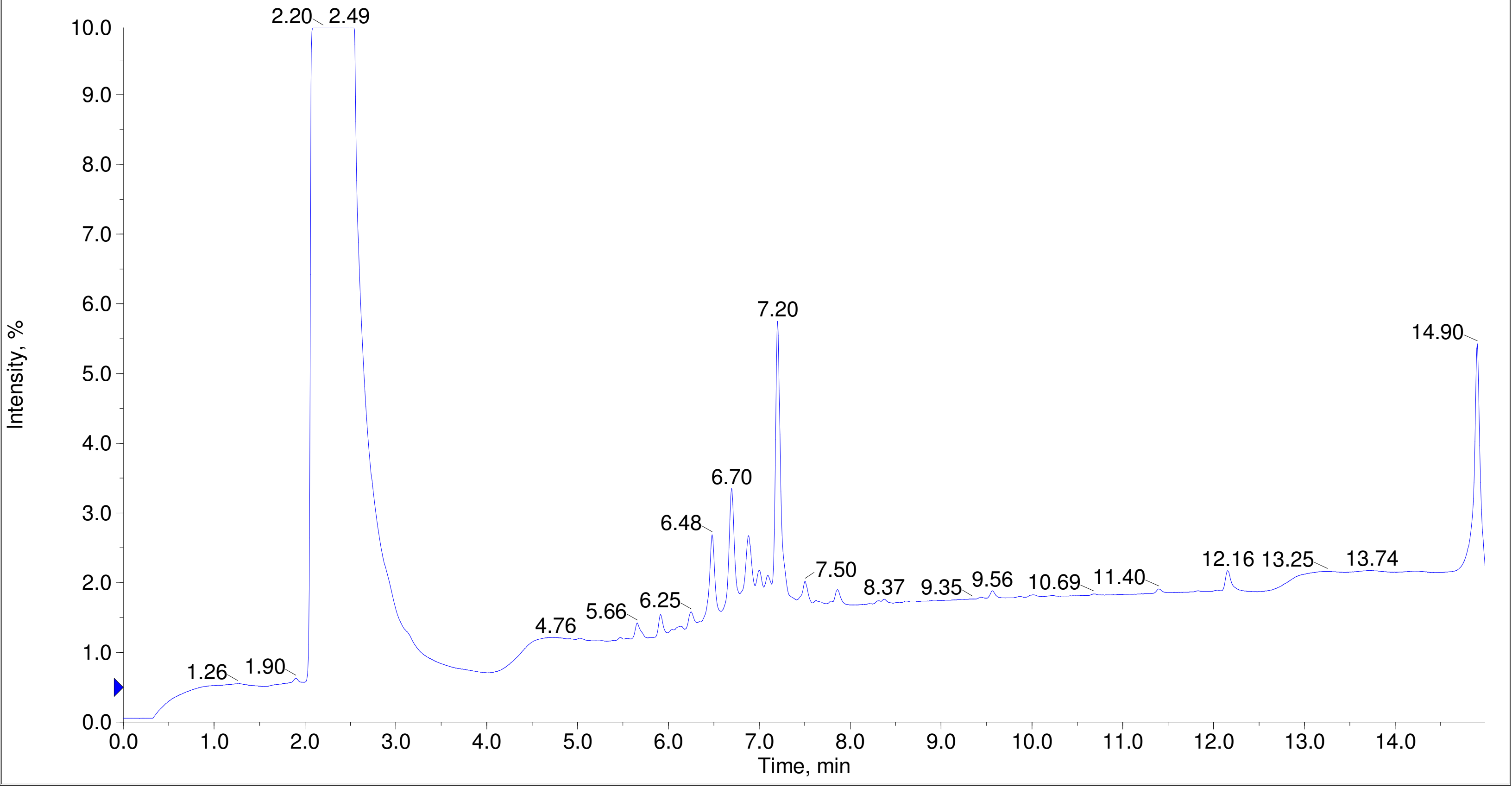

Printing Time: 3:47:26 PM | Post PM | Polarity/Scan Type: Positive Q1 MS
Printing Date: Tuesday, January 31, 2023 | | Sample Name: F7-35-P2--IS-OSF-num41-wellD5
Service Engineer Chris Maier | *Stanford U | *API 3000 SN D7640012

TIC of +Q1: from Sample 11 (F7-35-P2--IS-OSF-num41-wellD5) of Data20210708.wiff (Turbo Spray) Max. 5.1e8 cps.

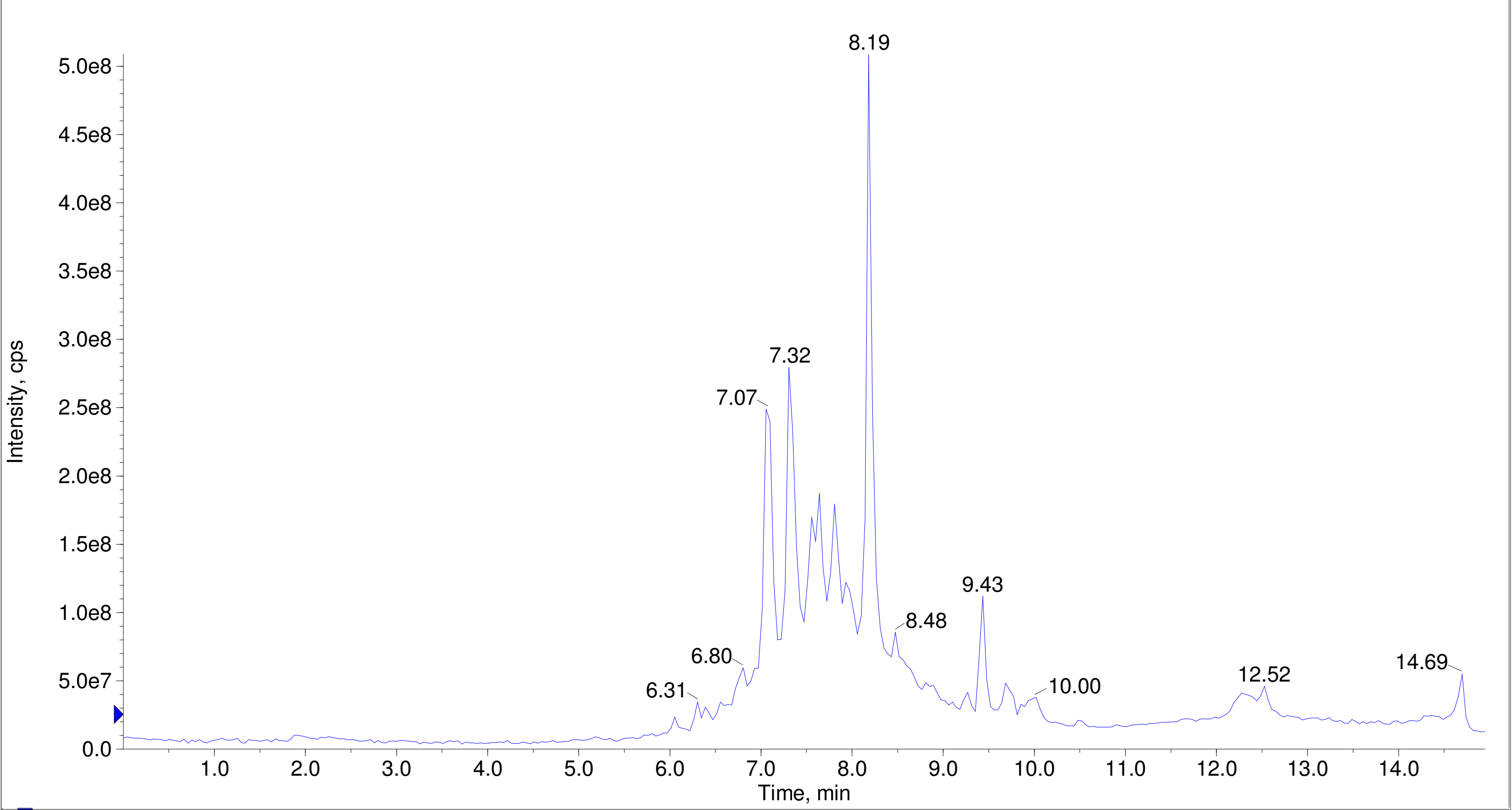


A/D Converter Channel 1 from Sample 11 (F7-35-P2--IS-OSF-num41-wellD5) of Data20210708.wiff Max. 10.0 %.

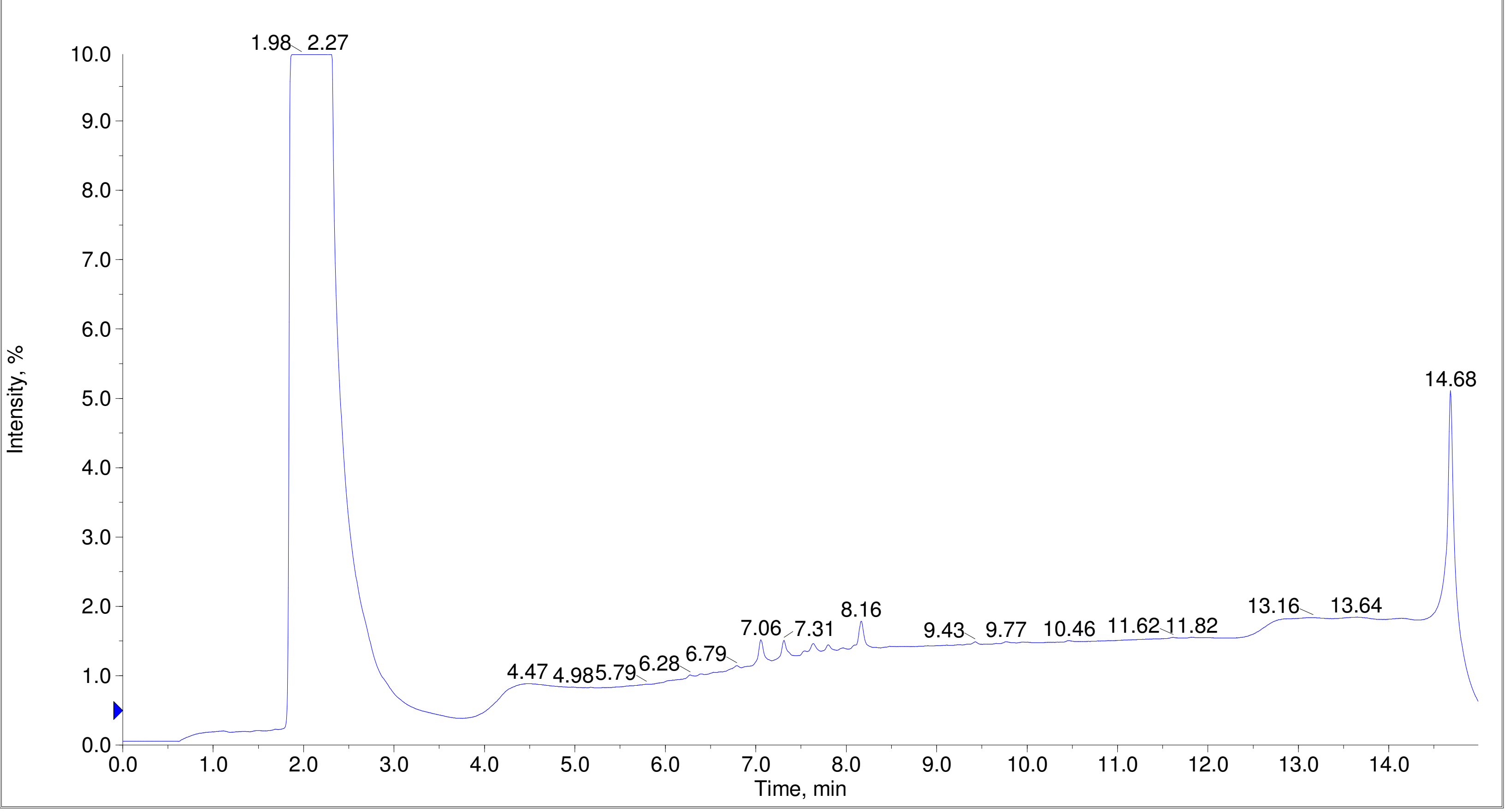

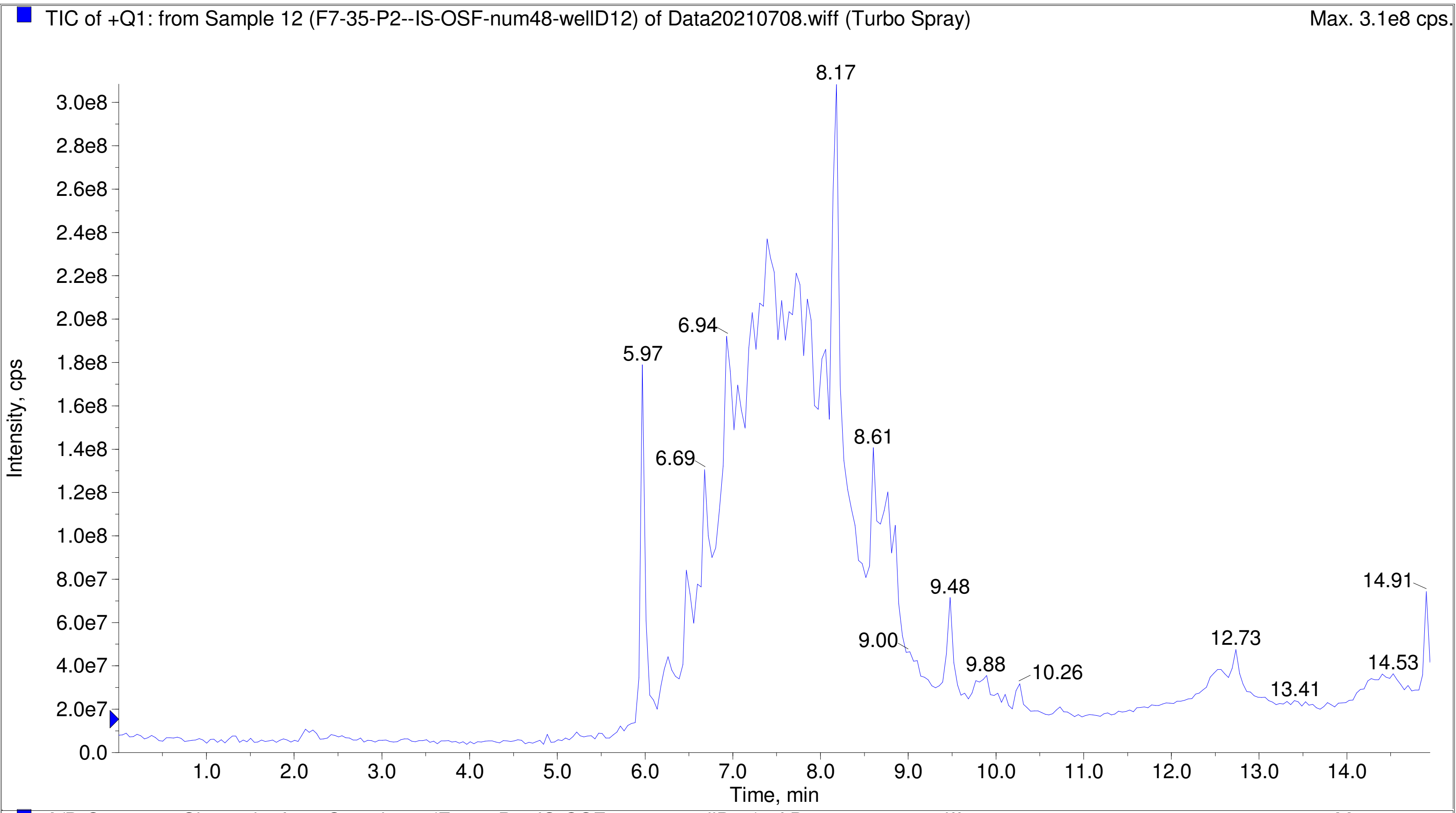
TIC of +Q1: from Sample 12 (F7-35-P2--IS-OSF-num48-wellD12) of Data20210708.wiff (Turbo Spray)
Max. 3.1e8 cps.
Intensity, cps
Time, min
8.17
6.94
5.97
6.69
8.61
9.48
9.00
9.88
10.26
12.73
13.41
14.91
14.53

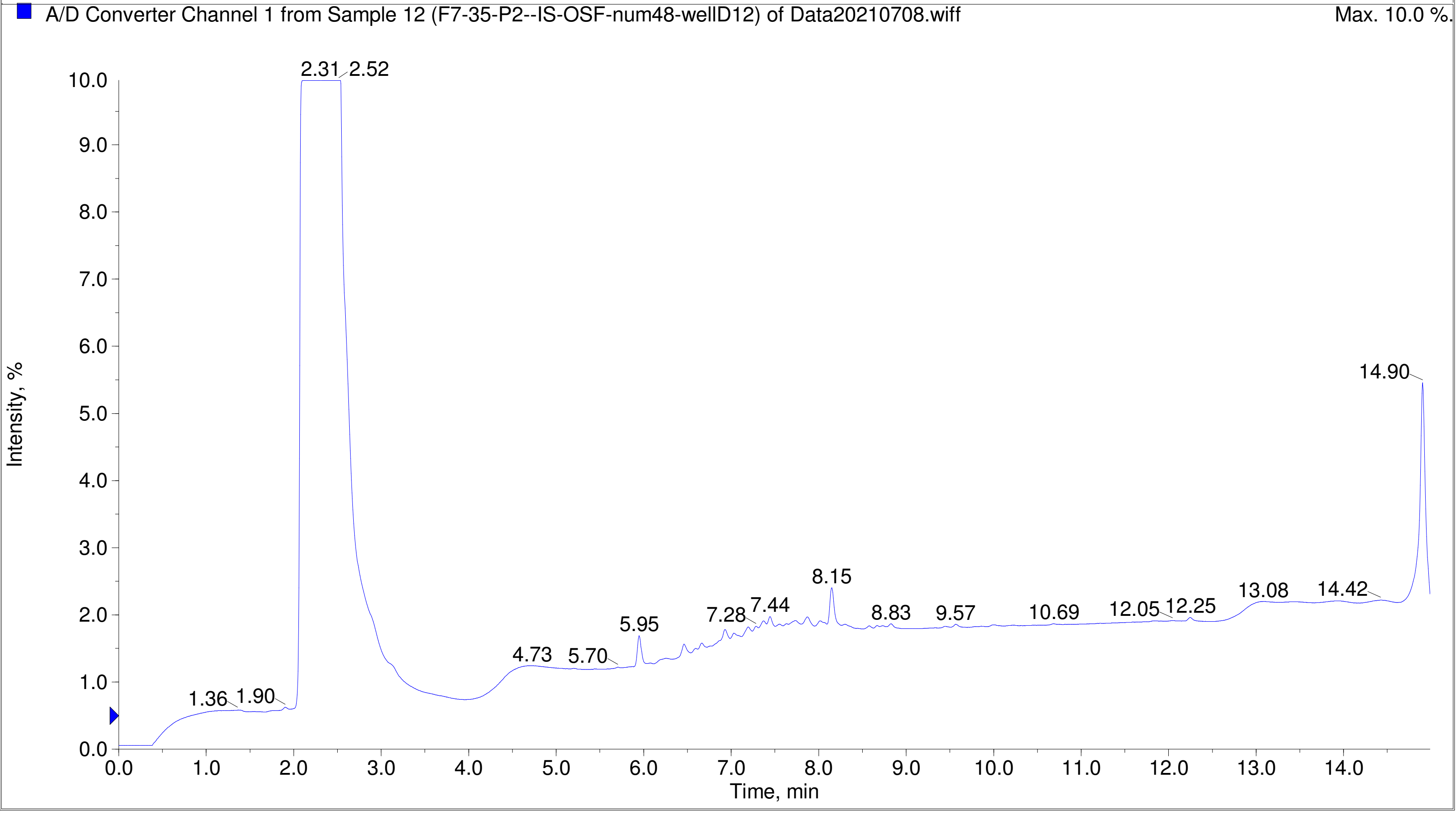
A/D Converter Channel 1 from Sample 12 (F7-35-P2--IS-OSF-num48-wellD12) of Data20210708.wiff
Max. 10.0 %.
Intensity, %
Time, min
2.31
2.52
1.36
1.90
4.73
5.70
5.95
7.28
7.44
8.15
8.83
9.57
10.69
12.05
12.25
13.08
14.42
14.90

Printing Time: 2:25:05 PM
Printing Date: Monday, January 30, 2023
Service Engineer Chris Maier

Post PM
*Stanford U

Polarity/Scan Type: Positive Q1 MS
Sample Name: F7-35-P3-JS-OSF-num2-A2
*API 3000 SN D7640012

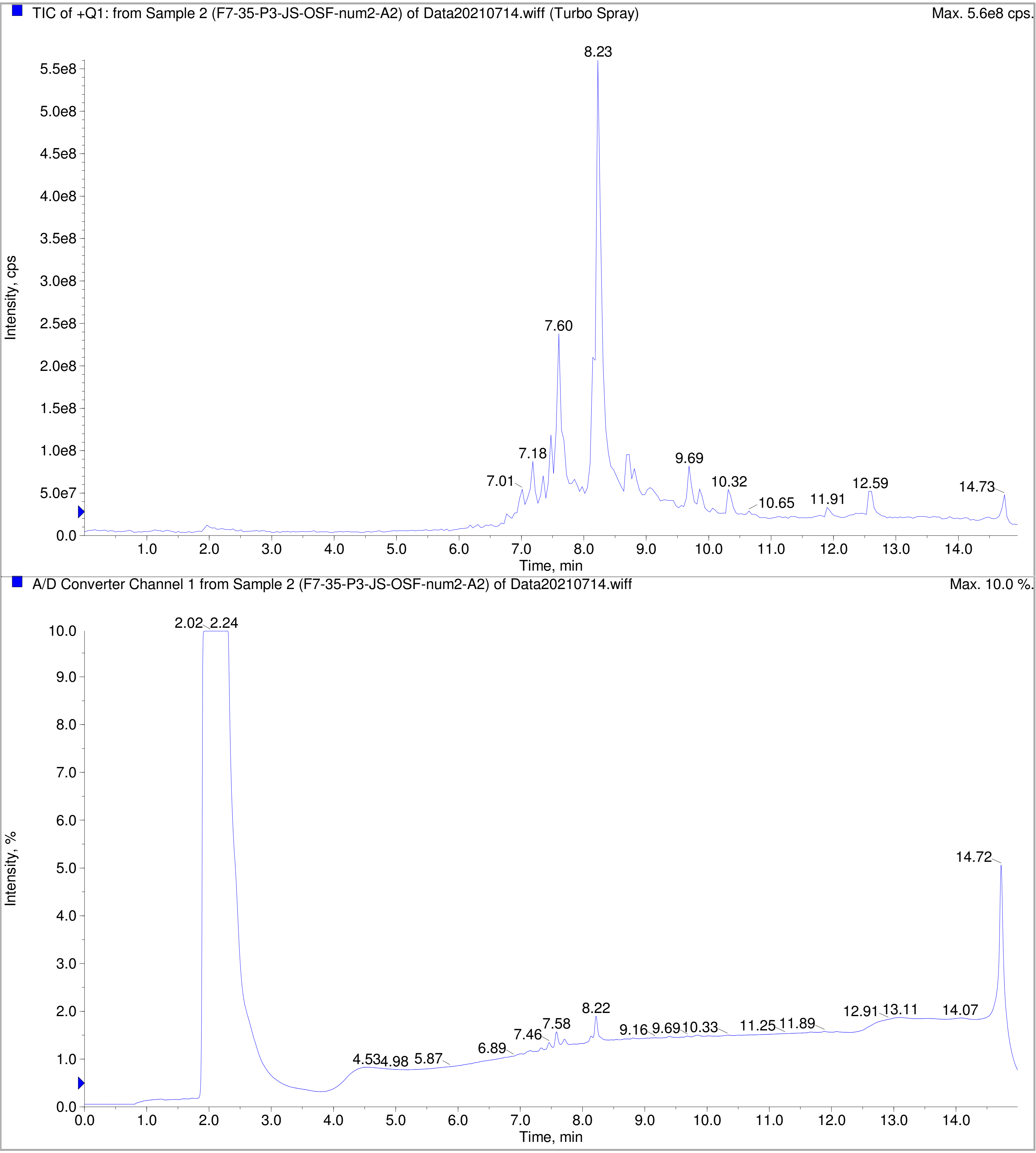

TIC of +Q1: from Sample 3 (F7-35-P3-JS-OSF-num5-A5) of Data20210714.wiff (Turbo Spray) Max. 2.7e8 cps.

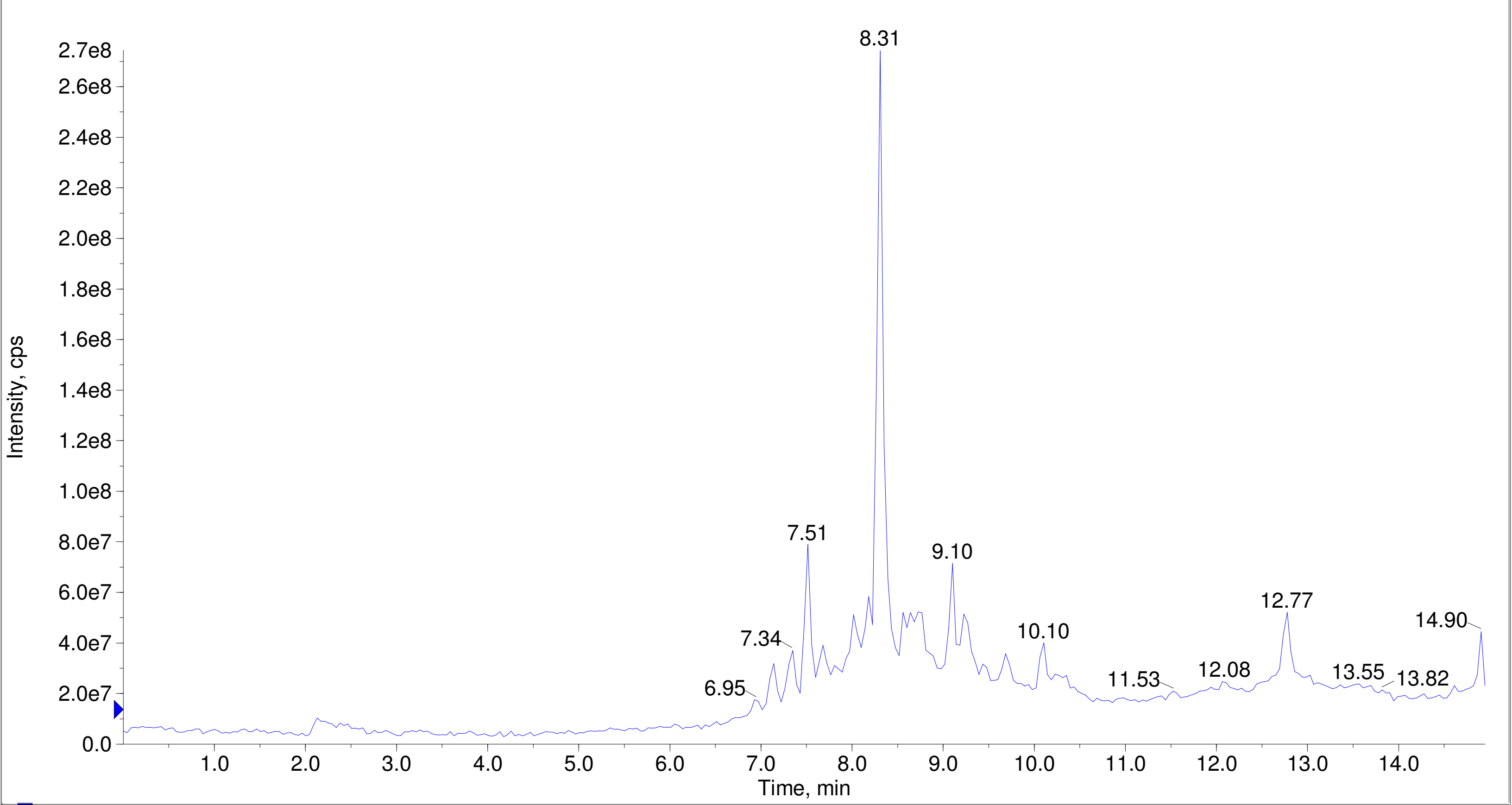


A/D Converter Channel 1 from Sample 3 (F7-35-P3-JS-OSF-num5-A5) of Data20210714.wiff Max. 10.0 %.

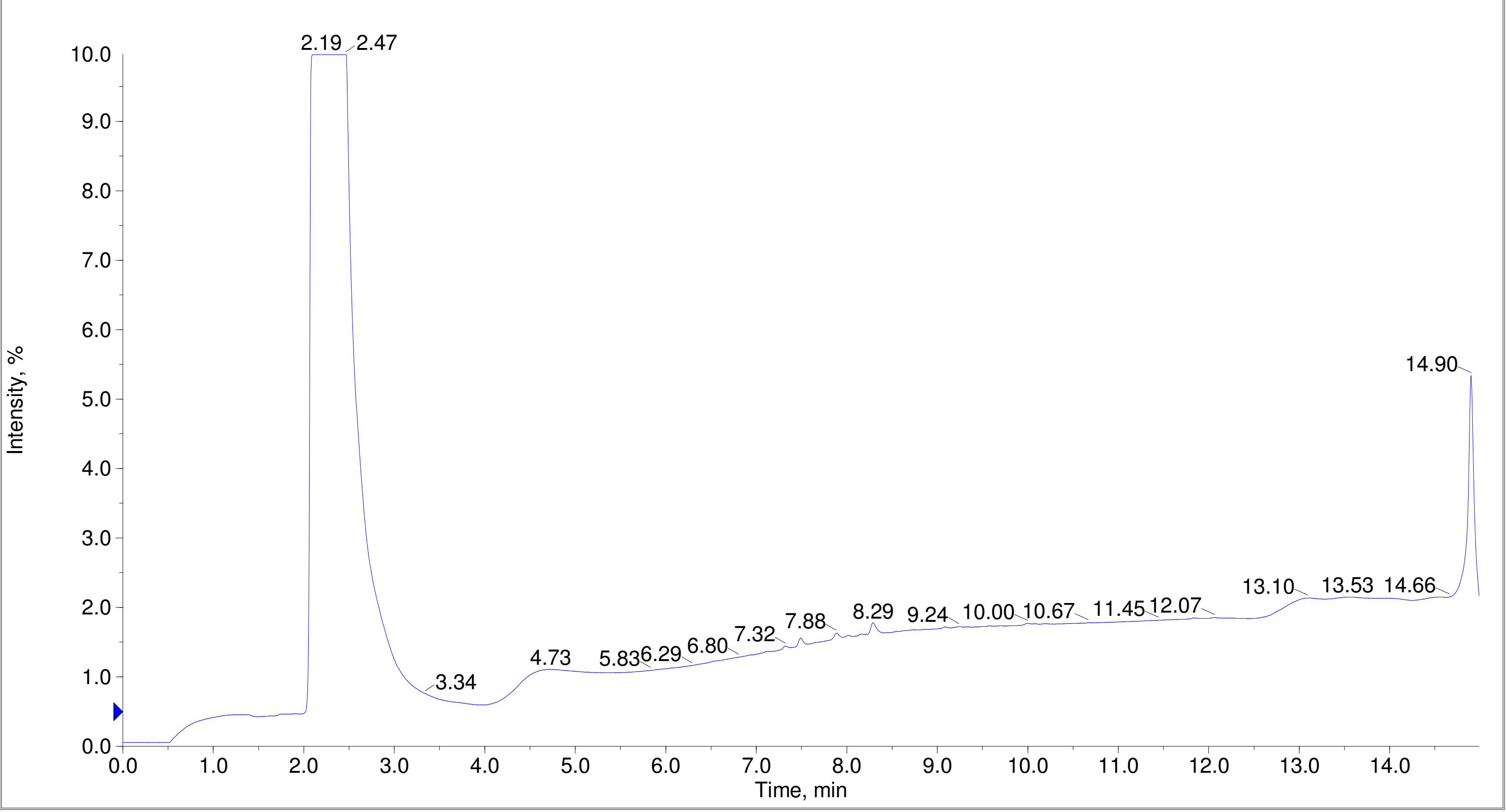

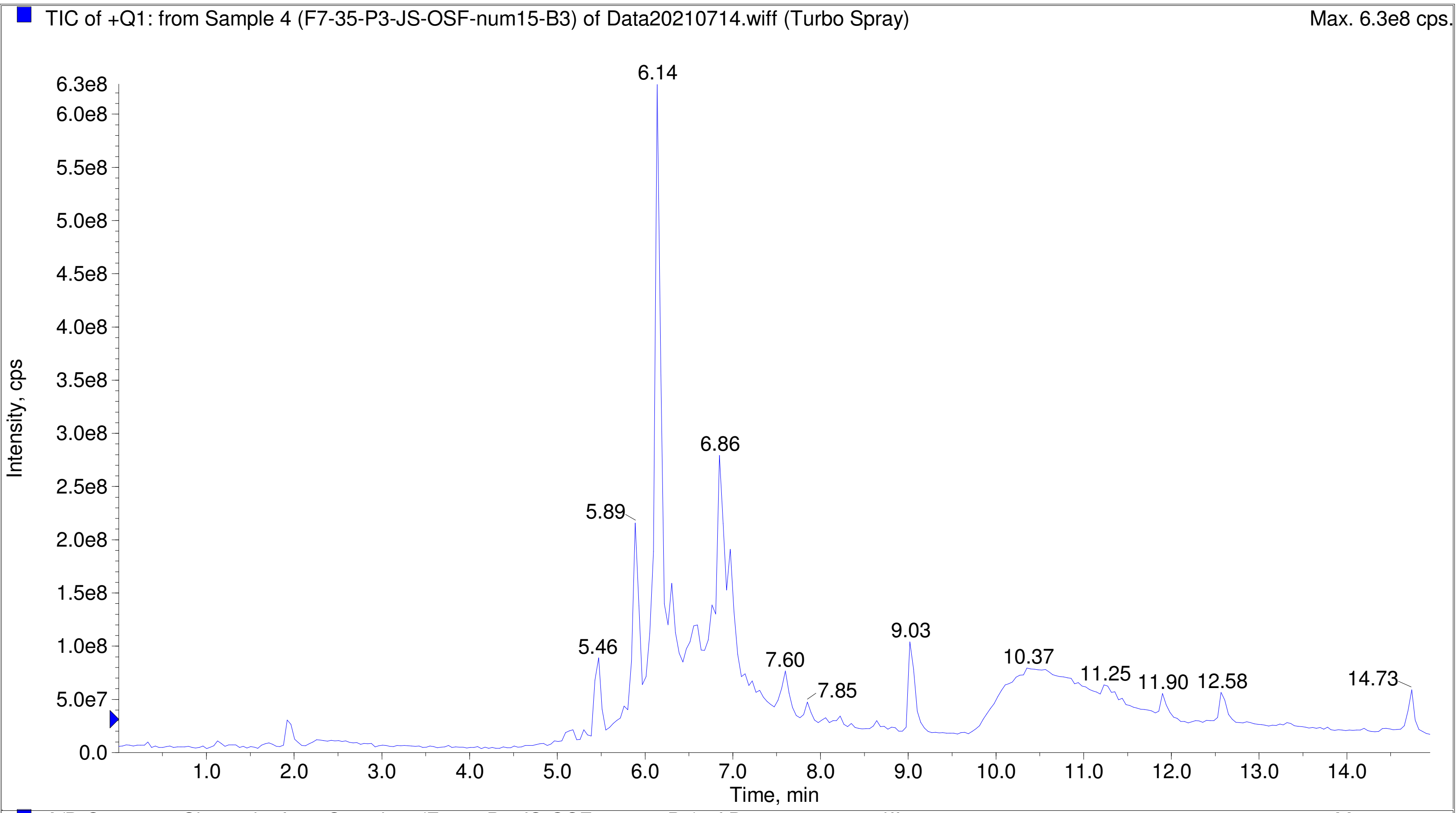
TIC of +Q1: from Sample 4 (F7-35-P3-JS-OSF-num15-B3) of Data20210714.wiff (Turbo Spray)
Max. 6.3e8 cps.
Intensity, cps
Time, min
6.14
6.86
5.89
5.46
7.60
7.85
9.03
10.37
11.25
11.90
12.58
14.73

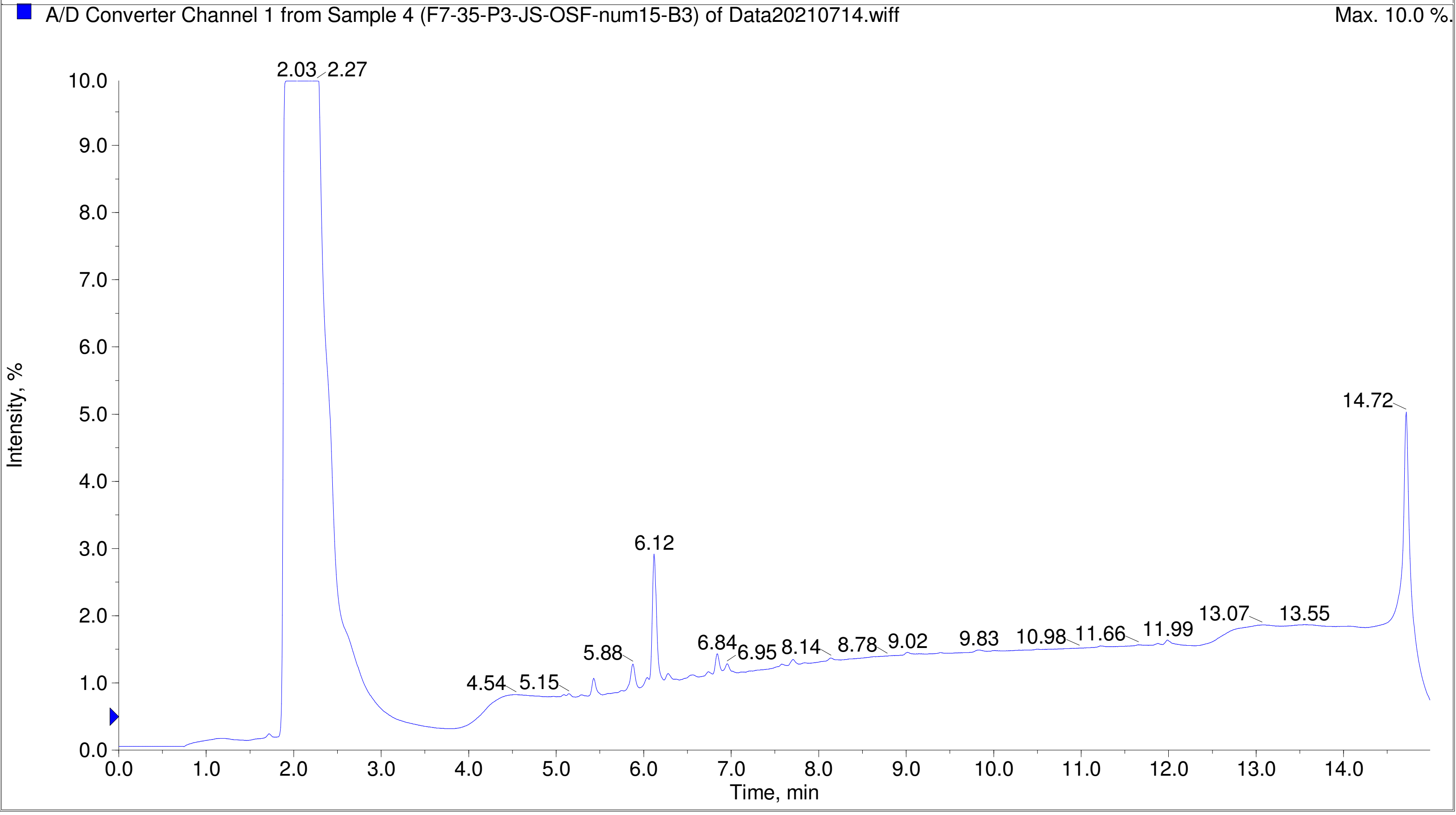
A/D Converter Channel 1 from Sample 4 (F7-35-P3-JS-OSF-num15-B3) of Data20210714.wiff
Max. 10.0 %.
Intensity, %
Time, min
2.03
2.27
14.72
6.12
5.88
4.54
5.15
6.84
6.95
8.14
8.78
9.02
9.83
10.98
11.66
11.99
13.07
13.55

Printing Time: 2:27:37 PM	Post PM	Polarity/Scan Type: Positive Q1 MS
Printing Date: Monday, January 30, 2023	Sample Name: F7-35-P3-JS-OSF-num19-B7
Service Engineer Chris Maier	*Stanford U	*API 3000 SN D7640012

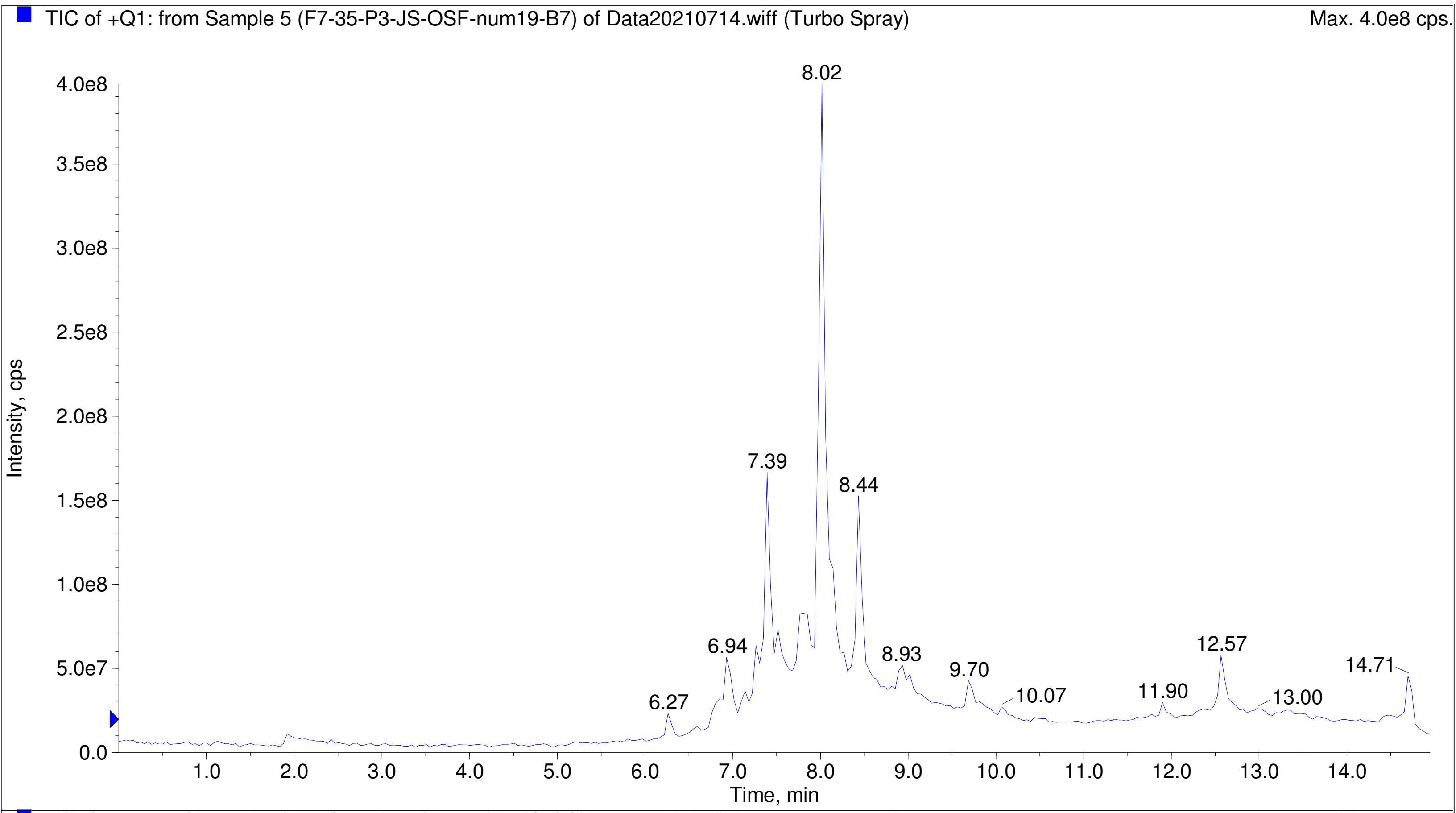


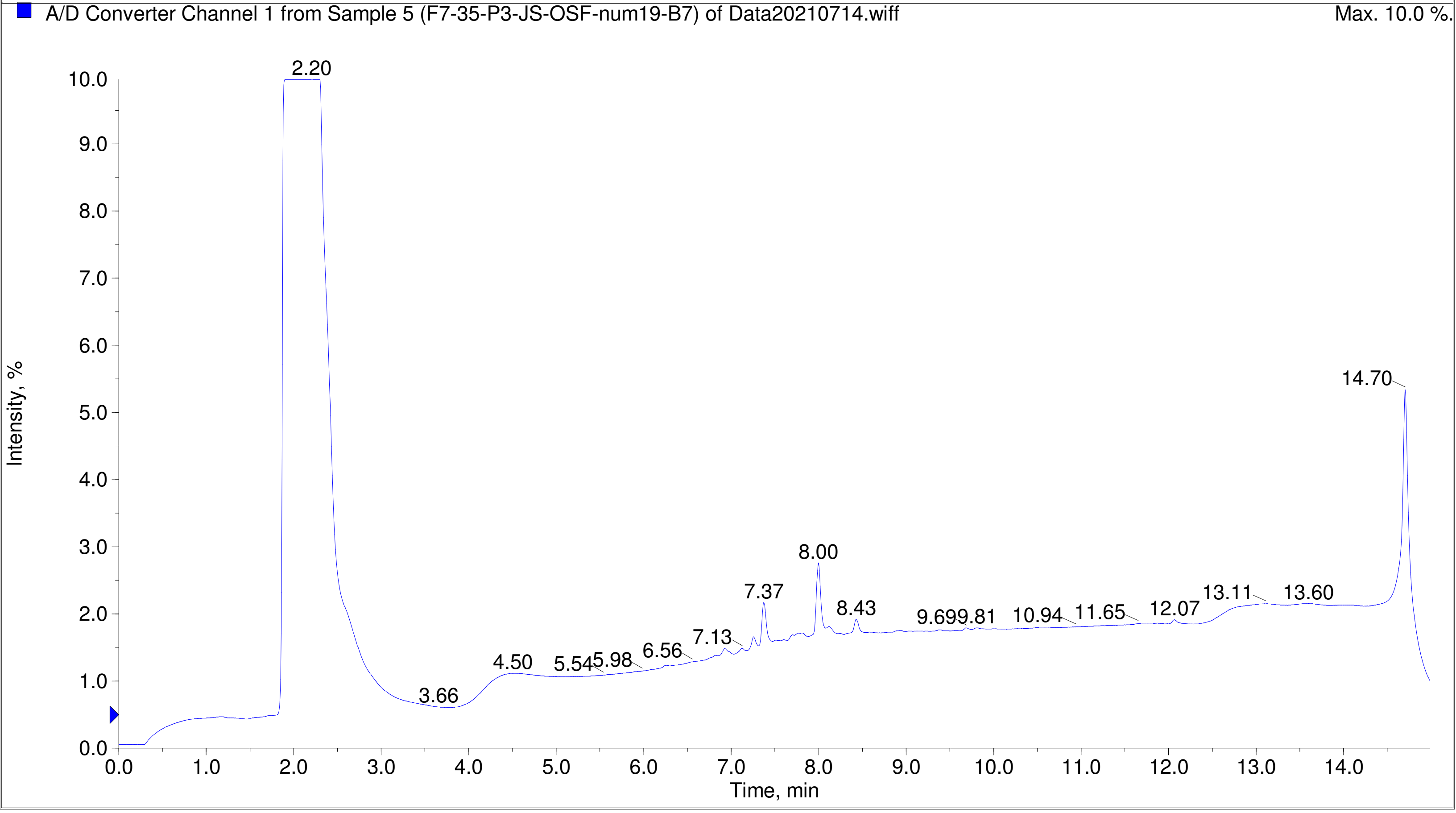

Printing Time: 2:28:30 PM
Printing Date: Monday, January 30, 2023
Service Engineer Chris Maier

Post PM
*Stanford U

Polarity/Scan Type: Positive Q1 MS
Sample Name: F7-35-P3-JS-OSF-num23-B11
*API 3000 SN D7640012

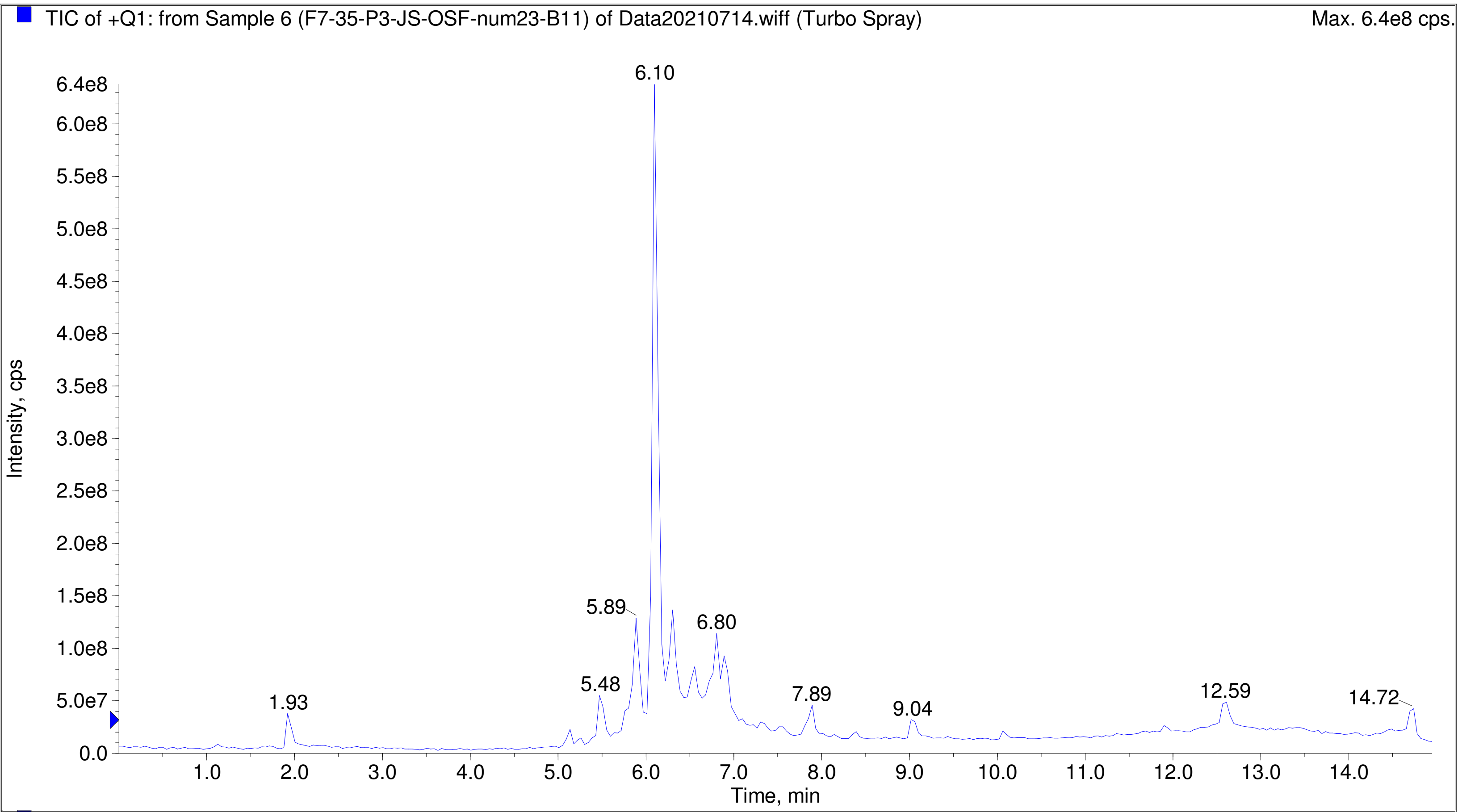


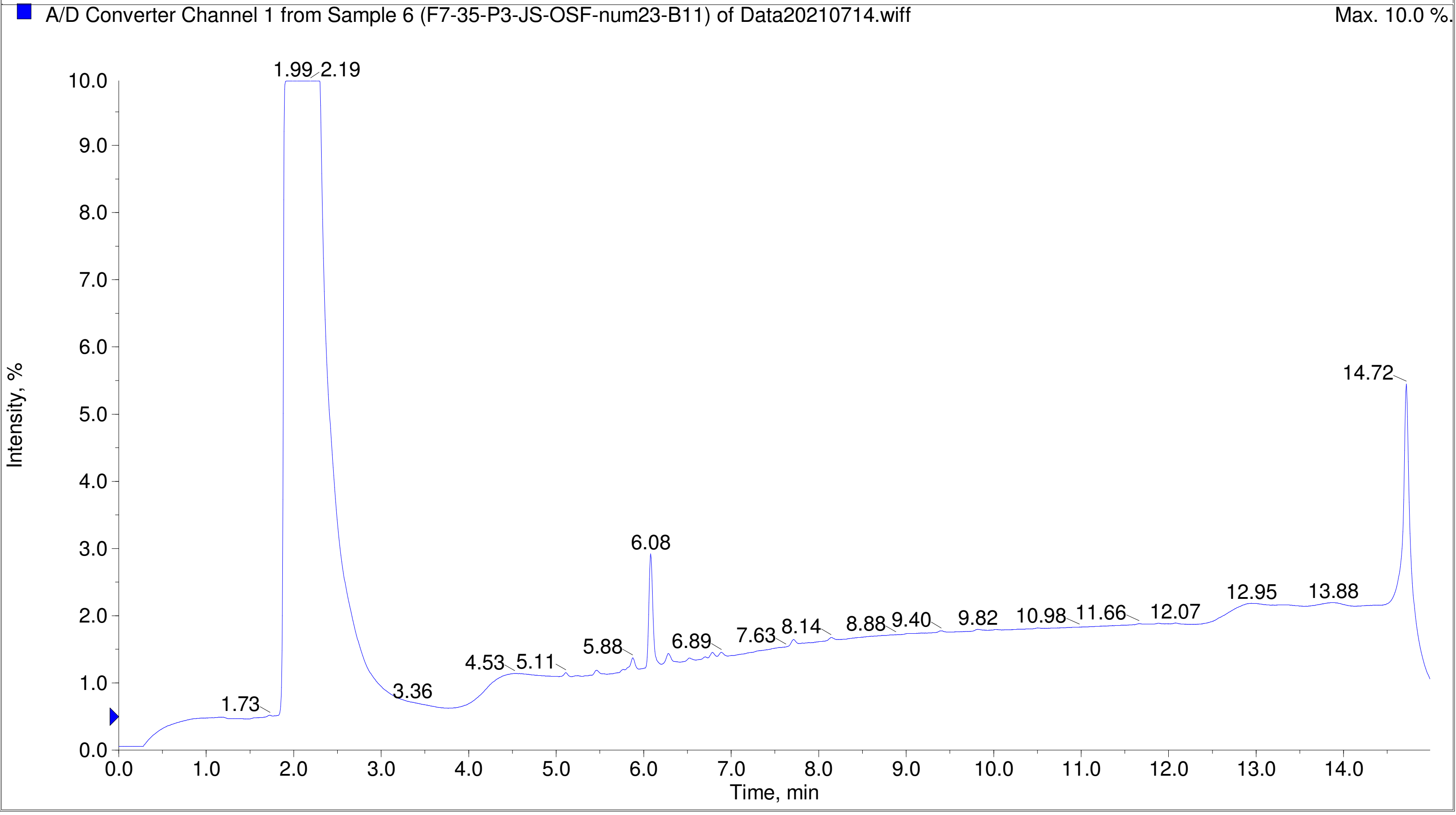

Printing Time: 3:52:31 PM　　Post PM　　Polarity/Scan Type: Positive Q1 MS
Printing Date: Tuesday, January 31, 2023　　Sample Name: F7-35-P3-JS-OSF-num32-C8
Service Engineer Chris Maier　　*Stanford U　　*API 3000 SN D7640012

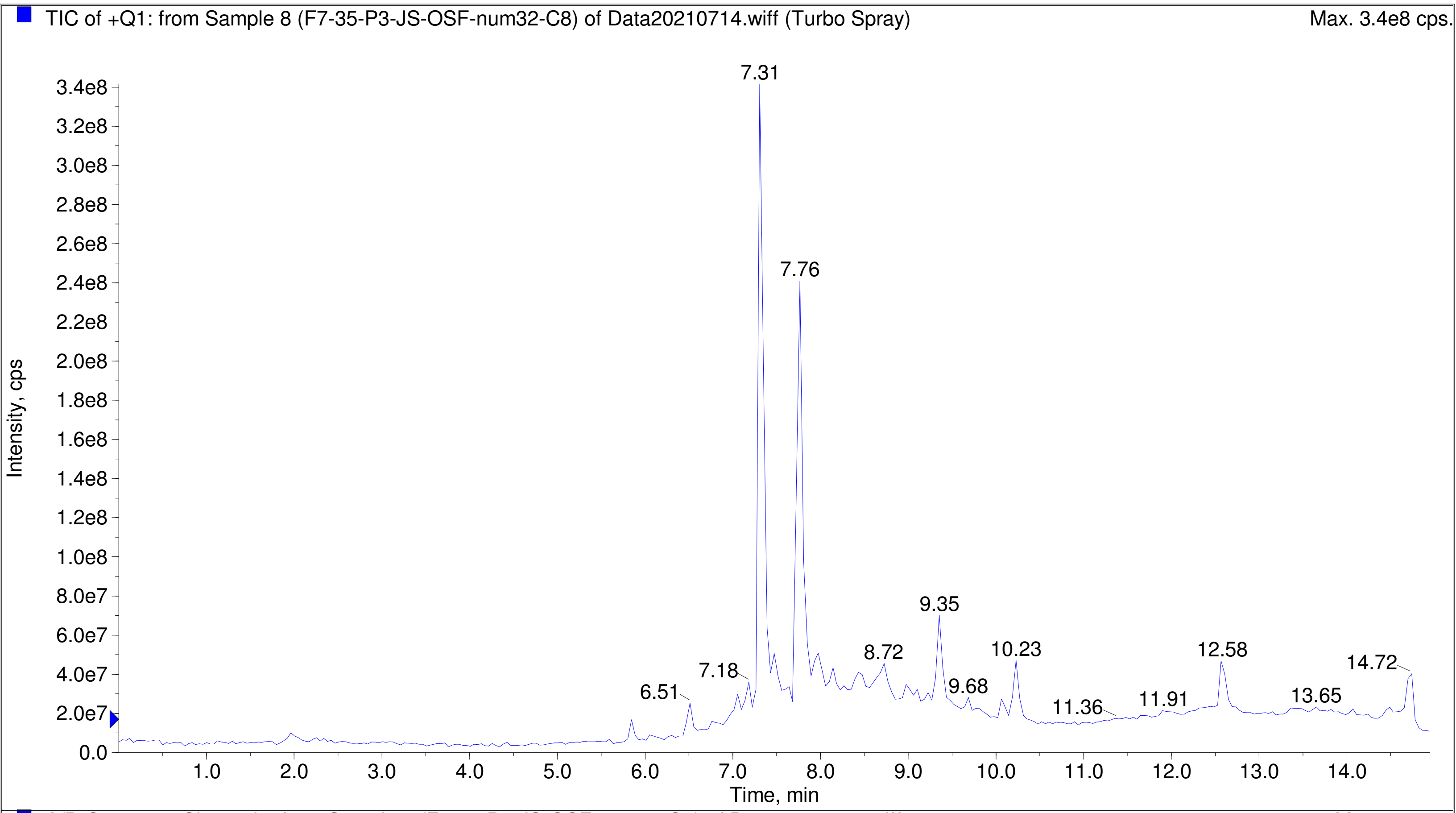


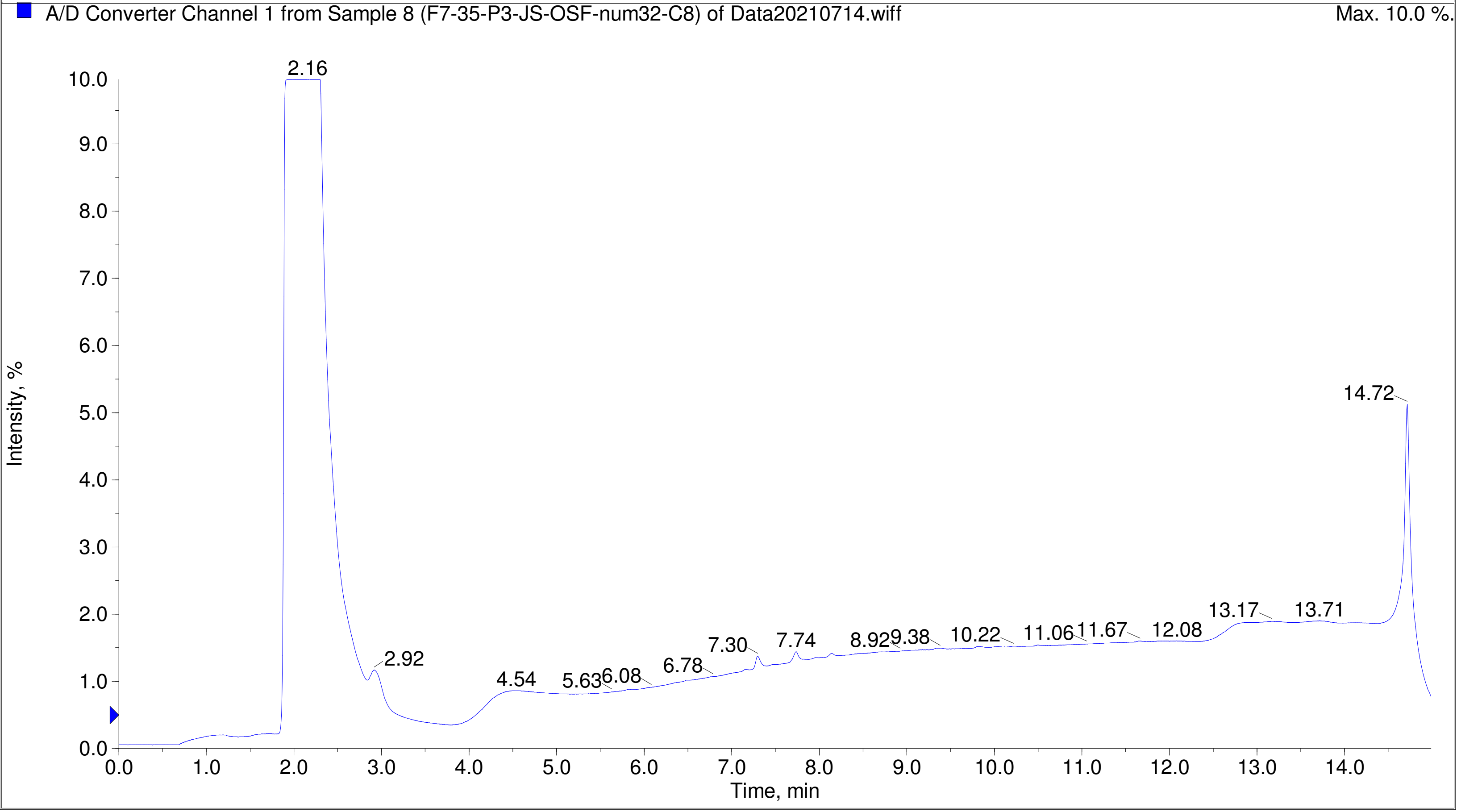

Printing Time: 3:53:59 PM
Printing Date: Tuesday, January 31, 2023
Service Engineer Chris Maier

Post PM
*Stanford U

Polarity/Scan Type: Positive Q1 MS
Sample Name: F7-35-P3-JS-OSF-num42-D6
*API 3000 SN D7640012

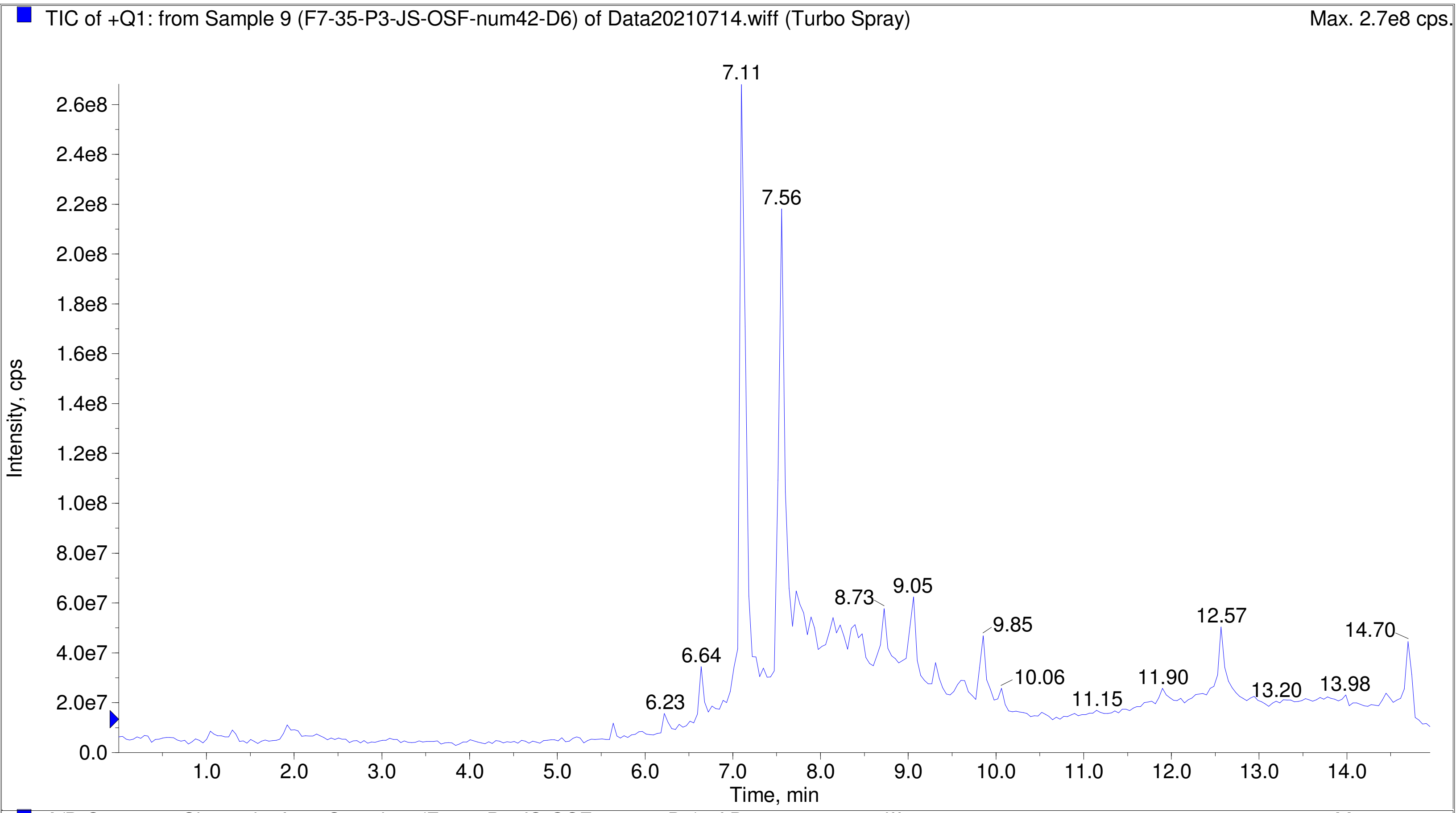


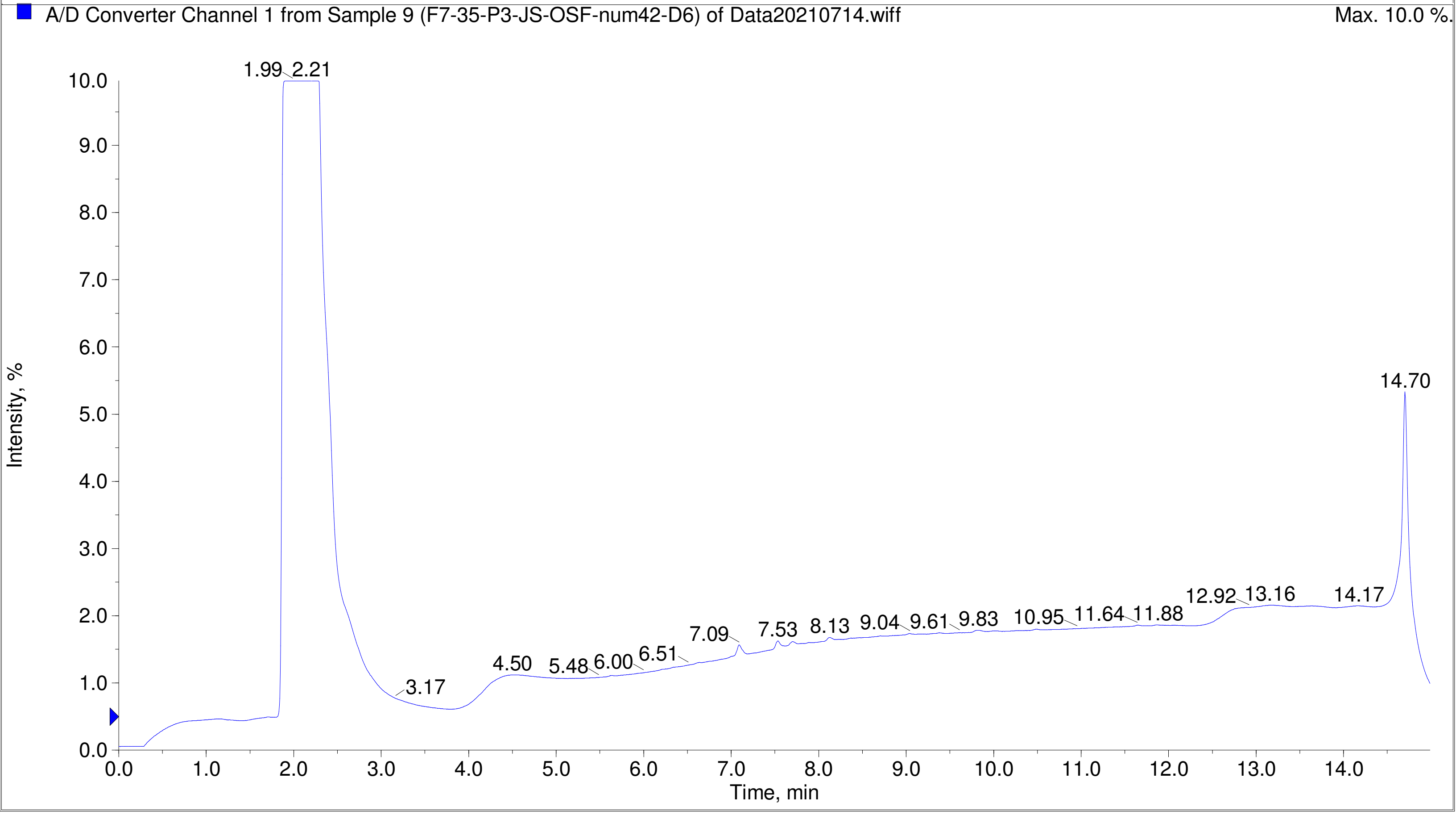

Printing Time: 3:55:21 PM　Post PM　Polarity/Scan Type: Positive Q1 MS
Printing Date: Tuesday, January 31, 2023　Sample Name: F7-35-P3-JS-OSF-num46-D10
Service Engineer Chris Maier　*Stanford U　*API 3000 SN D7640012

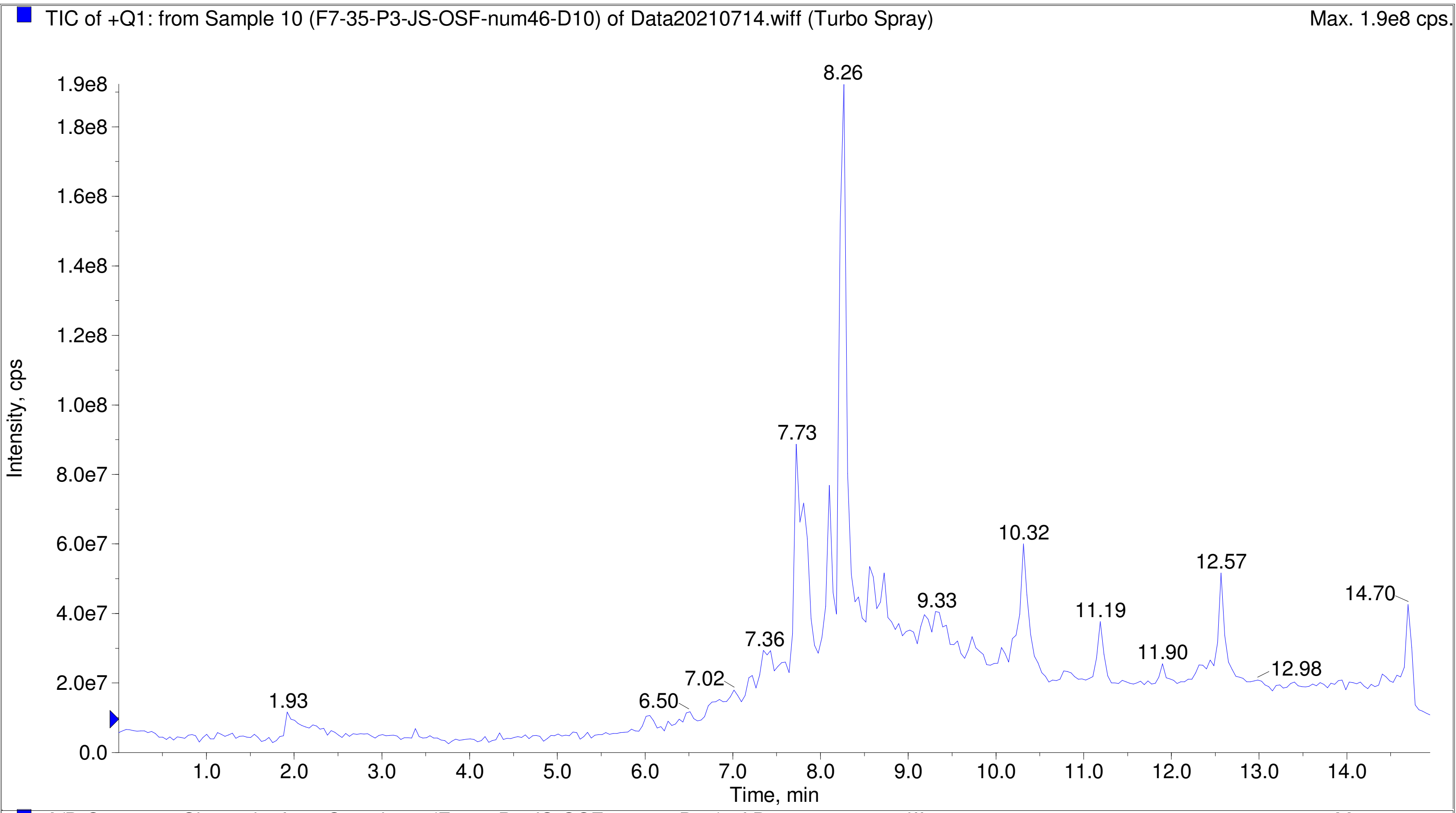


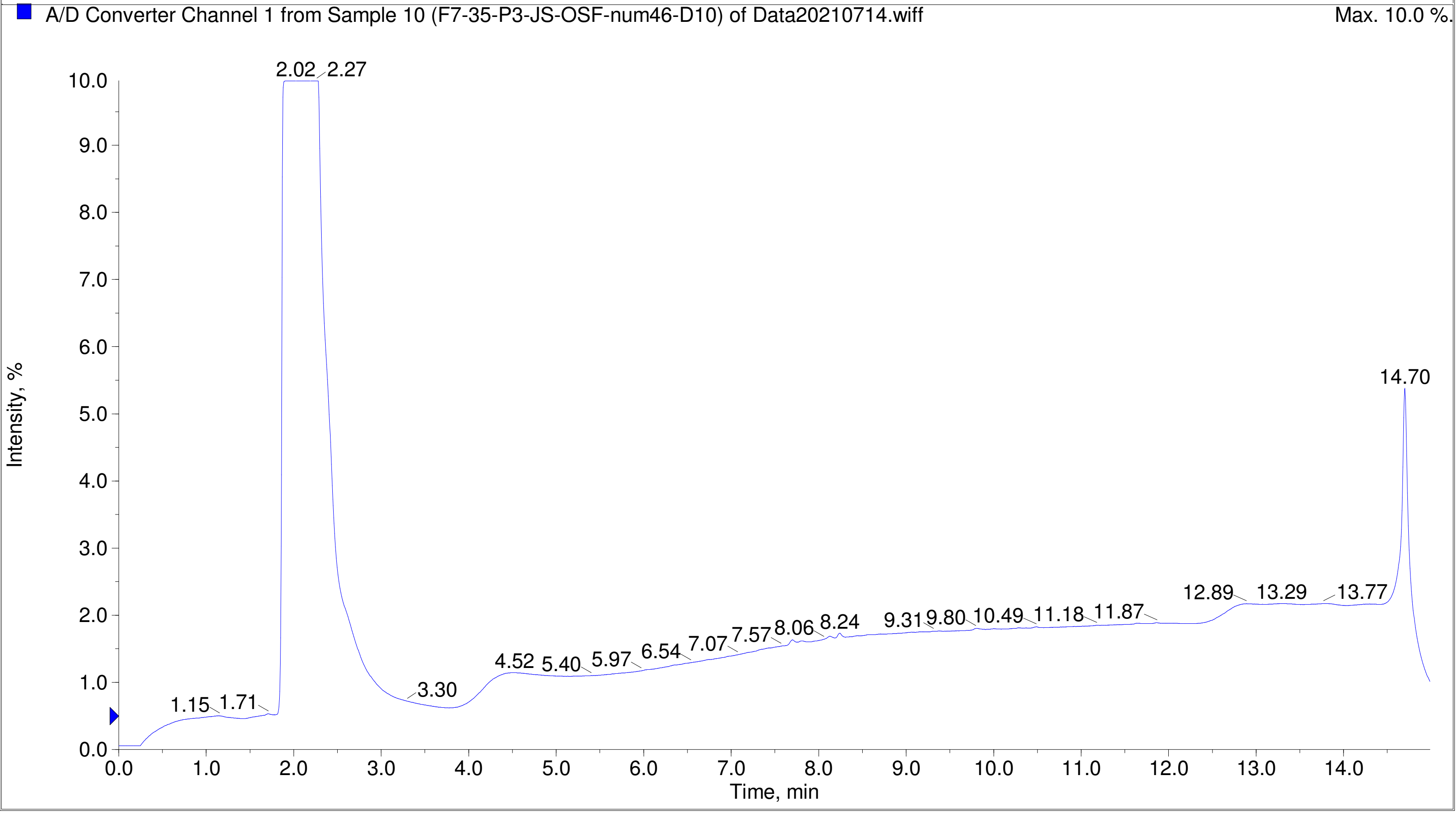

Printing Time: 3:56:34 PM  Post PM  Polarity/Scan Type: Positive Q1 MS
Printing Date: Tuesday, January 31, 2023  Sample Name: F7-35-P3-JS-OSF-num48-D12
Service Engineer Chris Maier  *Stanford U  *API 3000 SN D7640012

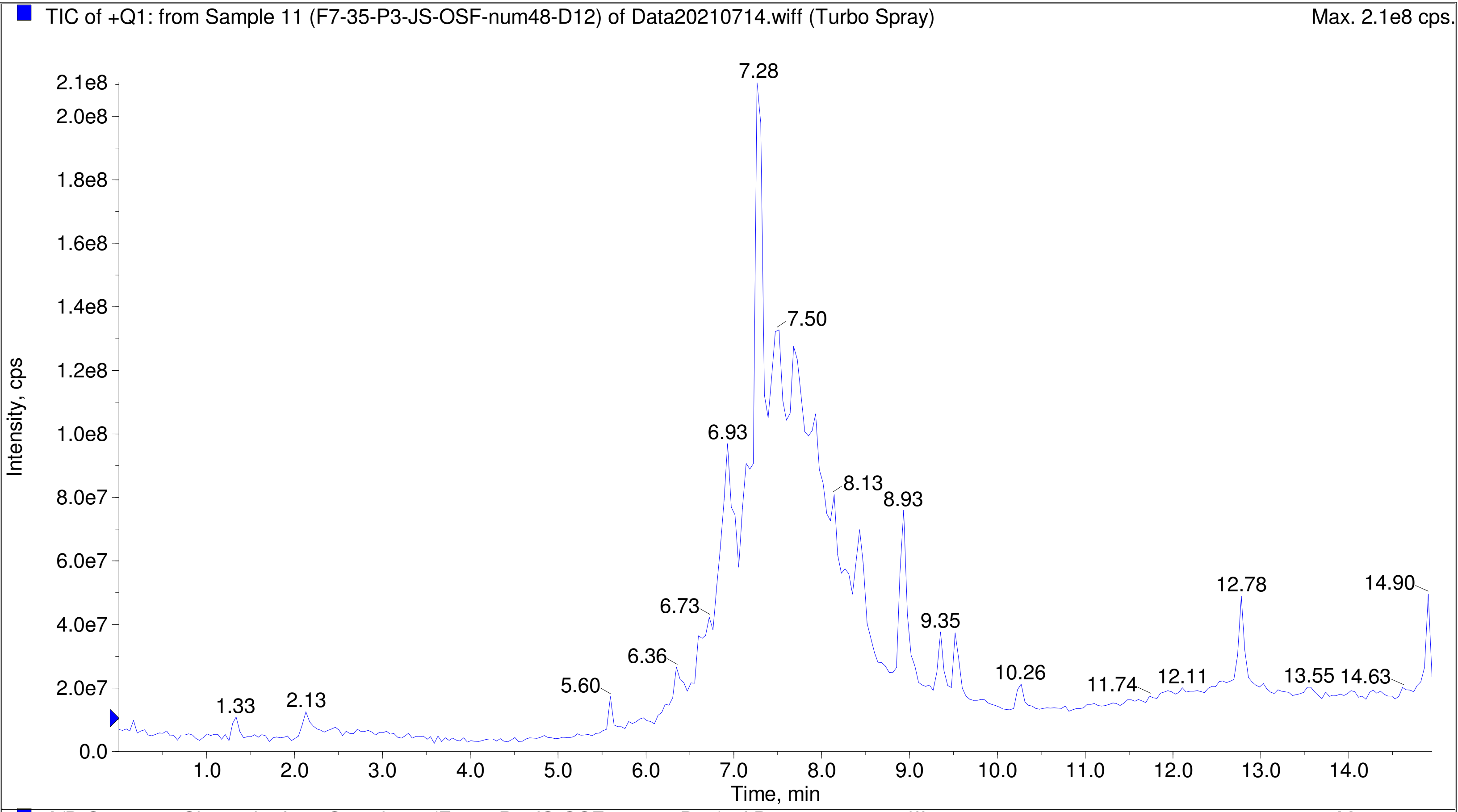


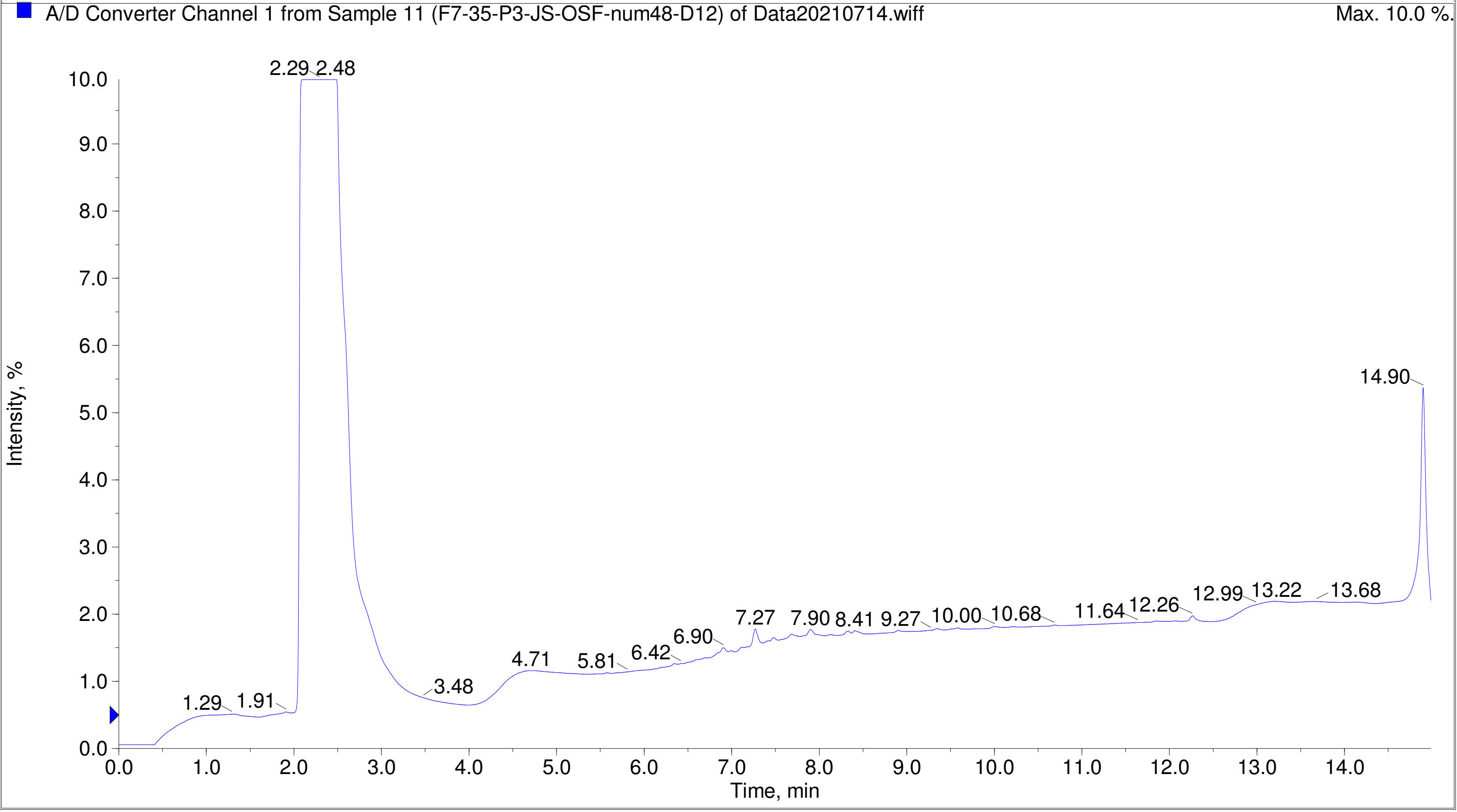

Printing Time: 3:57:28 PM    Post PM    Polarity/Scan Type: Positive Q1 MS
Printing Date: Tuesday, January 31, 2023    Sample Name: F7-35-P3-JS-OSF-num52-E4
Service Engineer Chris Maier    *Stanford U    *API 3000 SN D7640012

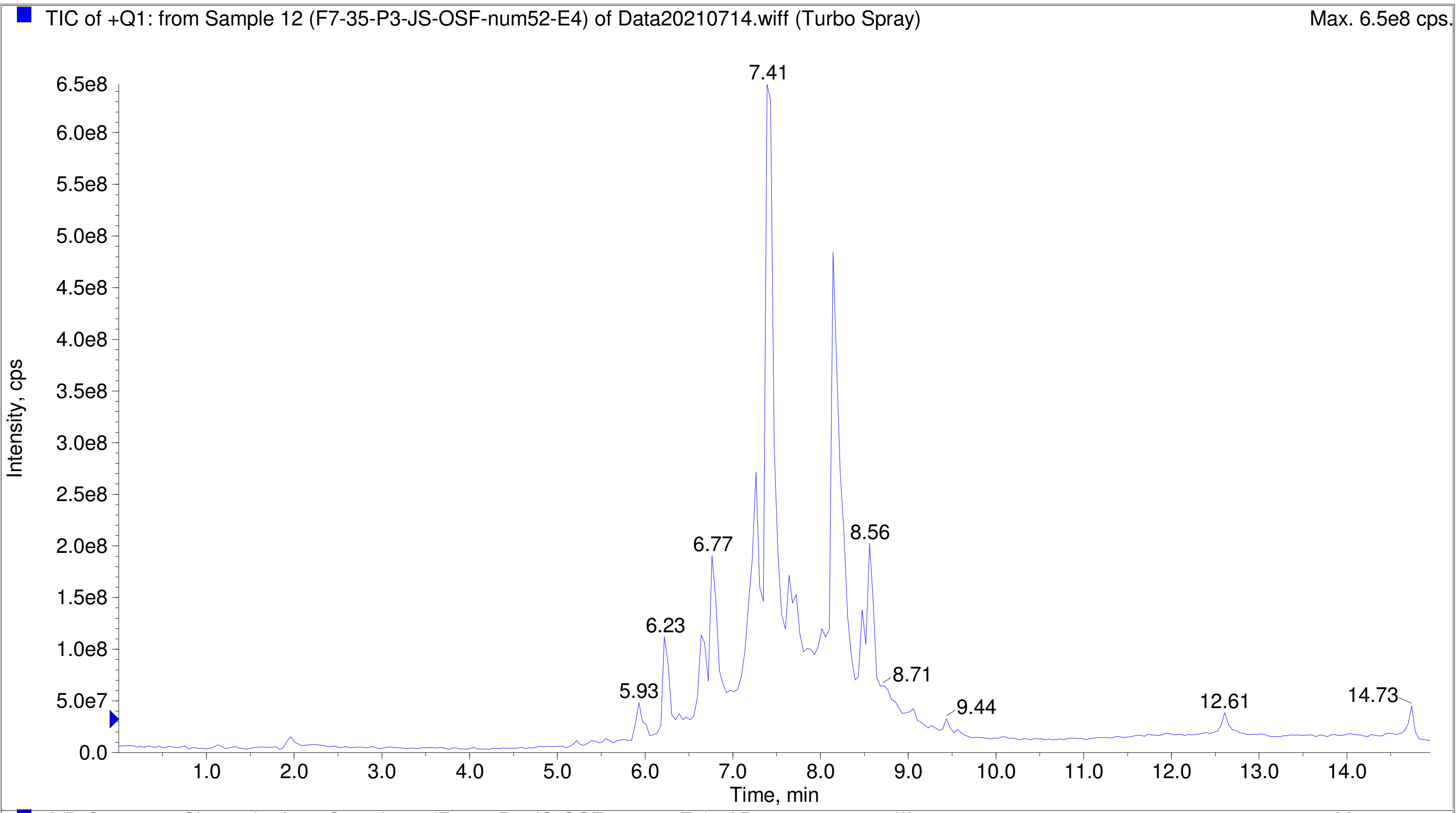


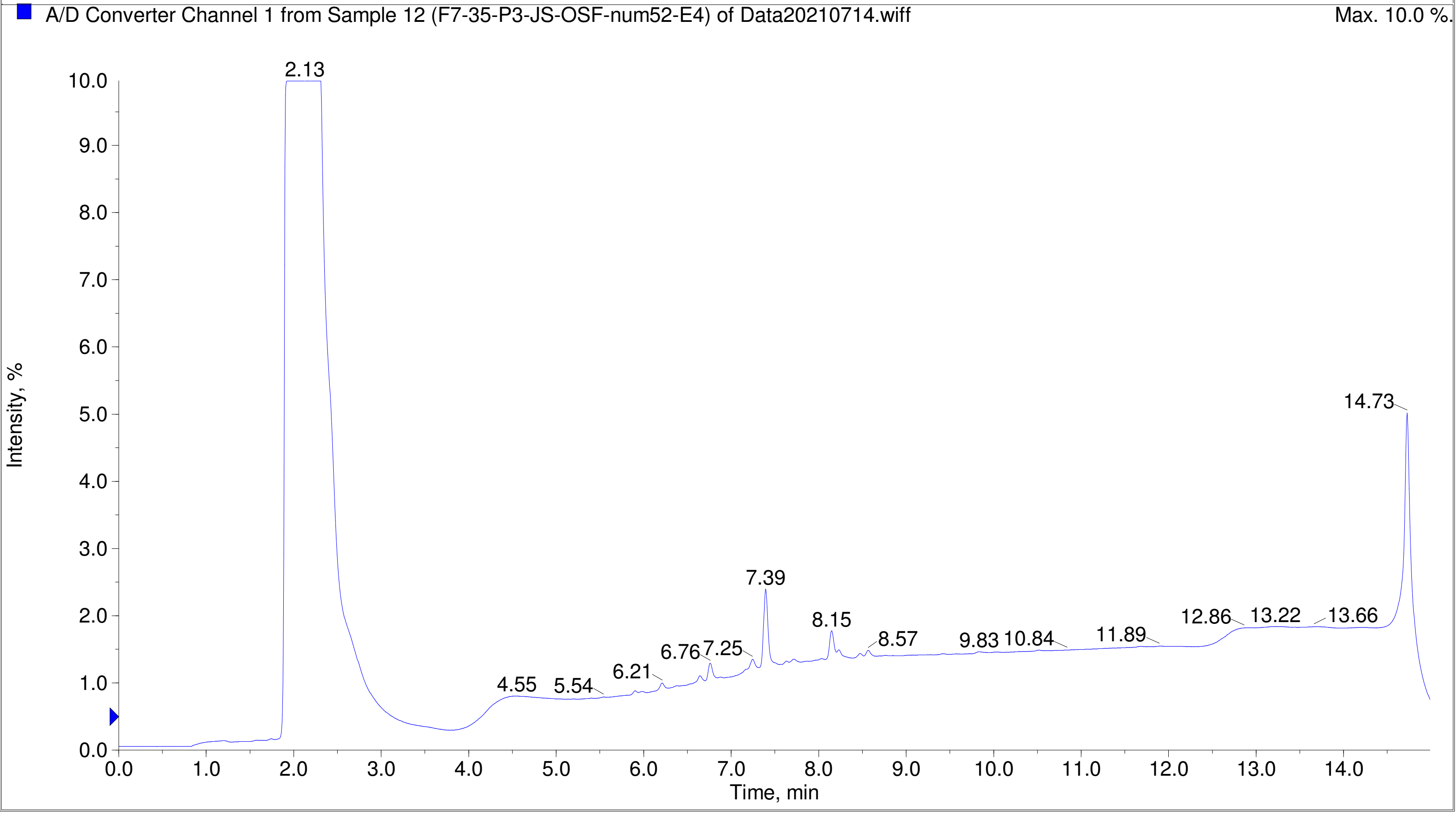

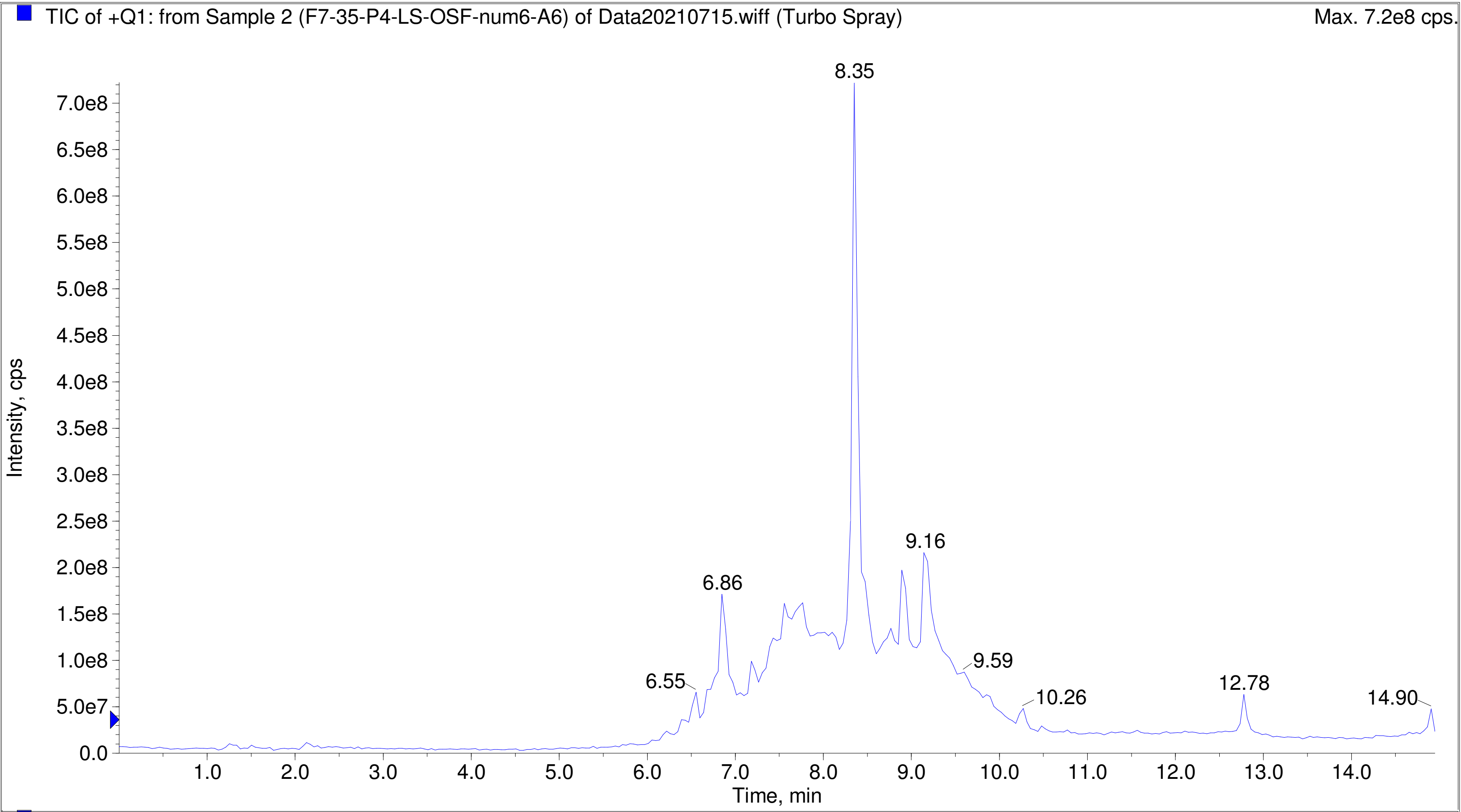
TIC of +Q1: from Sample 2 (F7-35-P4-LS-OSF-num6-A6) of Data20210715.wiff (Turbo Spray)
Max. 7.2e8 cps.
Intensity, cps
Time, min
8.35
9.16
6.86
6.55
9.59
10.26
12.78
14.90

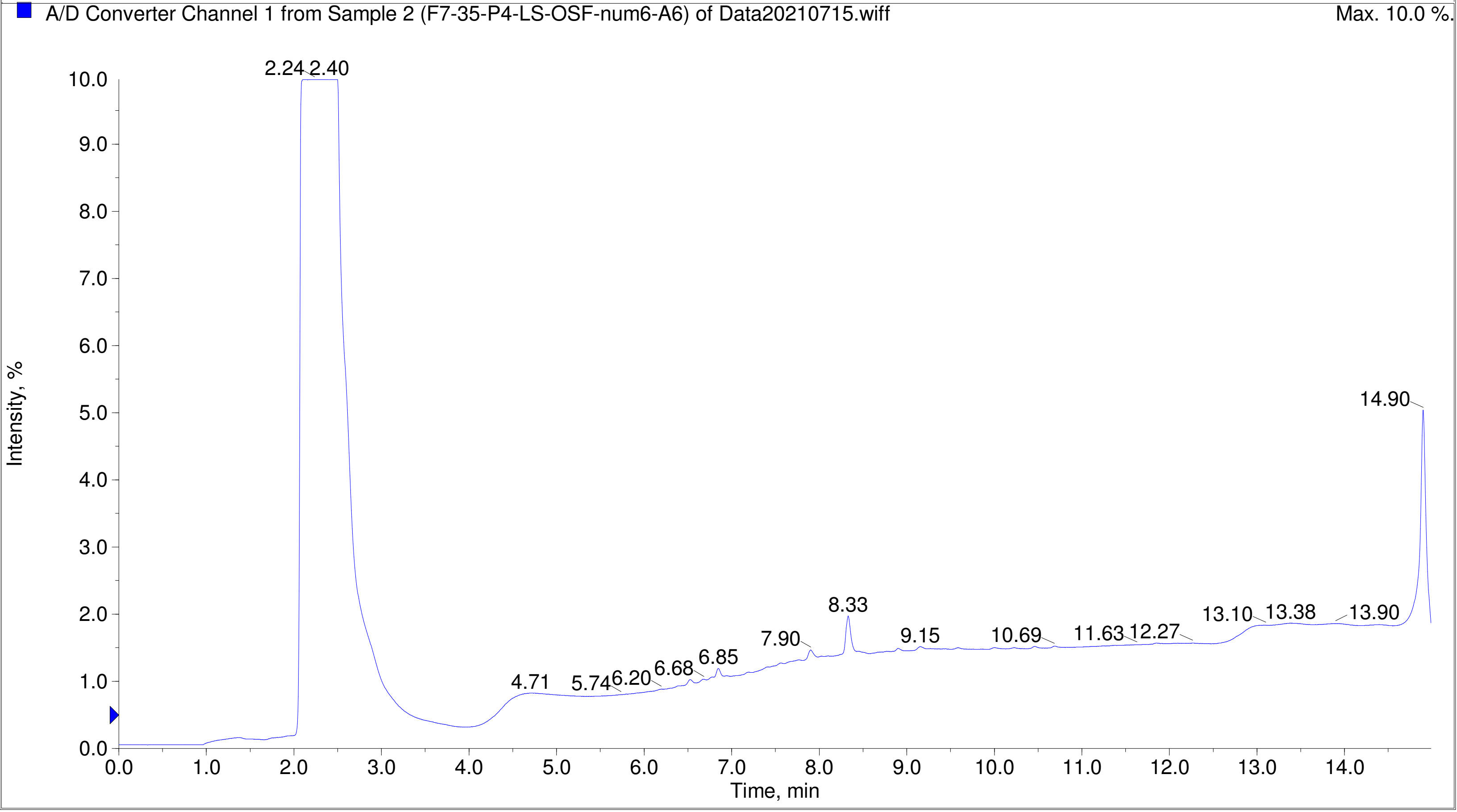
A/D Converter Channel 1 from Sample 2 (F7-35-P4-LS-OSF-num6-A6) of Data20210715.wiff
Max. 10.0 %.
Intensity, %
Time, min
2.24
2.40
14.90
8.33
13.10
13.38
13.90
7.90
9.15
10.69
11.63
12.27
6.85
6.68
6.20
5.74
4.71

Printing Time: 2:31:25 PM Post PM Polarity/Scan Type: Positive Q1 MS
Printing Date: Monday, January 30, 2023 Sample Name: F7-35-P4-LS-OSF-num10-A10
Service Engineer Chris Maier *Stanford U *API 3000 SN D7640012

TIC of +Q1: from Sample 3 (F7-35-P4-LS-OSF-num10-A10) of Data20210715.wiff (Turbo Spray) Max. 5.5e8 cps.

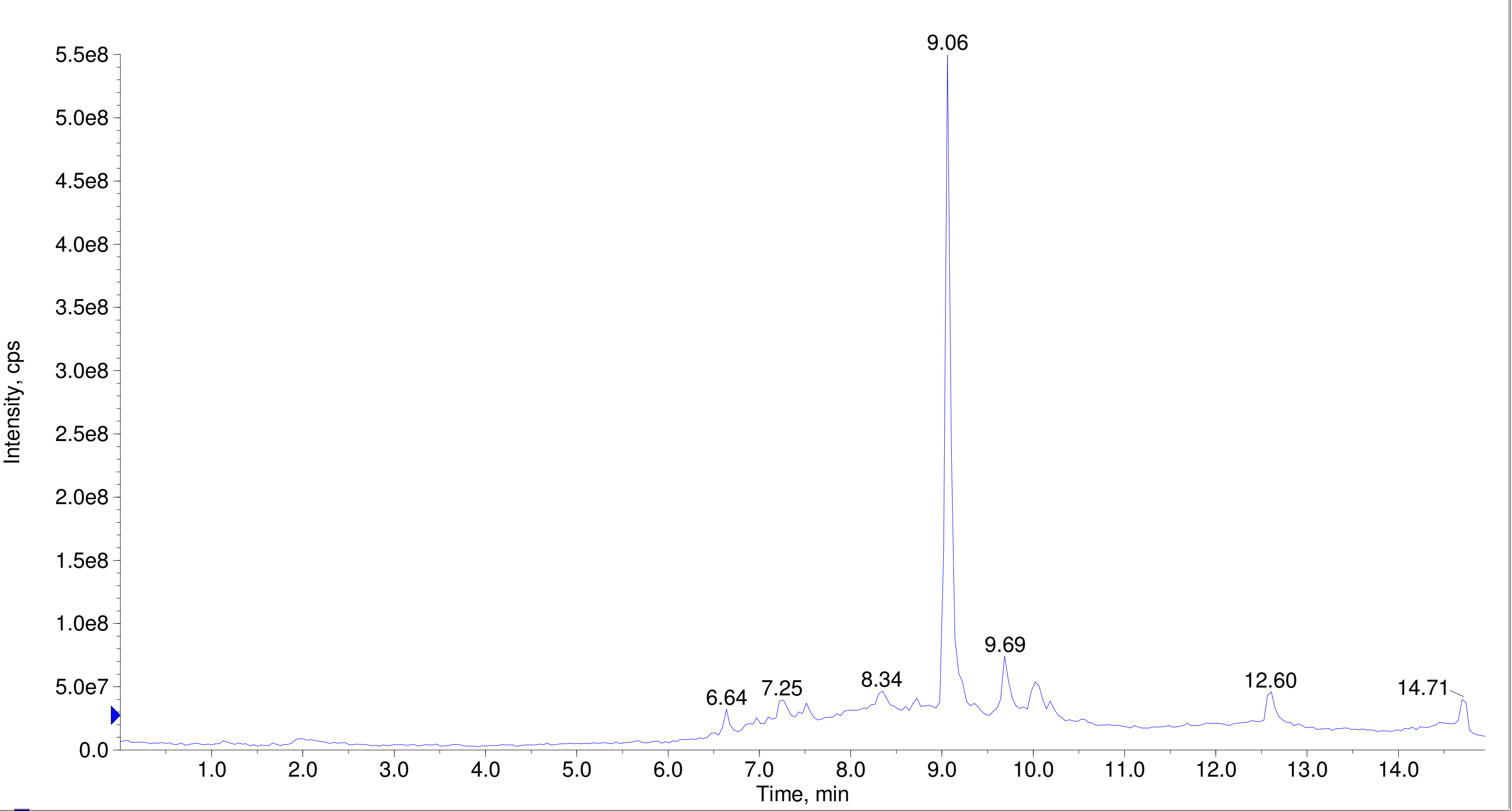


A/D Converter Channel 1 from Sample 3 (F7-35-P4-LS-OSF-num10-A10) of Data20210715.wiff Max. 10.0 %.

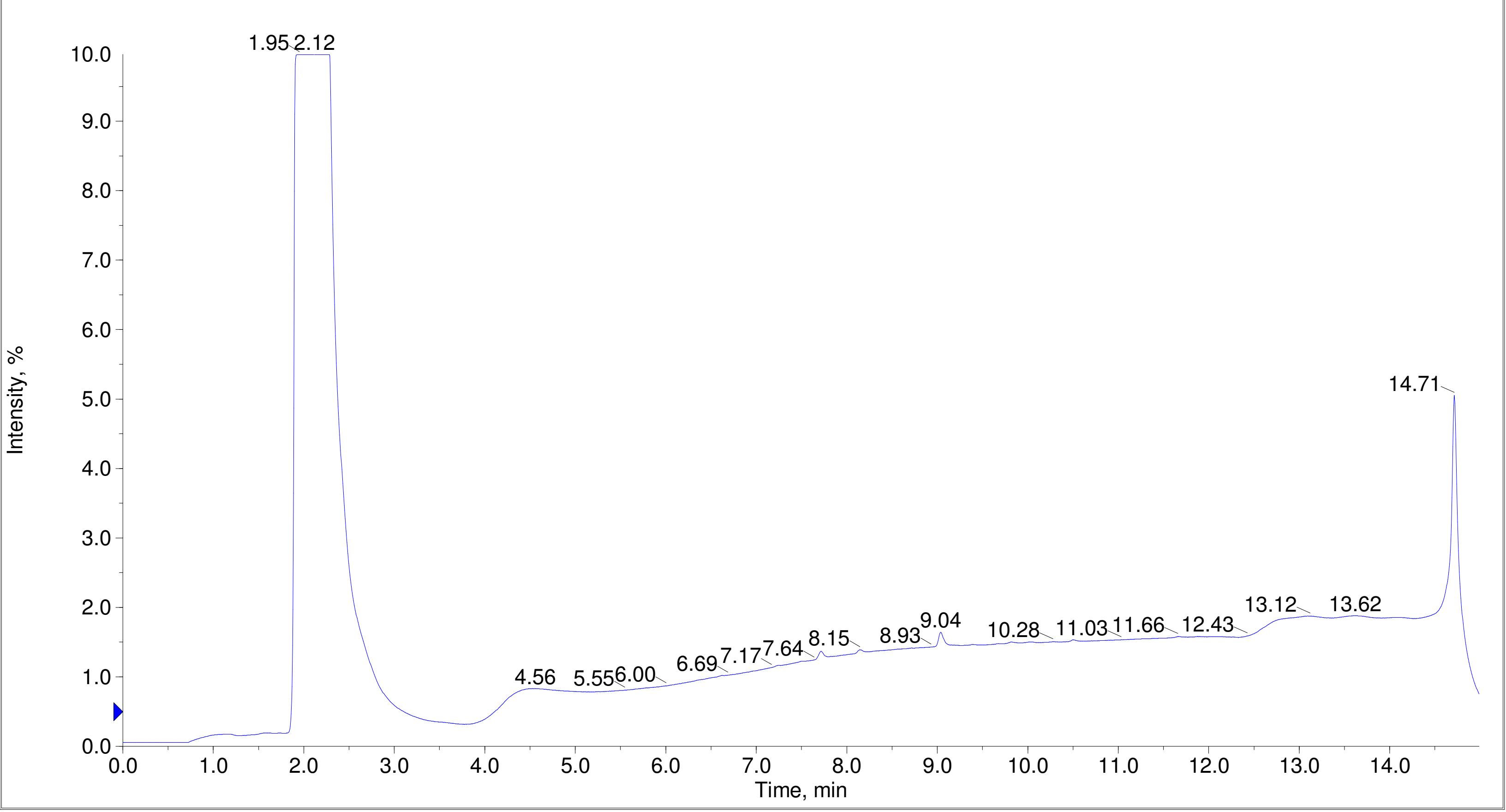

Printing Time: 2:38:07 PM  Post PM  Polarity/Scan Type: Positive Q1 MS
Printing Date: Monday, January 30, 2023  Sample Name: F7-35-P4-LS-OSF-num13-B1
Service Engineer Chris Maier  *Stanford U  *API 3000 SN D7640012

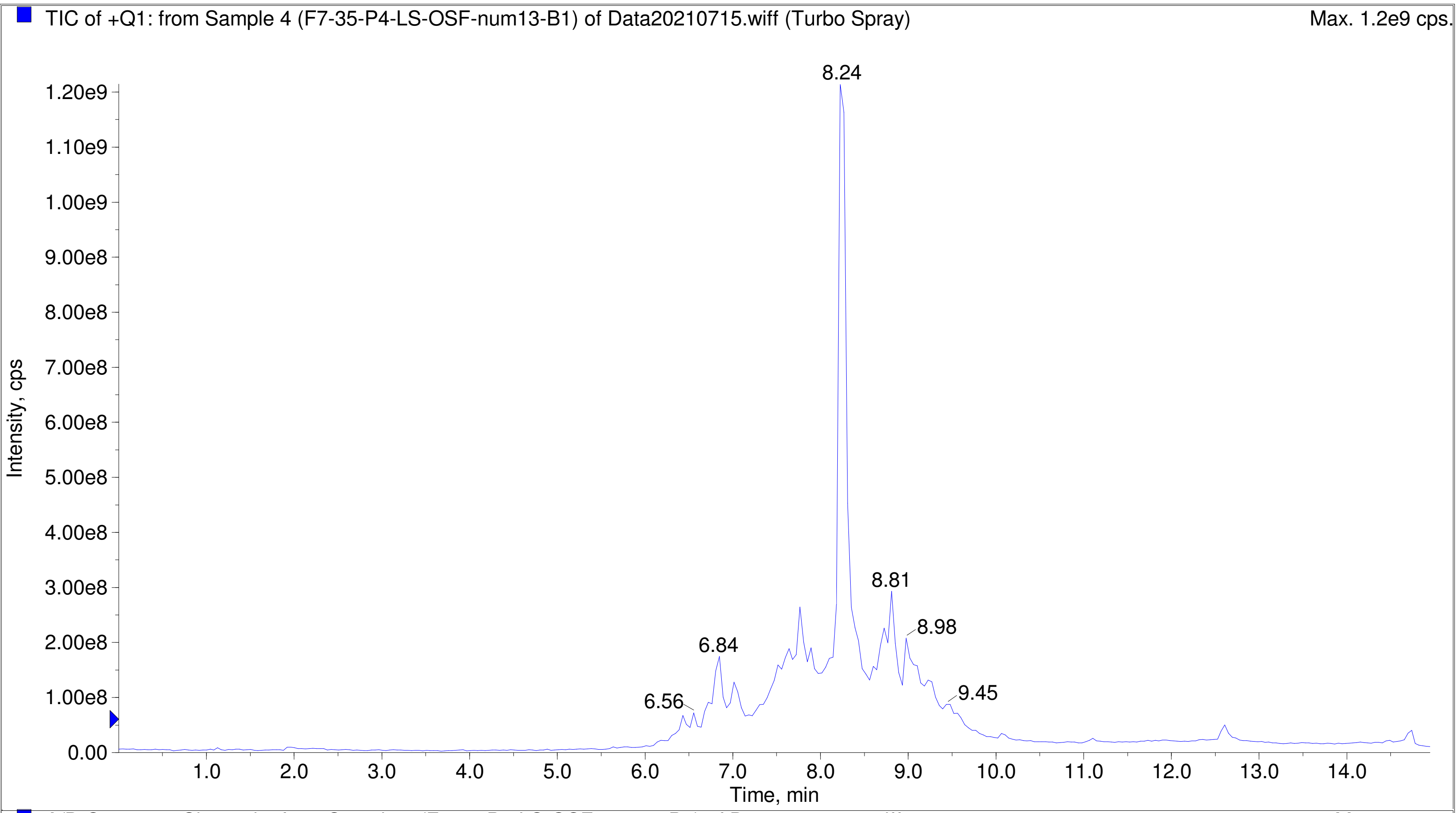


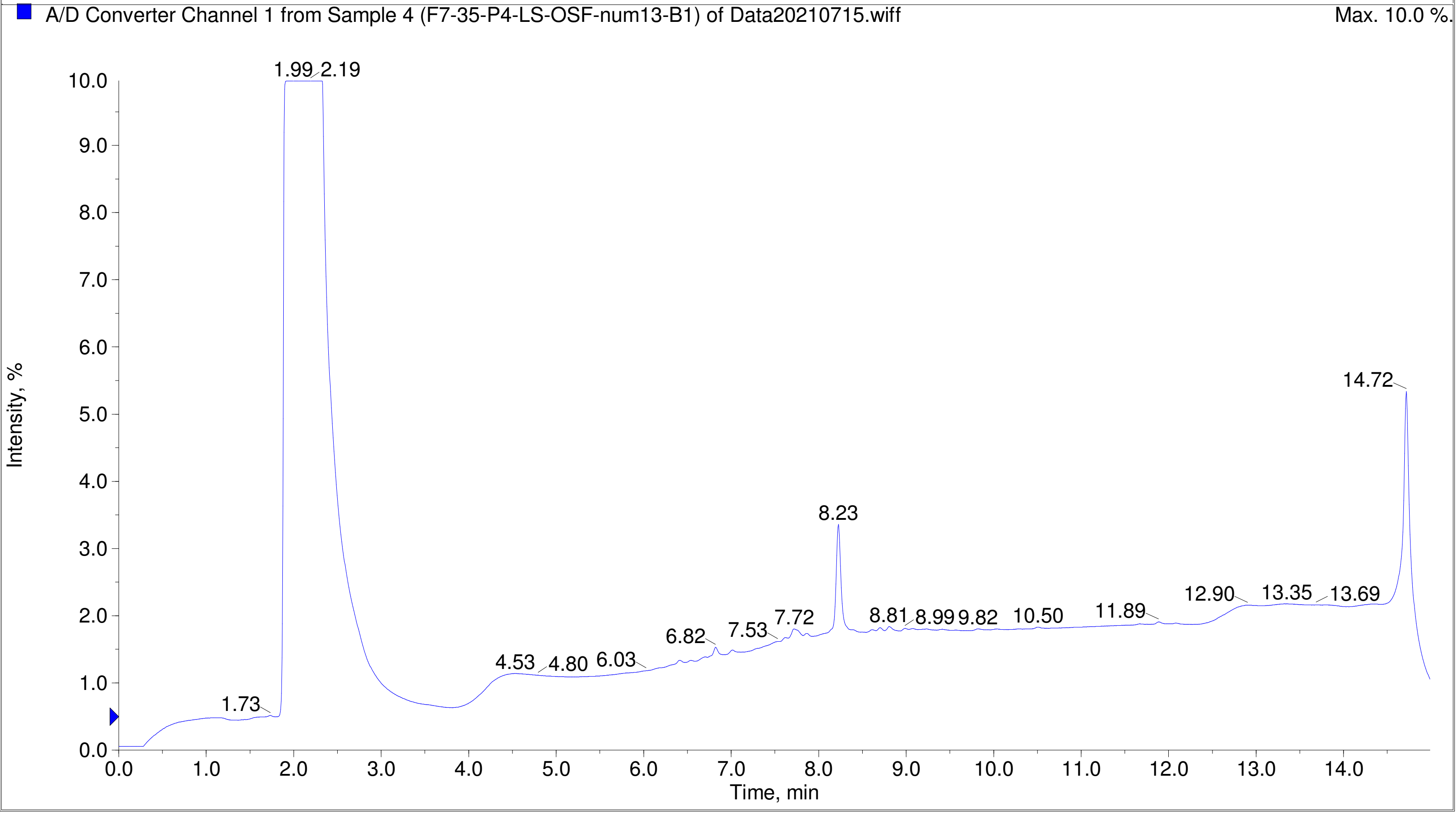

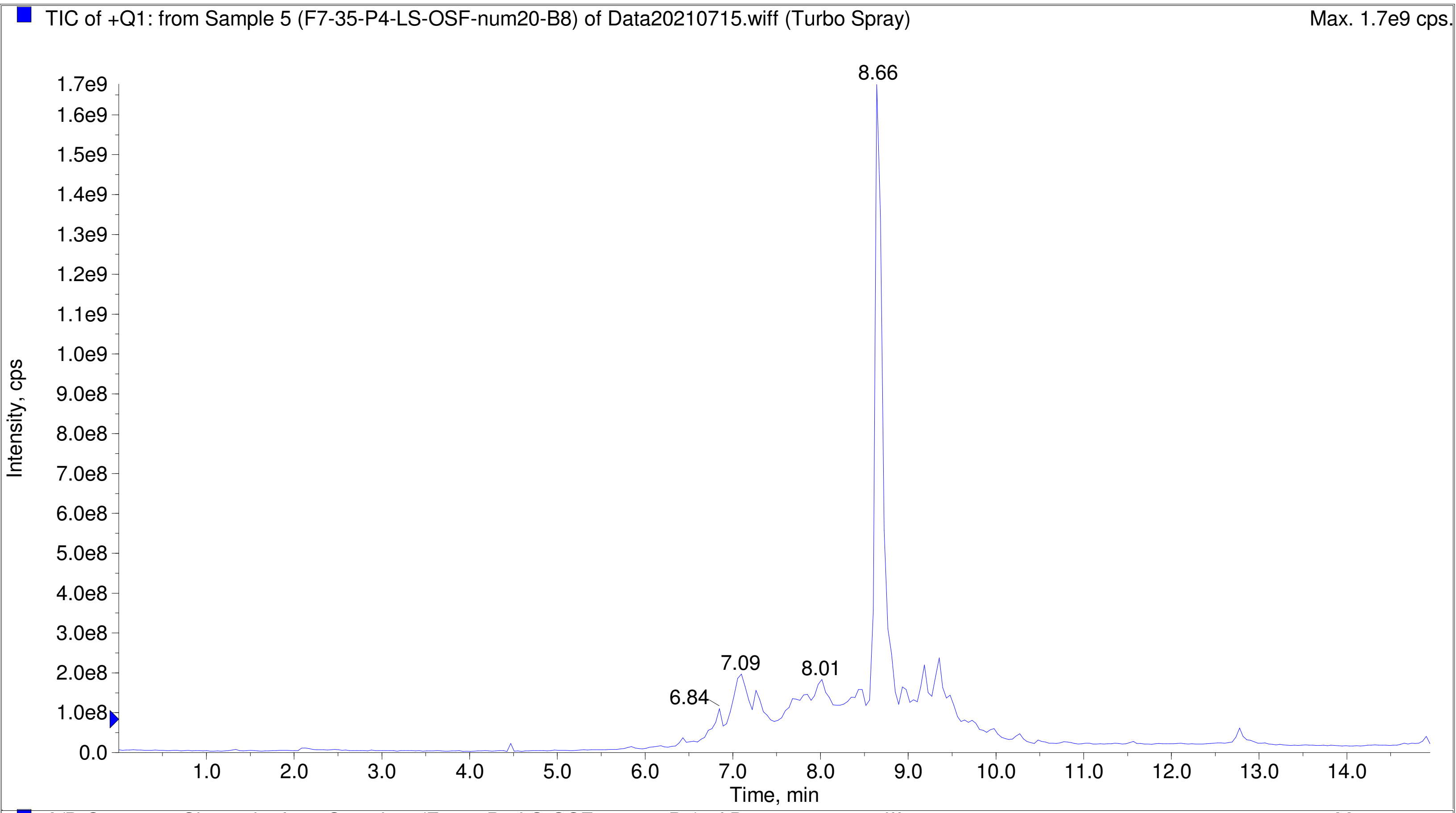
TIC of +Q1: from Sample 5 (F7-35-P4-LS-OSF-num20-B8) of Data20210715.wiff (Turbo Spray)
Max. 1.7e9 cps.
Intensity, cps
Time, min
8.66
7.09
8.01
6.84

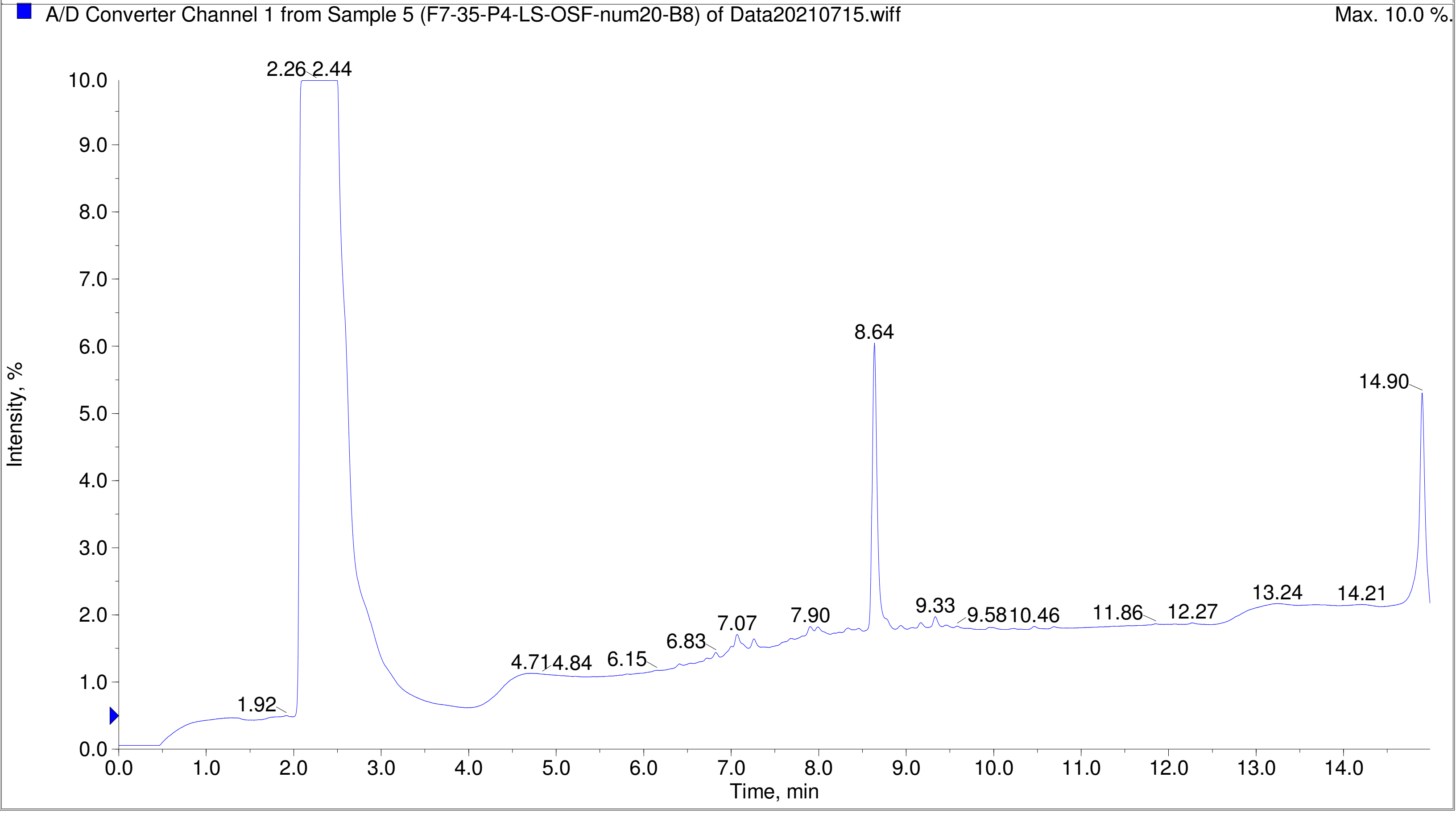
A/D Converter Channel 1 from Sample 5 (F7-35-P4-LS-OSF-num20-B8) of Data20210715.wiff
Max. 10.0 %.
Intensity, %
Time, min
2.26 2.44
8.64
14.90
1.92
4.71 4.84
6.15
6.83
7.07
7.90
9.33
9.58 10.46
11.86
12.27
13.24
14.21

Printing Time: 2:42:28 PM Post PM Polarity/Scan Type: Positive Q1 MS
Printing Date: Monday, January 30, 2023 Sample Name: F7-35-P4-LS-OSF-num23-B11
Service Engineer Chris Maier *Stanford U *API 3000 SN D7640012

TIC of +Q1: from Sample 6 (F7-35-P4-LS-OSF-num23-B11) of Data20210715.wiff (Turbo Spray) Max. 1.3e9 cps.

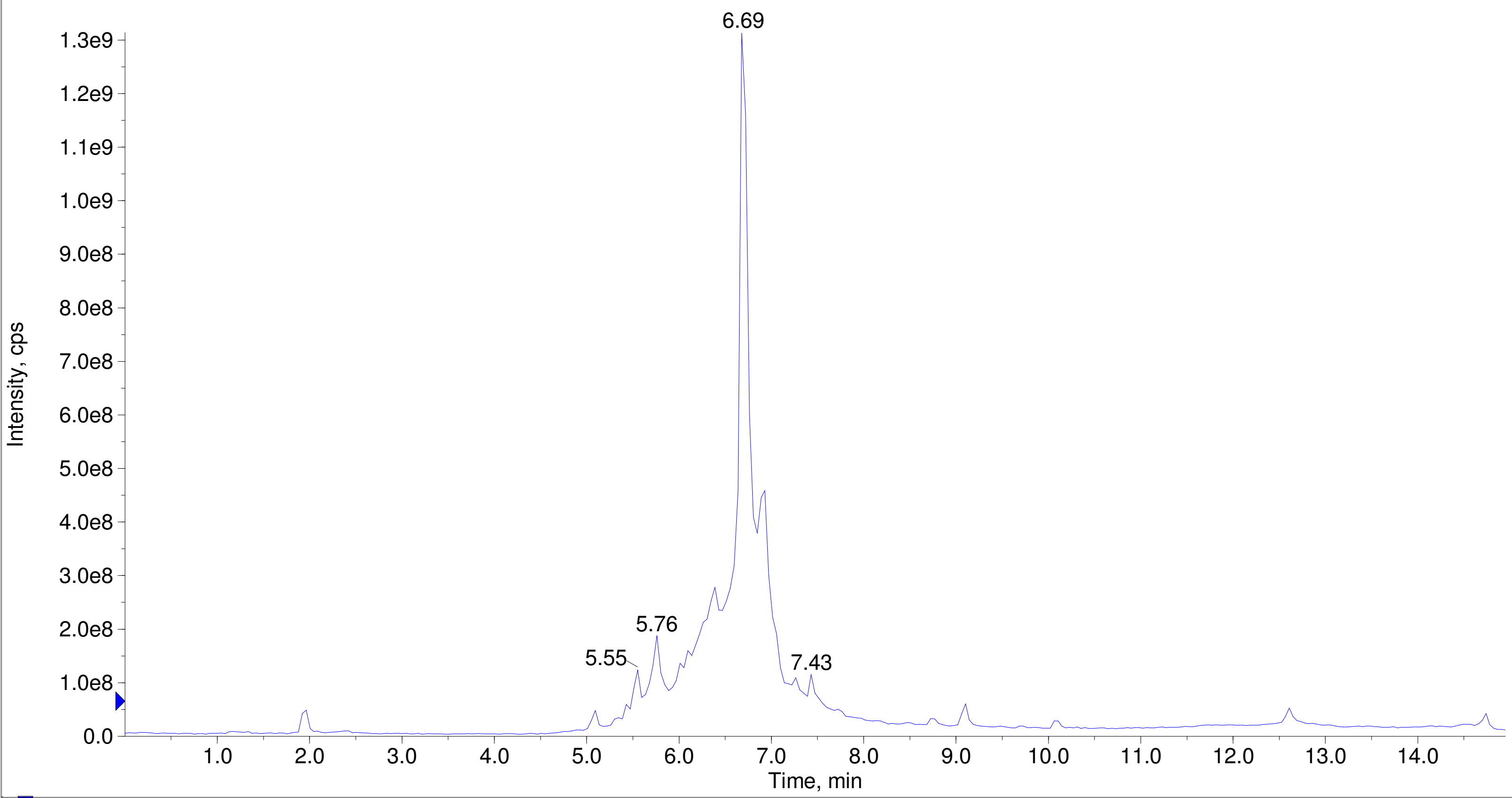


A/D Converter Channel 1 from Sample 6 (F7-35-P4-LS-OSF-num23-B11) of Data20210715.wiff Max. 10.0 %.

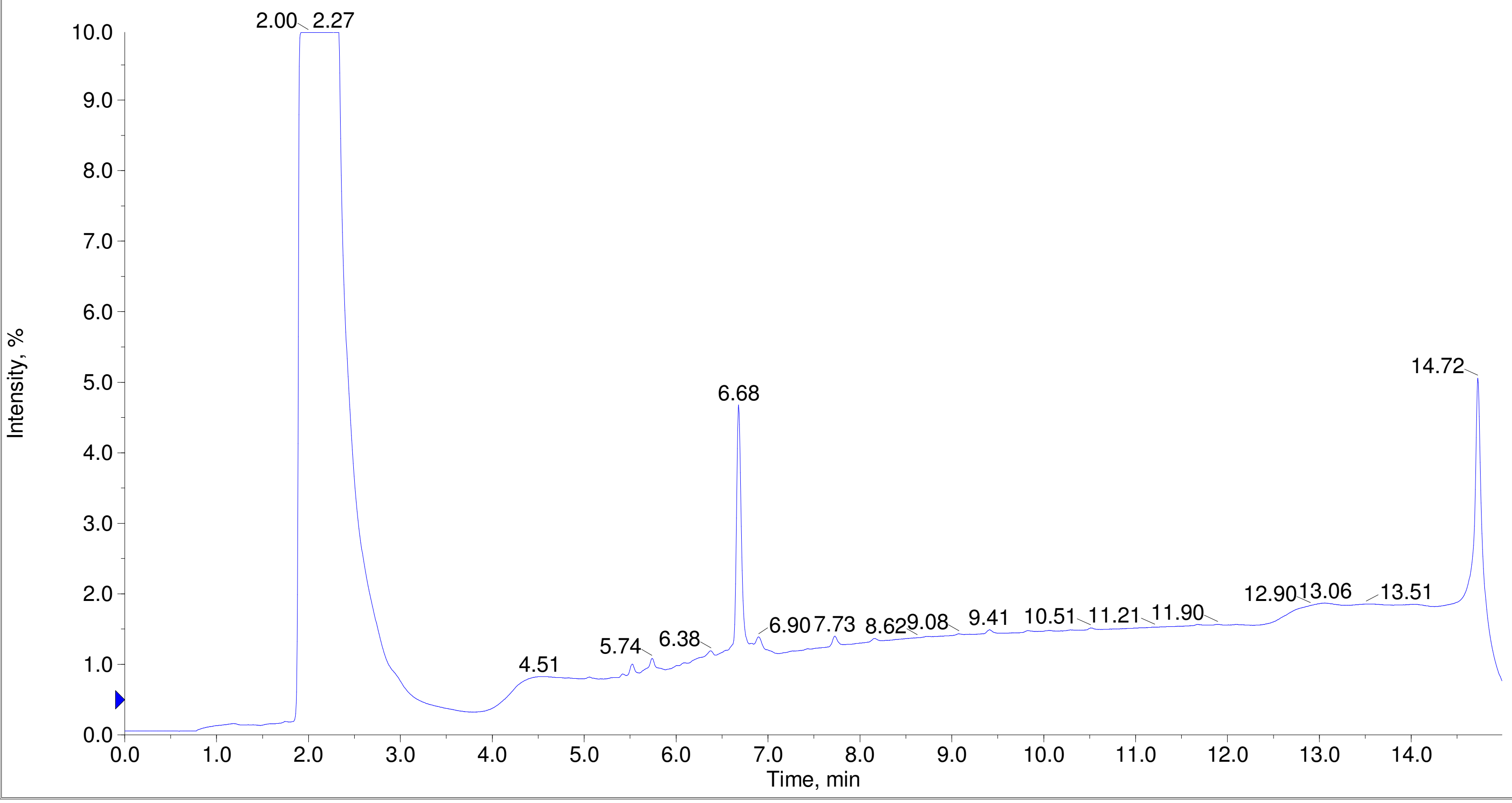

Printing Time: 4:00:09 PM
Printing Date: Tuesday, January 31, 2023
Service Engineer Chris Maier

Post PM
*Stanford U

Polarity/Scan Type: Positive Q1 MS
Sample Name: F7-35-P4-LS-OSF-num27-C3
*API 3000 SN D7640012

TIC of +Q1: from Sample 8 (F7-35-P4-LS-OSF-num27-C3) of Data20210715.wiff (Turbo Spray)    Max. 7.4e8 cps.

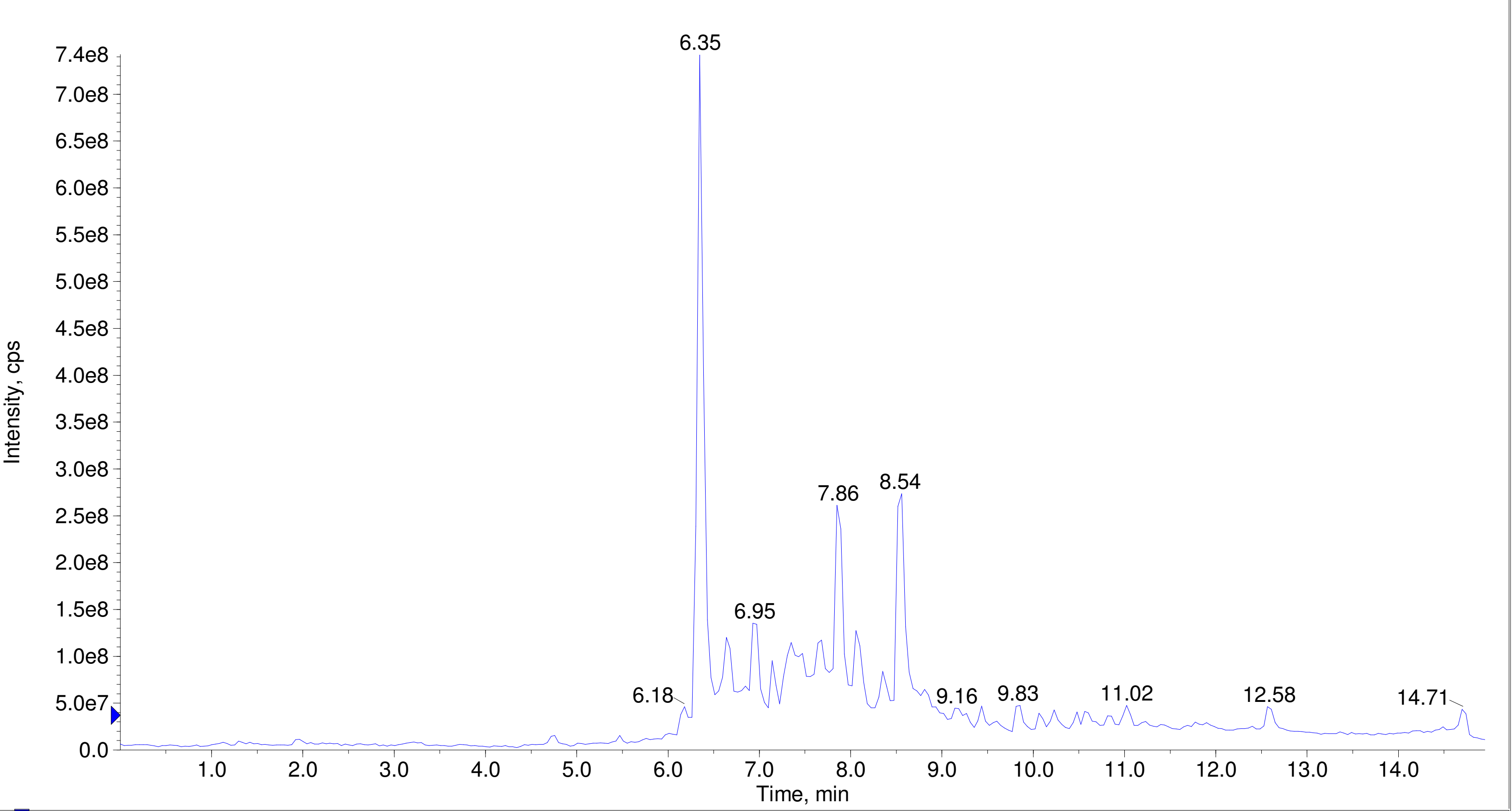


A/D Converter Channel 1 from Sample 8 (F7-35-P4-LS-OSF-num27-C3) of Data20210715.wiff    Max. 10.0 %.

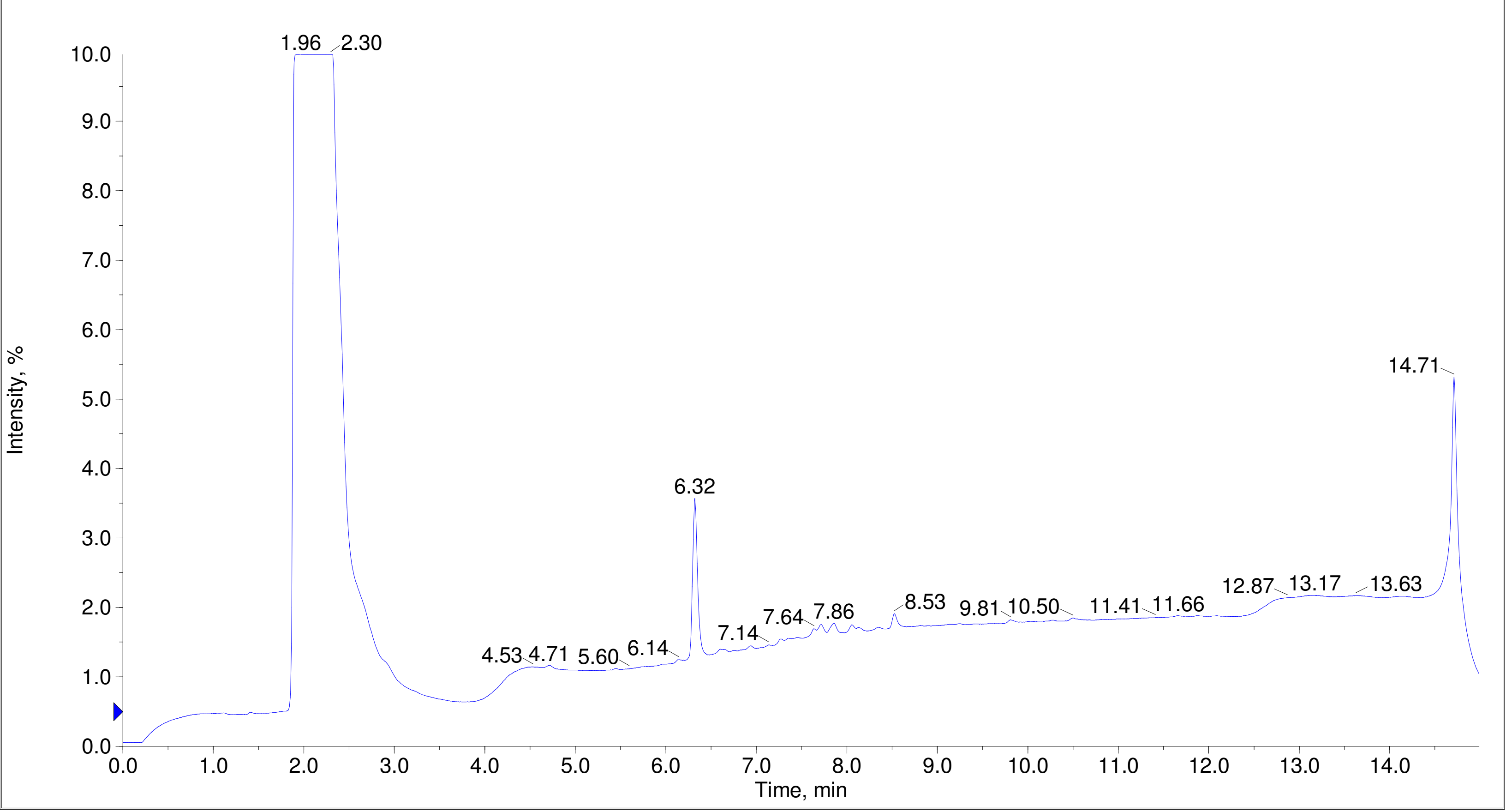

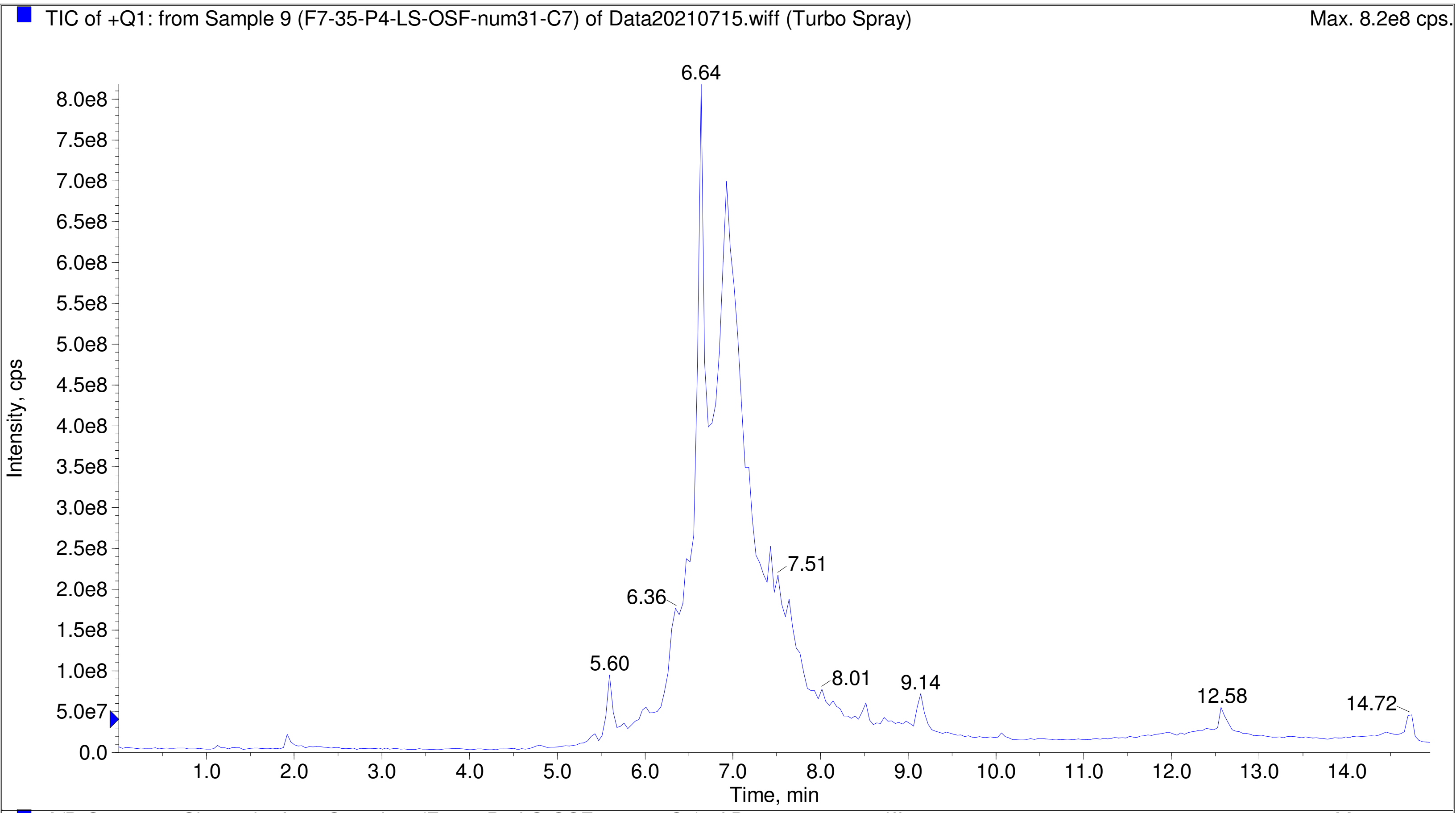
TIC of +Q1: from Sample 9 (F7-35-P4-LS-OSF-num31-C7) of Data20210715.wiff (Turbo Spray)
Max. 8.2e8 cps.
Intensity, cps
Time, min
6.64
7.51
6.36
5.60
8.01
9.14
12.58
14.72

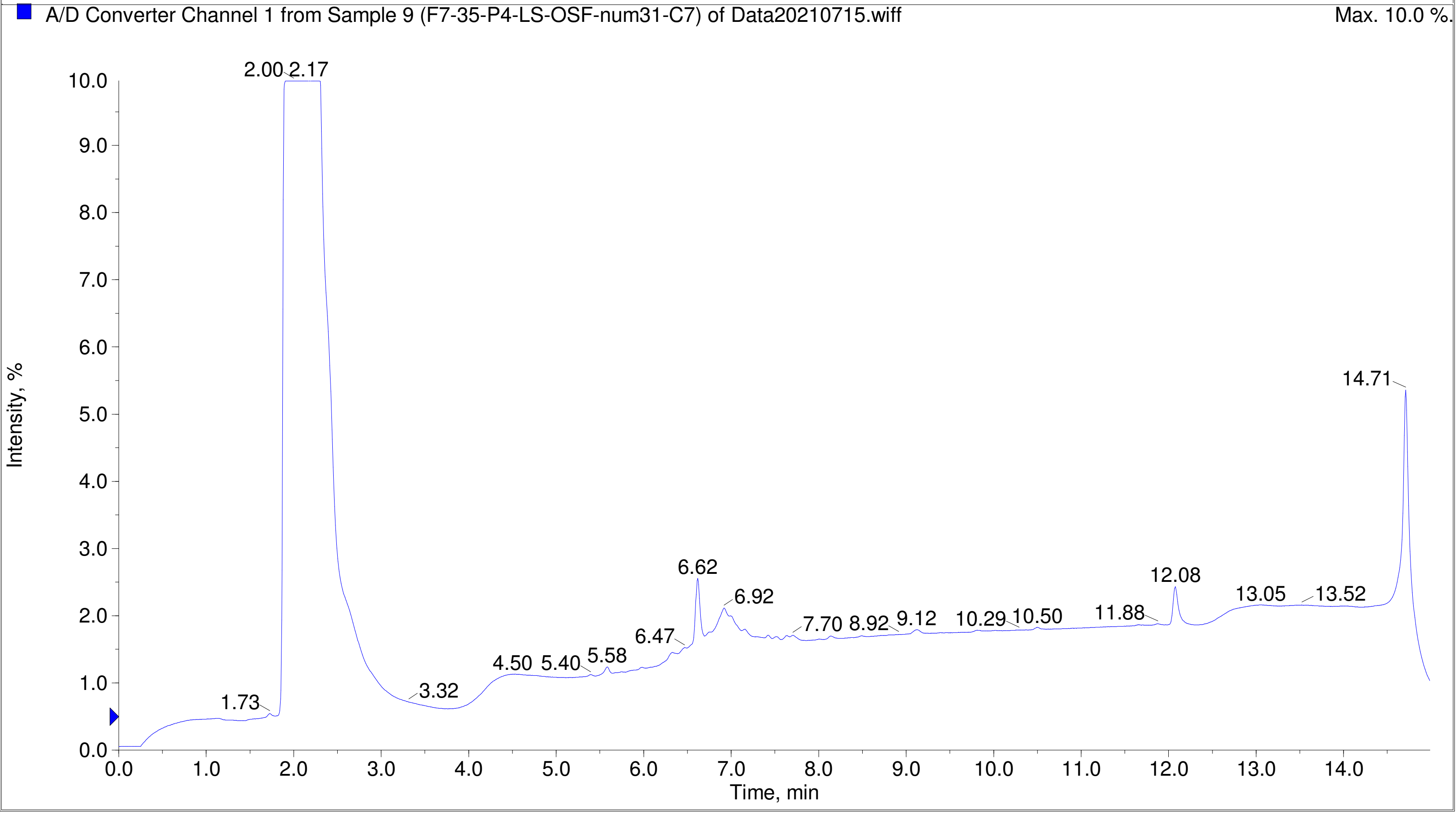
A/D Converter Channel 1 from Sample 9 (F7-35-P4-LS-OSF-num31-C7) of Data20210715.wiff
Max. 10.0 %.
Intensity, %
Time, min
2.00
2.17
14.71
6.62
6.92
12.08
13.05
13.52
7.70
8.92
9.12
10.29
10.50
11.88
6.47
4.50
5.40
5.58
3.32
1.73

Printing Time: 4:02:49 PM | Post PM | Polarity/Scan Type: Positive Q1 MS
Printing Date: Tuesday, January 31, 2023 | | Sample Name: F7-35-P4-LS-OSF-num42-D6
Service Engineer Chris Maier | *Stanford U | *API 3000 SN D7640012

TIC of +Q1: from Sample 10 (F7-35-P4-LS-OSF-num42-D6) of Data20210715.wiff (Turbo Spray) Max. 5.5e8 cps.

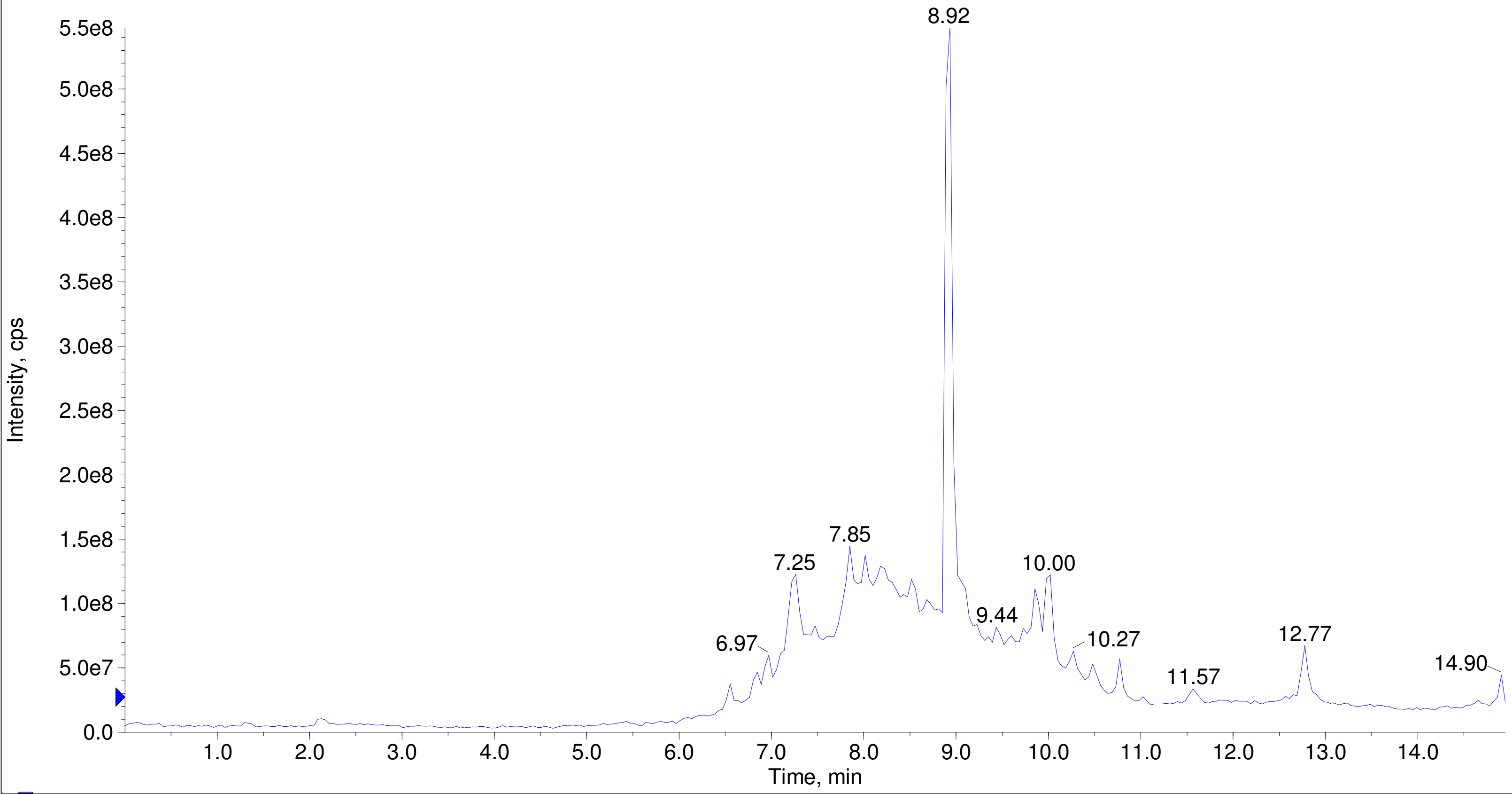


A/D Converter Channel 1 from Sample 10 (F7-35-P4-LS-OSF-num42-D6) of Data20210715.wiff Max. 10.0 %.

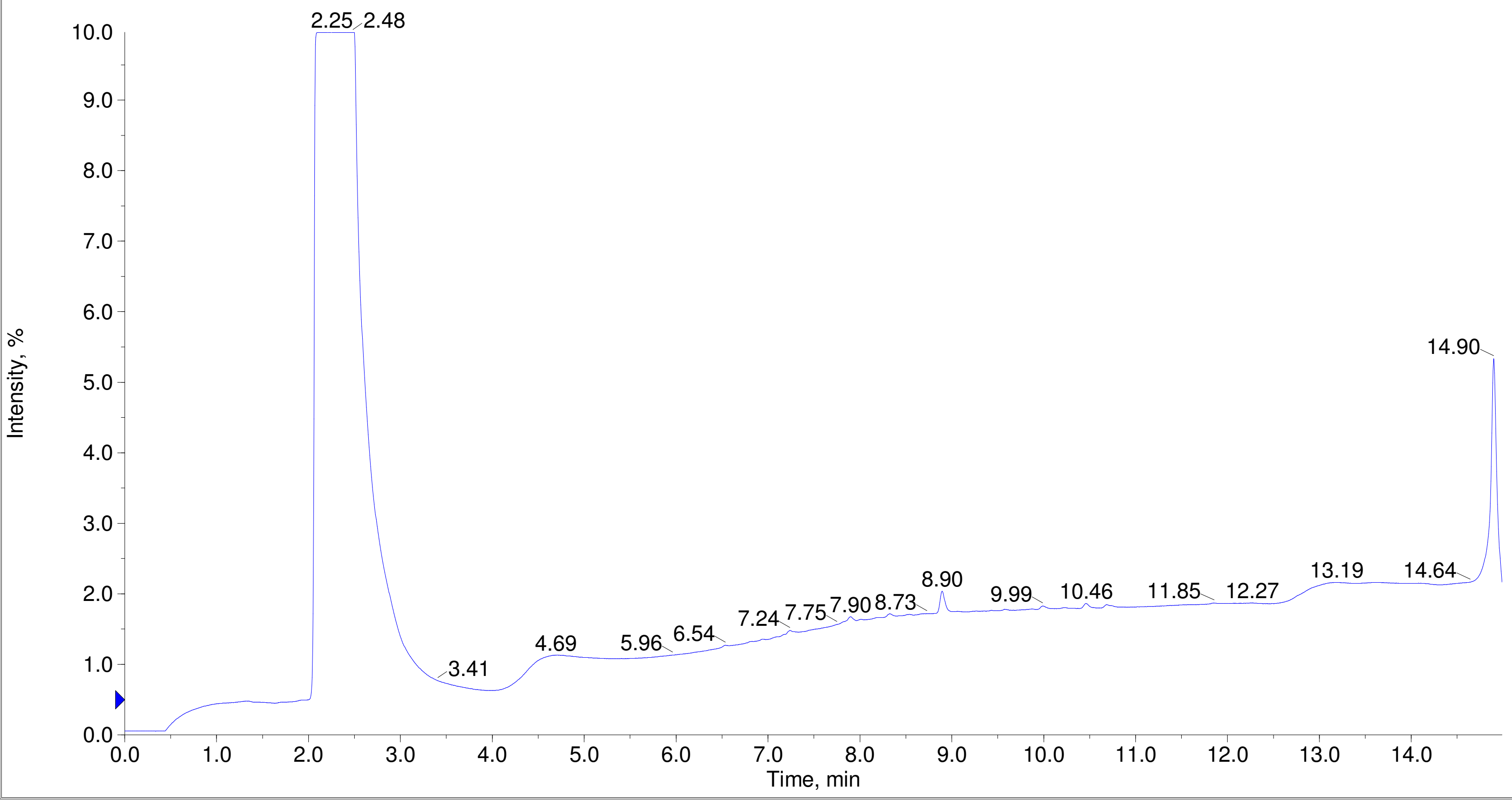

Printing Time: 4:03:59 PM
Printing Date: Tuesday, January 31, 2023
Service Engineer Chris Maier

Post PM
*Stanford U

Polarity/Scan Type: Positive Q1 MS
Sample Name: F7-35-P4-LS-OSF-num48-D12
*API 3000 SN D7640012

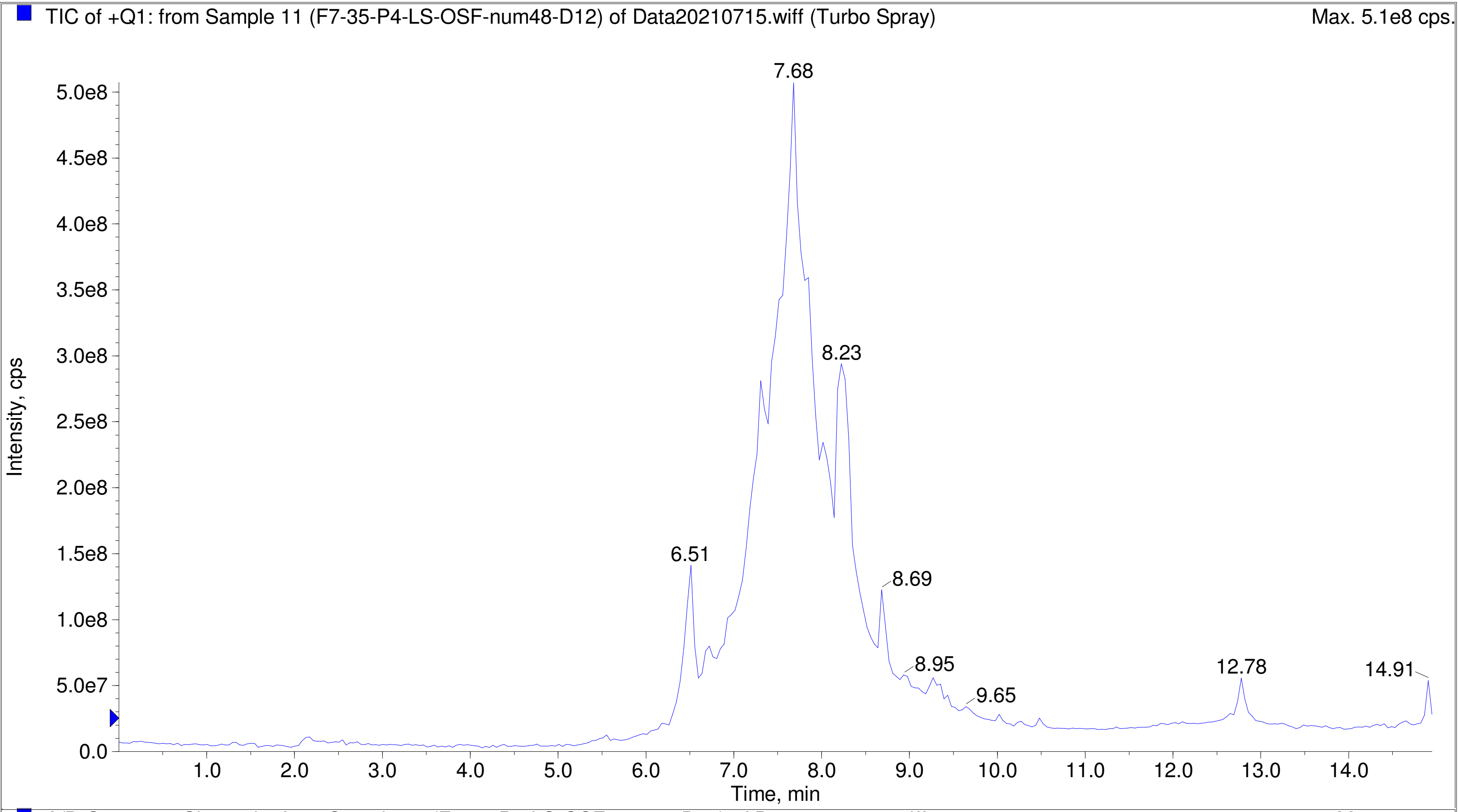


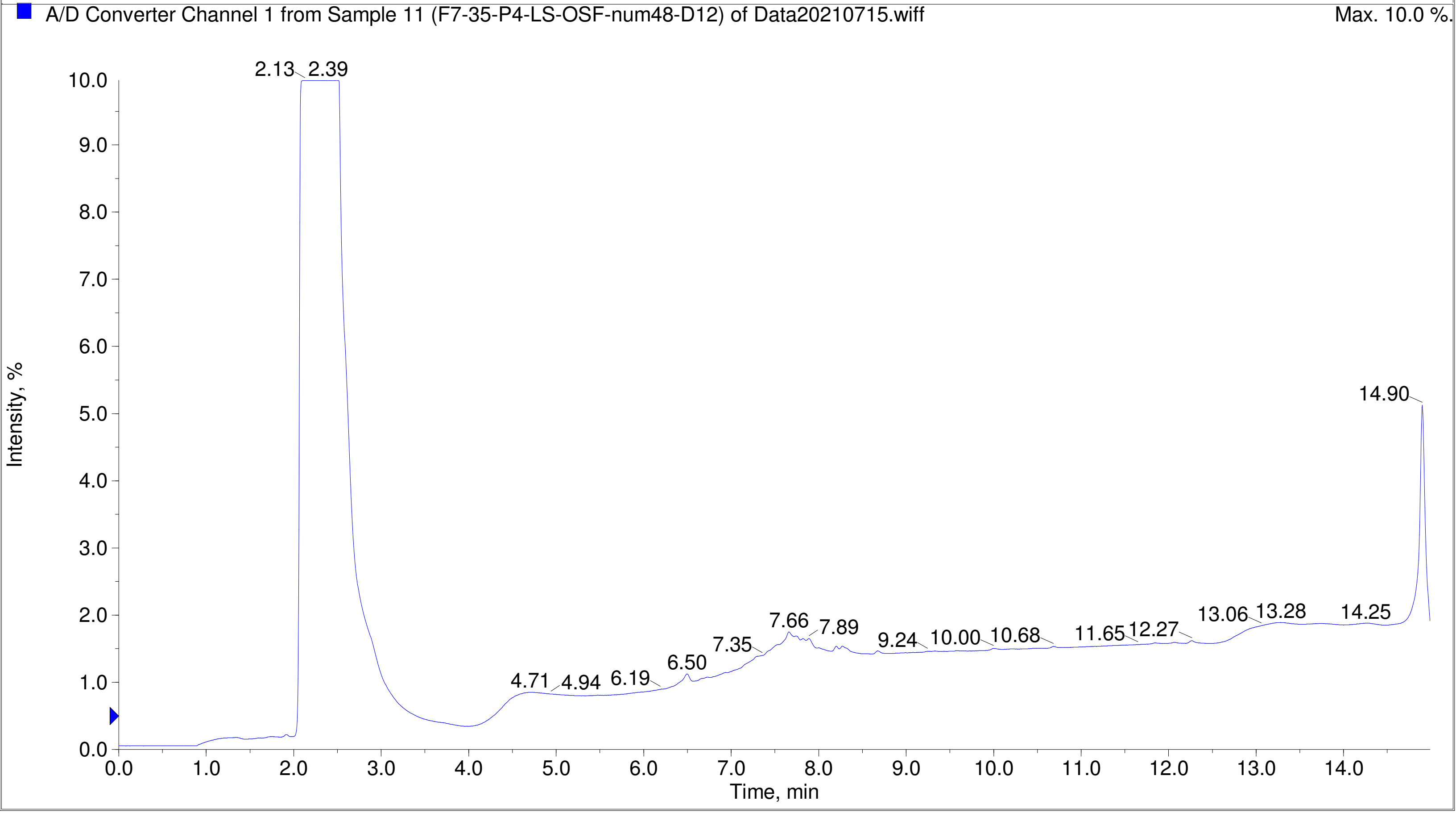

Printing Time: 4:04:59 PM Post PM Polarity/Scan Type: Positive Q1 MS
Printing Date: Tuesday, January 31, 2023 Sample Name: F7-35-P4-LS-OSF-num52-E4
Service Engineer Chris Maier *Stanford U *API 3000 SN D7640012

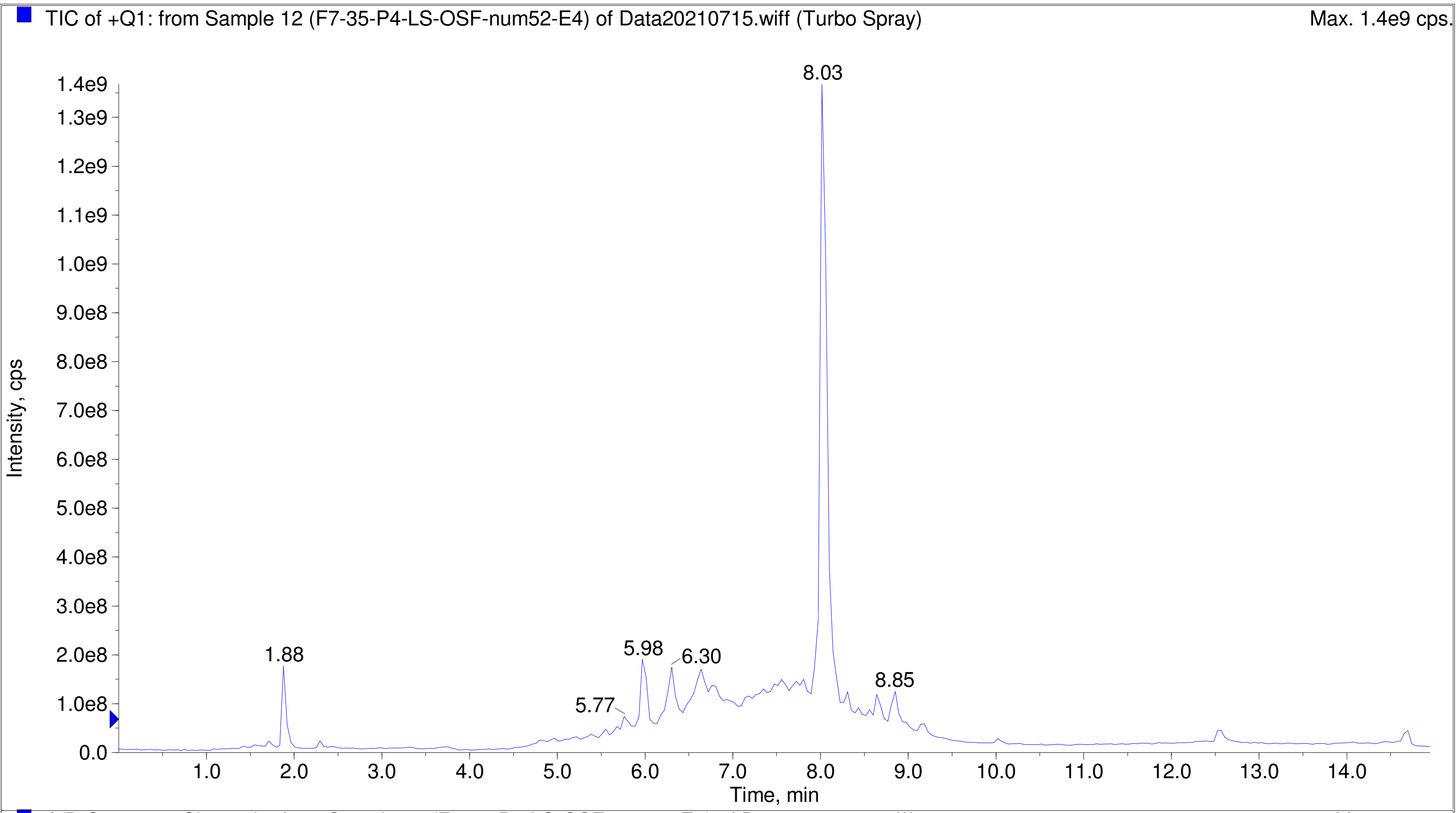


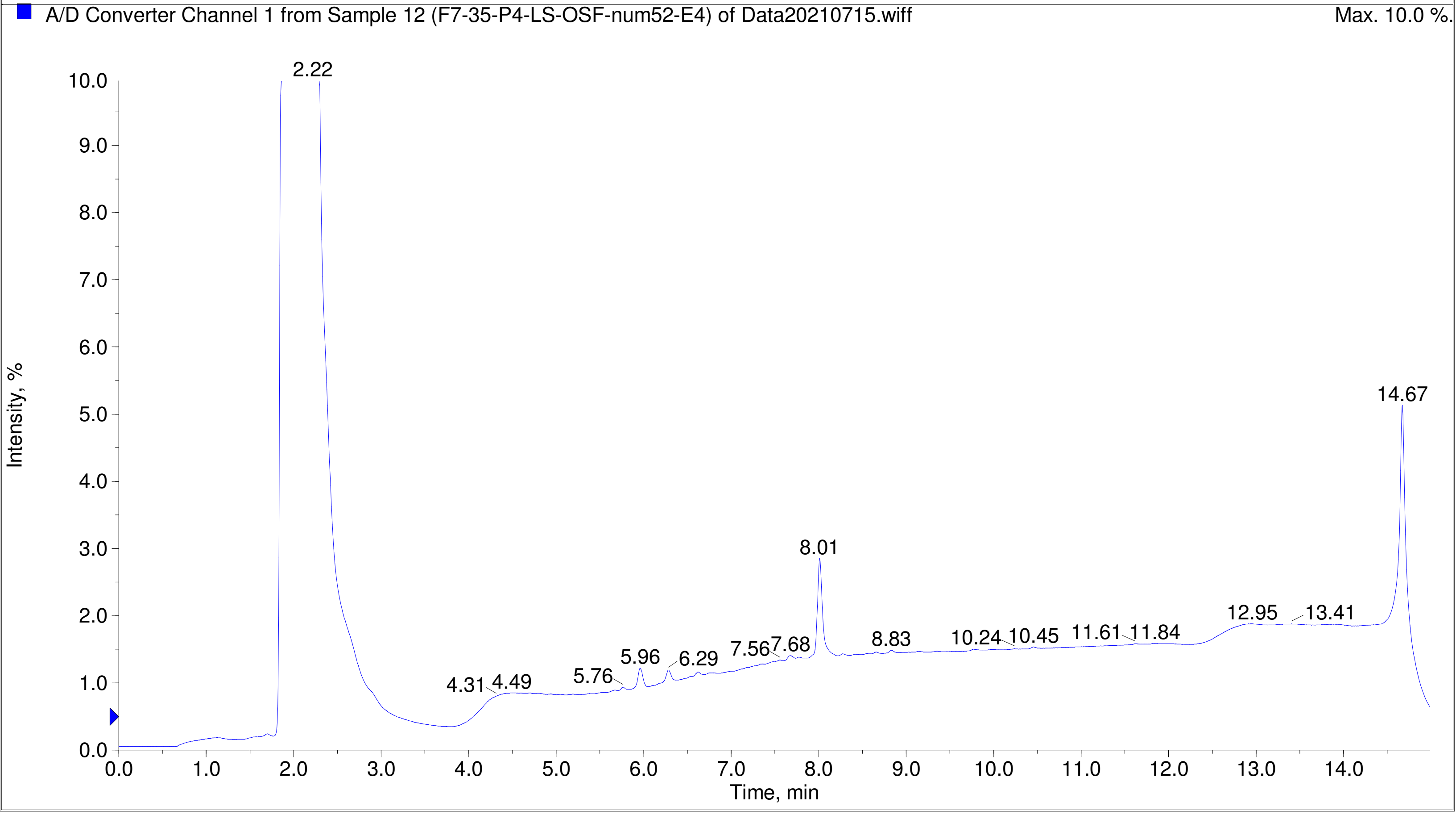

Printing Time: 2:43:12 PM Post PM Polarity/Scan Type: Positive Q1 MS
Printing Date: Monday, January 30, 2023 Sample Name: F7-35-P5-KS-OSF-num1-A1
Service Engineer Chris Maier *Stanford U *API 3000 SN D7640012

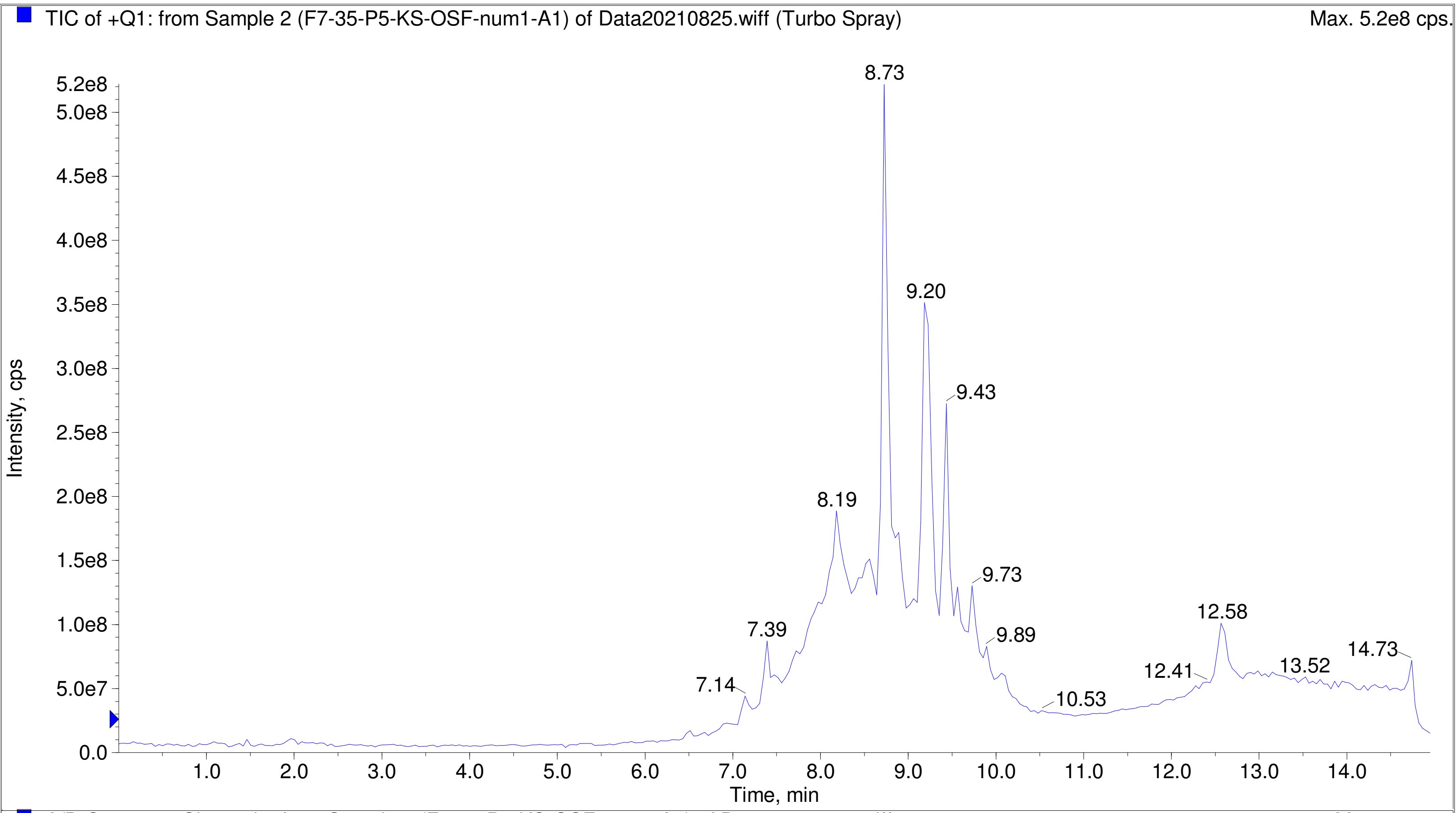


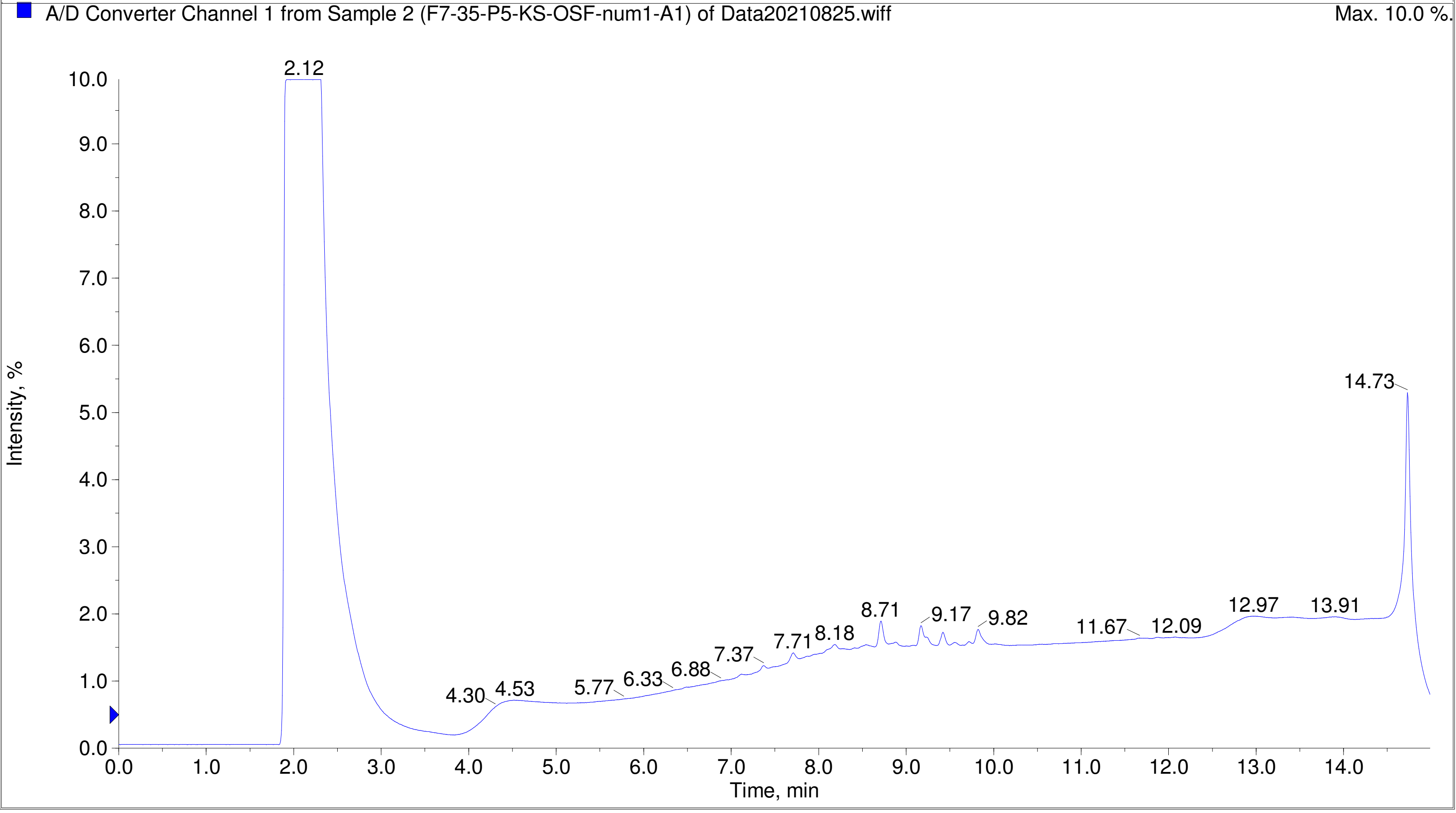

TIC of +Q1: from Sample 3 (F7-35-P5-KS-OSF-num3-A3) of Data20210825.wiff (Turbo Spray) Max. 6.7e8 cps.

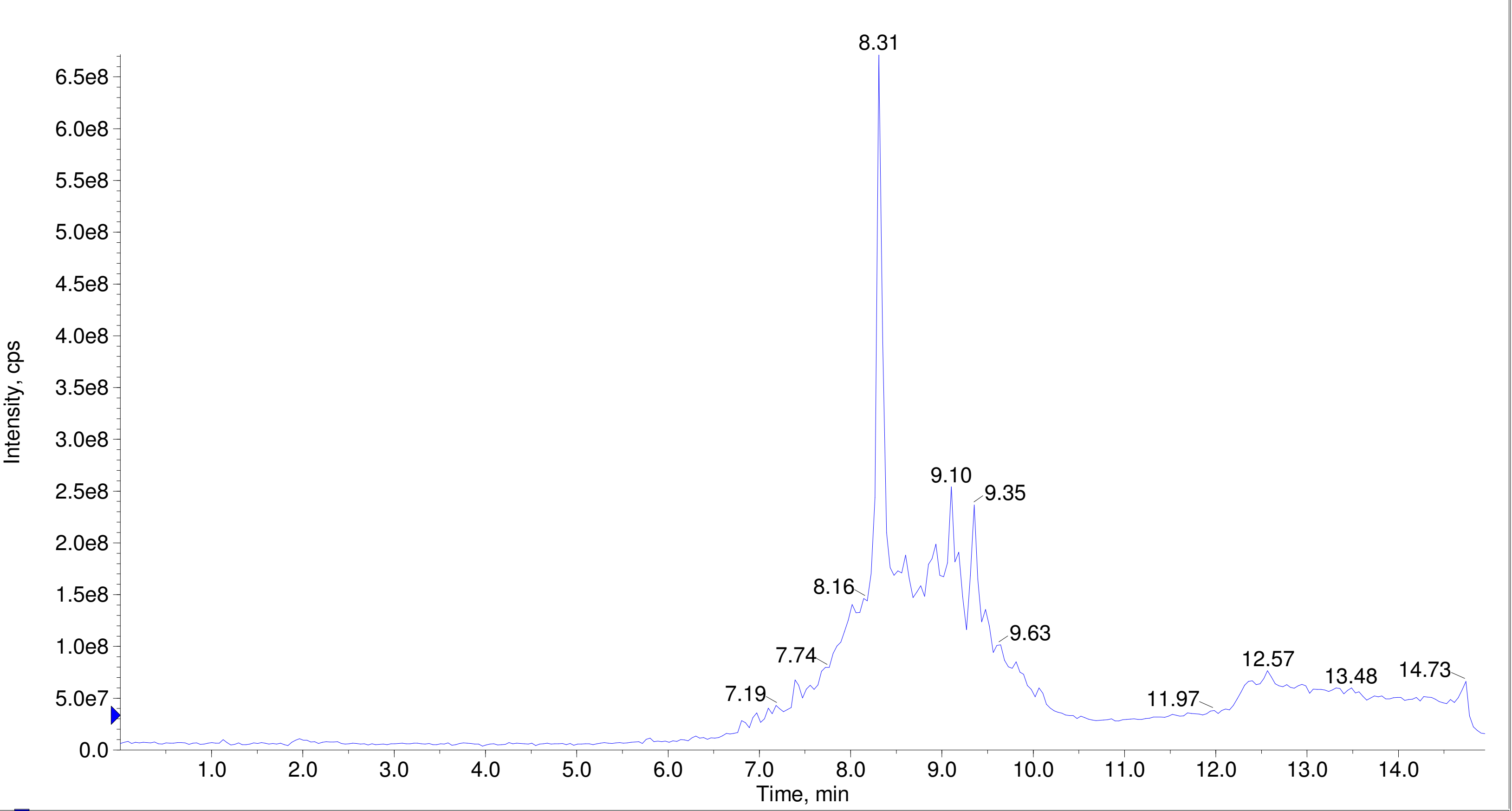


A/D Converter Channel 1 from Sample 3 (F7-35-P5-KS-OSF-num3-A3) of Data20210825.wiff Max. 10.0 %.

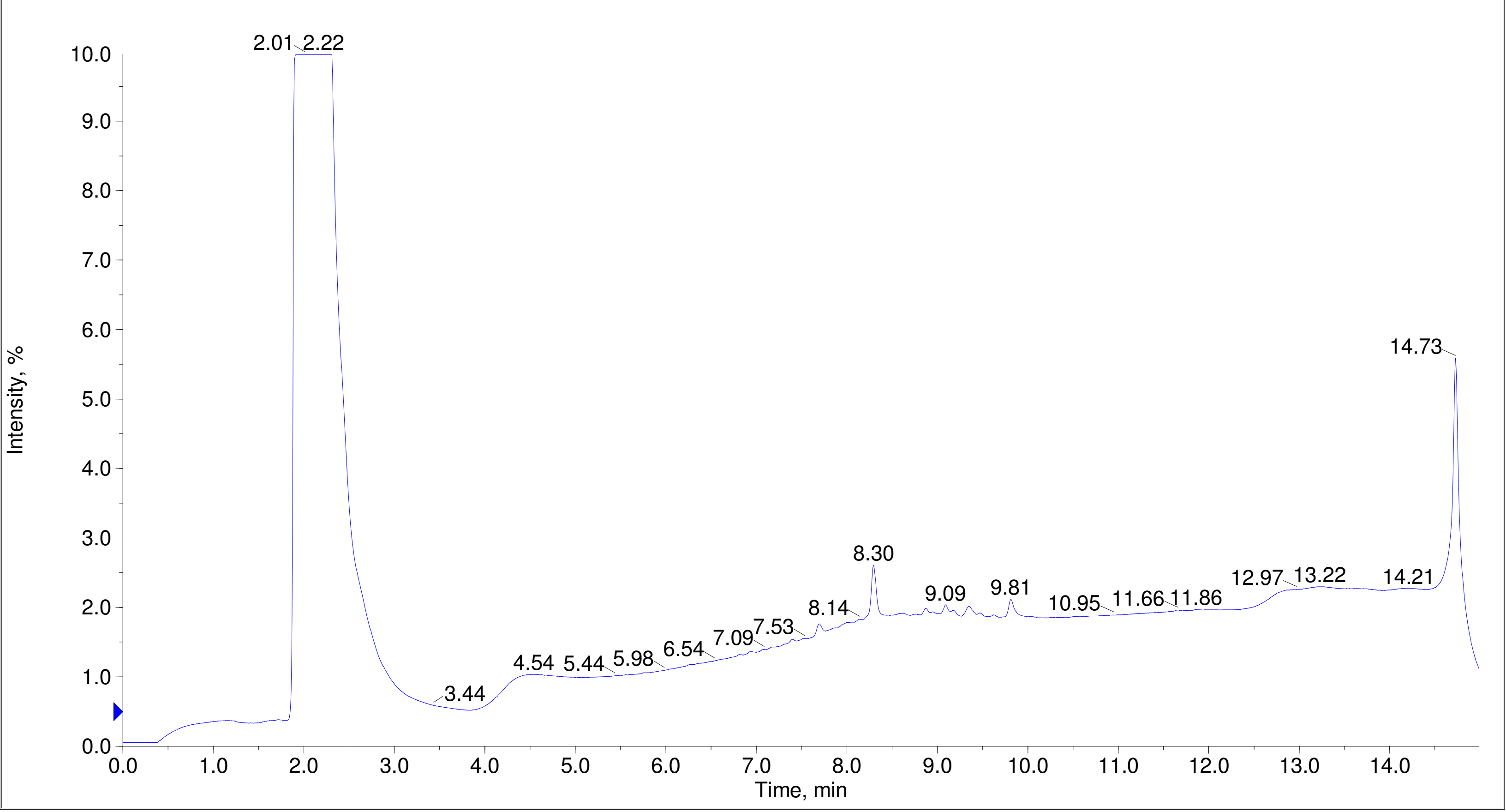

Printing Time: 2:46:17 PM | Post PM | Polarity/Scan Type: Positive Q1 MS
Printing Date: Monday, January 30, 2023 | Sample Name: F7-35-P5-KS-OSF-num10-A10
Service Engineer Chris Maier | *Stanford U | *API 3000 SN D7640012

TIC of +Q1: from Sample 4 (F7-35-P5-KS-OSF-num10-A10) of Data20210825.wiff (Turbo Spray) Max. 6.6e8 cps.

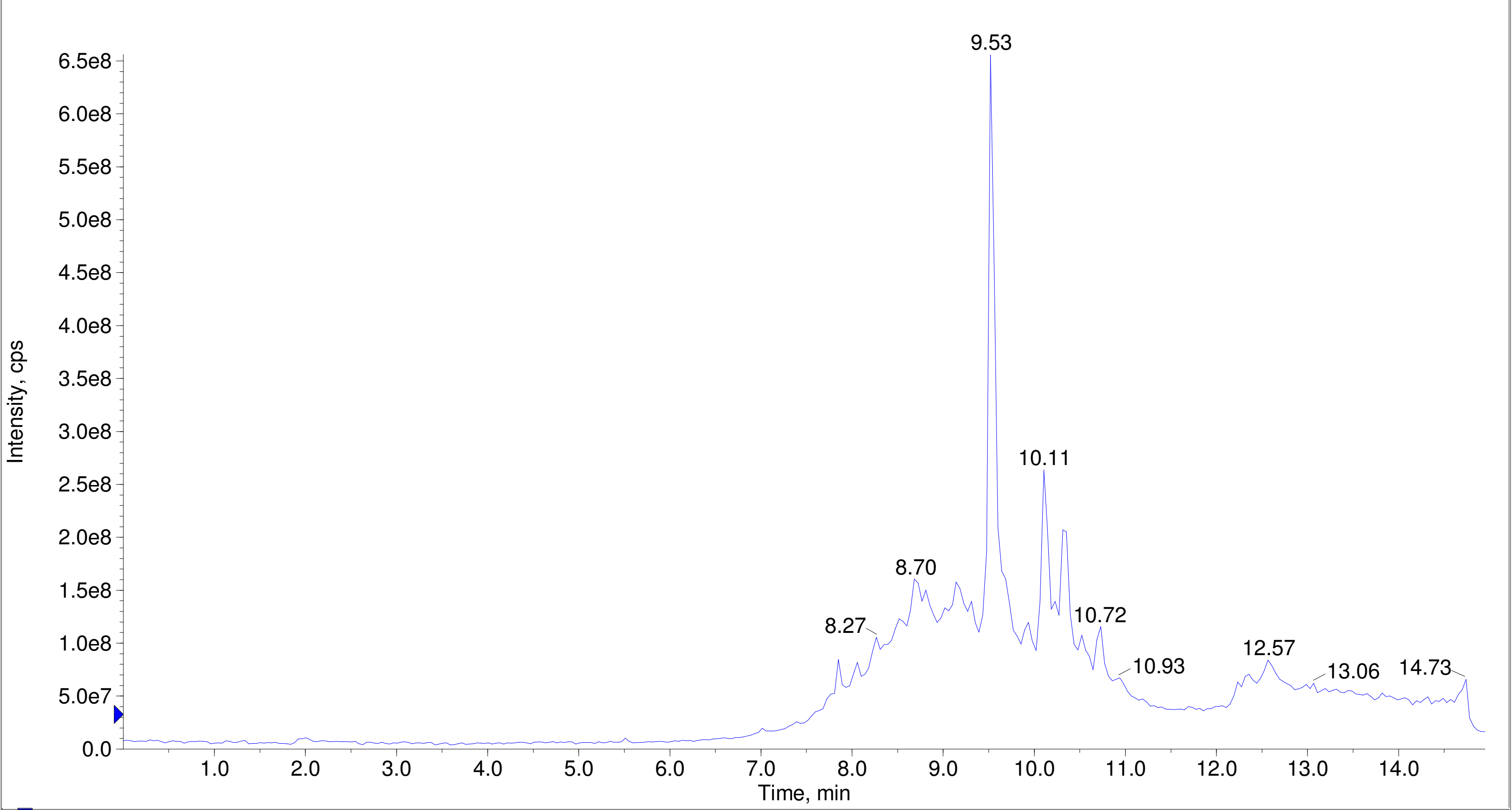


A/D Converter Channel 1 from Sample 4 (F7-35-P5-KS-OSF-num10-A10) of Data20210825.wiff Max. 10.0 %.

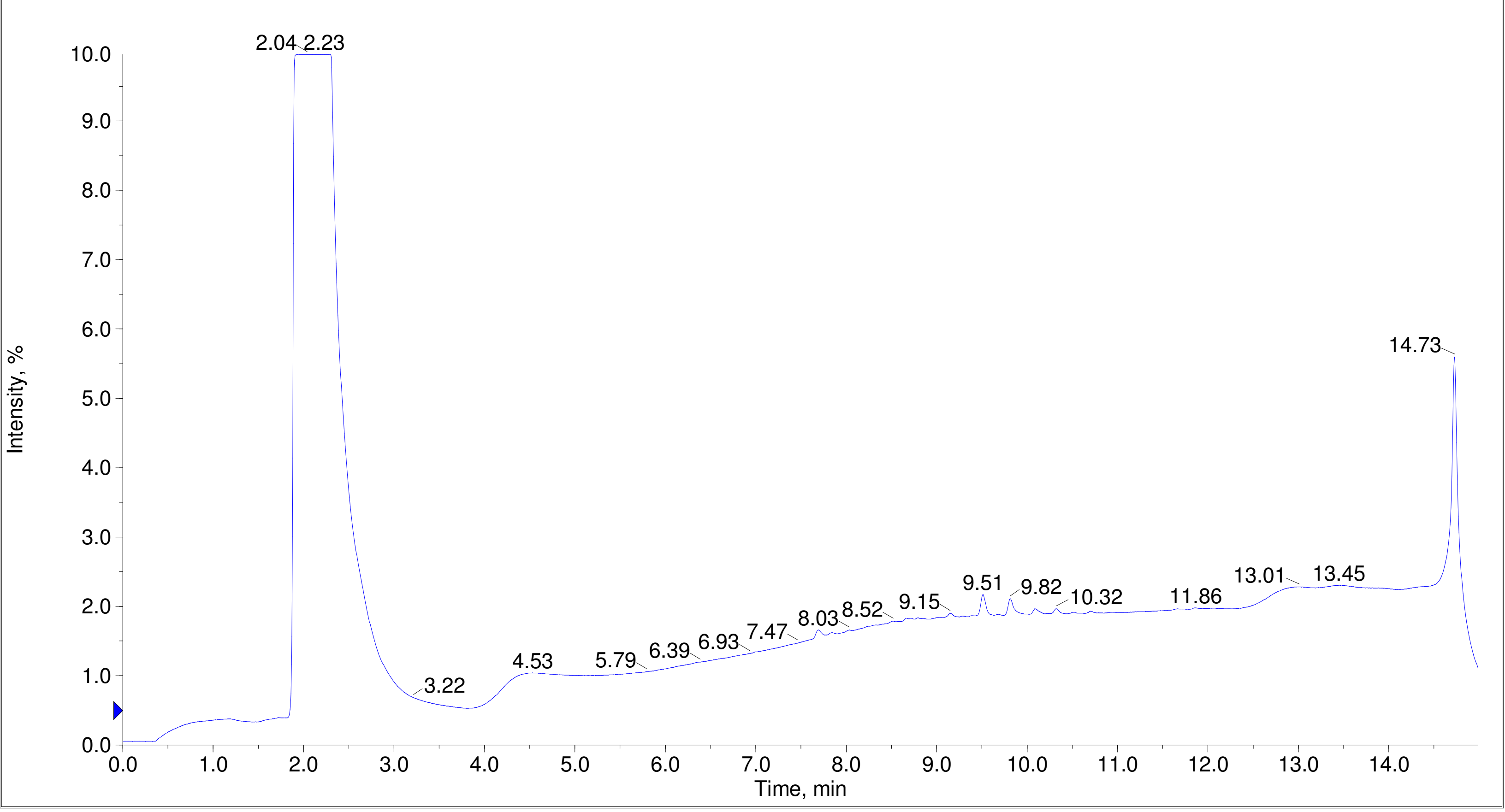

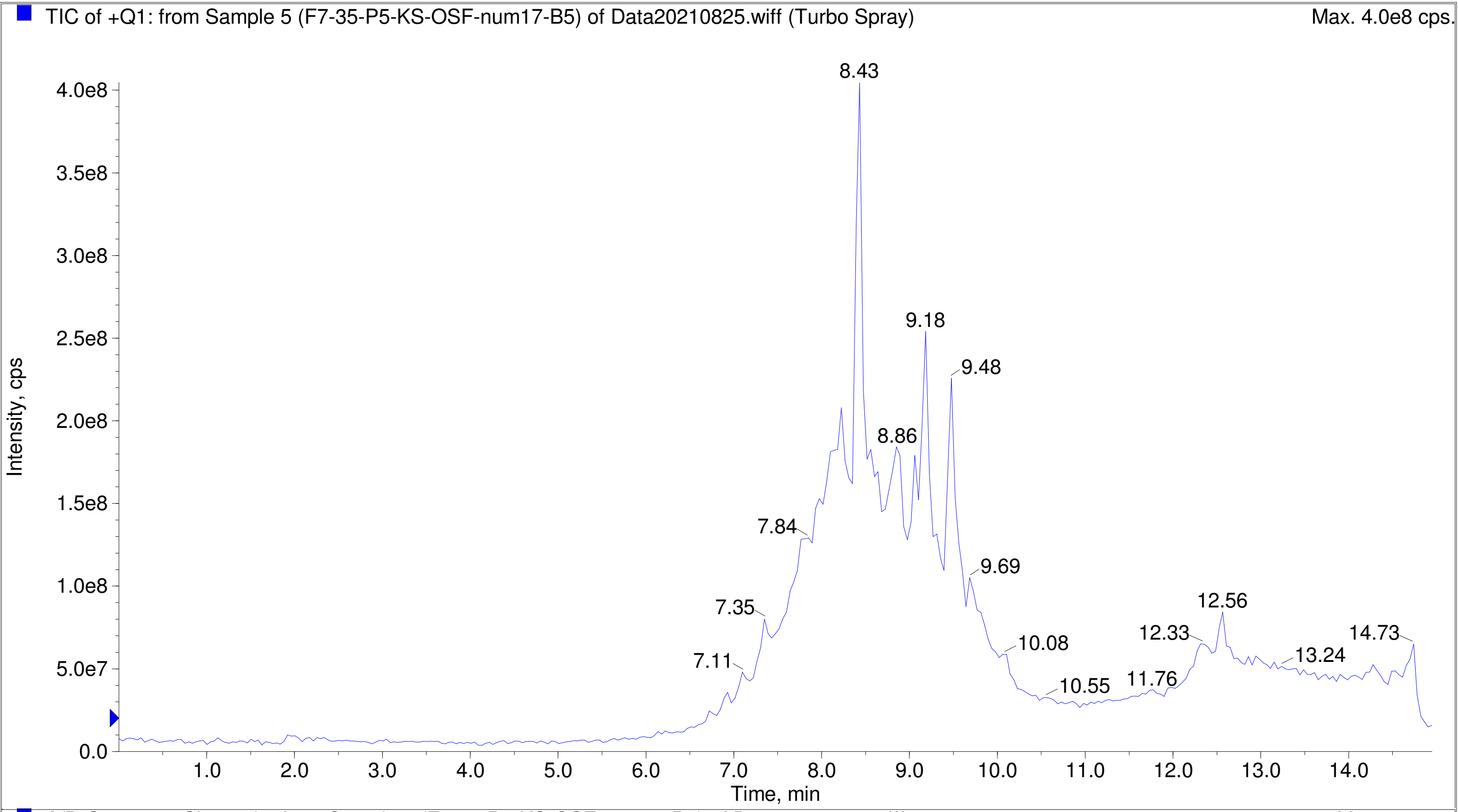
TIC of +Q1: from Sample 5 (F7-35-P5-KS-OSF-num17-B5) of Data20210825.wiff (Turbo Spray)
Max. 4.0e8 cps.
Intensity, cps
Time, min
8.43
9.18
9.48
8.86
7.84
9.69
7.35
12.56
7.11
10.08
12.33
14.73
13.24
10.55
11.76

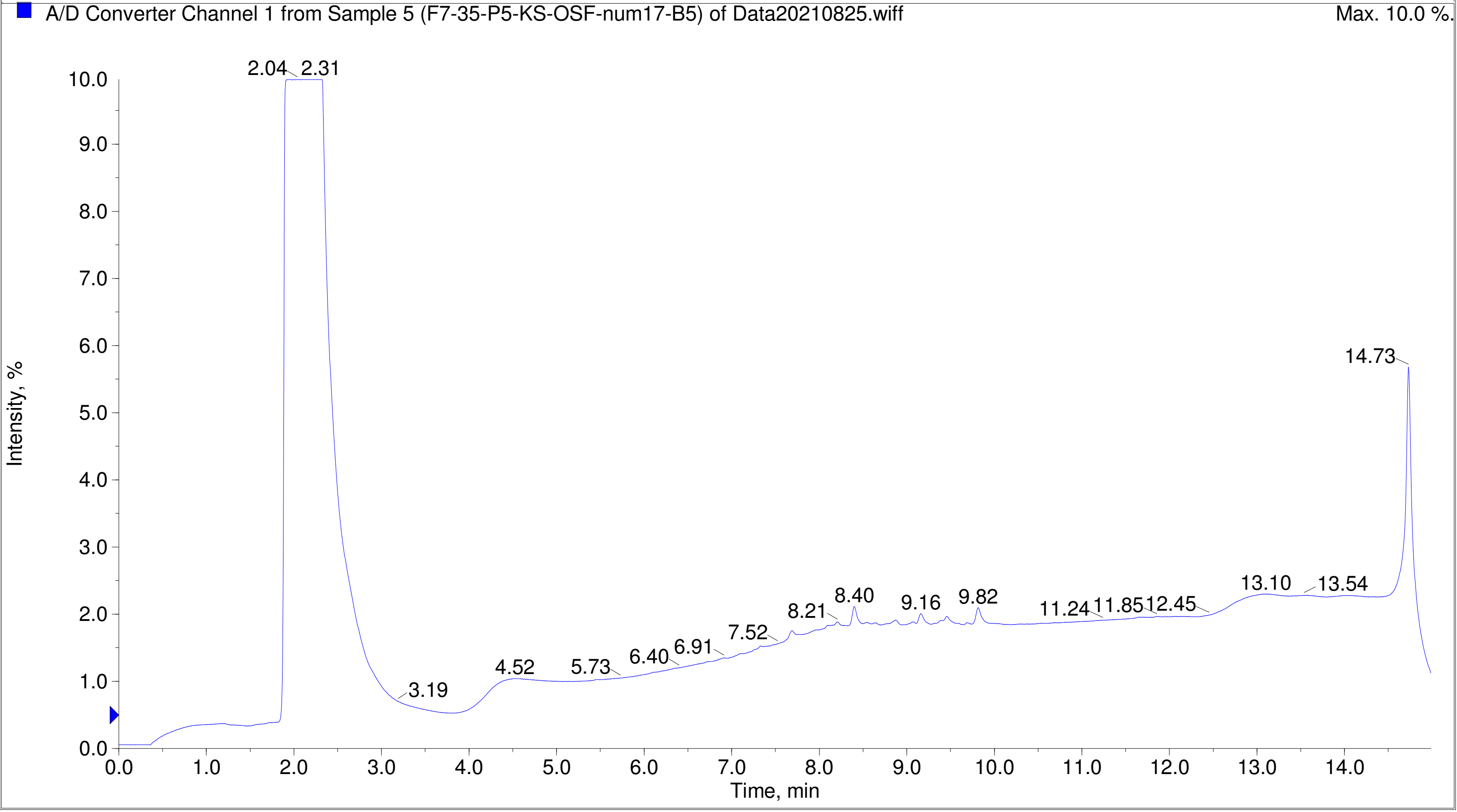
A/D Converter Channel 1 from Sample 5 (F7-35-P5-KS-OSF-num17-B5) of Data20210825.wiff
Max. 10.0 %.
Intensity, %
Time, min
2.04
2.31
14.73
3.19
4.52
5.73
6.40
6.91
7.52
8.21
8.40
9.16
9.82
11.24
11.85
12.45
13.10
13.54

Printing Time: 2:50:06 PM
Printing Date: Monday, January 30, 2023
Service Engineer Chris Maier

Post PM

*Stanford U

Polarity/Scan Type: Positive Q1 MS
Sample Name: F7-35-P5-KS-OSF-num24-B12
*API 3000 SN D7640012

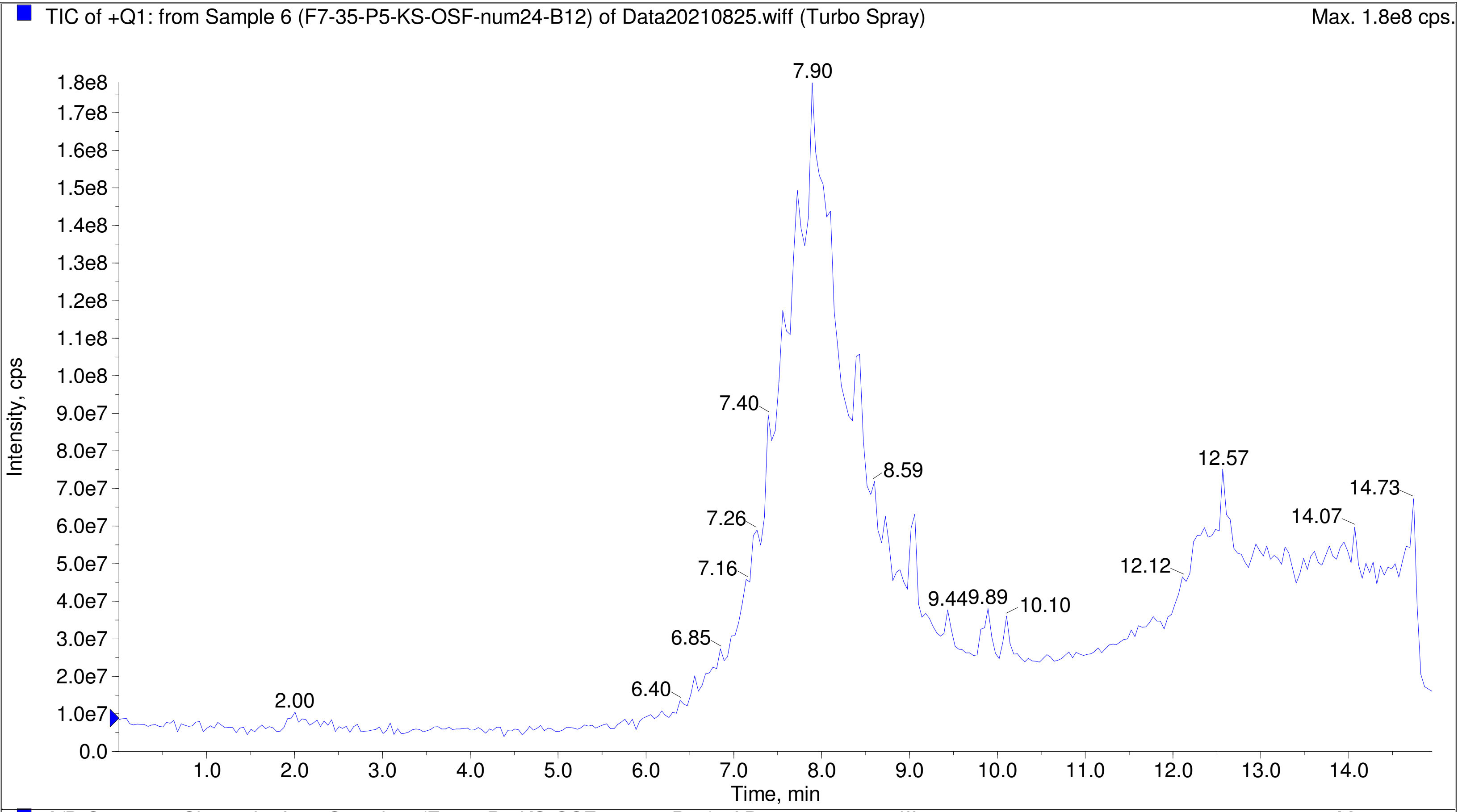


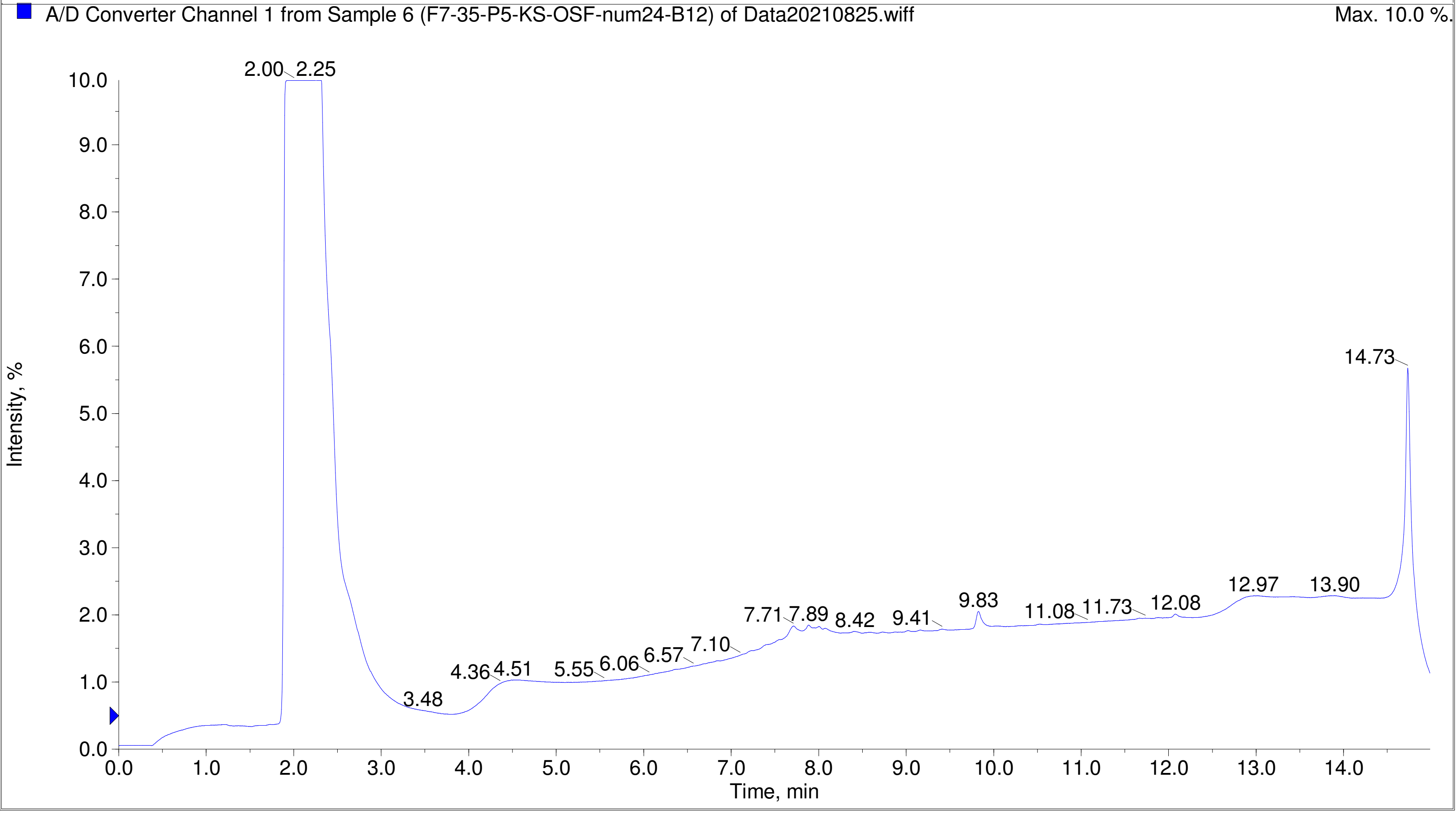

Printing Time: 4:06:42 PM
Printing Date: Tuesday, January 31, 2023
Service Engineer Chris Maier

Post PM
*Stanford U

Polarity/Scan Type: Positive Q1 MS
Sample Name: F7-35-P5-KS-OSF-num30-C6
*API 3000 SN D7640012

TIC of +Q1: from Sample 8 (F7-35-P5-KS-OSF-num30-C6) of Data20210825.wiff (Turbo Spray)

Max. 5.3e8 cps.

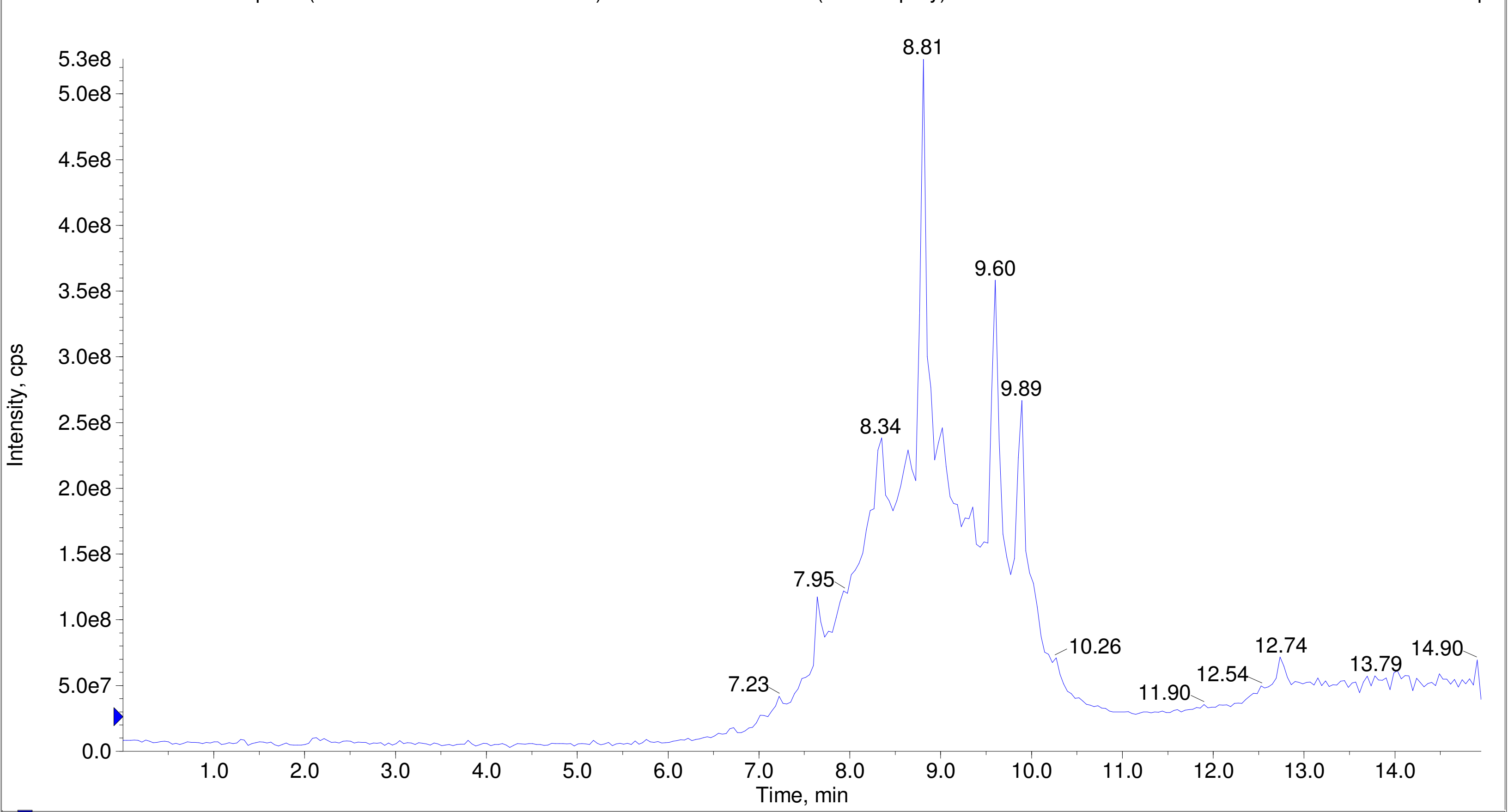


A/D Converter Channel 1 from Sample 8 (F7-35-P5-KS-OSF-num30-C6) of Data20210825.wiff

Max. 10.0 %.

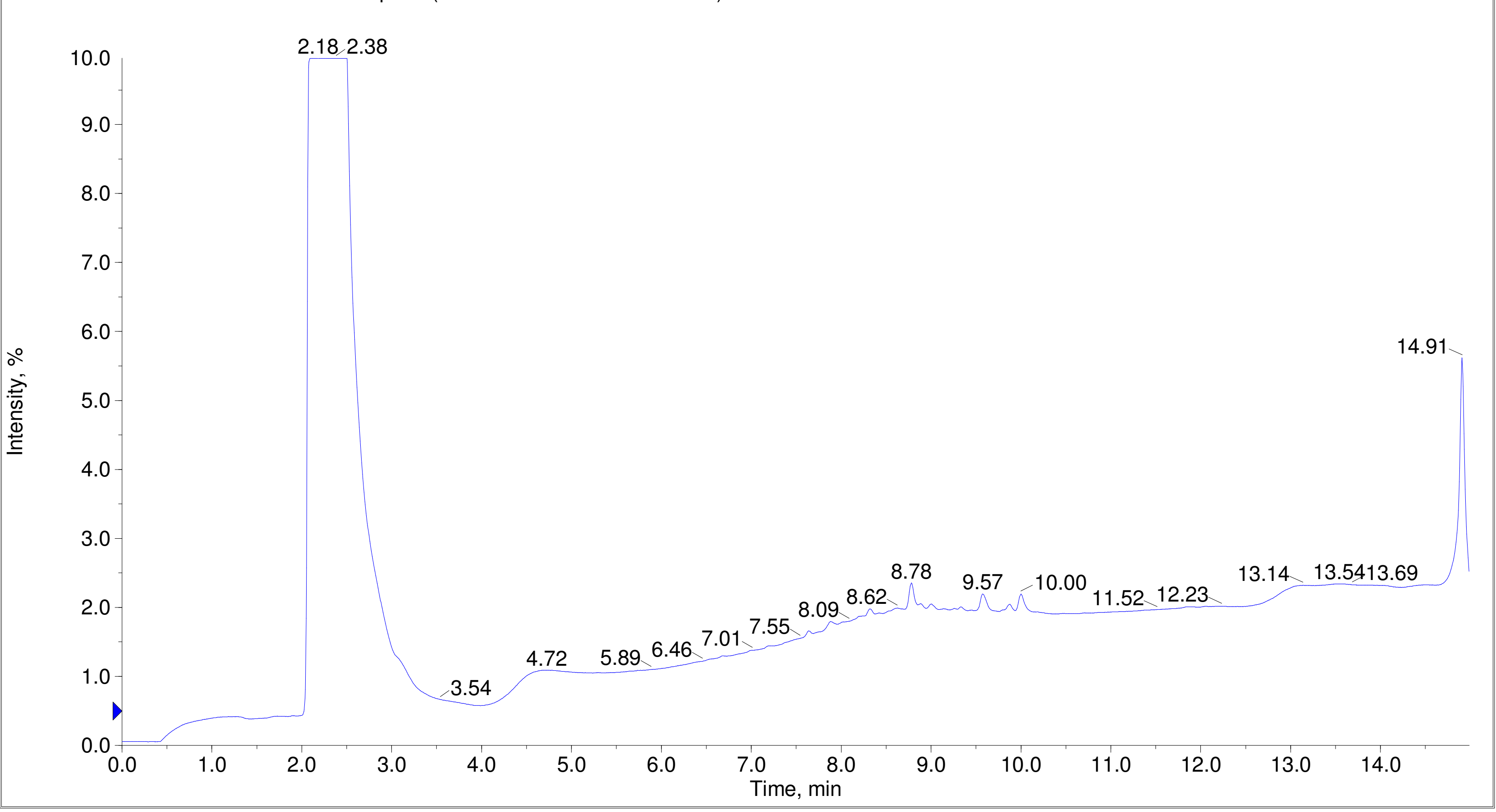

Printing Time: 4:08:38 PM Post PM Polarity/Scan Type: Positive Q1 MS
Printing Date: Tuesday, January 31, 2023 Sample Name: F7-35-P5-KS-OSF-num32-C8
Service Engineer Chris Maier *Stanford U *API 3000 SN D7640012

TIC of +Q1: from Sample 9 (F7-35-P5-KS-OSF-num32-C8) of Data20210825.wiff (Turbo Spray) Max. 4.9e8 cps.

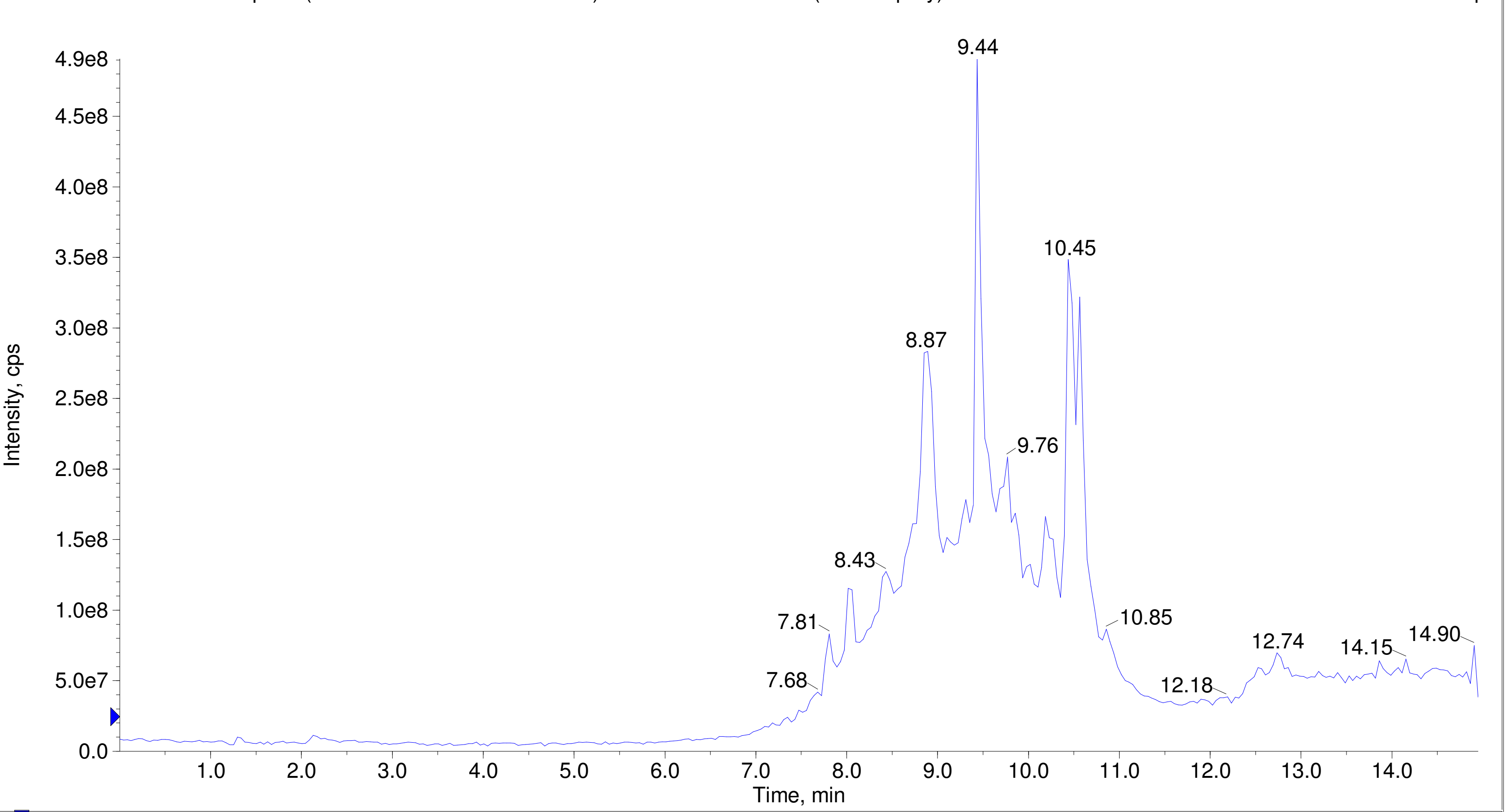


A/D Converter Channel 1 from Sample 9 (F7-35-P5-KS-OSF-num32-C8) of Data20210825.wiff Max. 10.0 %.

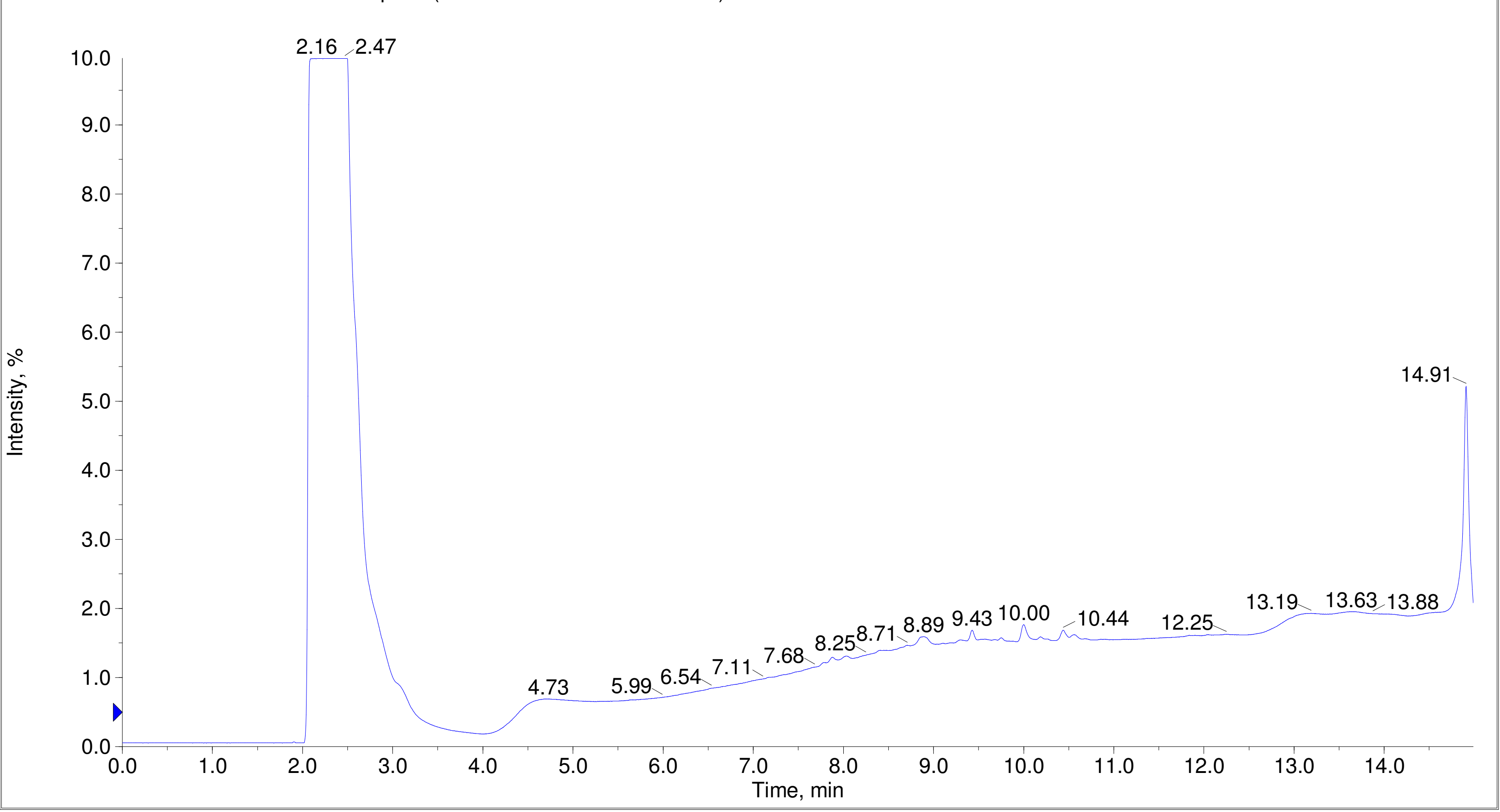

TIC of +Q1: from Sample 10 (F7-35-P5-KS-OSF-num39-D3) of Data20210825.wiff (Turbo Spray) Max. 5.0e8 cps.

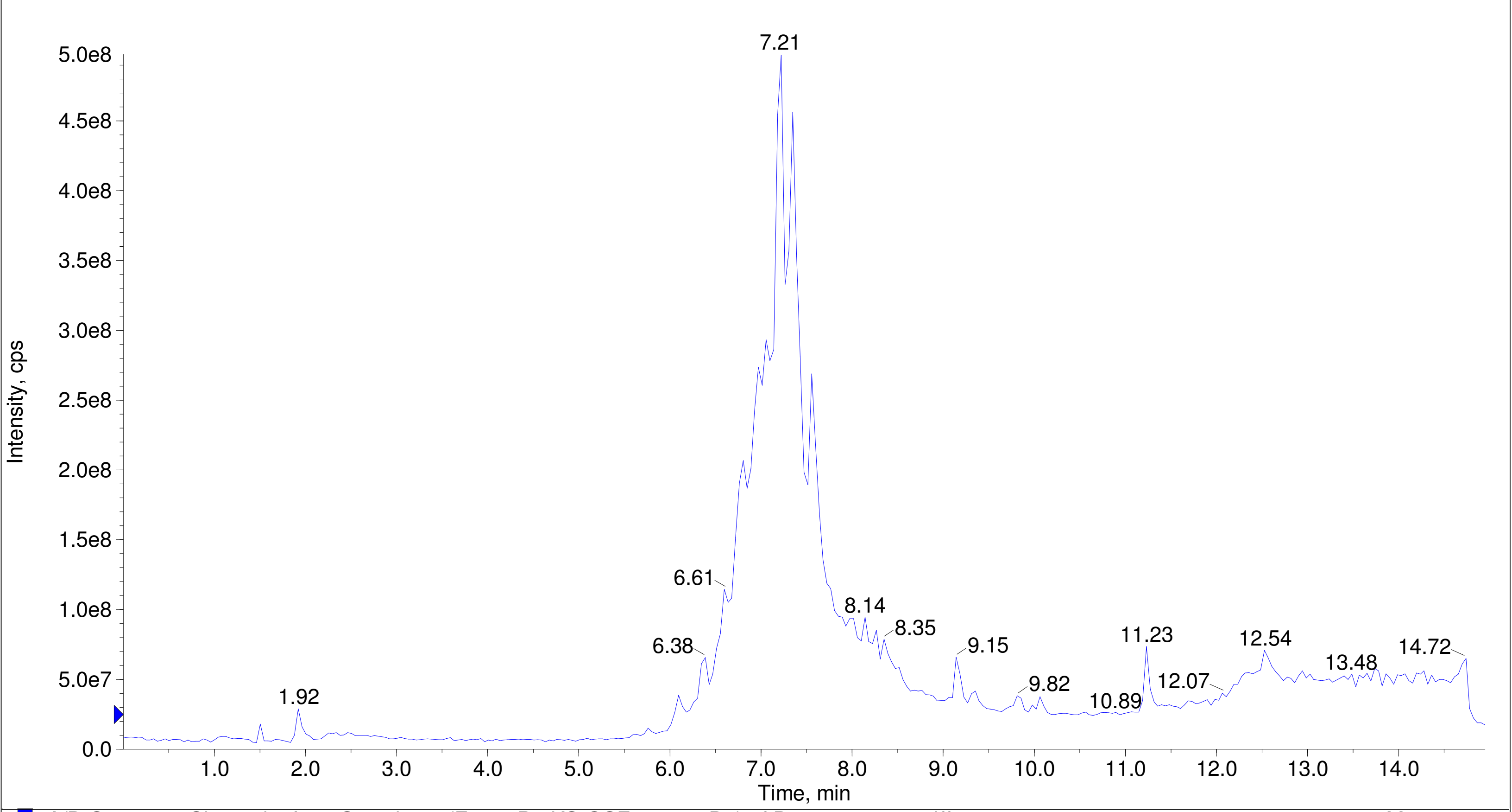


A/D Converter Channel 1 from Sample 10 (F7-35-P5-KS-OSF-num39-D3) of Data20210825.wiff Max. 10.0 %.

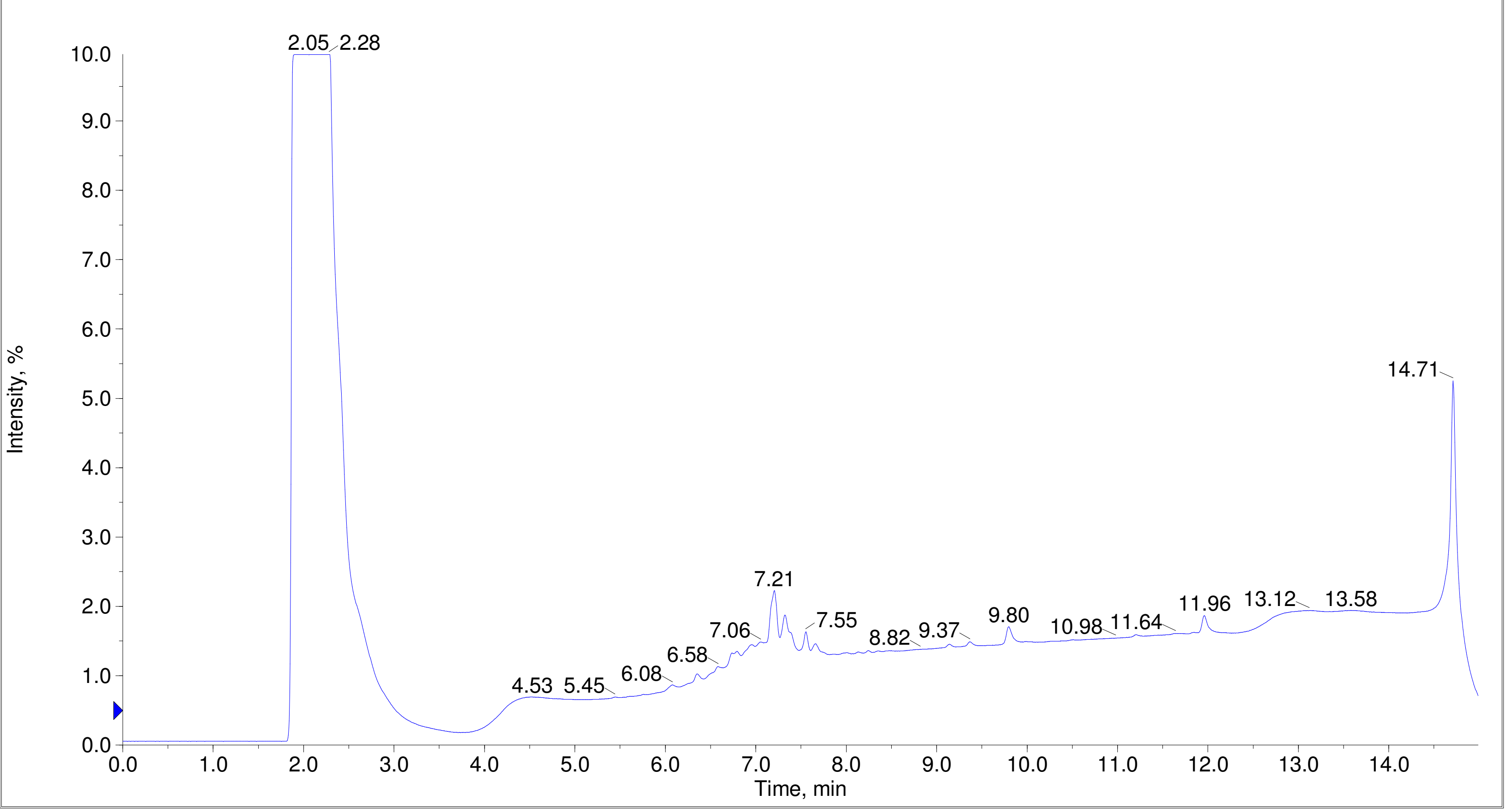

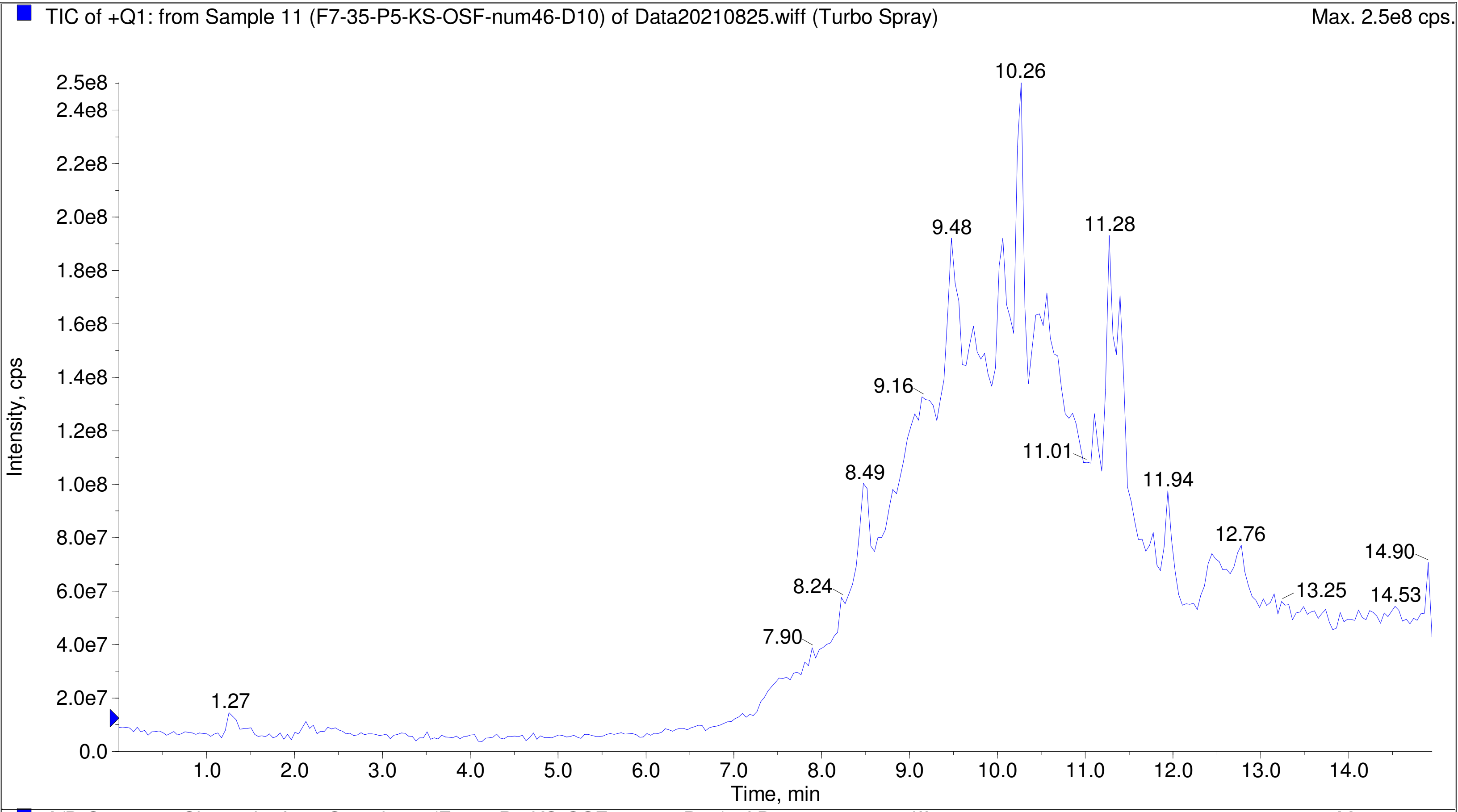
TIC of +Q1: from Sample 11 (F7-35-P5-KS-OSF-num46-D10) of Data20210825.wiff (Turbo Spray)
Max. 2.5e8 cps.
Intensity, cps
Time, min
10.26
9.48
11.28
9.16
11.01
8.49
11.94
12.76
14.90
8.24
13.25
14.53
7.90
1.27

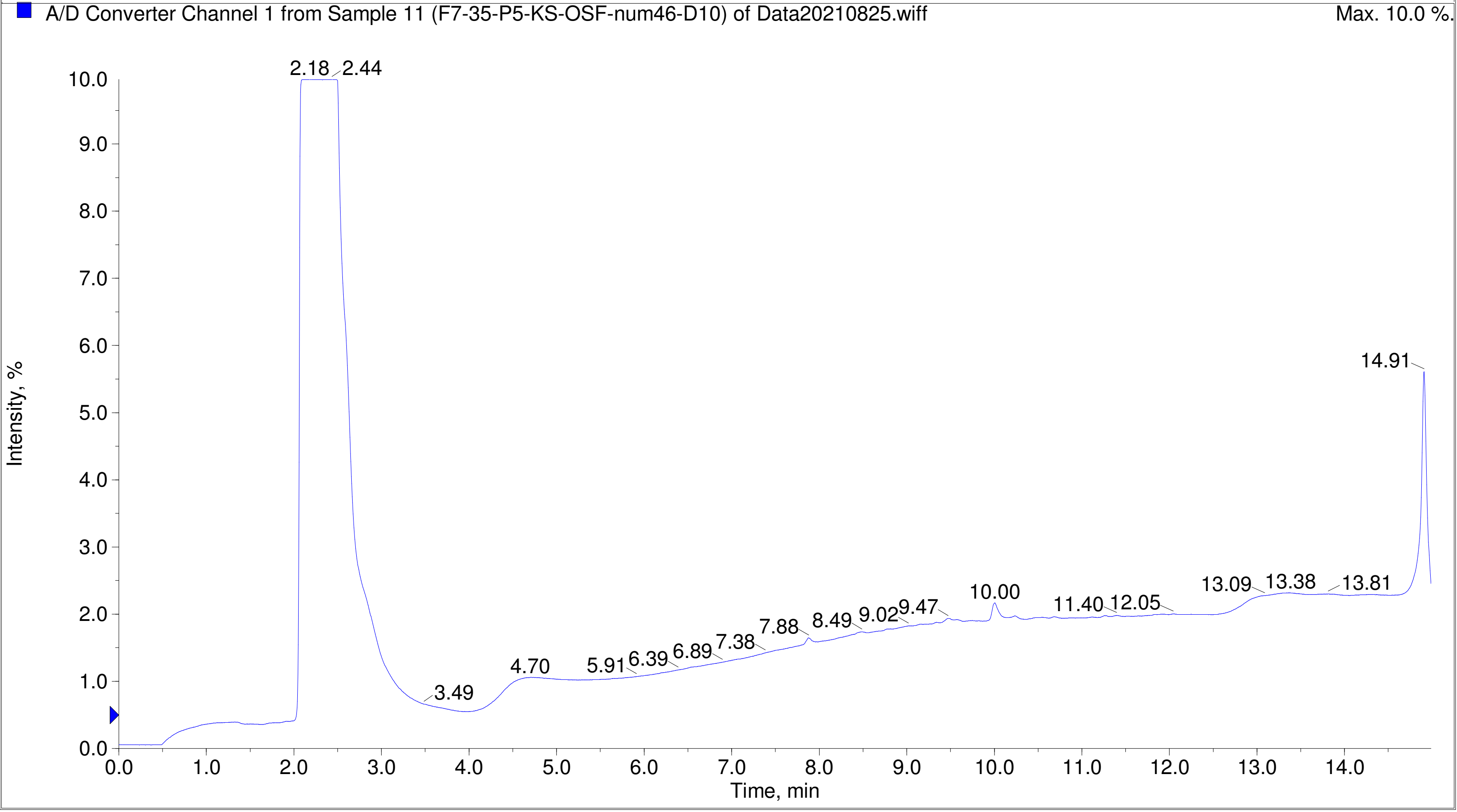
A/D Converter Channel 1 from Sample 11 (F7-35-P5-KS-OSF-num46-D10) of Data20210825.wiff
Max. 10.0 %.
Intensity, %
Time, min
2.18
2.44
14.91
10.00
13.09
13.38
13.81
11.40
12.05
7.88
8.49
9.02
9.47
7.38
6.89
6.39
5.91
4.70
3.49

Printing Time: 4:11:59 PM Post PM Polarity/Scan Type: Positive Q1 MS
Printing Date: Tuesday, January 31, 2023 Sample Name: F7-35-P5-KS-OSF-num49-E1
Service Engineer Chris Maier *Stanford U *API 3000 SN D7640012

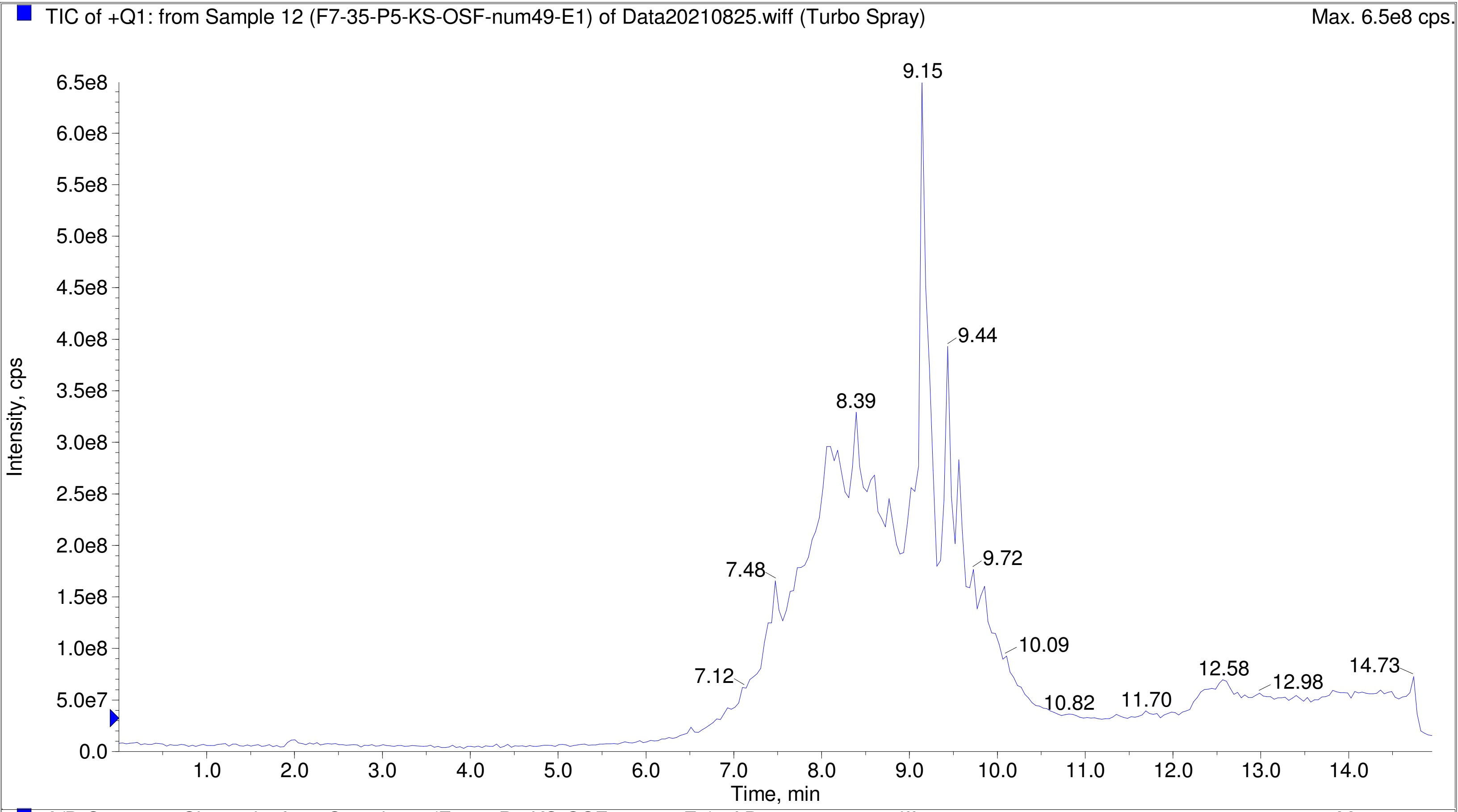


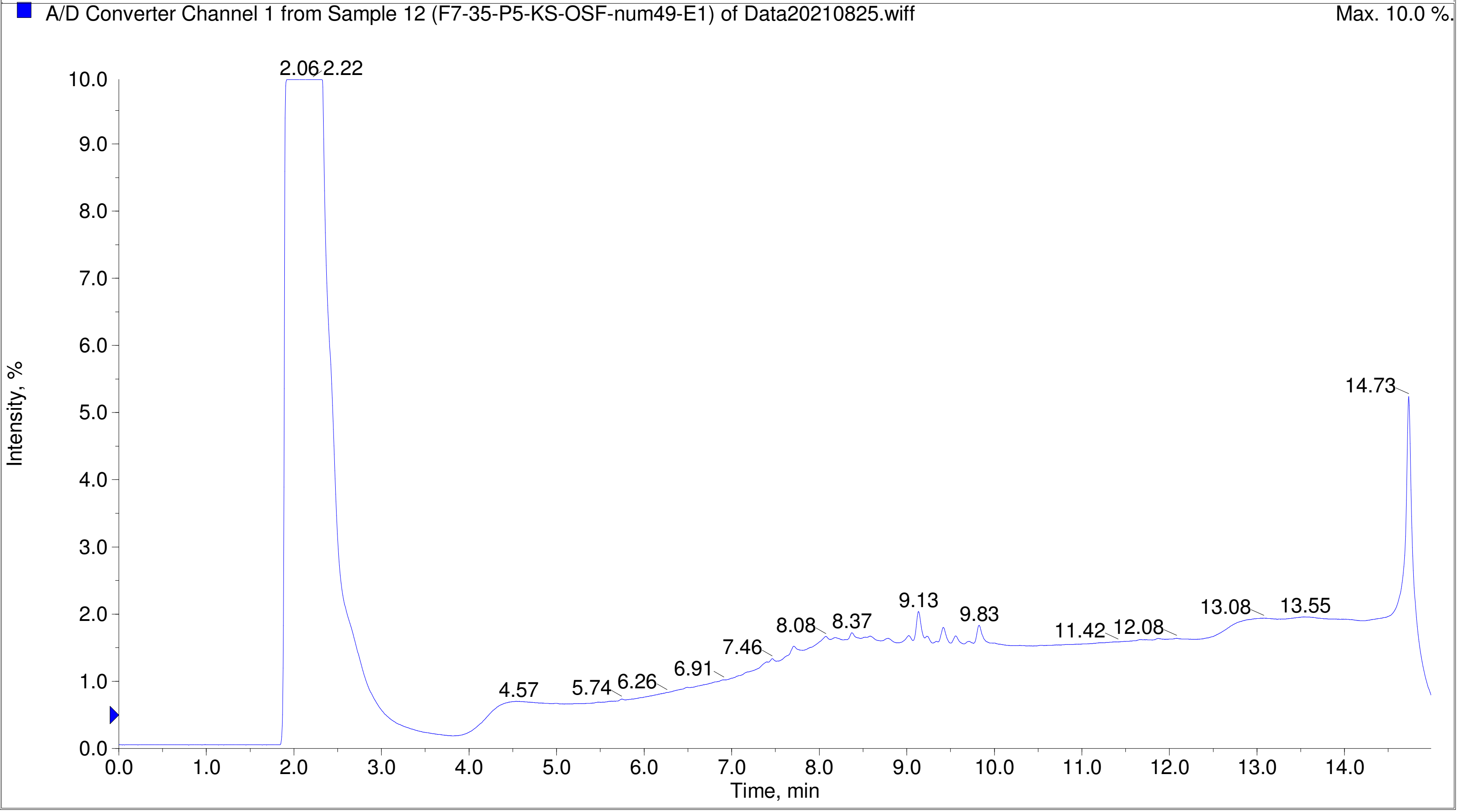

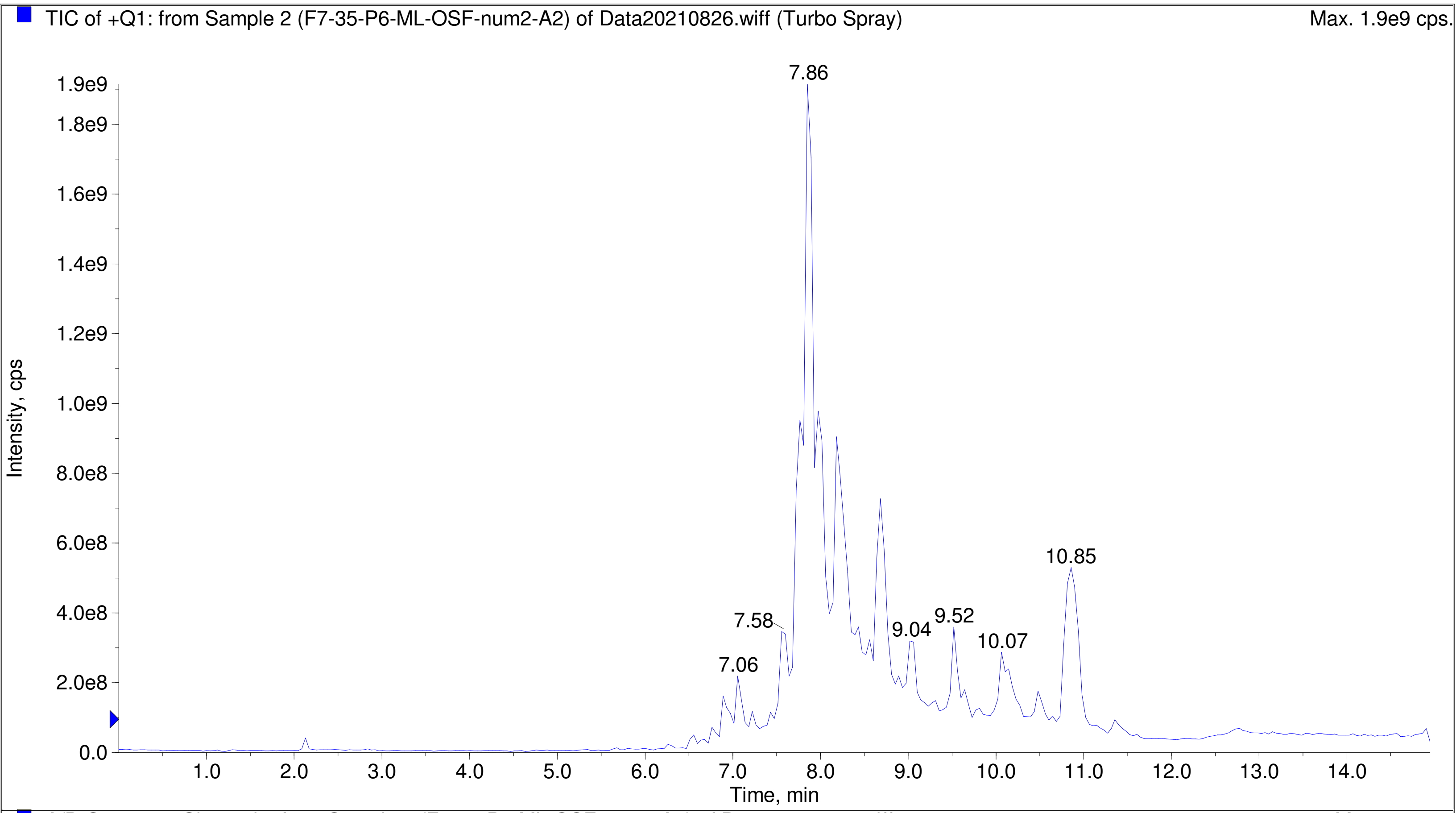

TIC of +Q1: from Sample 2 (F7-35-P6-ML-OSF-num2-A2) of Data20210826.wiff (Turbo Spray)
Max. 1.9e9 cps.
Intensity, cps
Time, min
7.86
7.06
7.58
9.04
9.52
10.07
10.85


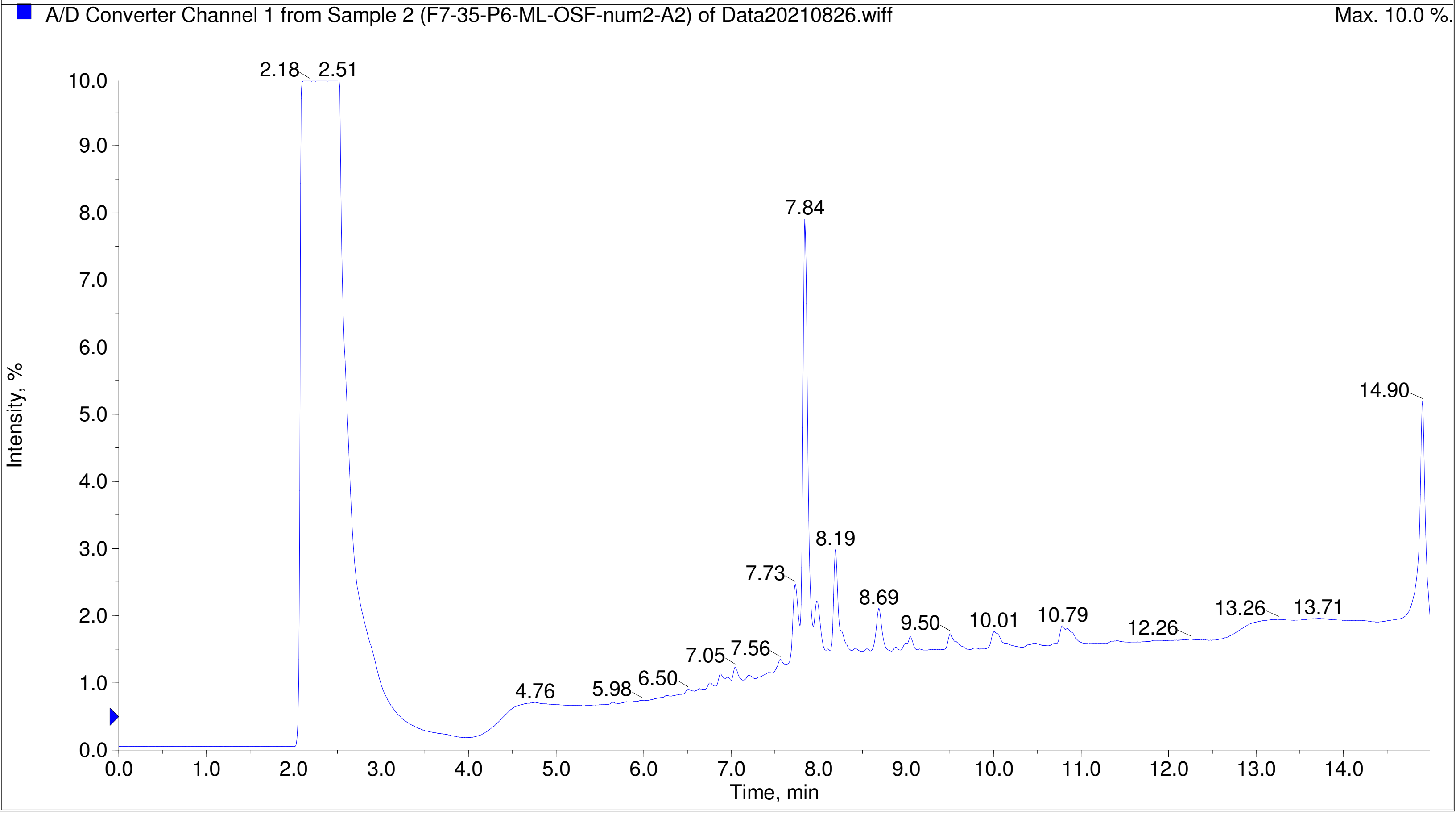

A/D Converter Channel 1 from Sample 2 (F7-35-P6-ML-OSF-num2-A2) of Data20210826.wiff
Max. 10.0 %.
Intensity, %
Time, min
2.18
2.51
4.76
5.98
6.50
7.05
7.56
7.73
7.84
8.19
8.69
9.50
10.01
10.79
12.26
13.26
13.71
14.90

Printing Time: 3:12:25 PM
Printing Date: Monday, January 30, 2023
Service Engineer Chris Maier

Post PM
*Stanford U

Polarity/Scan Type: Positive Q1 MS
Sample Name: F7-35-P6-ML-OSF-num7-A7
*API 3000 SN D7640012

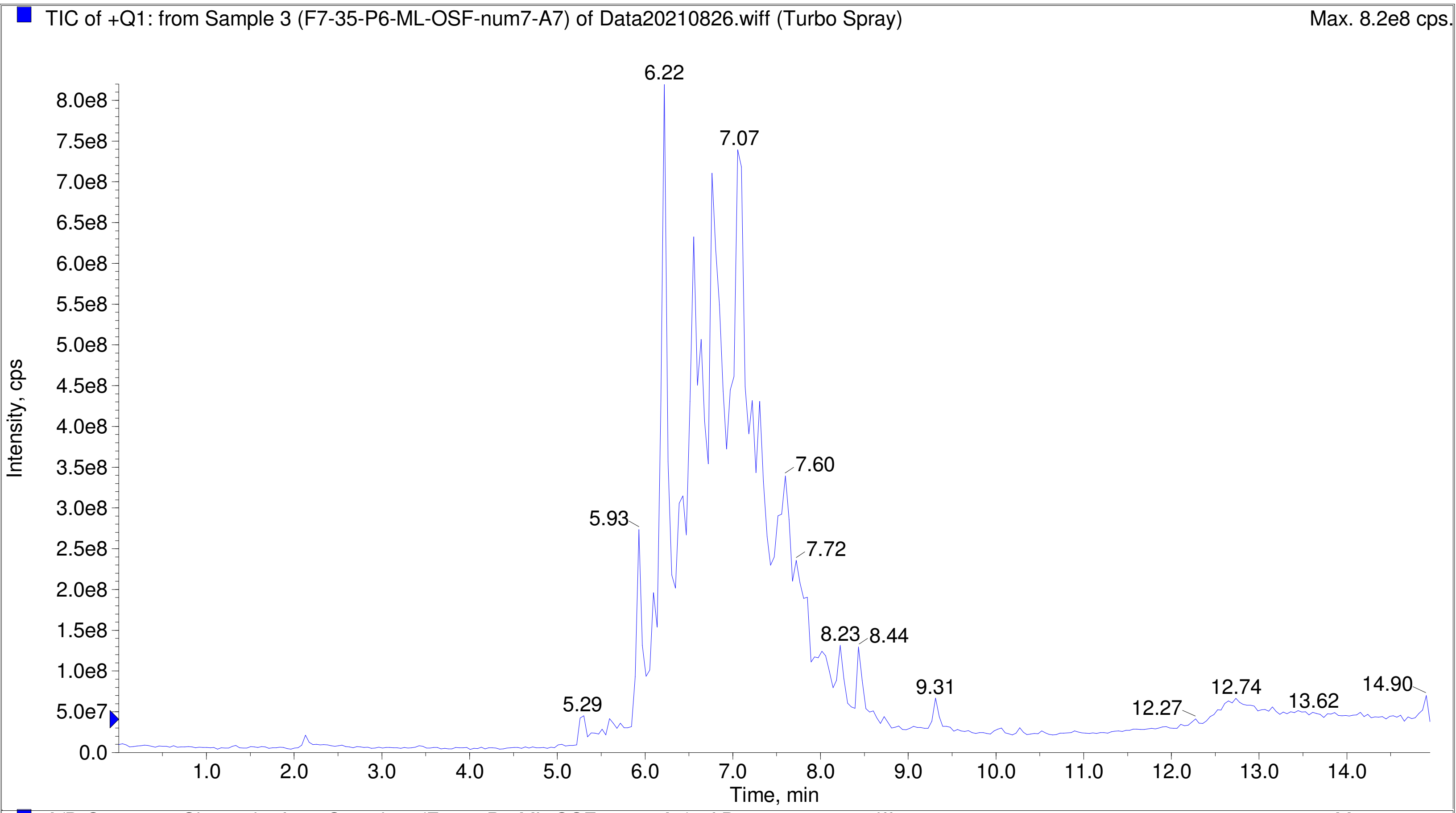


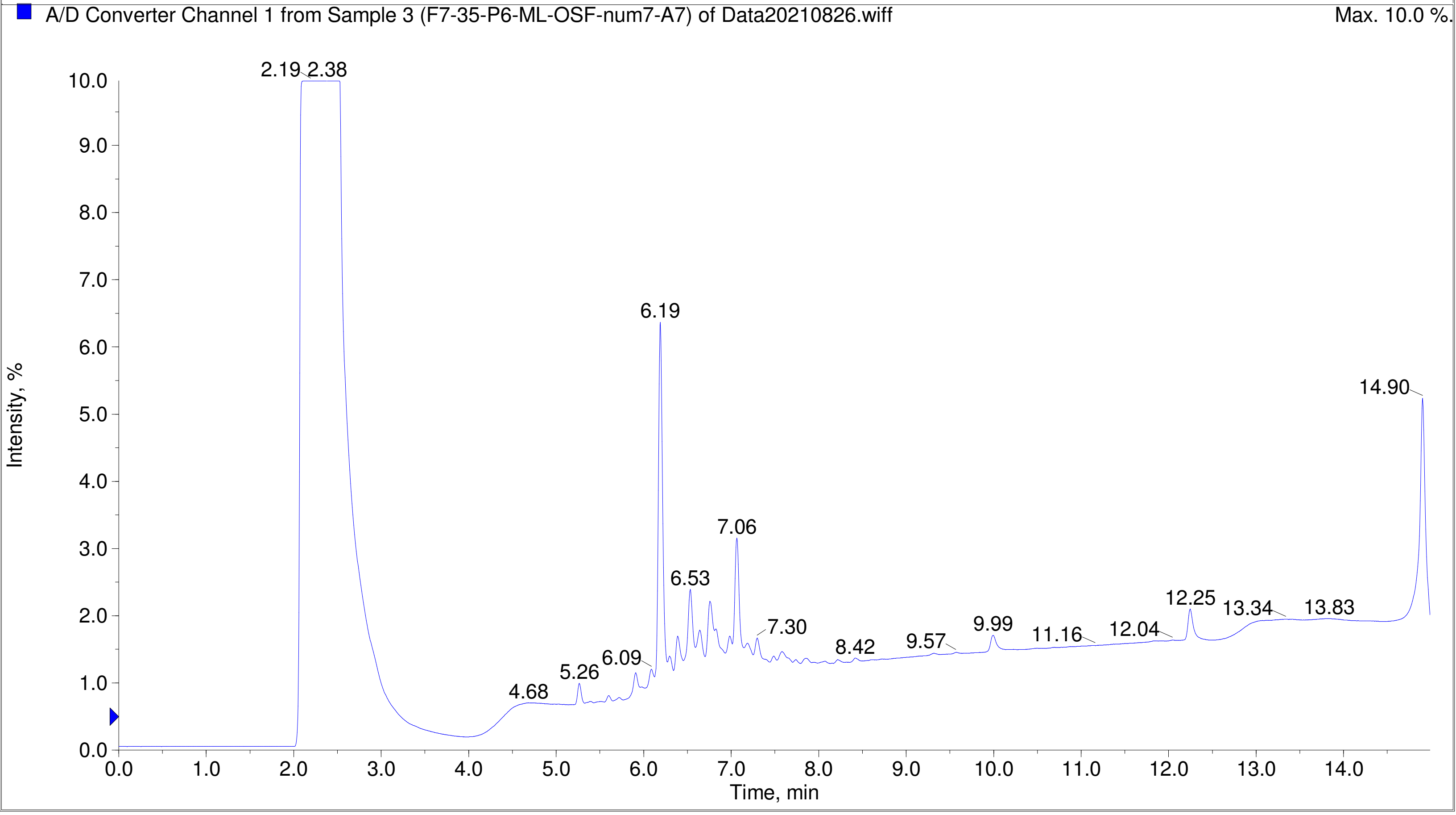

TIC of +Q1: from Sample 4 (F7-35-P6-ML-OSF-num14-B2) of Data20210826.wiff (Turbo Spray) Max. 1.6e9 cps.

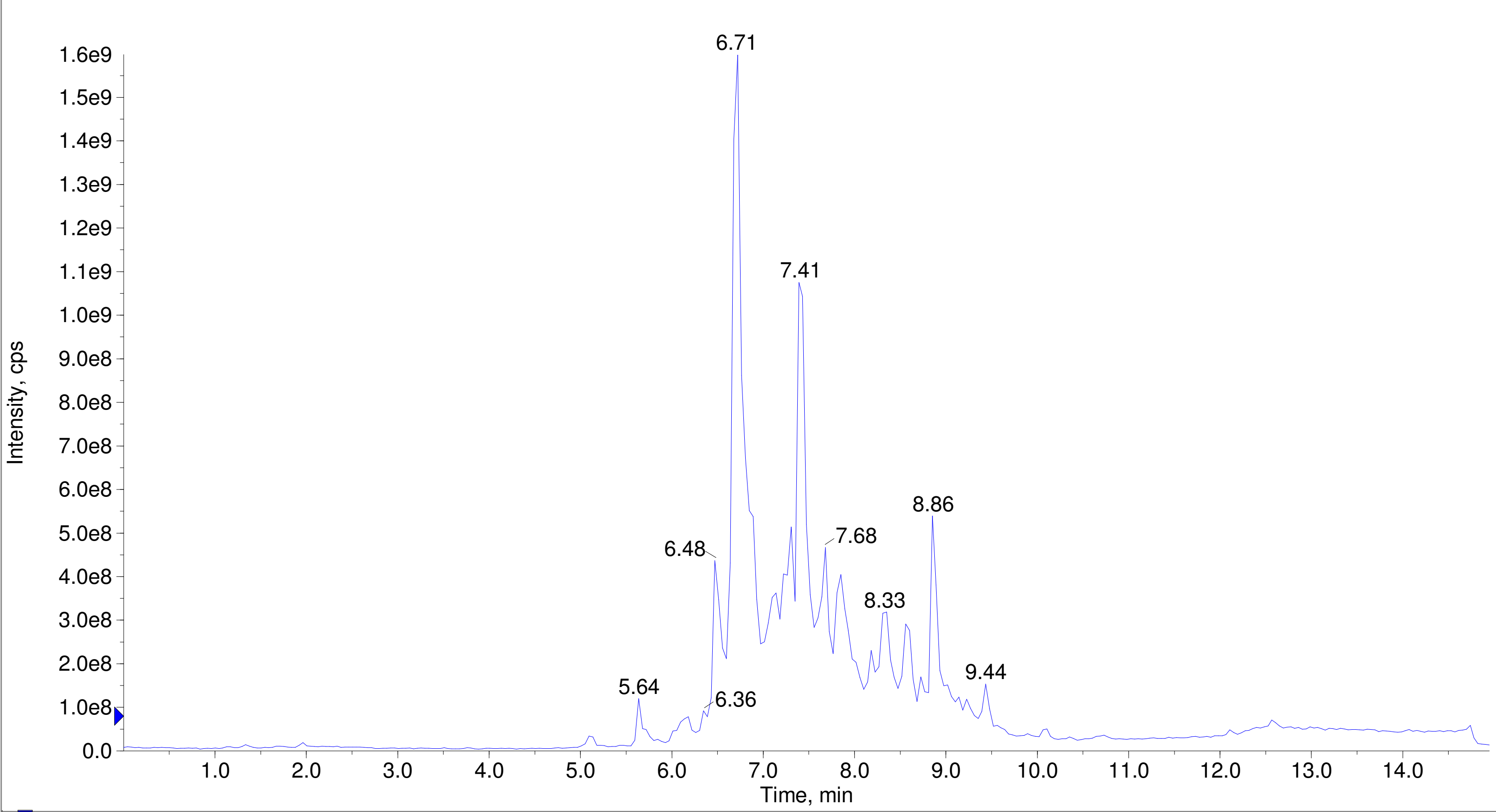


A/D Converter Channel 1 from Sample 4 (F7-35-P6-ML-OSF-num14-B2) of Data20210826.wiff Max. 10.0 %.

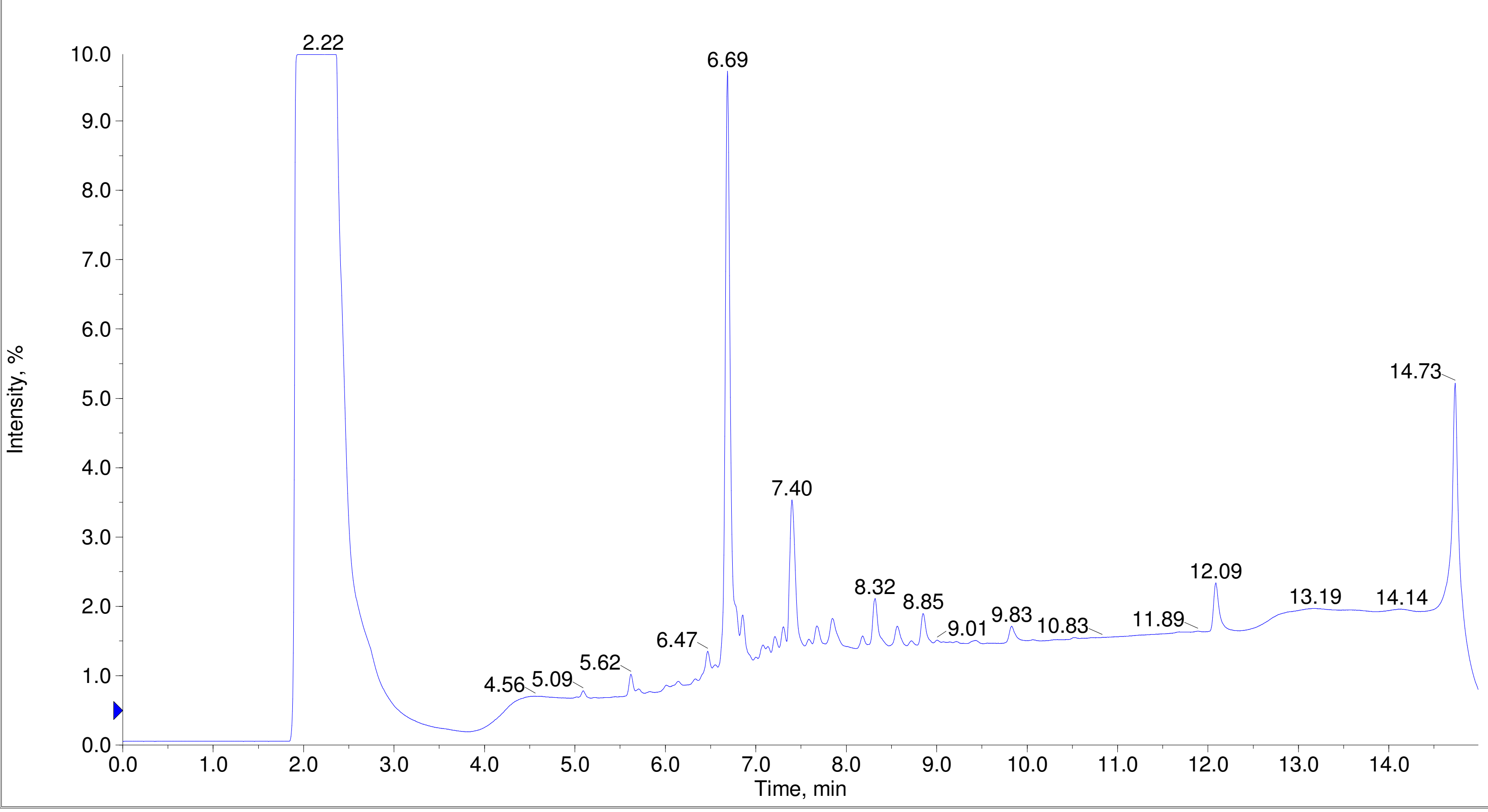

Printing Time: 3:15:26 PM
Printing Date: Monday, January 30, 2023
Service Engineer Chris Maier

Post PM
*Stanford U

Polarity/Scan Type: Positive Q1 MS
Sample Name: F7-35-P6-ML-OSF-num18-B6
*API 3000 SN D7640012

TIC of +Q1: from Sample 5 (F7-35-P6-ML-OSF-num18-B6) of Data20210826.wiff (Turbo Spray) Max. 2.1e9 cps.

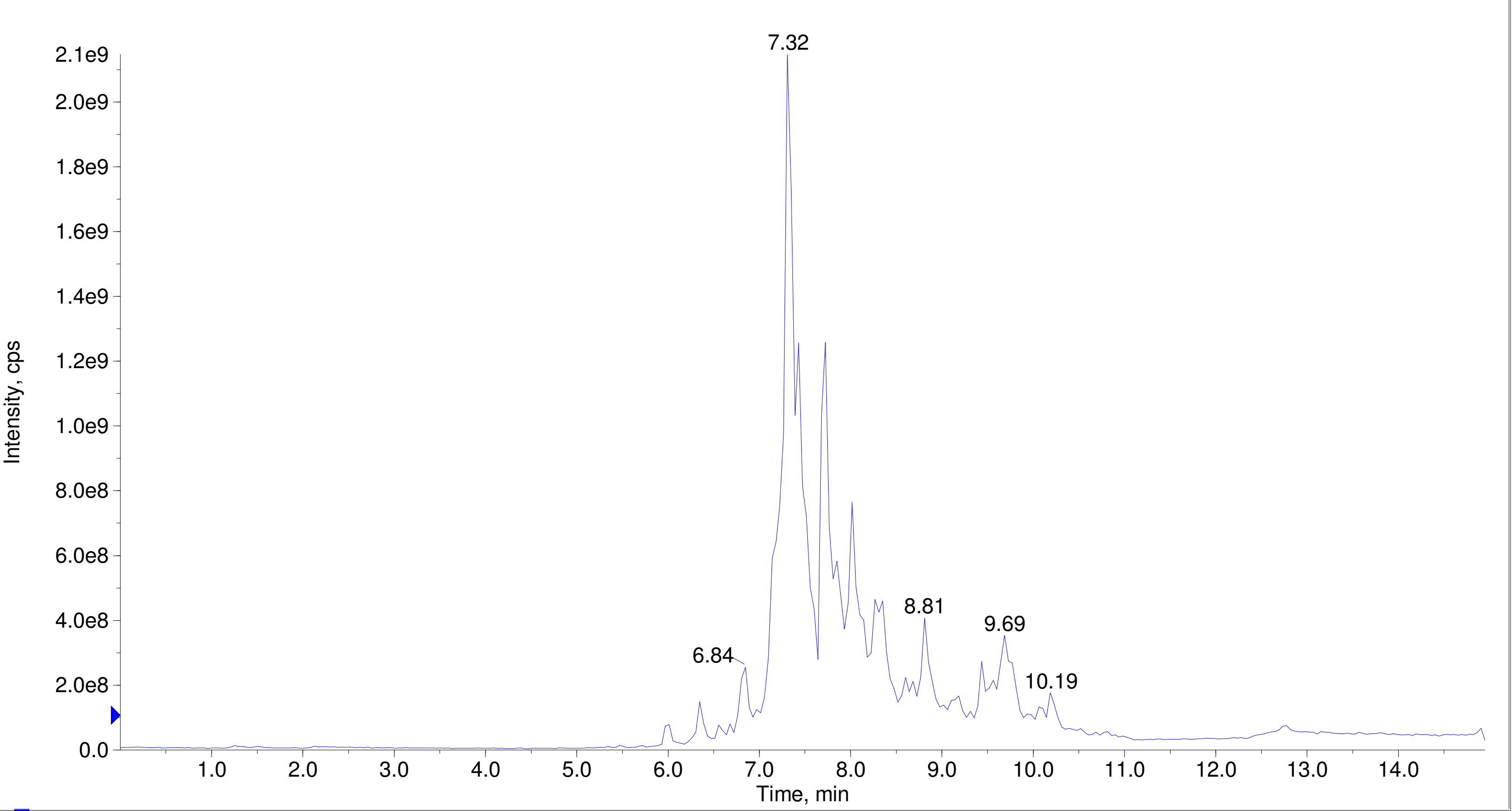


A/D Converter Channel 1 from Sample 5 (F7-35-P6-ML-OSF-num18-B6) of Data20210826.wiff Max. 10.0 %.

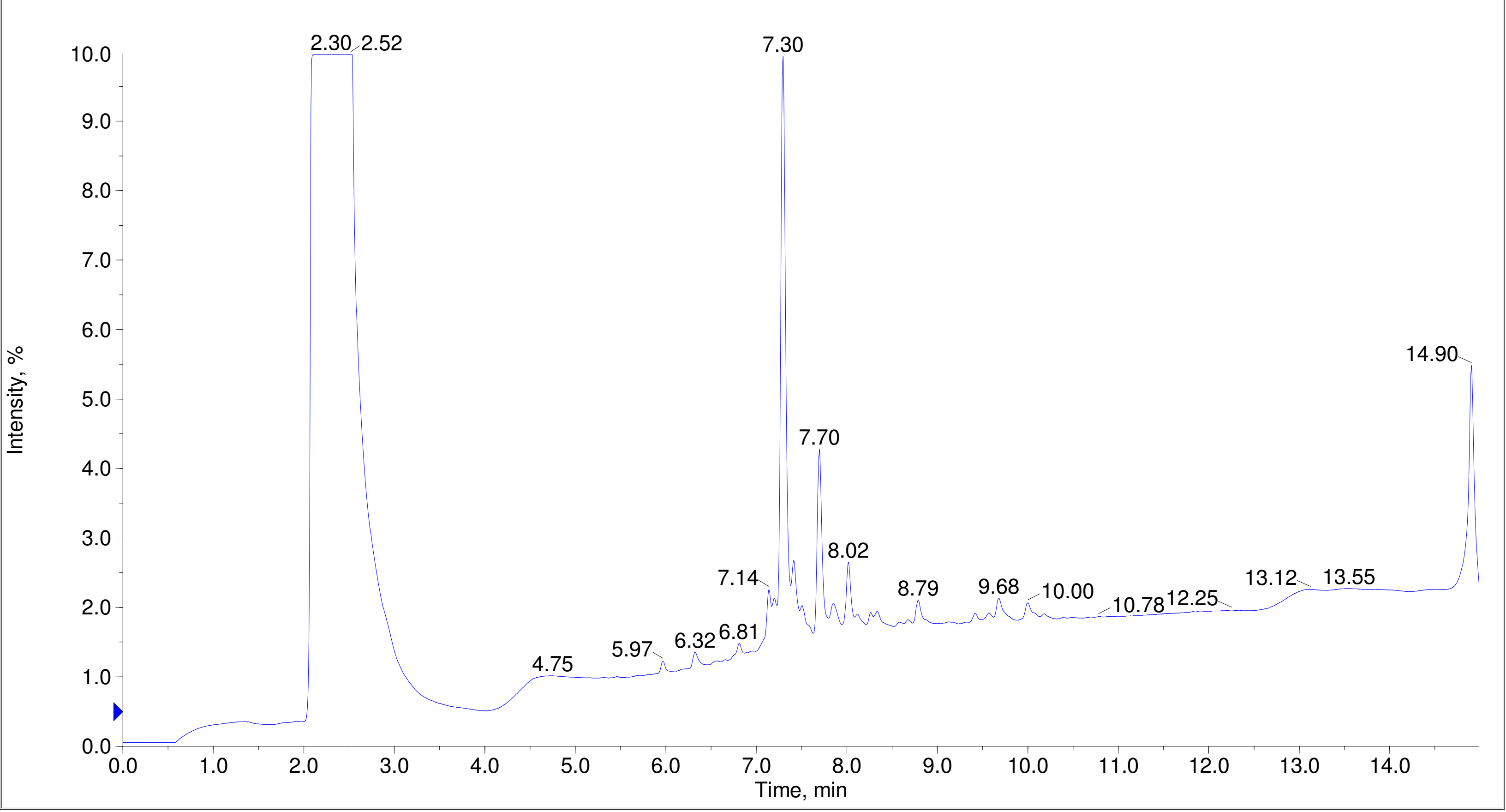

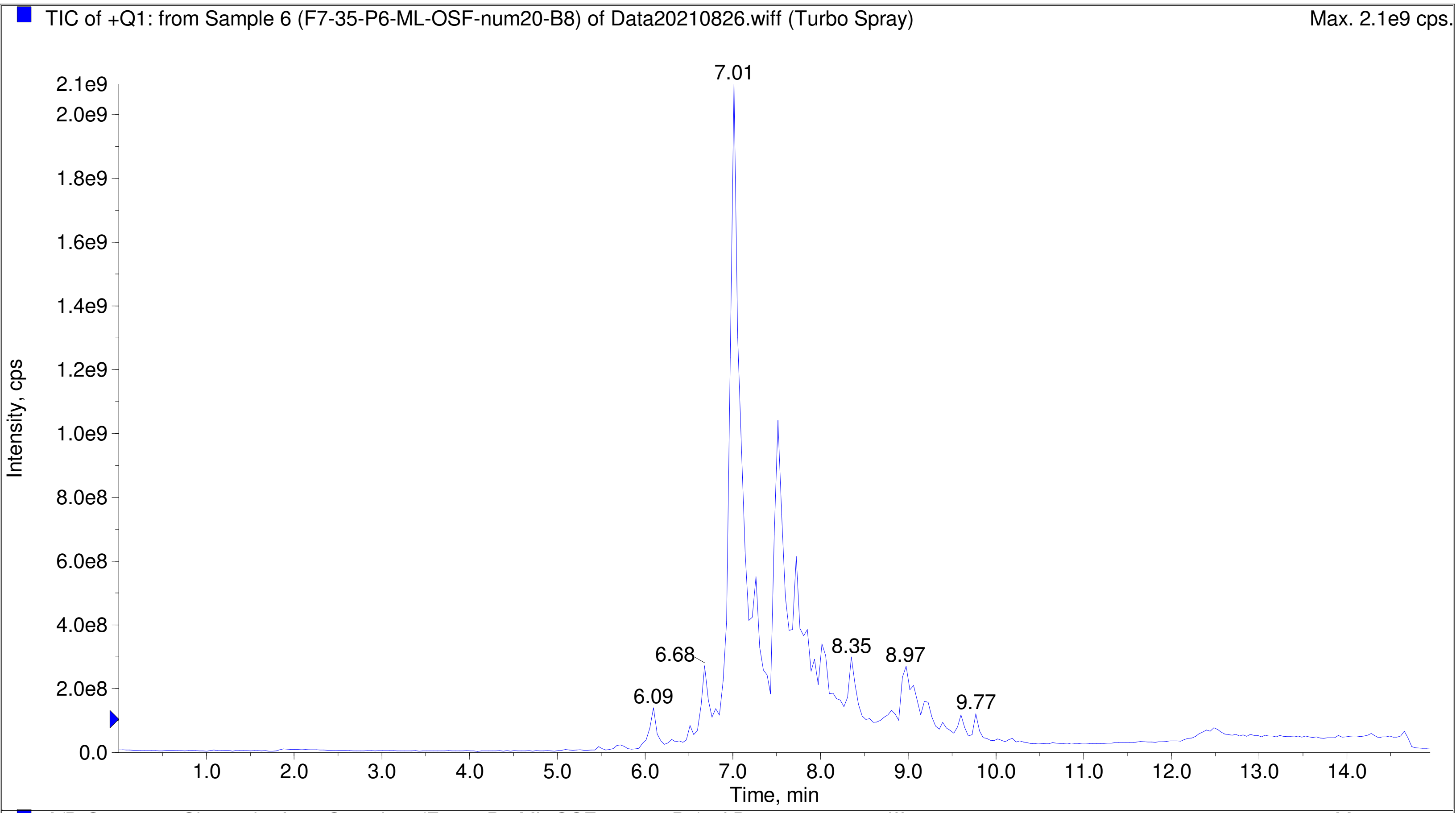
TIC of +Q1: from Sample 6 (F7-35-P6-ML-OSF-num20-B8) of Data20210826.wiff (Turbo Spray)
Max. 2.1e9 cps.
Intensity, cps
Time, min
7.01
6.09
6.68
8.35
8.97
9.77

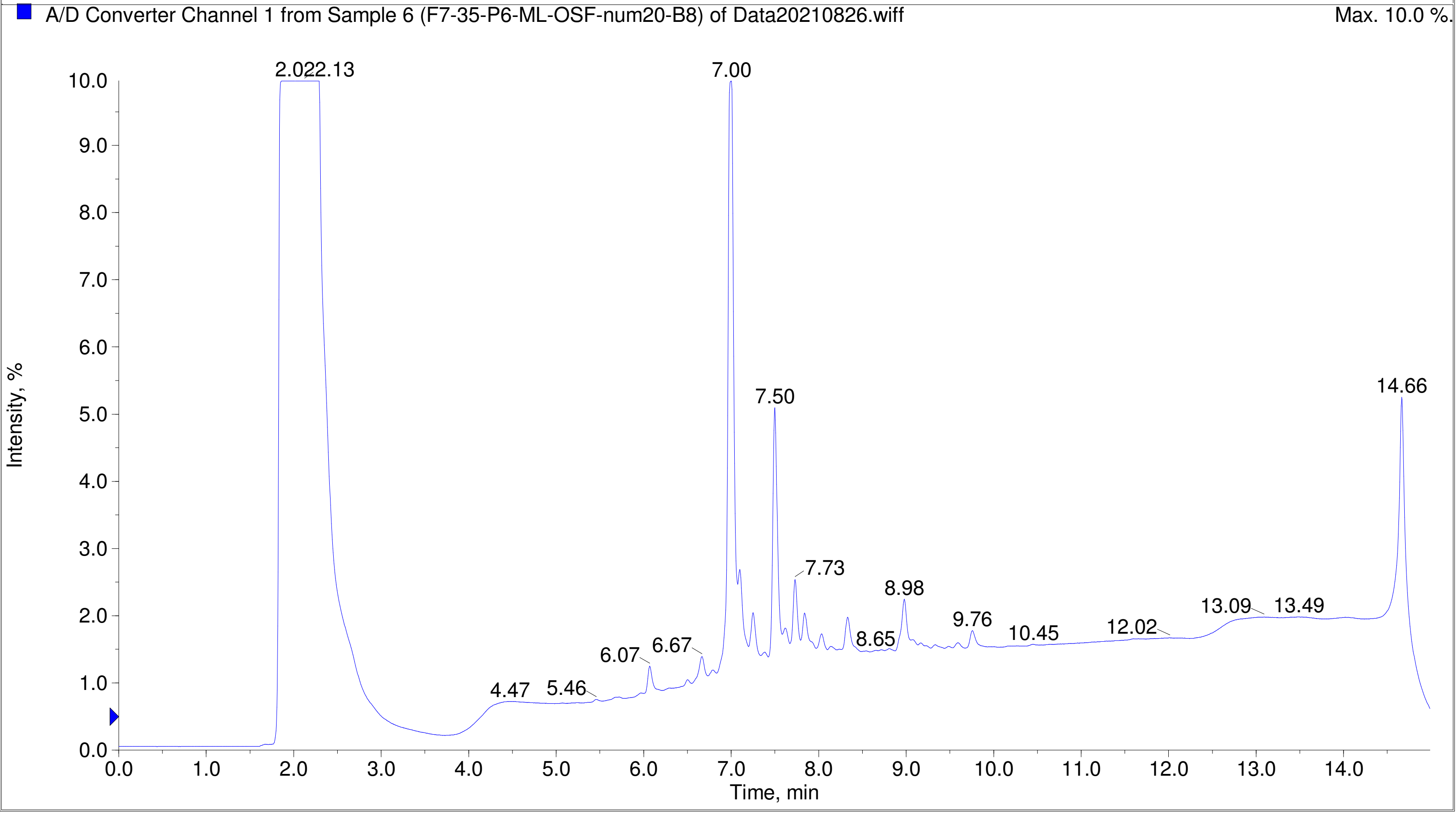
A/D Converter Channel 1 from Sample 6 (F7-35-P6-ML-OSF-num20-B8) of Data20210826.wiff
Max. 10.0 %.
Intensity, %
Time, min
2.022.13
7.00
7.50
14.66
7.73
8.98
9.76
10.45
12.02
13.09
13.49
8.65
6.07
6.67
4.47
5.46

Printing Time: 4:14:52 PM
Printing Date: Tuesday, January 31, 2023
Service Engineer Chris Maier

Post PM
*Stanford U

Polarity/Scan Type: Positive Q1 MS
Sample Name: F7-35-P6-ML-OSF-num21-B9
*API 3000 SN D7640012

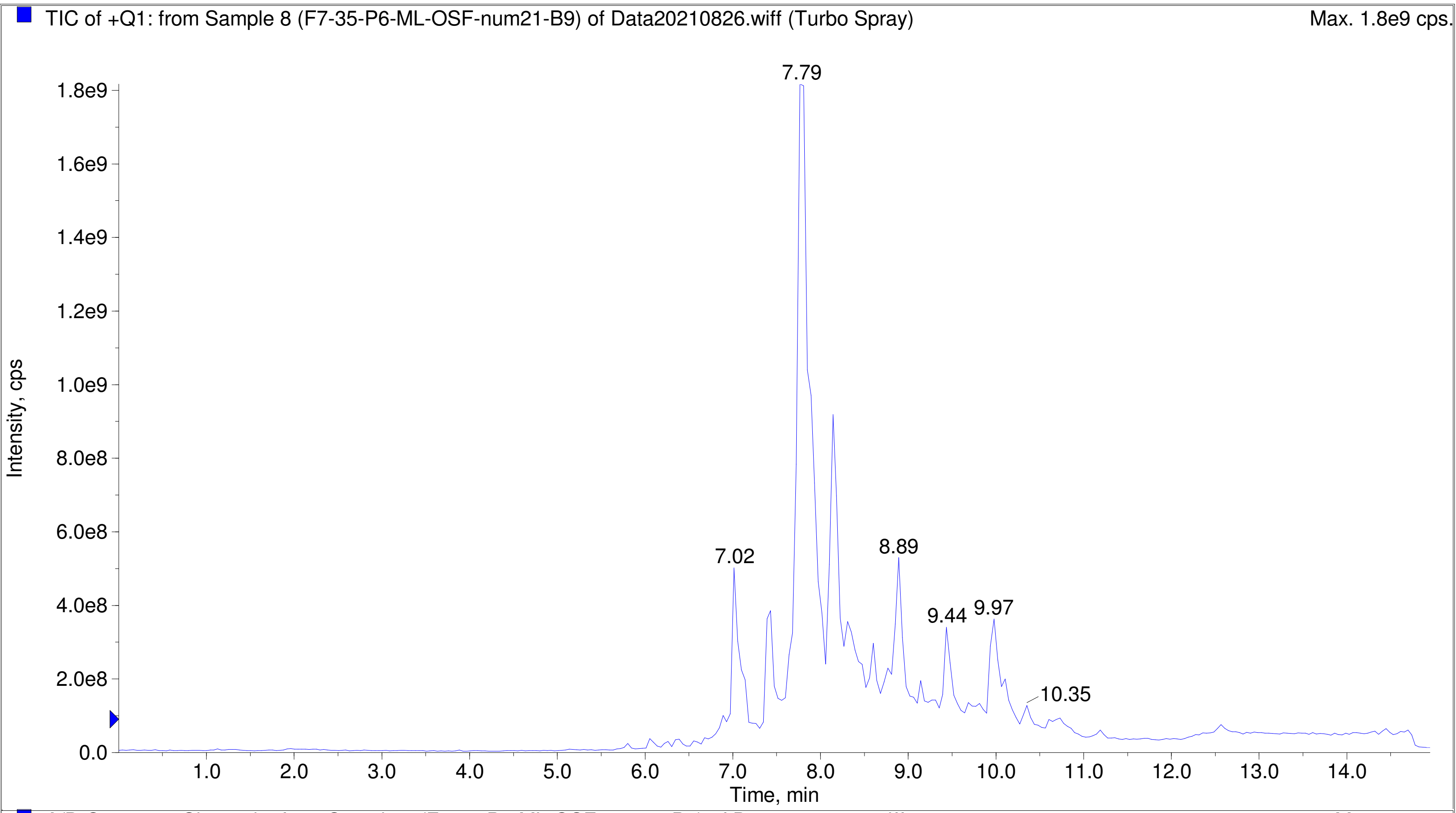


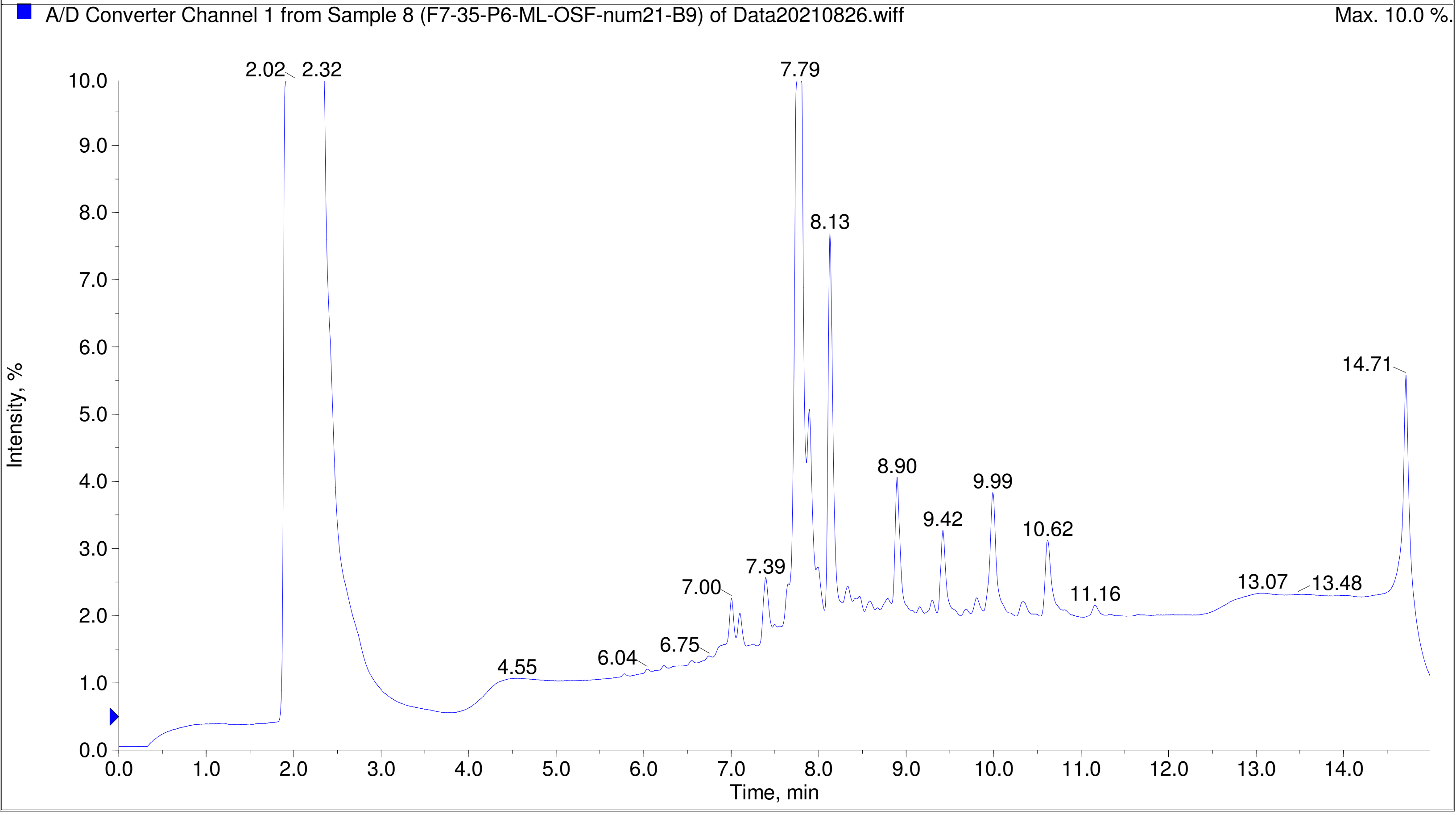

Printing Time: 4:17:00 PM Post PM Polarity/Scan Type: Positive Q1 MS
Printing Date: Tuesday, January 31, 2023 Sample Name: F7-35-P6-ML-OSF-num34-C10
Service Engineer Chris Maier *Stanford U *API 3000 SN D7640012

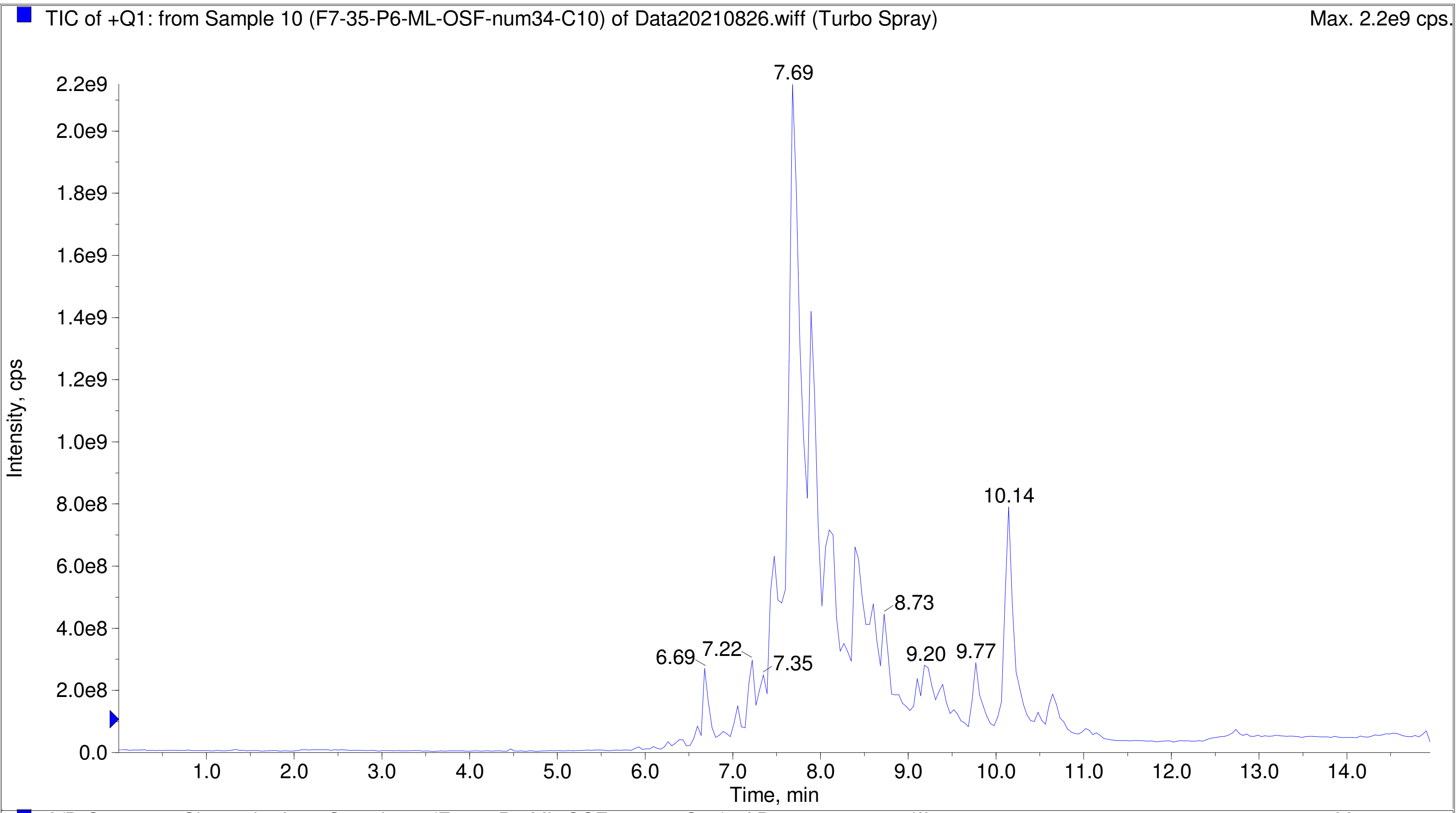


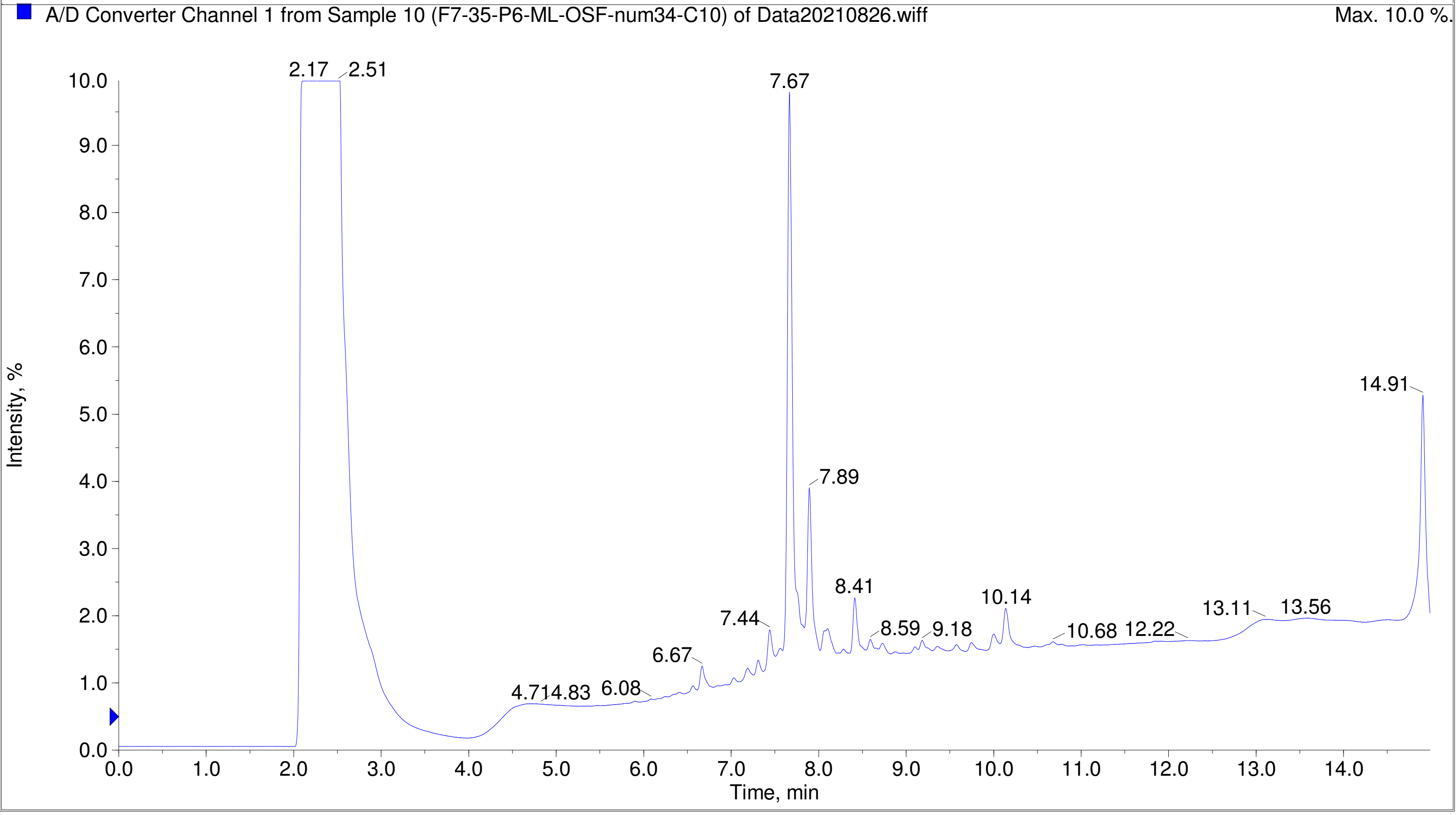

Printing Time: 4:17:56 PM
Printing Date: Tuesday, January 31, 2023
Service Engineer Chris Maier

Post PM

*Stanford U

Polarity/Scan Type: Positive Q1 MS
Sample Name: F7-35-P6-ML-OSF-num36-C12
*API 3000 SN D7640012

TIC of +Q1: from Sample 11 (F7-35-P6-ML-OSF-num36-C12) of Data20210826.wiff (Turbo Spray) Max. 2.1e9 cps.

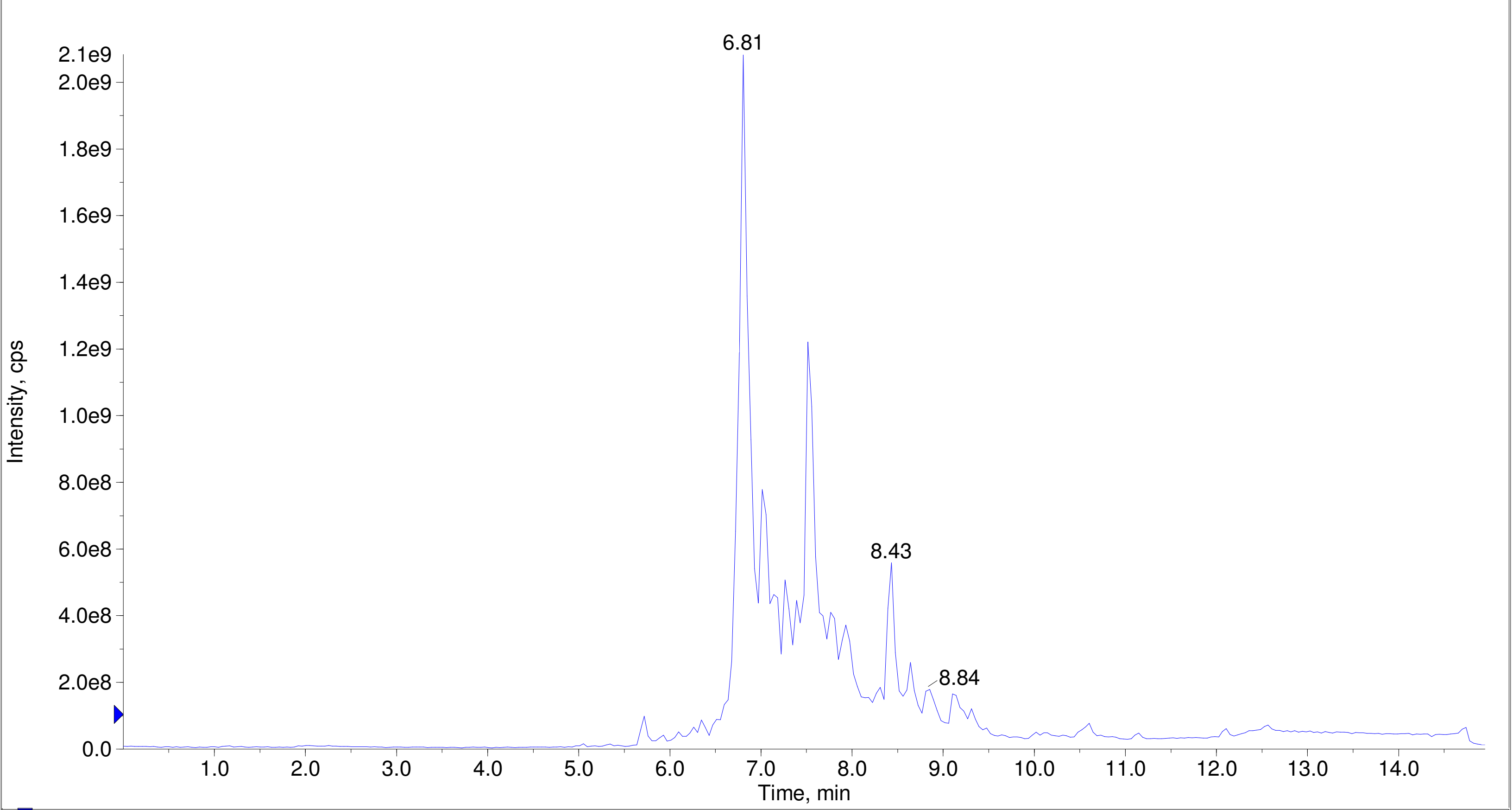


A/D Converter Channel 1 from Sample 11 (F7-35-P6-ML-OSF-num36-C12) of Data20210826.wiff Max. 10.0 %.

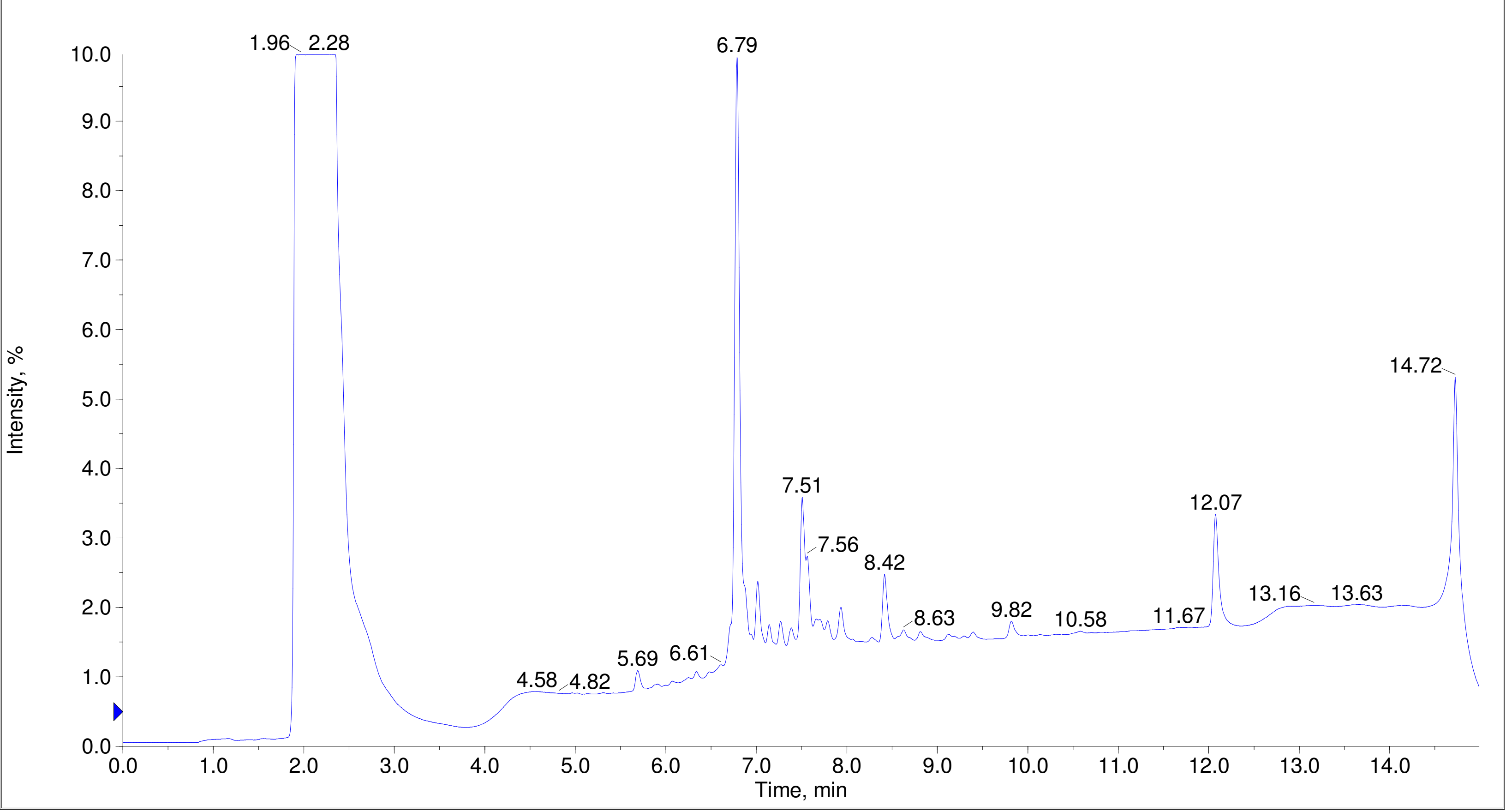

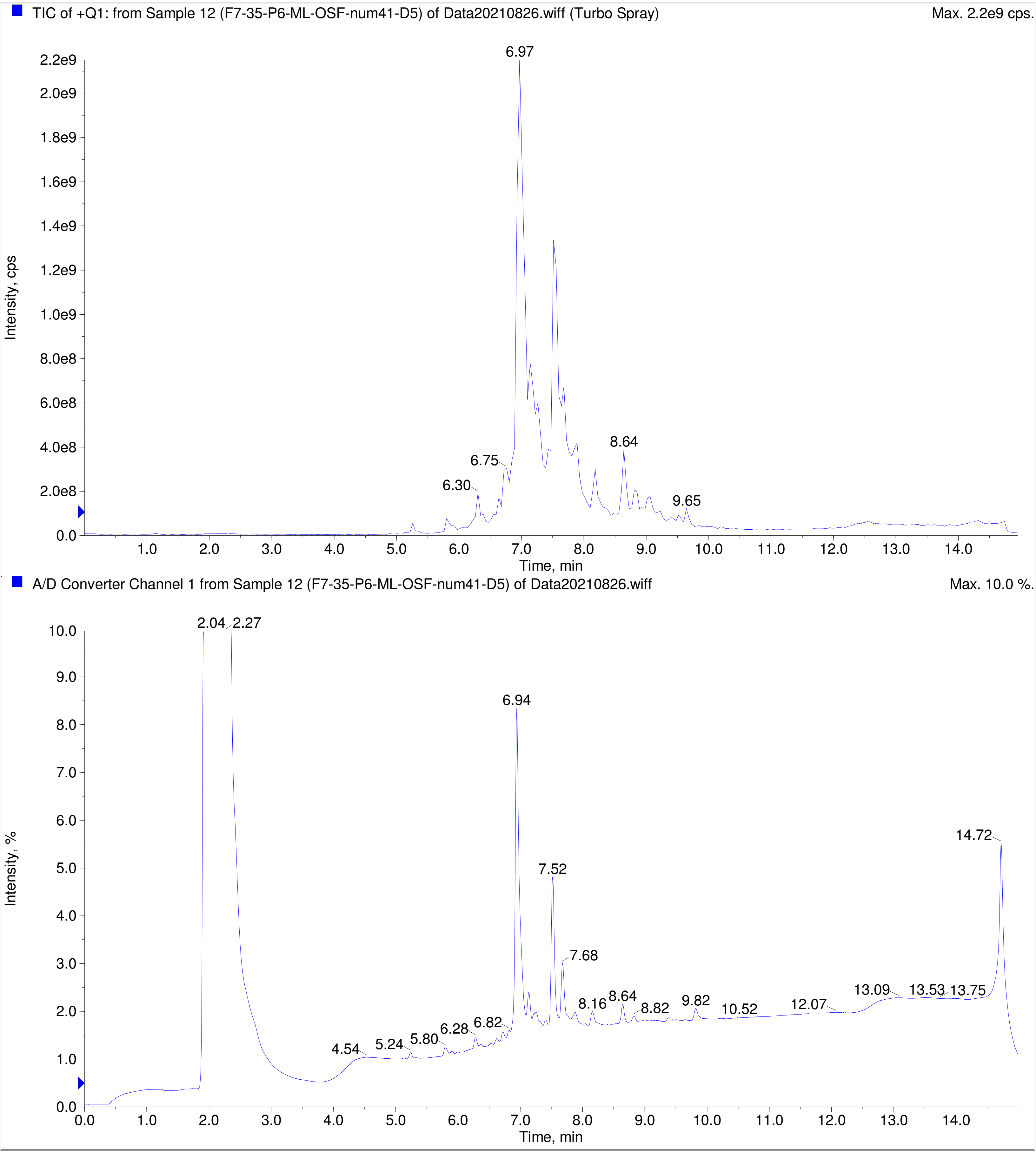
TIC of +Q1: from Sample 12 (F7-35-P6-ML-OSF-num41-D5) of Data20210826.wiff (Turbo Spray)
Max. 2.2e9 cps.
Intensity, cps
Time, min
6.97
6.75
6.30
8.64
9.65
A/D Converter Channel 1 from Sample 12 (F7-35-P6-ML-OSF-num41-D5) of Data20210826.wiff
Max. 10.0 %.
Intensity, %
Time, min
2.04
2.27
6.94
7.52
7.68
14.72
4.54
5.24
5.80
6.28
6.82
8.16
8.64
8.82
9.82
10.52
12.07
13.09
13.53
13.75

TIC of +Q1: from Sample 2 (F7-35-P7-IL_OSF-num2-A2) of Data20210827.wiff (Turbo Spray) Max. 7.1e8 cps.

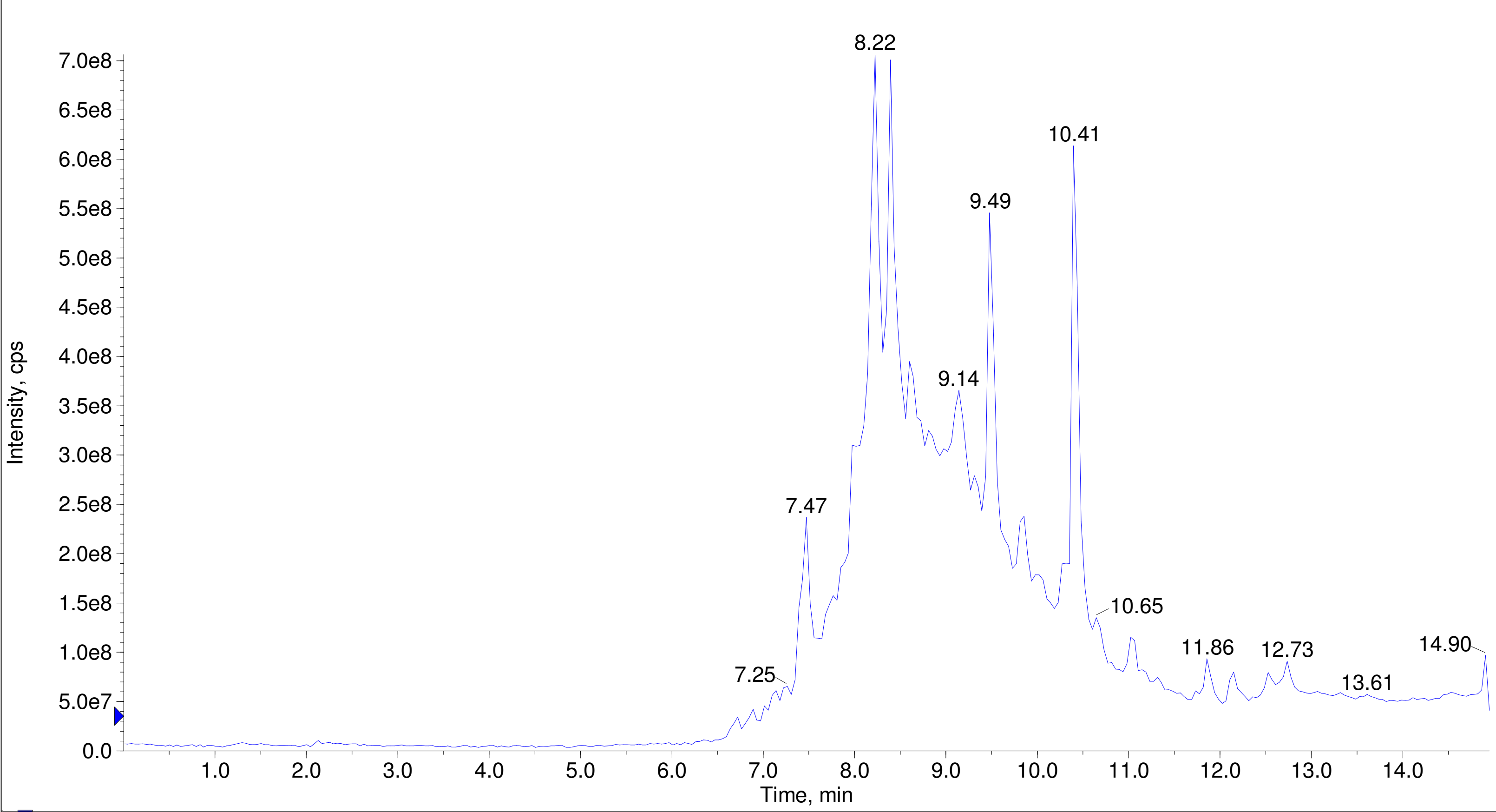


A/D Converter Channel 1 from Sample 2 (F7-35-P7-IL_OSF-num2-A2) of Data20210827.wiff Max. 10.0 %.

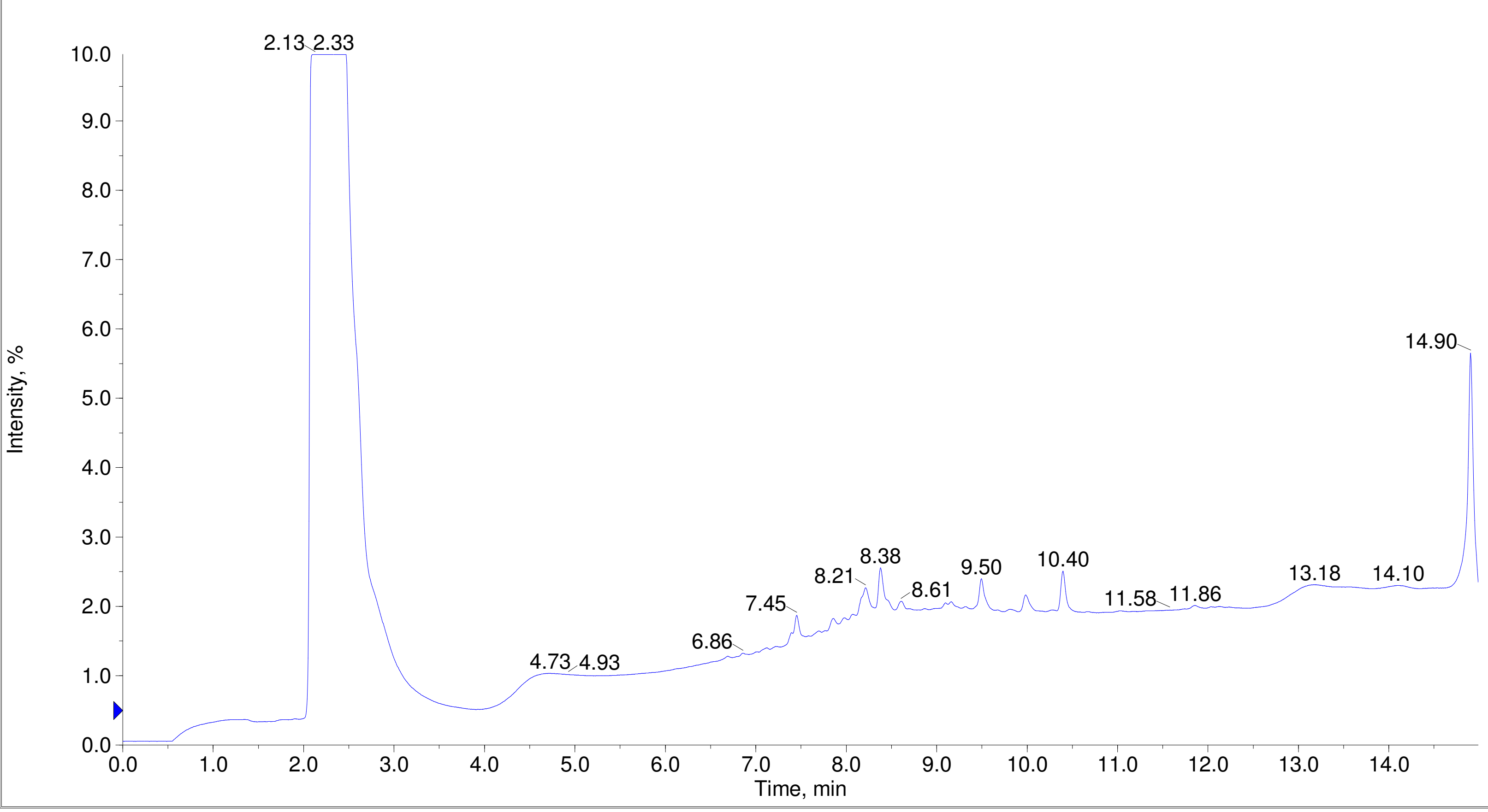

Printing Time: 3:22:39 PM    Post PM    Polarity/Scan Type: Positive Q1 MS
Printing Date: Monday, January 30, 2023    Sample Name: F7-35-P7-IL_OSF-num6-A6
Service Engineer Chris Maier    *Stanford U    *API 3000 SN D7640012

TIC of +Q1: from Sample 3 (F7-35-P7-IL_OSF-num6-A6) of Data20210827.wiff (Turbo Spray)    Max. 3.0e8 cps.

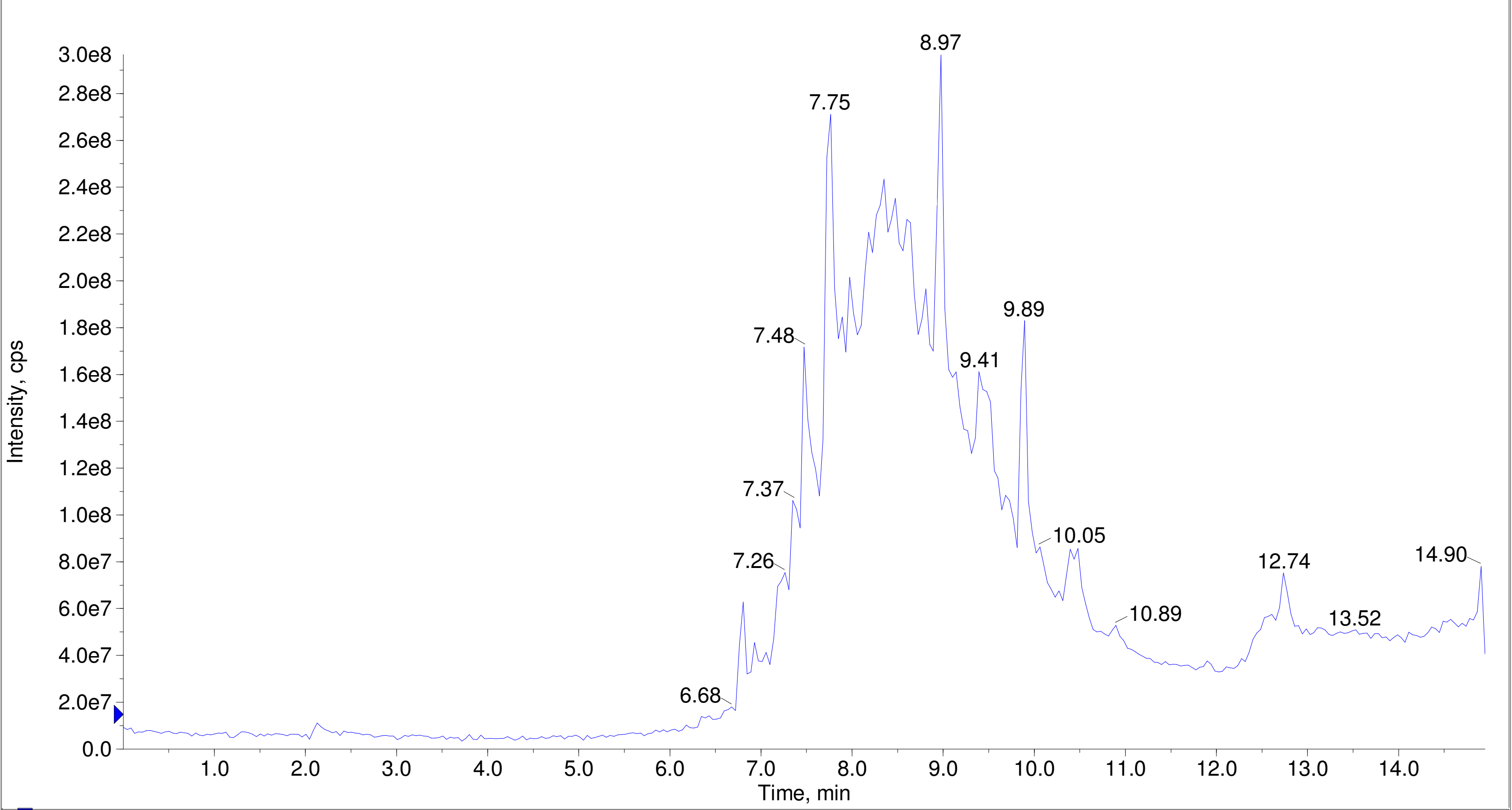


A/D Converter Channel 1 from Sample 3 (F7-35-P7-IL_OSF-num6-A6) of Data20210827.wiff    Max. 10.0 %.

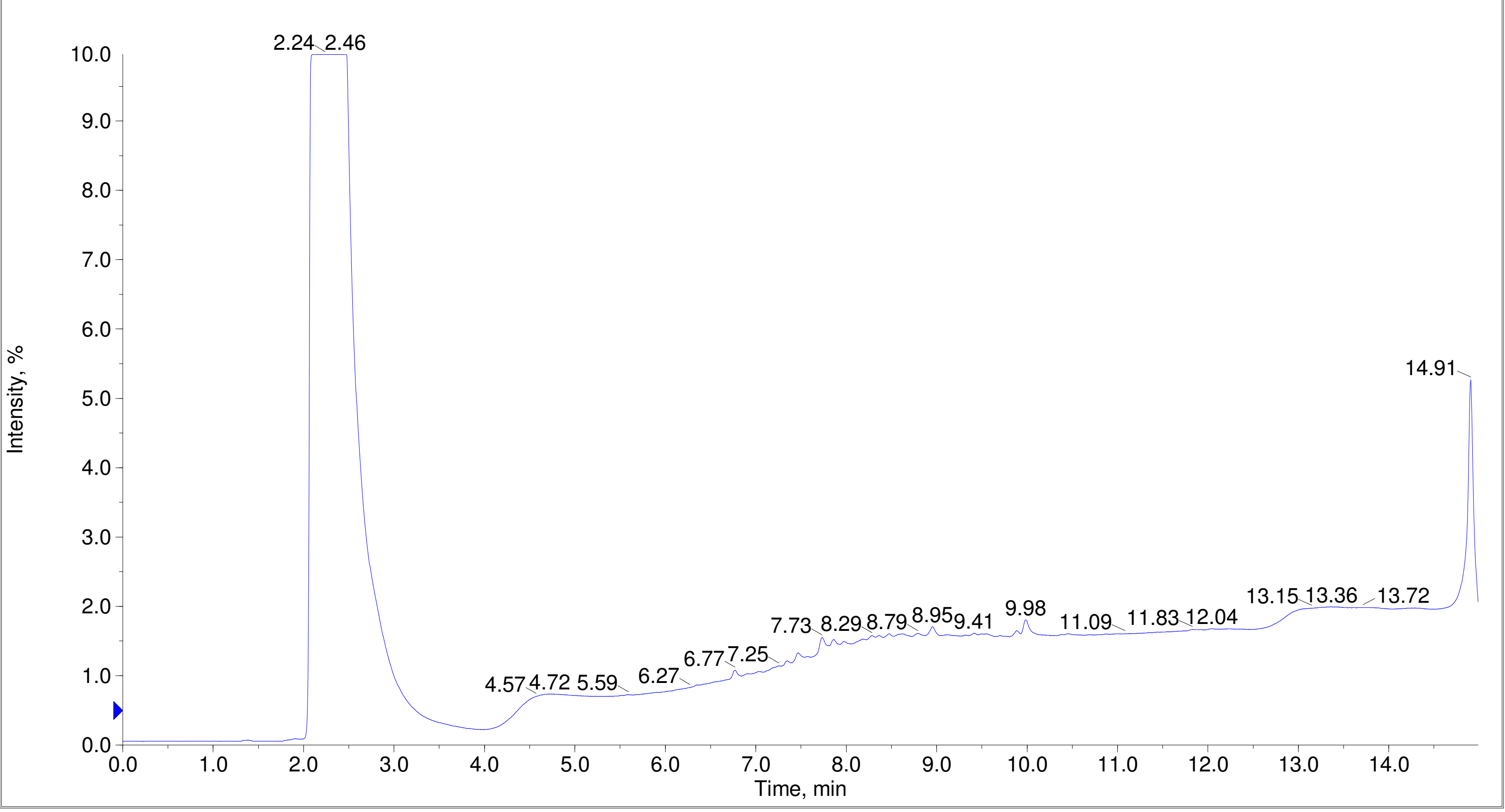

Printing Time: 3:23:48 PM    Post PM    Polarity/Scan Type: Positive Q1 MS
Printing Date: Monday, January 30, 2023    Sample Name: F7-35-P7-IL_OSF-num12-A12
Service Engineer Chris Maier    *Stanford U    *API 3000 SN D7640012

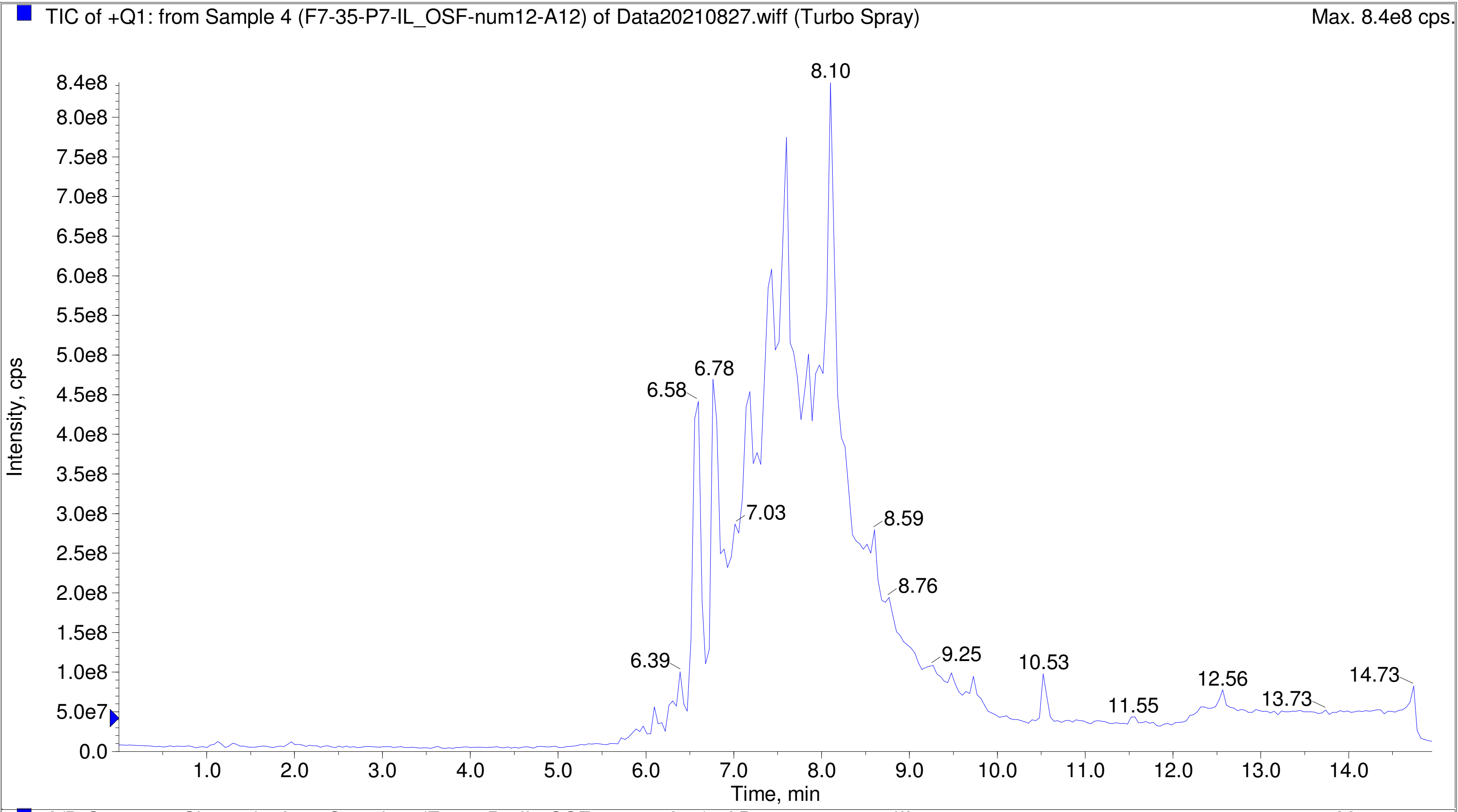


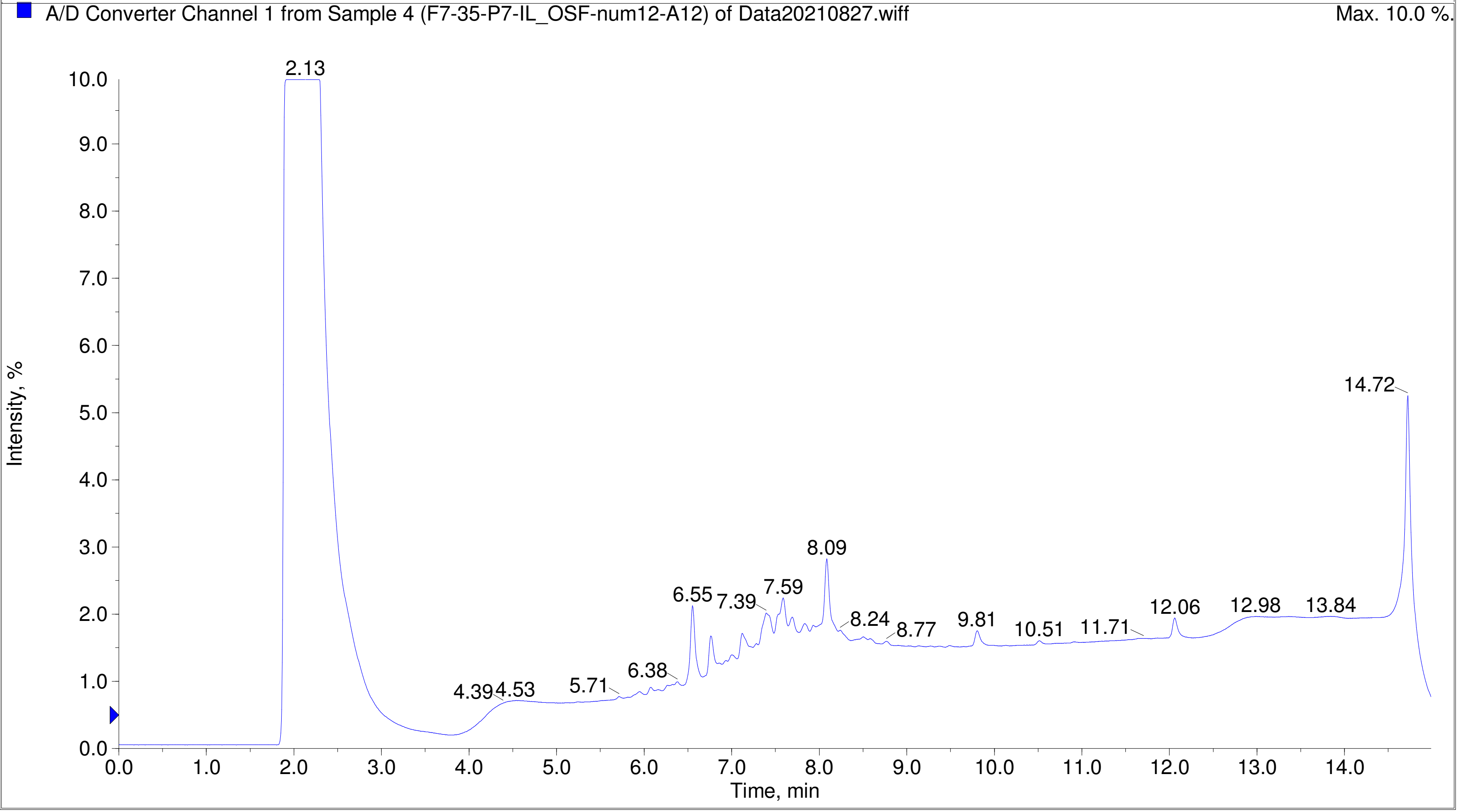

Printing Time: 3:22:09 PM Post PM Polarity/Scan Type: Positive Q1 MS
Printing Date: Monday, January 30, 2023 Sample Name: F7-35-P7-IL_OSF-num20-B8
Service Engineer Chris Maier *Stanford U *API 3000 SN D7640012

TIC of +Q1: from Sample 5 (F7-35-P7-IL_OSF-num20-B8) of Data20210827.wiff (Turbo Spray) Max. 6.2e8 cps.

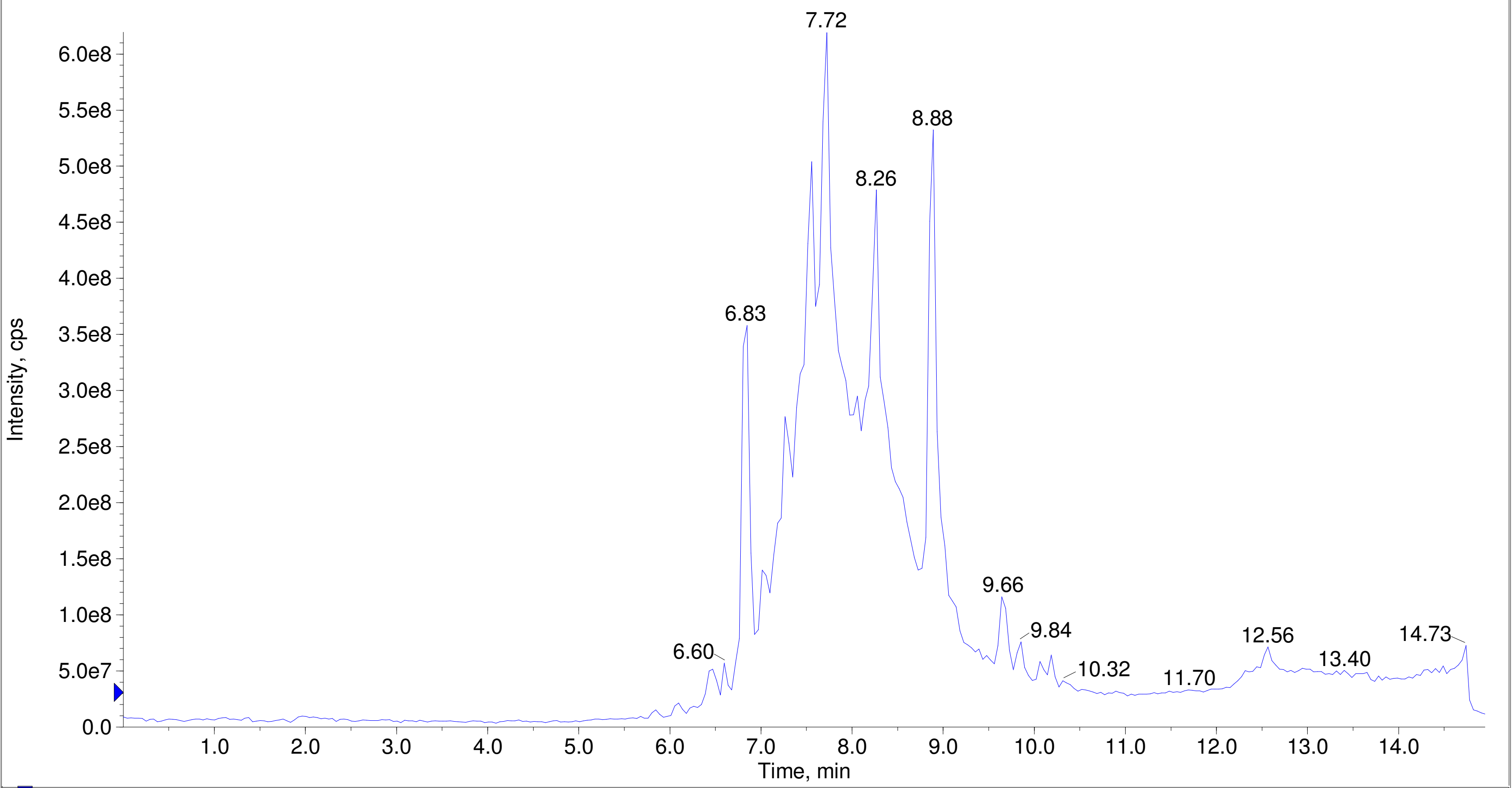


A/D Converter Channel 1 from Sample 5 (F7-35-P7-IL_OSF-num20-B8) of Data20210827.wiff Max. 10.0 %.

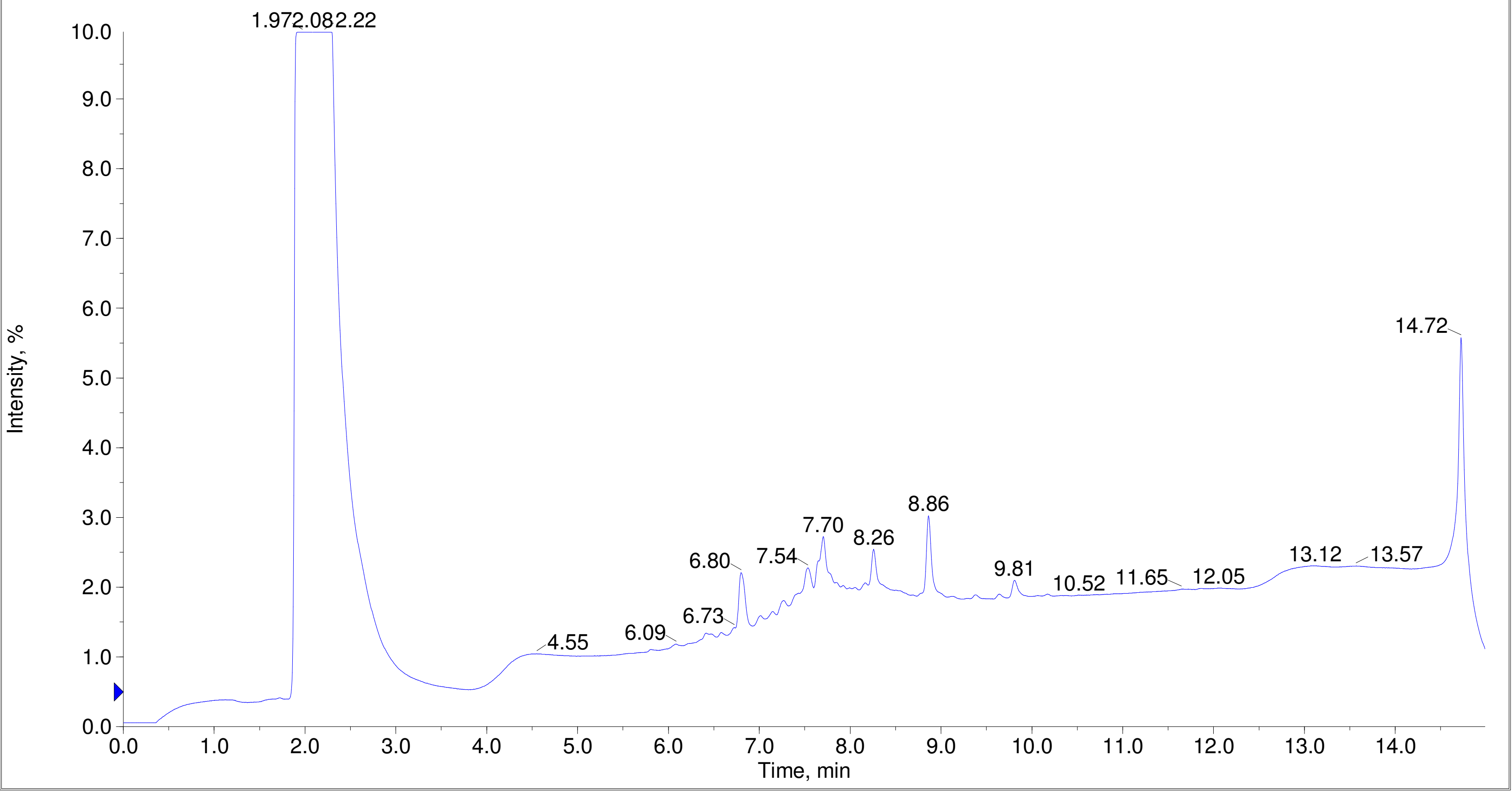

Printing Time: 3:21:35 PM
Printing Date: Monday, January 30, 2023
Service Engineer Chris Maier

Post PM
*Stanford U

Polarity/Scan Type: Positive Q1 MS
Sample Name: F7-35-P7-IL_OSF-num21-B9
*API 3000 SN D7640012

TIC of +Q1: from Sample 6 (F7-35-P7-IL_OSF-num21-B9) of Data20210827.wiff (Turbo Spray) Max. 5.2e8 cps.

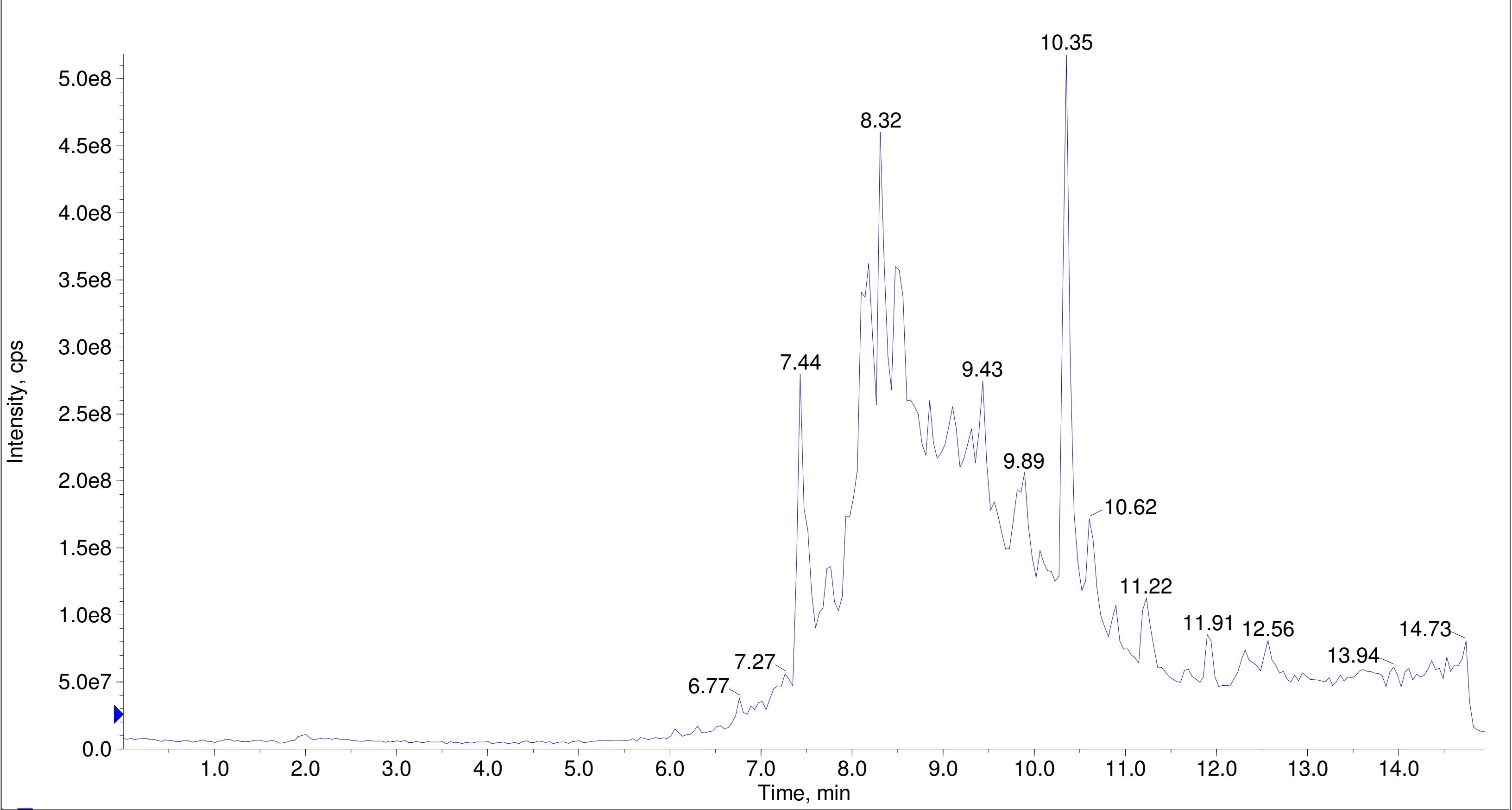


A/D Converter Channel 1 from Sample 6 (F7-35-P7-IL_OSF-num21-B9) of Data20210827.wiff Max. 10.0 %.

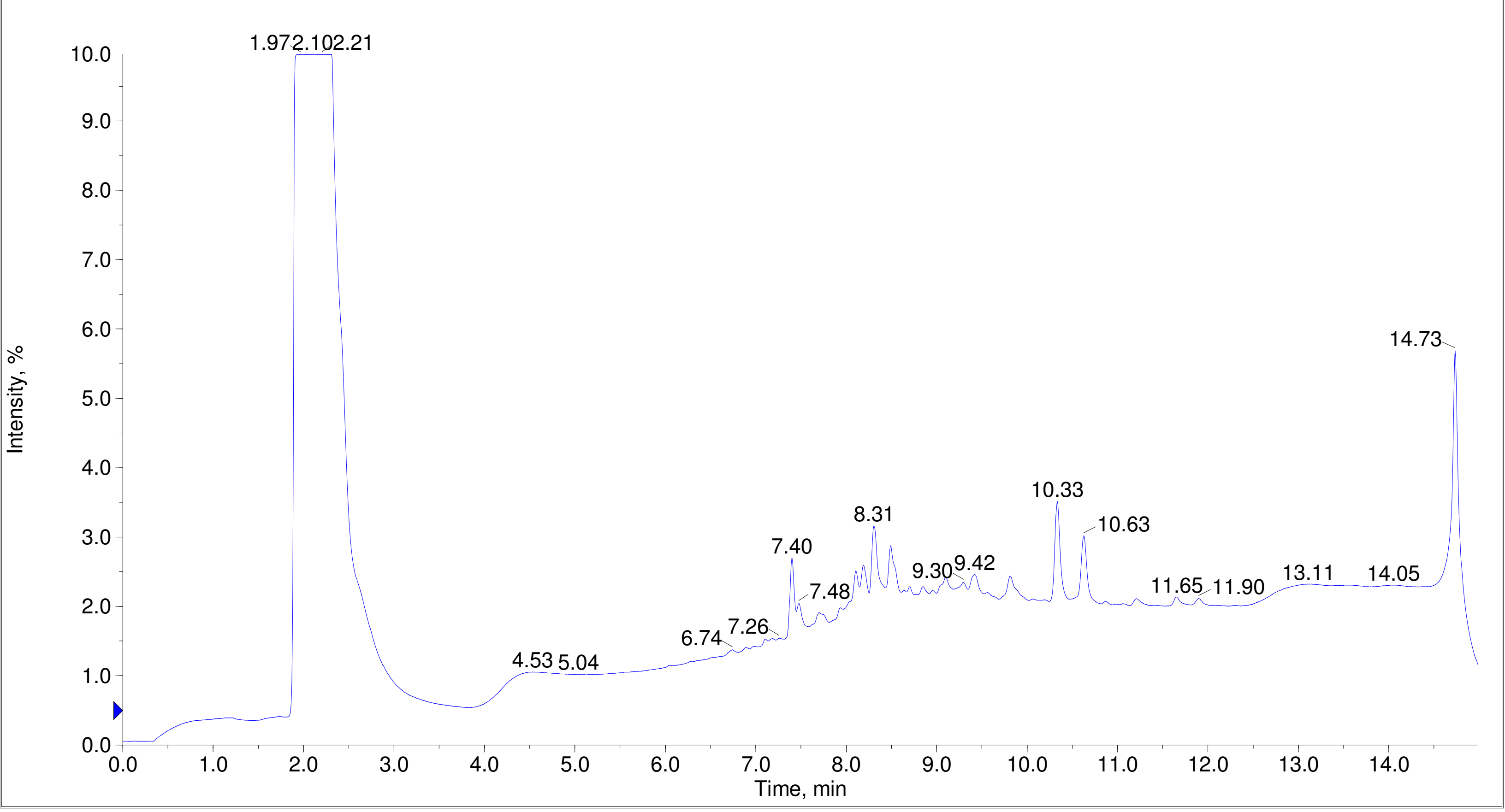

Printing Time: 4:22:51 PM Post PM Polarity/Scan Type: Positive Q1 MS
Printing Date: Tuesday, January 31, 2023 Sample Name: F7-35-P7-IL_OSF-num23-B11
Service Engineer Chris Maier *Stanford U *API 3000 SN D7640012

TIC of +Q1: from Sample 8 (F7-35-P7-IL_OSF-num23-B11) of Data20210827.wiff (Turbo Spray) Max. 6.5e8 cps.

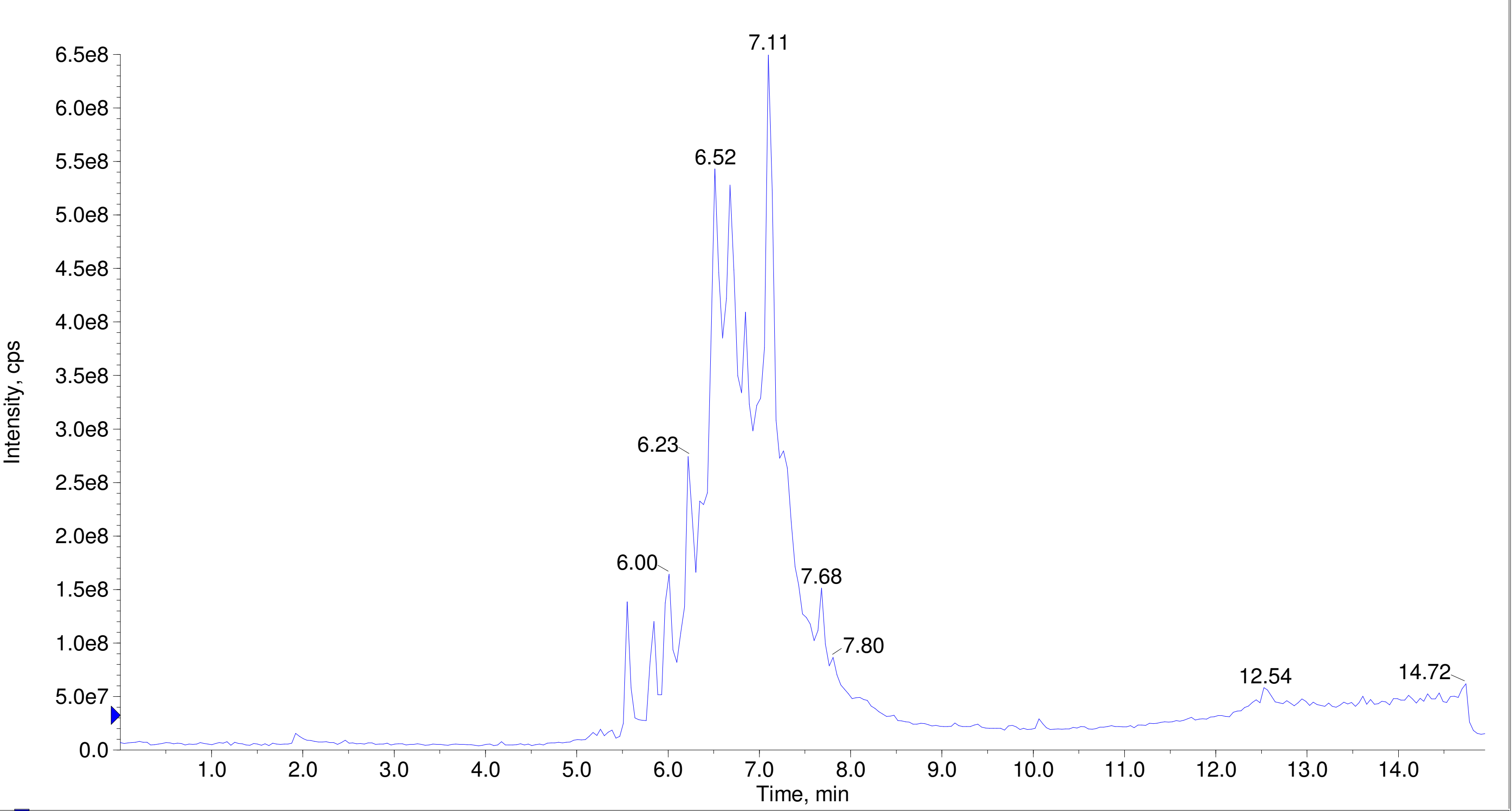


A/D Converter Channel 1 from Sample 8 (F7-35-P7-IL_OSF-num23-B11) of Data20210827.wiff Max. 10.0 %.

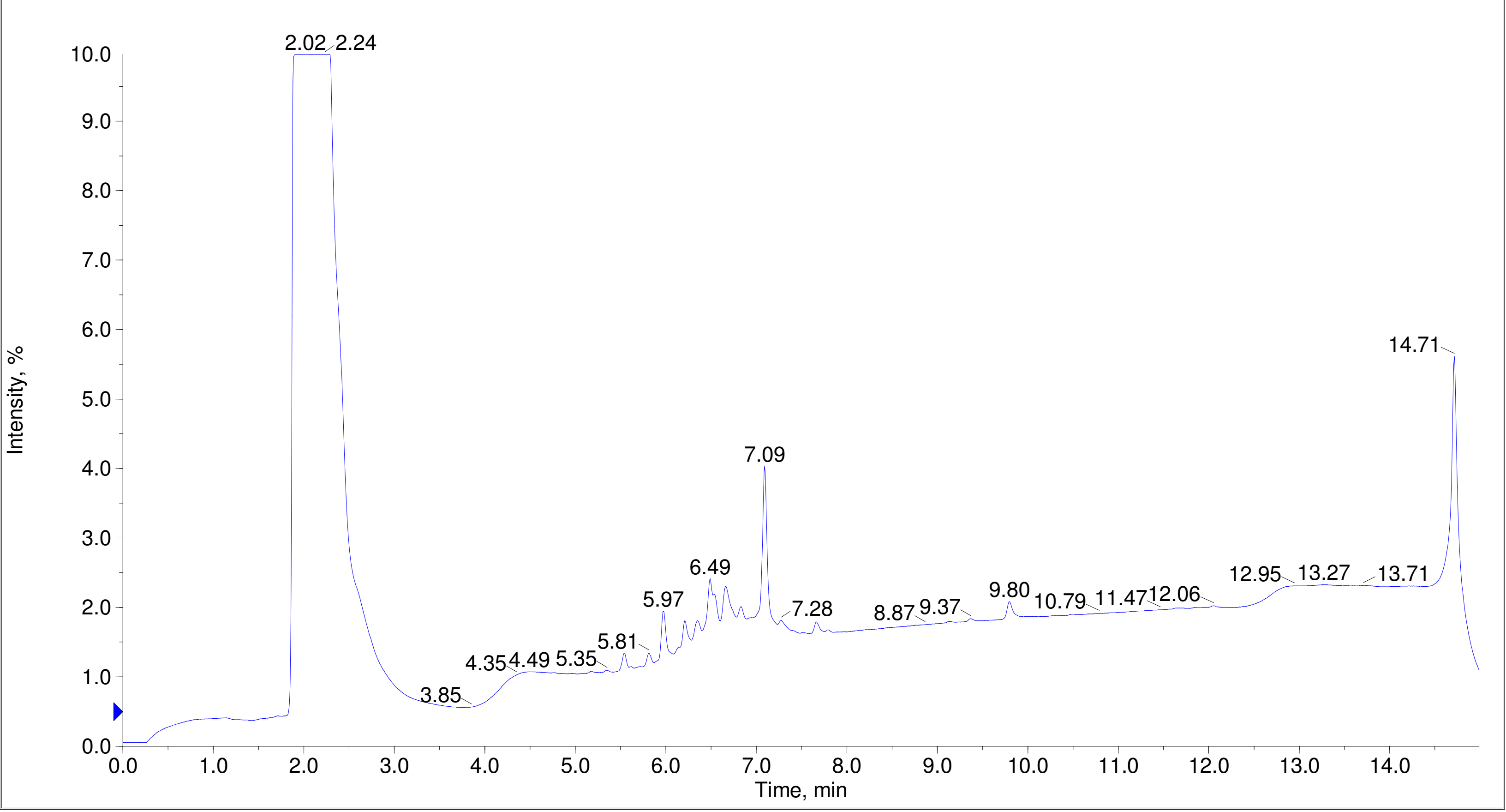

Printing Time: 4:24:19 PM
Printing Date: Tuesday, January 31, 2023
Service Engineer Chris Maier

Post PM
*Stanford U

Polarity/Scan Type: Positive Q1 MS
Sample Name: F7-35-P7-IL_OSF-num27-C3
*API 3000 SN D7640012

TIC of +Q1: from Sample 9 (F7-35-P7-IL_OSF-num27-C3) of Data20210827.wiff (Turbo Spray)

Max. 1.1e9 cps.

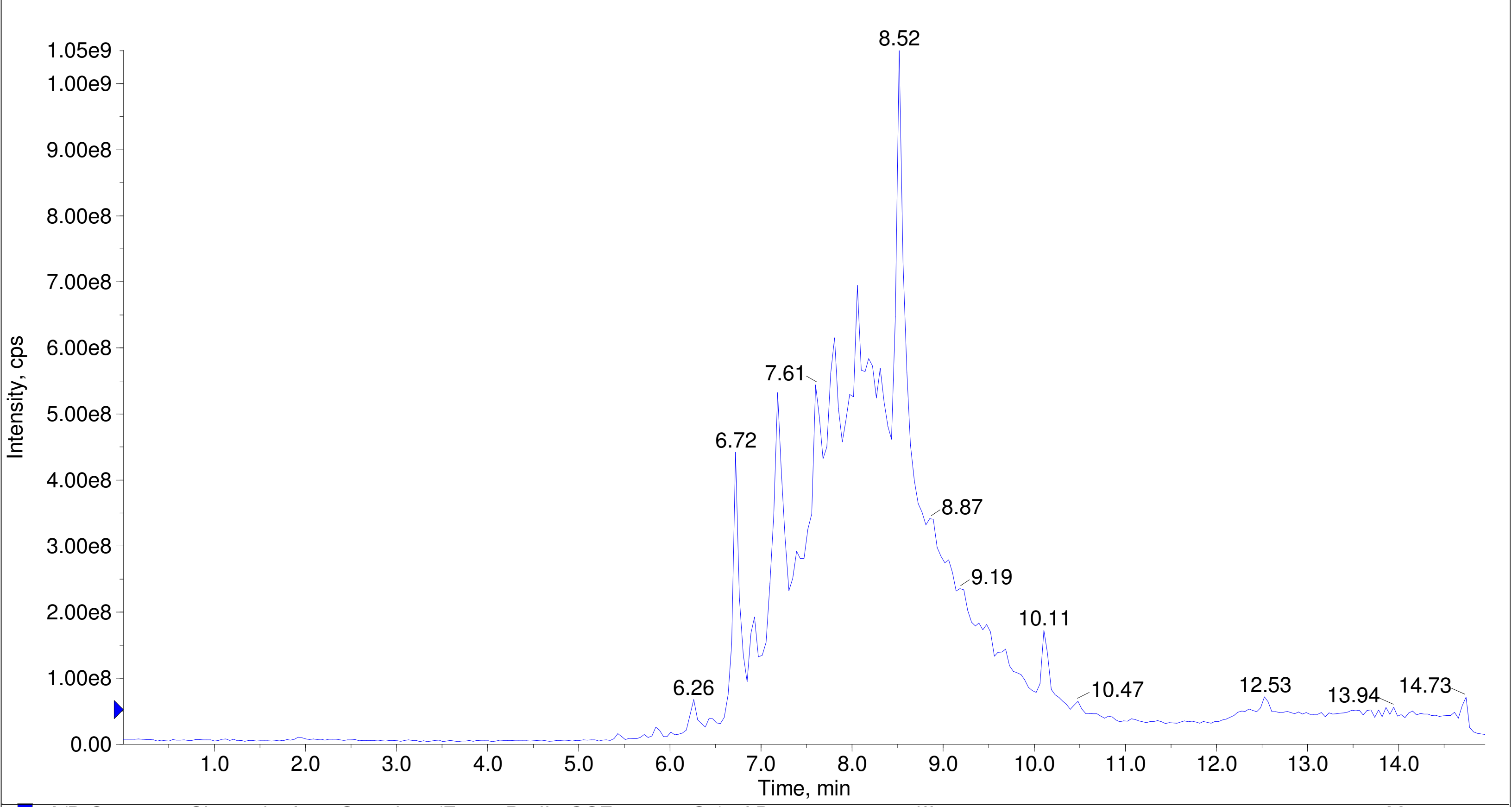


A/D Converter Channel 1 from Sample 9 (F7-35-P7-IL_OSF-num27-C3) of Data20210827.wiff

Max. 10.0 %.

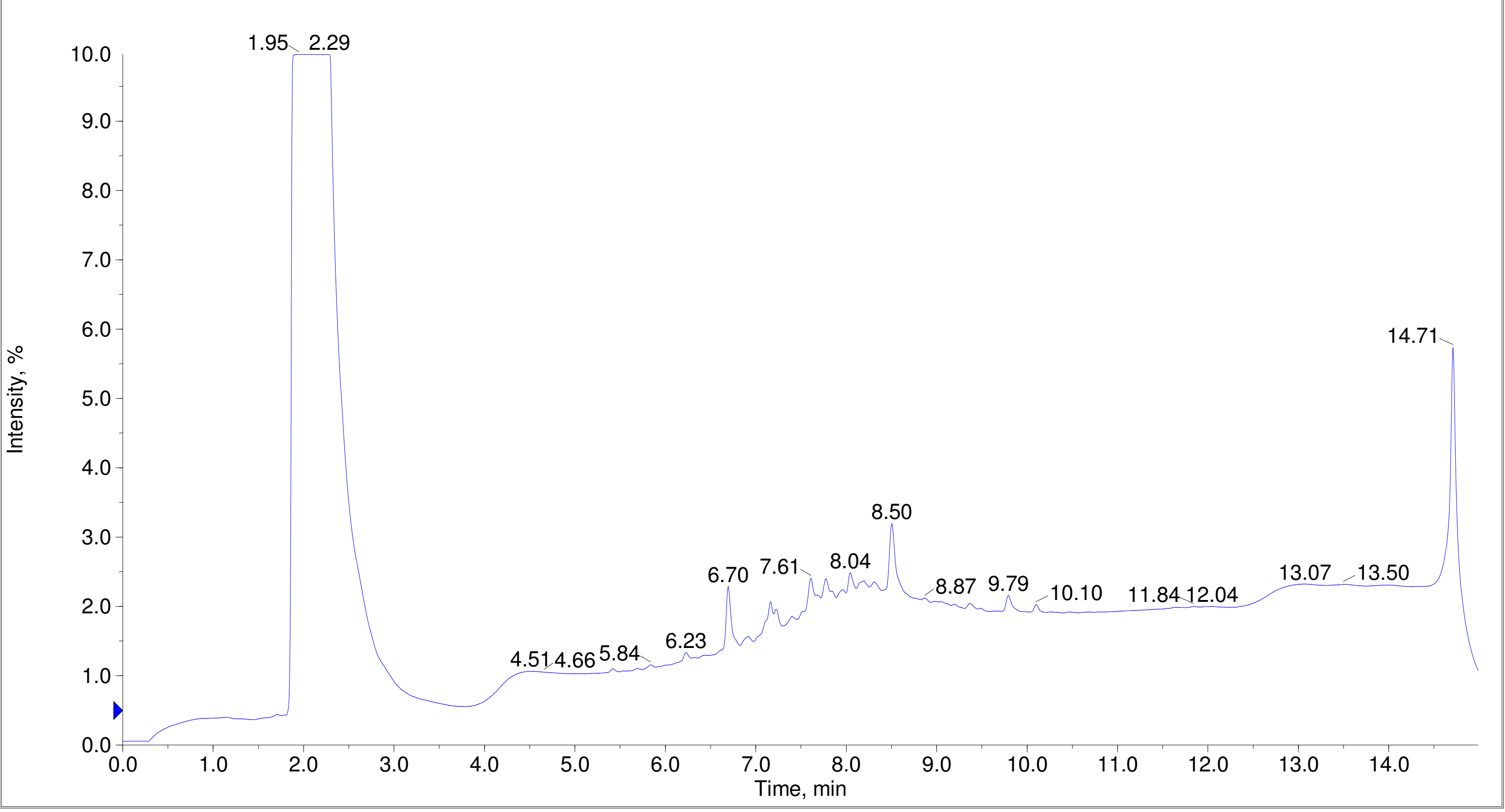

Printing Time: 4:25:30 PM
Printing Date: Tuesday, January 31, 2023
Service Engineer Chris Maier

Post PM
*Stanford U

Polarity/Scan Type: Positive Q1 MS
Sample Name: F7-35-P7-IL_OSF-num29-C5
*API 3000 SN D7640012

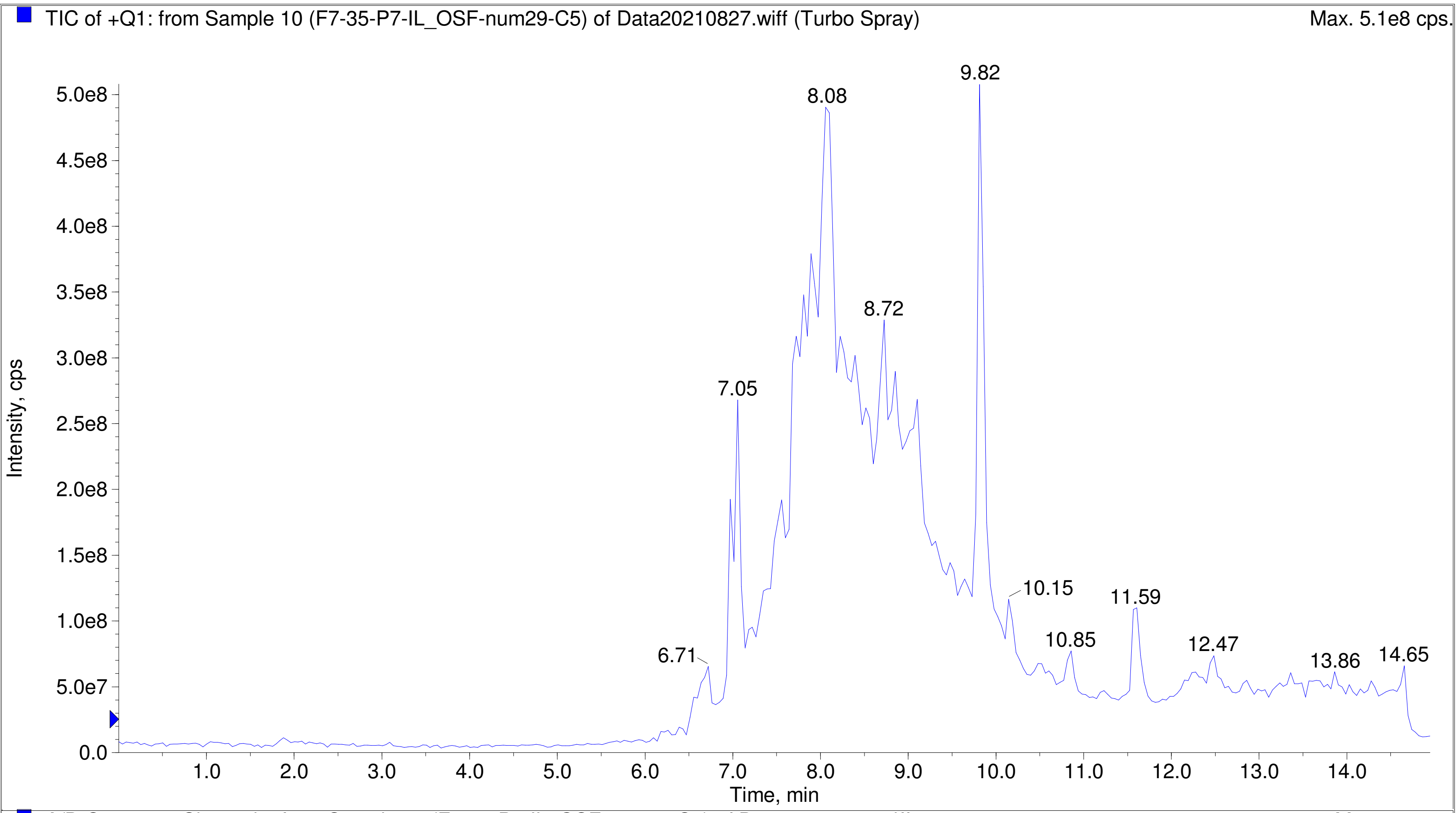


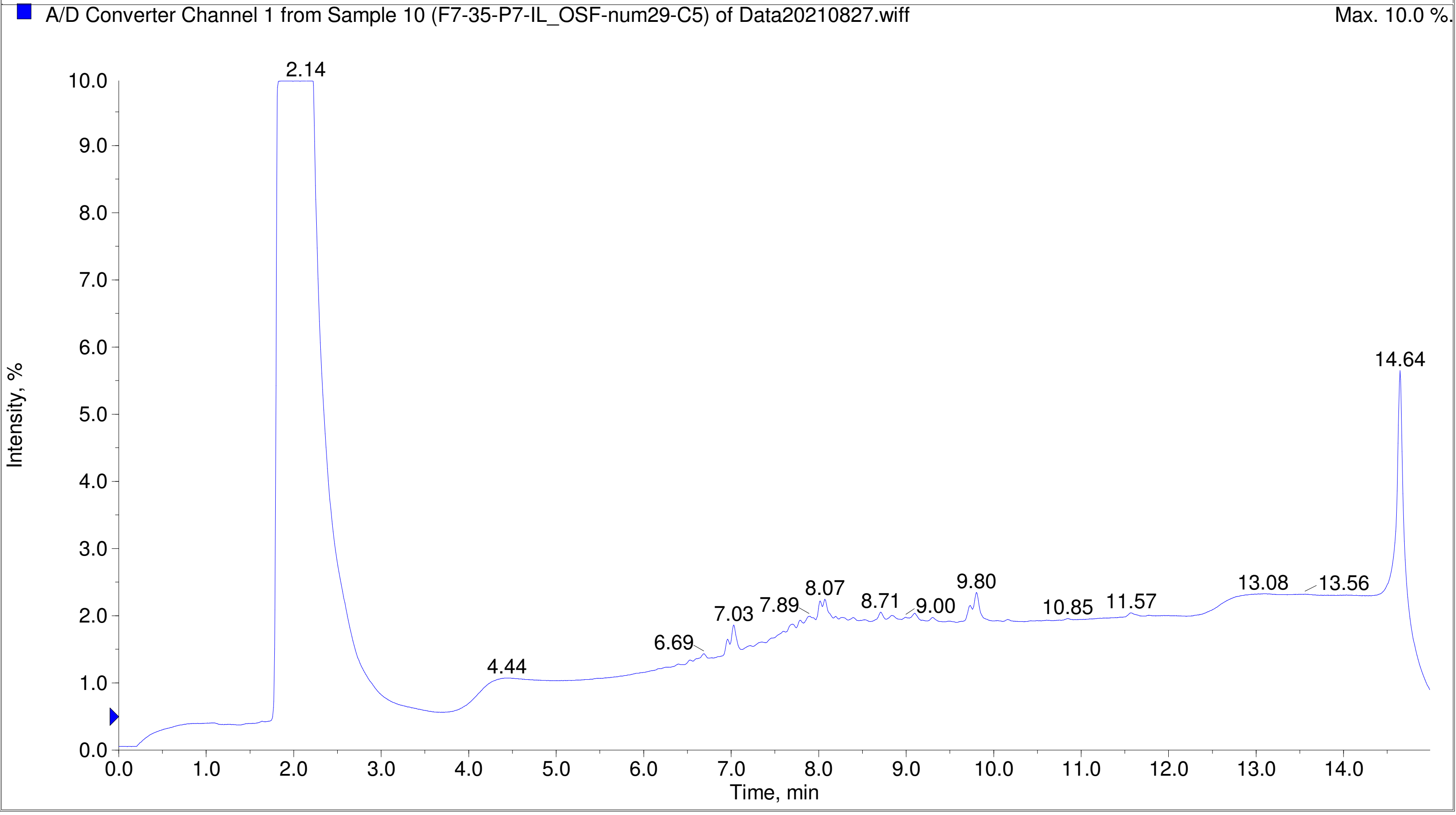

Printing Time: 4:26:42 PM Post PM Polarity/Scan Type: Positive Q1 MS
Printing Date: Tuesday, January 31, 2023 Sample Name: F7-35-P7-IL_OSF-num40-D4
Service Engineer Chris Maier *Stanford U *API 3000 SN D7640012

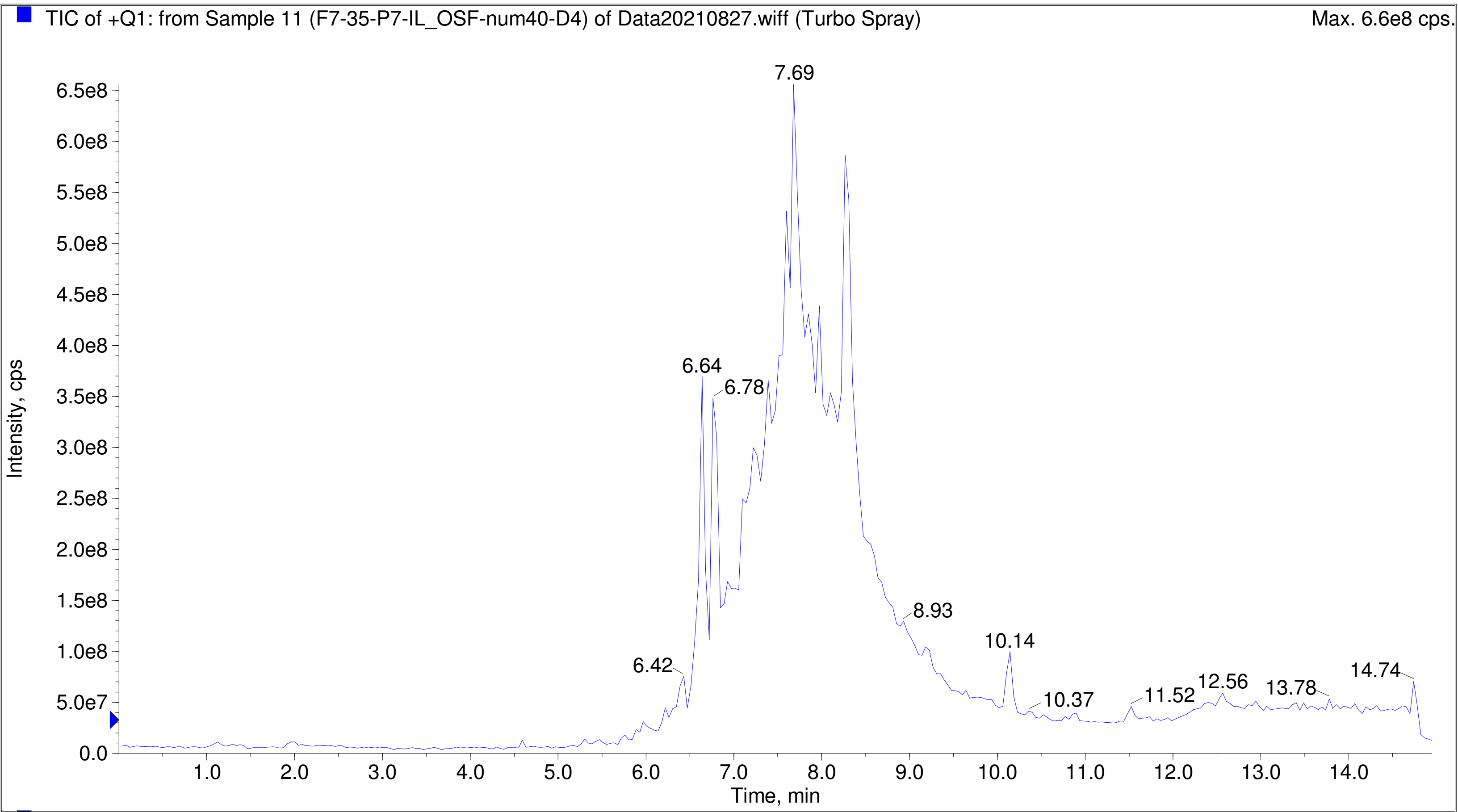


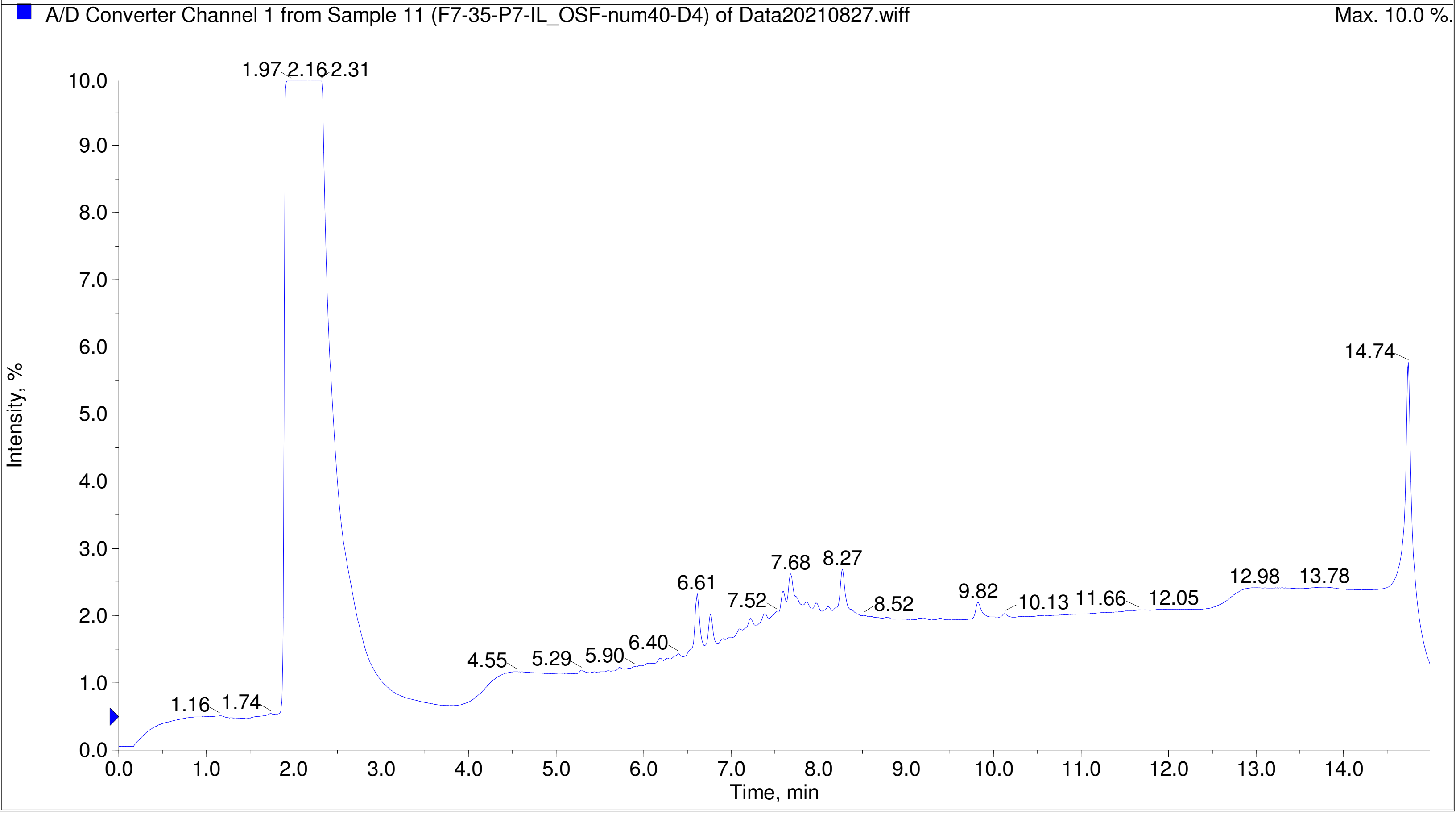

Printing Time: 4:27:29 PM
Printing Date: Tuesday, January 31, 2023
Service Engineer Chris Maier

Post PM

*Stanford U

Polarity/Scan Type: Positive Q1 MS
Sample Name: F7-35-P7-IL_OSF-num41-D5
*API 3000 SN D7640012

TIC of +Q1: from Sample 12 (F7-35-P7-IL_OSF-num41-D5) of Data20210827.wiff (Turbo Spray) Max. 7.6e8 cps.

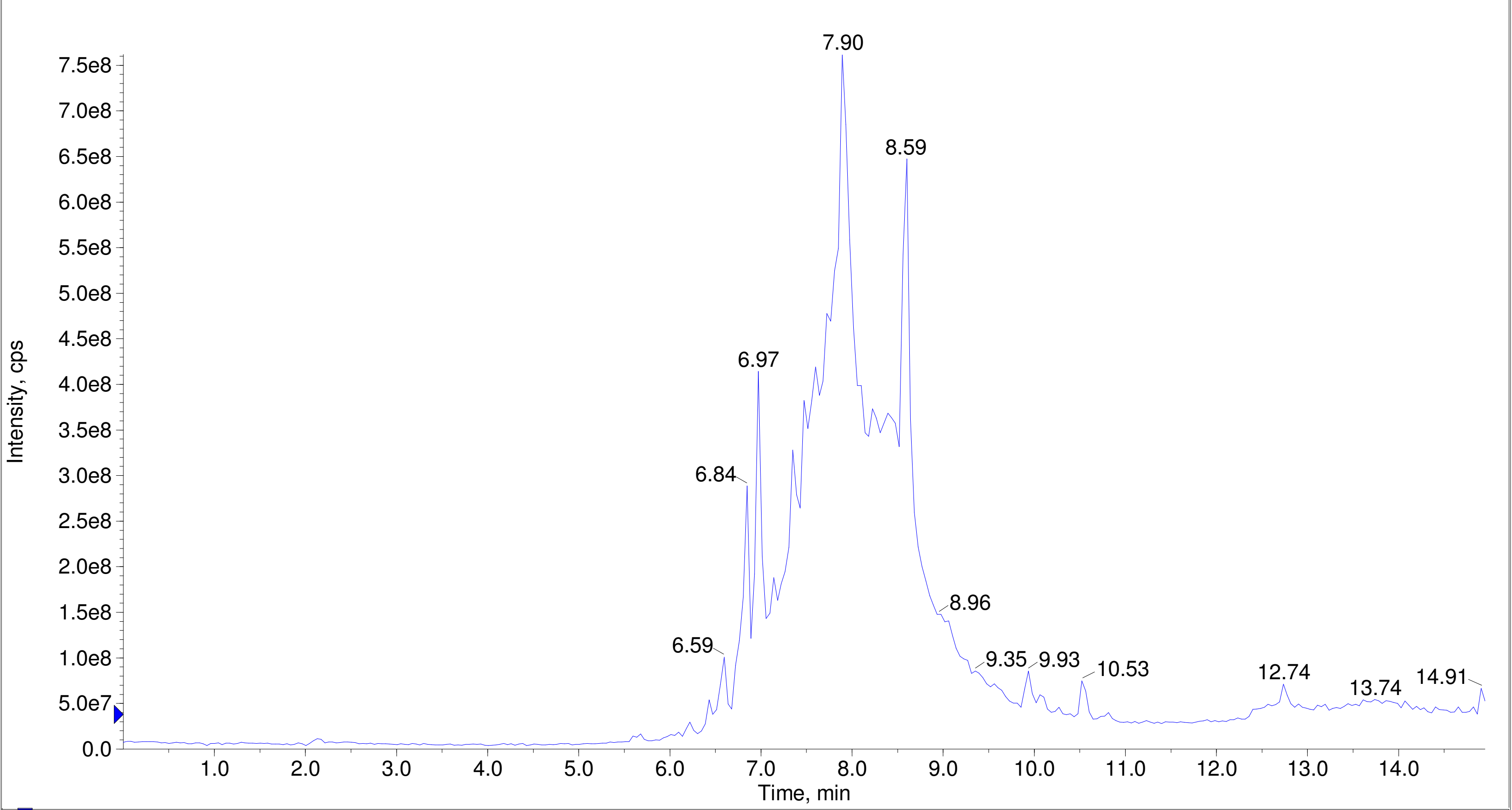


A/D Converter Channel 1 from Sample 12 (F7-35-P7-IL_OSF-num41-D5) of Data20210827.wiff Max. 10.0 %.

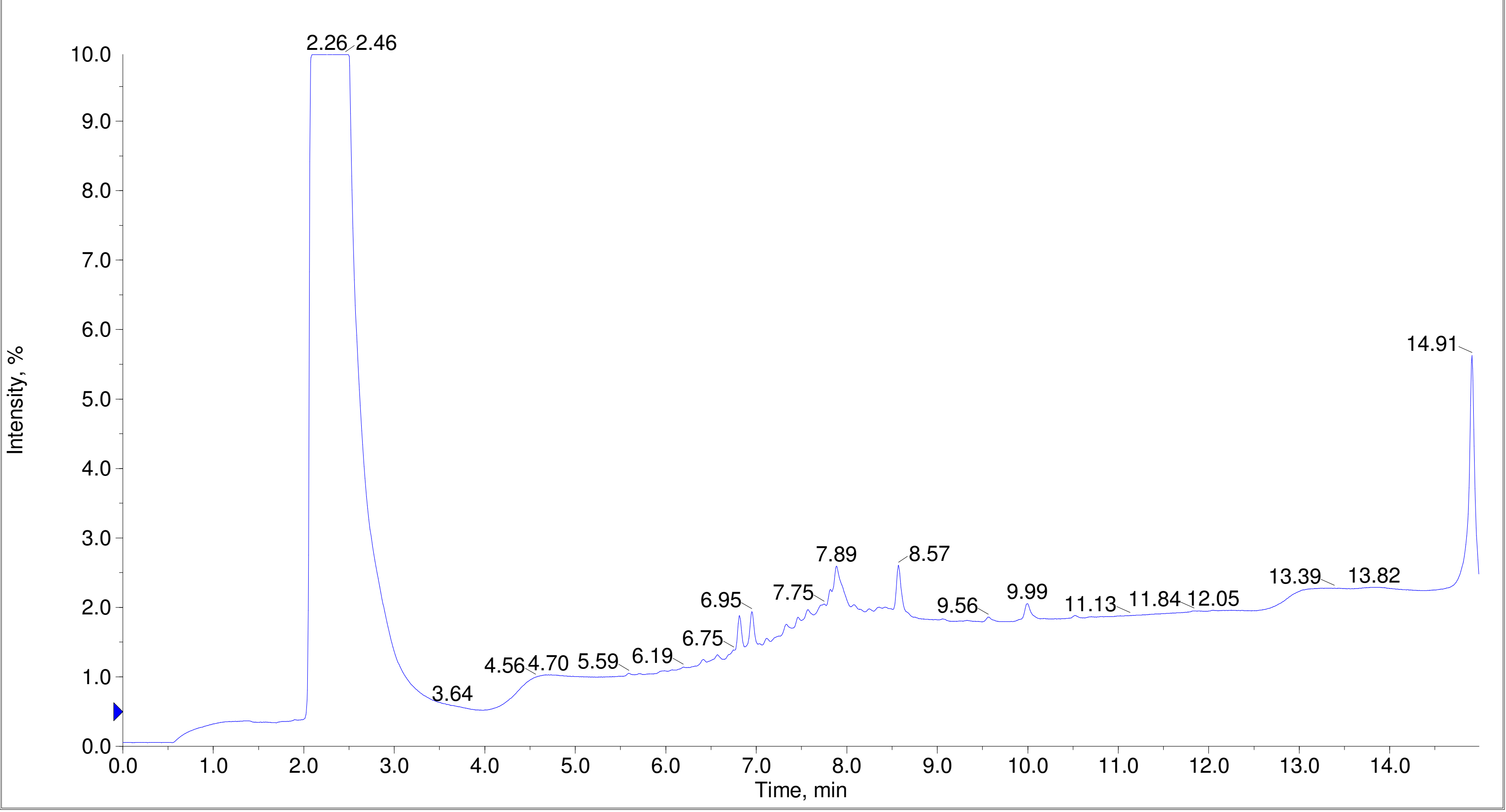
